# A high content microscopy screening identifies new genes involved in cell width control in *Bacillus subtilis*

**DOI:** 10.1101/2021.06.10.444761

**Authors:** Dimitri Juillot, Charlène Cornilleau, Nathalie Deboosere, Cyrille Billaudeau, Parfait Evouna-Mengue, Véronique Lejard, Priscille Brodin, Rut Carballido-López, Arnaud Chastanet

## Abstract

How cells control their shape and size is a fundamental question of biology. In most bacteria, cell shape is imposed by the peptidoglycan (PG) polymeric meshwork that surrounds the cell. Thus, bacterial cell morphogenesis results from the coordinated action of the proteins assembling and degrading the PG shell. Remarkably, during steady-state growth, most bacteria maintain a defined shape along generations, suggesting that error-proof mechanisms tightly control the process. In the rod-shaped model for Gram-positive bacteria *Bacillus subtilis*, the average cell length varies as a function of the growth rate but the cell diameter remains constant throughout the cell cycle and across growth conditions. Here, in an attempt to shed light on the cellular circuits controlling bacterial cell width, we developed a screen to identify genetic determinants of cell width in *B. subtilis*. Using high-content screening (HCS) fluorescence microscopy and semi-automated measurement of single-cell dimensions, we screened a library of ~ 4000 single knockout mutants. We identified 13 mutations significantly altering cell diameter, in genes that belong to several functional groups. In particular, our results indicate that metabolism plays a major role in cell width control in *B. subtilis*.

## Introduction

The bacterial landscape displays a rich variety of cell shapes, which are usually highly conserved at the single bacterial species level (1). The rationale behind a specific shape and its selective value remains speculative in most cases (1), as well as the molecular mechanisms that enable a specific shape to be determined and maintained across generations.

The shape of most bacterial cells directly depends on the shape of their cell wall (CW). The CW is primarily composed of a peptidoglycan (PG) scaffold that forms a rigid shell responsible for the mechanical properties of the cell envelope. In Gram-positive (G(+)) bacteria, the CW additionally contains PG-linked glycopolymers, the most abundant being the teichoic acids (TAs) (2). The PG sacculus is a contiguous matrix of linear sugar strands cross-linked by peptide bridges (3). Rod-shaped bacteria like *Bacillus subtilis* and *Escherichia coli*, the models for G(+) and Gram-negative (G(−)) bacteria respectively, use two different PG-synthesizing machineries: the divisome and the elongasome (4, 5). The divisome is required to build the septum at the sites of division, which upon cell separation will become the new polar caps of the resulting daughter cells. The elongasome synthesizes the cylindrical sidewall during cell elongation. The latter comprises two machineries working semi-independently: one involving aPBPs (class A penicillin binding proteins), bifunctional enzymes with transpeptidase (TP) and transglycosylase (TG) activity, and one named the “Rod complex”, which contains the SEDS-family RodA TG acting in concert with bPBPs (class B PBPs) carrying mono-functional TP activity such as PBP2A and PbpH in *B. subtilis* (5, 6). The prevailing model postulates that the Rod complex processively and directionally inserts glycan strands around the cell circumference, building the bulk of the PG meshwork, while aPBPs perform limited and localized, unoriented strands insertion (6–8). In agreement with this model, in *B. subtilis* aPBPs are dispensable (9–11) while most PG synthases of the Rod complex are essential, like RodA (6, 9, 12), or co-essential like PBP2A and PbpH (13). This essentiality reflects that a failure in the proper establishment of the PG mesh compromises cellular integrity.

In addition to TP and TG enzymes, the Rod complex also includes the essential MreC and MreD morphogenetic proteins, which are presumed regulators of the activity of the complex (4, 14), and actin-like MreB proteins, which are believed to orient the circumferential motion of the complex (5, 15). The *B. subtilis* genome encodes three MreB paralogs, the essential MreB and Mbl, and MreBH, which becomes essential in the absence of the other two paralogs, in the absence of aPBPs, under stress conditions and at low Mg^2+^ concentrations (11, 16, 17). RodZ, a protein of unknown function, is also a component of the Rod complex shown to be critical for rod shape maintenance in the G(−) bacteria *Caulobacter crescentus* and *E. coli*, and essential only in *C. crescentus* (18–21). The involvement of RodZ in shape control and its essentiality are less clearly established in G(+) bacteria. Described as essential in *B. subtilis* in an early report (22), several *rodZ* insertional or deletion mutants have been reported since, displaying minimal shape defects (23–25).

It has long been known that rod-shaped bacteria vary their size depending on the growth conditions and in particular on nutrient availability (26, 27). Rapidly growing cells have a bigger volume that slowly growing cells, a relationship often referred to as the (nutrient) “growth law” (for a review on this topic, see (28) or the very detailed (29)). However, while in *E. coli* cell width varies greatly (up to 100%) and concomitantly with cell length (26, 30–32), *B. subtilis* cells adjust their length but maintain a virtually constant diameter regardless of the growth conditions (32–36). This remarkable consistency suggests that cell width is a physiological parameter somehow encrypted in the genome of *B. subtilis*, and that it must be carefully monitored during growth to correct for potential deviations to its nominal value. Yet how rod-shaped bacteria check and balance their diameter remains unclear. Recently, Garner and co-workers showed that the cell diameter results from the balance between the opposite activities of the Rod and aPBPs elongation machineries (7). They proposed a model in which aPBPs-mediated isotropic insertion of unoriented strands into the PG meshwork enlarges the cell cylinder while Rod complex-mediated organized circumferential insertion of PG strands reduces it (7).

According to this model, the observation of thinner *B. subtilis* cells in the absence of aPBPs (37–39), can be explained as the result of the imbalance of the aPBP/Rod complex activities (7). Albeit thinner, cells that rely on the Rod complex for growth retain nevertheless their rod shape, indicating that the ‘check & balance’ process of cell width control is still in place. Conversely, reduced activity of the Rod complex leads to the opposite imbalance, driving to an increased cell diameter (7). In absence of the essential (or co-essential) component(s) of the Rod complex, this ultimately leads to spherical cells, as exemplified by the depletion of RodA, MreC, MreD, PBP2A/PbpH or MreB/Mbl/MreBH (13, 40–43). In agreement with this model, most genes reported to affect cell width in *B. subtilis* are directly involved in CW homeostasis, affecting one of the competing PG-synthesizing machineries, PG hydrolysis (required to allow PG expansion) or TAs synthesis (Table S1). Other genes previously reported to affect width encode proteins whose absence perturbs the production or the localization of the latter (Table S1).

Here, we aimed at identifying at the genome scale level additional determinants of cell width control during rapid exponential growth. We screened a complete organized collection of *B. subtilis* deletion mutants (24) using High-Content Screening microscopes (HCSm). Our protocol for mid-throughput analysis allowed us to uncover several new genes that may work to maintain cell diameter. These are involved in several cellular processes including CW synthesis, cell division, metabolism, and translation, suggesting that cell width homeostasis results from the combined action of several cellular circuits. Among these, our analysis suggests that metabolism and CW homeostasis are the two main routes affecting cell width.

## Results

### *Bacillus subtilis* cells display limited width variability during rapid exponential growth

It has long been accepted that, in contrast to *E. coli*, the cell diameter of *B. subtilis* cells remains virtually constant regardless of the growth rate (32, 33). We wondered how variable the cell diameter could be in isogenic *B. subtilis* populations during fast exponential growth. We used MicrobeJ, a Fiji plugin (44–46), to perform cell segmentation and quantify cell diameter (see ‘Material and methods’, Table S2, Sup. Information). We first compared six independently acquired datasets of wild type *B. subtilis* cells grown to exponential phase in rich LB medium. We observed that the average width remained remarkably constant between experiments (variability below 2 %; Fig. 1A and Table S3). Also, the cell-to-cell variability (standard deviation) of the measured width in each population remained low, ranging from 0.071 to 0.089 µm across the different replicates (Table S3). These variations might reflect true differences of cell diameter or just the error of our measurements, but in either case variability was low. This reproducibility allowed us to expect the detection of potentially small variations of cell diameter between mutant strains. We next compared the diameter of *B. subtilis* cells exponentially growing in two different media, rich (LB) and poor (S), and thus supporting different growth rates. In agreement with previous reports (32, 33, 36), we found no significant difference of width between cells grown in rich and poor media (Fig. 1B). The cell-to-cell variability was similar in the two media, indicating that this variability is independent of the growth rate. Taken together, these experiments indicated that *B. subtilis* exerts a tight control over its diameter, whose variability remains below 2 % in average across conditions and replicates.

**Figure 1.**
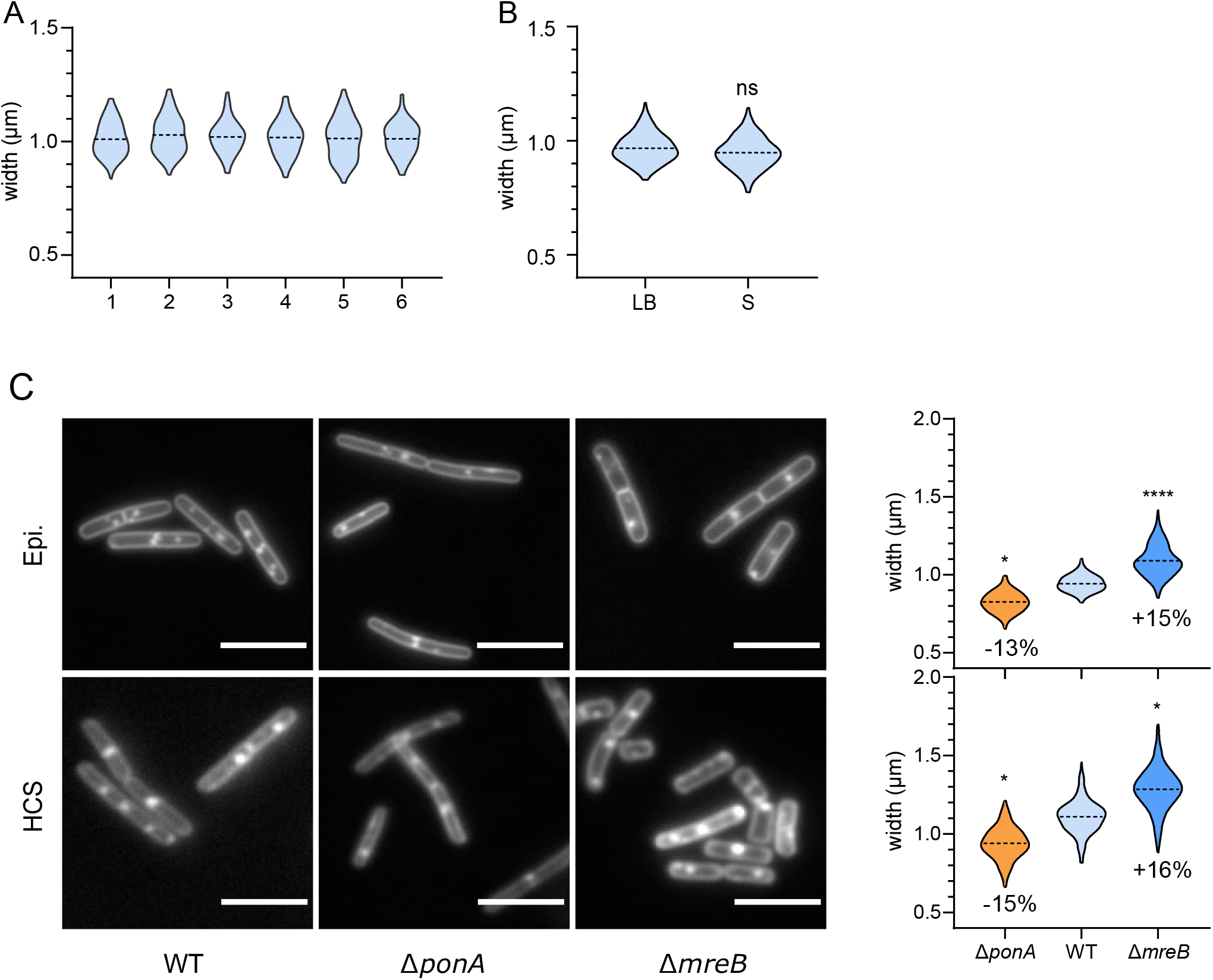
Discrimination of diameter-control deficient *B. subtilis* mutants: a proof of concept. **A**. Comparison of cell width distribution of six independent cultures of fixed wild type *B. subtilis* cells grown in rich (LB) medium, observed on an epifluorescent microscope. One-way ANOVA statistical analysis showed no significant differences between the replicates (Table S3). **B**. Comparative cell width distributions of fixed wild type *B. subtilis* cells grown in rich (LB) and minimal (S) media (epifluorescent microscope). **C**. Qualitative (images) and quantitative (distribution of measured cell width) comparisons of data acquired on a wide-field epifluorescence microscope and a confocal HCSm, using the wild type, Δ*mreB* and Δ*ponA* mutant strains of *B. subtilis*. Fluorescent images were acquired on cells grown to mid exponential phase (0.2<OD600 nm<0.3), fixed and stained with the FM1-43fx membrane dye. Discrete fluorescent foci result from cell fixation. Scale bar, 5 µm. Width distributions are displayed as violin plots with the broken line indicating the mean. Statistical analyses were performed as described in the method section ‘statistical analysis’. When significant, the difference between the means, expressed as a percent, is indicated on the plots. B and C are compilations of at least two independent experiments.

### HCS microscopy allows screening for small phenotypic variations of cell width

We next defined conditions that would minimize false positives in a microscopy-based screen of a genome-scale deletion library of *B. subtilis*. First, cells were fixed to obtain snapshots of their dimensions during exponential growth. Fixation induces a slight reduction of cell width relative to live cells (Fig. S1A), but prevents issues resulting from the time required for the preparation and imaging of 96-well plates with HCSm. Second, the growth medium was supplemented with 20 mM of MgSO_4_ to prevent potential inaccurate estimation of the cell diameter of mutants displaying irregular shapes or lysing. In *B. subtilis*, millimolar concentrations of magnesium in the growth medium are known to reduce the activity of PG hydrolases (47) and to alleviate the morphological defects of mutants affected in PG synthesis (48–50), allowing propagation of otherwise lethal mutations. Importantly, in the presence of high magnesium these mutants display a normal rod shape but still present an abnormal width (48, 49). Addition of Mg^2+^ to the growth medium slightly reduced the average width of wild type cells (Fig. S1B), as previously reported (37). Our ability to detect these slight width differences when cells were either fixed (Fig. S1A) or grown in high magnesium (Fig. S1B) confirmed the sensitivity of our assay to detect small variations of average width between populations.

To further demonstrate the sensitivity of our assay, we tested the *mreB* and *ponA* null mutants, known to be wider and thinner, respectively, than wild type cells (38, 51). As shown in Figure 1C, the altered width of Δ*mreB* and Δ*ponA* mutants was unambiguously detected when cells were grown in high Mg^2+^, fixed and observed in either our conventional epifluorescence microscope or the HCSm. The cell-to-cell variability and the average cell widths noticeably increased when measurements were performed on HCSm-acquired images but the relative difference of width between the two mutant strains and the wild type were perfectly conserved (Fig. 1C). These control experiments showed that mutants affected for the control of width could be identified in our medium-throughput HCSm approach.

Next, we screened the complete *B. subtilis* kanamycin-marked ordered deletion library (BKK) (24), which contains 3 983 single-gene deletion mutants (~93 % open reading frames coverage) of the parental 168 strain (GenBank Al009126) (Fig. 2, see ‘Materials and methods’ for details). In order to prevent plate-to-plate fluctuations and to compare the widths of the mutants across plates, the width of each mutant was expressed relative to the <u>a</u>verage cell <u>d</u>iameter per <u>p</u>late (ADP, see ‘Materials and methods’). The average of the ADPs of the 48 plates (Fig. S2A) and the average cell width of the wild type strain grown and imaged in the same conditions showed no significant difference (Fig. S2B). For each single mutant, we calculated the delta between its average width and the ADP of its plate (Table S4). The 3 983 Δwidth obtained display a Gaussian distribution, spreading from −13.9 to +23.4 % but with 90 % of the values comprised in a narrow +/−5 % variation to the mean (Fig. 3A). Next, we arbitrarily set up a cutoff of the 1 % most affected strains (0.5 % largest and 0.5 % thinnest) (Fig. 3A). The 40 mutants selected displayed a difference in diameter ranging from 8.9 to 23.4 % to that of their ADP (Table S5, ‘screening step’). Using low-throughput epifluorescence microscopy imaging and the wild type *B. subtilis* strain as a reference, we checked the cell width phenotype of the selected mutants (Fig. 2, see also ‘Materials and methods’), while the deletion in each mutant was verified by PCR. Two of the strains in the collection were wild type for the tested loci (Table S5) and were discarded for further analysis. A quarter of the mutants displayed a Δwidth ≤ 2%, i.e. equivalent to the variability between wild type replicates (Fig. 1A; Table S3), suggesting that our HCSm analysis yielded some false positives (Table S5, ‘checking step’). All 38 confirmed-knockout mutants were nevertheless back-crossed into the wild type background (Fig. 2, Table S6) before attempting further characterization, in order to exclude phenotypes unlinked to the candidate gene deletions.

**Figure 2.**
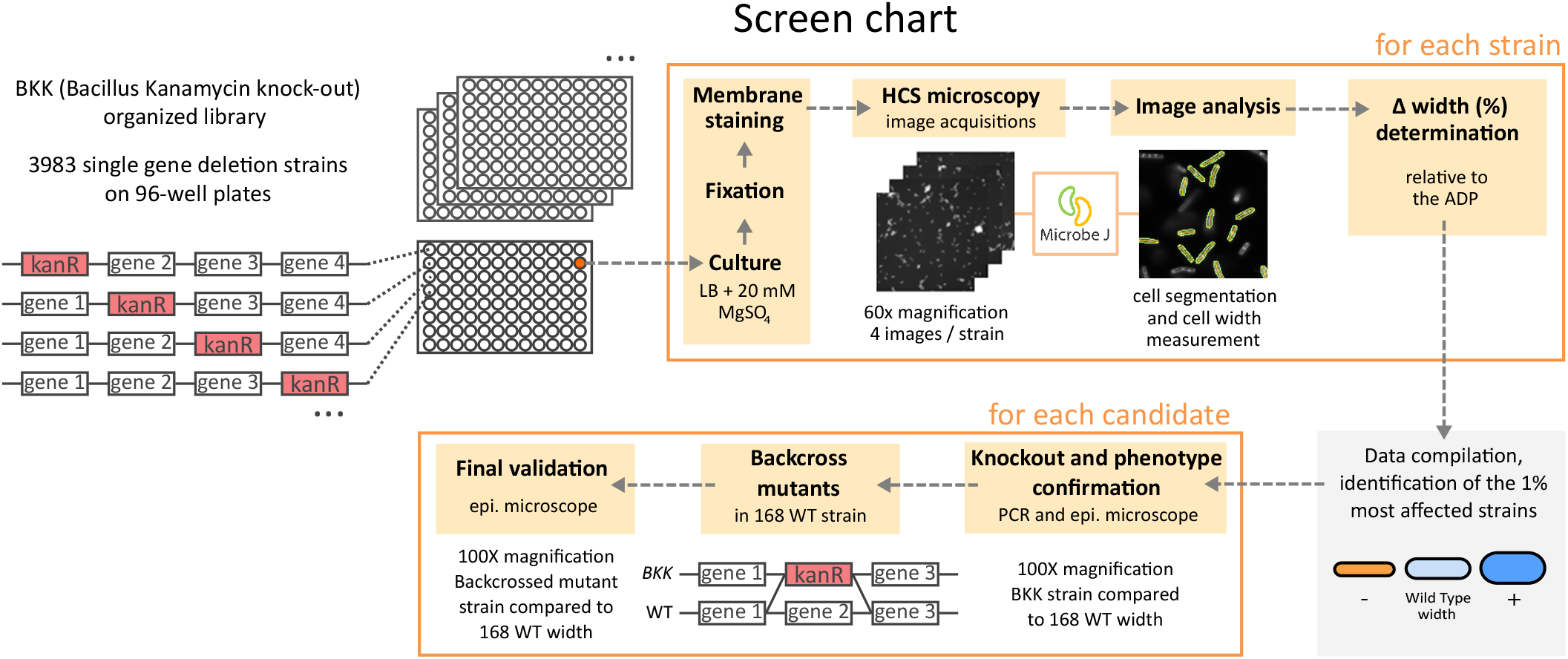
Protocol summary of the screening process. Screening of the *B. subtilis* BKK collection arrayed in 96-well plates was based on automated image acquisition using an HCSm. The MicrobeJ plugin of Fiji was used for cell segmentation and width measurements. Average cell diameter of each mutant was compared to the average diameter of all cells on the plate (ADP). Candidates where confirmed by measuring their diameter on images acquired with an epifluorescence microscope, relative to the wild type strain. The selected mutants were then backcrossed into a wild type background before final width determination over triplicate experiments.

**Figure 3.**
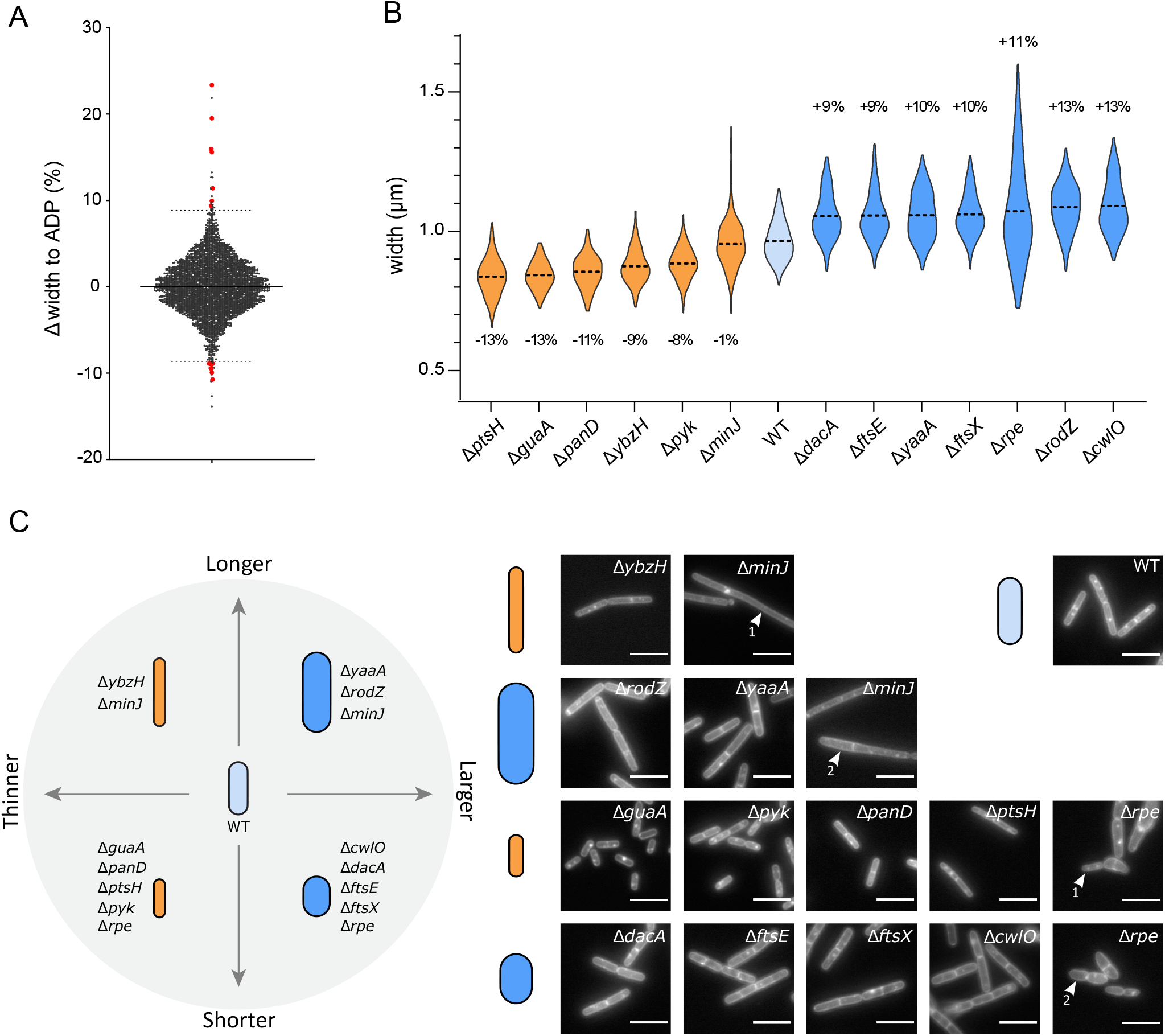
The screen reveals 13 mutants with a cell width variation > 8 % relative to the wild-type. **A**. Width difference (Δwidth) of each mutant relative to the ADP of its plate. Dotted lines indicate the cut-offs for the 0.5 % largest (top) and thinnest (bottom) mutants. Red dots mark the 13 mutants with confirmed diameter defects after deletion were backcrossed into the wild-type strain. **B**. Cell width distributions of the 13 selected (backcrossed) mutants. Orange and blue plots correspond to thinner and wider mutants respectively, compared to their parental wild type strain (light blue). Broken line: mean. Data are compilations of three independent experiments. The difference between the means of each mutants and the wild-type is indicated, in %. Statistical significances between the mutants and the wild type strain width were calculated using nested t-tests (see Table 1 for P-values). All differences were significant except for the mean width of the Δ*minJ* mutant. **C**. Phenotypes of the backcrossed mutants segregate into four classes based on their width and length defects. Δ*minJ* and Δ*rpe* mutants form both thinner (arrowhead, 1) and larger (arrowhead, 2) cells. Displayed are images of FM1-43fx membrane-labelled fixed cells. Scale bar, 5µm.

### Cell wall and central carbon metabolism genes are linked to cell width control

Next, we carefully measured the diameter of the backcrossed mutants (Table S5, ‘post-backcross step’). A large reduction of the Δwidth compared to that of the parental strain was confirmed for most of them. Choosing a stringent Δwidth cutoff of 8%, we selected 12 mutants significantly and reproducibly (over 3 independent experiments) wider (7) or thinner (5) (Fig. 3B, 3C, Table 1 and Table S5). We additionally kept the Δ*minJ* mutant despite its non-significant Δwidth (−1 %) because of the peculiar uneven width affecting some of the cells in this mutant (Fig. 3C). The width phenotypes were conserved for all 13 mutants when grown without magnesium supplementation (Table S5, ‘backcross strains without Mg^2+^’), confirming that in contrast to cell bulging, swelling and lysis (observed as consequences of CW synthesis impairment), magnesium cannot rescue the alteration of width. This suggests that cell width alteration does not result from uncontrolled PG hydrolytic activity.

**Table 1.** Cellular parameters of the confirmed width-control defficient strains

| Gene | Functional category<sup>4</sup> | $\delta$ length<sup>1</sup> (%) | P-value<sup>2</sup> | $\delta$ width<sup>1</sup> (%) | P-value<sup>2</sup> | $\delta$ GT<sup>3</sup> (min) | P-value<sup>2</sup> |
| --- | --- | --- | --- | --- | --- | --- | --- |
| *minJ* | cell division | 120.6 | **** | -1.0 | ns | 5,6 | * |
| *ybzH* | unknown (putative TR) | 12.9 | **** | -9.3 | **** | 1,7 | * |
| *rodZ* | CW homeostasis | 10.7 | ** | 12.5 | **** | -1,1 | ns |
| *yaaA* | translation | 4.4 | ** | 9.7 | *** | -0,7 | ns |
| *wt* | - | 0,0 | - | 0,0 | - | 0,0 | - |
| *cwlO* | CW homeostasis | -5.2 | ns | 13.1 | *** | 0,3 | * |
| *ftsX* | CW homeostasis | -8.6 | ** | 10.0 | **** | -0,7 | ns |
| *ftsE* | CW homeostasis | -10.9 | **** | 9.4 | **** | -0,7 | ns |
| *dacA* | CW homeostasis | -11.7 | **** | 9.3 | **** | -1,1 | ns |
| *ptsH* | central C metabolism | -12.2 | **** | -13.3 | **** | 17,6 | * |
| *panD* | central C metabolism | -12.5 | **** | -11.4 | **** | 11,9 | * |
| *rpe* | central C metabolism | -13.6 | *** | 11.0 | **** | 12,8 | * |
| *pyk* | central C metabolism | -21.4 | **** | -8.3 | *** | 36,5 | * |
| *guaA* | purine metabolism | -36.7 | **** | -12.7 | **** | 38,1 | * |

Colors of numbers indicate positively (purple) or negatively (orange) affected value in width or length, or increased or decreased generation time, compared to the wild type strain
1: difference (in %) relative to wild type, average of 3 independent pooled replicates
2: statistical significance estimated by nested t-test ( $\delta$ length, $\delta$ width) or Mann-Whitney test (GT)
P-values are displayed as follow: \*\*\*\* = $P < 0.0001$ ; \*\*\* = $0.0001 < P < 0.001$ ; \*\* = $0.001 < P < 0.01$ ; \* = $0.01 < P < 0.05$ ; ns = $P > 0.05$ .
3: difference of generation time (GT) relative to the wild type, averages of 4 independent experiments
4: according to Subtiwiki (Zhu, 2018)

Among the 13 selected mutants (Fig. 3B and C and Table 1), we identified 9 new genes affecting cell width of *B. subtilis* (*ptsH, guaA, panD, ybzH, pyk, yaaA, minJ, dacA, rpe*) and confirmed 4 others (*rodZ, cwlO, ftsE*, and *ftsX*) previously reported to be affected in cell diameter (Table S1; ‘this study’). Note that despite not being in the top 1 % of genes retained for further analysis, *ponA* still displayed a significantly reduced width in the first step of our screen (Table S4), as expected (Fig. 1C). However, *mreBH, lytE* and *rny* (*ymdA*) did not display a significant width difference in our experimental conditions (Table S4). It should be noted too that a *rodZ* null mutant is present in the BKK library (24) even though the *rodZ* gene was originally reported to be essential in *B. subtilis* (22). We addressed this apparent discrepancy and showed that *rodZ* is not essential for growth in *B. subtilis*, at least in the experimental conditions tested. We also confirmed that Δ*rodZ* cells display division defects (52) and found that they display shape alterations in some media, and that this phenotype is influenced by the parental genetic background (see Sup. Information and Fig. S3).

The 9 new cell width determinants identified in our screen belong to different functional categories (Table 1). Interestingly, only one of them, *dacA* (encoding PBP5, a bPBP involved in PG maturation (53, 54)), is directly involved in CW homeostasis. The most represented functional category among our newly identified width-deficient mutants is the metabolism (Table 1). One gene, *guaA*, is involved in purine nucleotide synthesis (encoding the GMP synthetase (55)), and four are part of the central carbon metabolism (Fig. S4): *pyk*, specifying the pyruvate kinase acting in glycolysis (56); *ptsH*, encoding HPr, a component of the sugar phosphotransferase system (PTS) (57); *panD*, involved in coenzyme A biosynthesis (58); and *rpe* (*yloR*), predicted to encode the ribulose-P-epimerase (Rpe) of the pentose phosphate pathway. Although the role of *rpe* has not yet been investigated in *B. subtilis*, the prediction got a score >99.91 % using a hidden Markov model-based homology prediction tool (HHpred; (59, 60)). Noteworthy, while most mutants involved in CW synthesis were wider, all metabolism mutants but *rpe* were thinner (Fig. 3B, 3C; Table 1). Out of the three remaining genes selected, one is involved in cell division (*minJ* (61, 62)), one in translation (*yaaA*, encoding a ribosome assembly factor (63)), and one is annotated as a putative transcriptional regulator (*ybzH*, (64)). Using the HHpred homology prediction tool (59, 60), we confirmed that *ybzH* encodes a probable HTH-type transcriptional regulator sharing strong structural resemblance with proteins of the ArsR and GntR families or transcriptional repressors. Regulators of the ArsR-type are involved in the stress-response to heavy-metal ions and GntR-family members in various metabolic pathways including fatty acid, amino acid, or gluconate metabolism (65, 66).

### *ΔminJ* and Δ*rpe* display a phenotype of cell diameter instability

The mutant strains selected in our screen displayed a thinner or a larger mean diameter relative to wild type cells. Although their mean width differs from the wild type, most of these mutants still control their diameter to maintain it constant over generations. However, two of these mutants, Δ*rpe* and Δ*minJ*, displayed a distinctive large dispersion of width values (Fig 3B, Table S5).

The Δ*rpe* mutant from the BKK library (BKK15790) was first selected based on its reduced width (−11.4 %) during the HCSm analysis (Table S5). Surprisingly, once backcrossed, the Δ*rpe* strain (RCL856) displayed the opposite phenotype with an increased width (+11 %) (Fig. 4A; Table S5), nonetheless indicative of a defect of width control. Another striking difference between the two strains was their width dispersion (Fig. 4A). While the BKK Δ*rpe* showed thin and regular cell diameters with a dispersion of values similar to the control strain, widths of the backcrossed mutant displayed the largest variability of our dataset (Fig. 4A, Fig. 3B and Table S5). Furthermore, the backcrossed Δ*rpe* mutant formed small slow growing colonies while its BKK parent did not (Fig. S5). Taken together, these results suggest that the Δ*rpe* mutant present in our BKK collection had acquired some suppressor mutation(s), partially restoring its growth and reducing its width variability.

**Figure 4.**
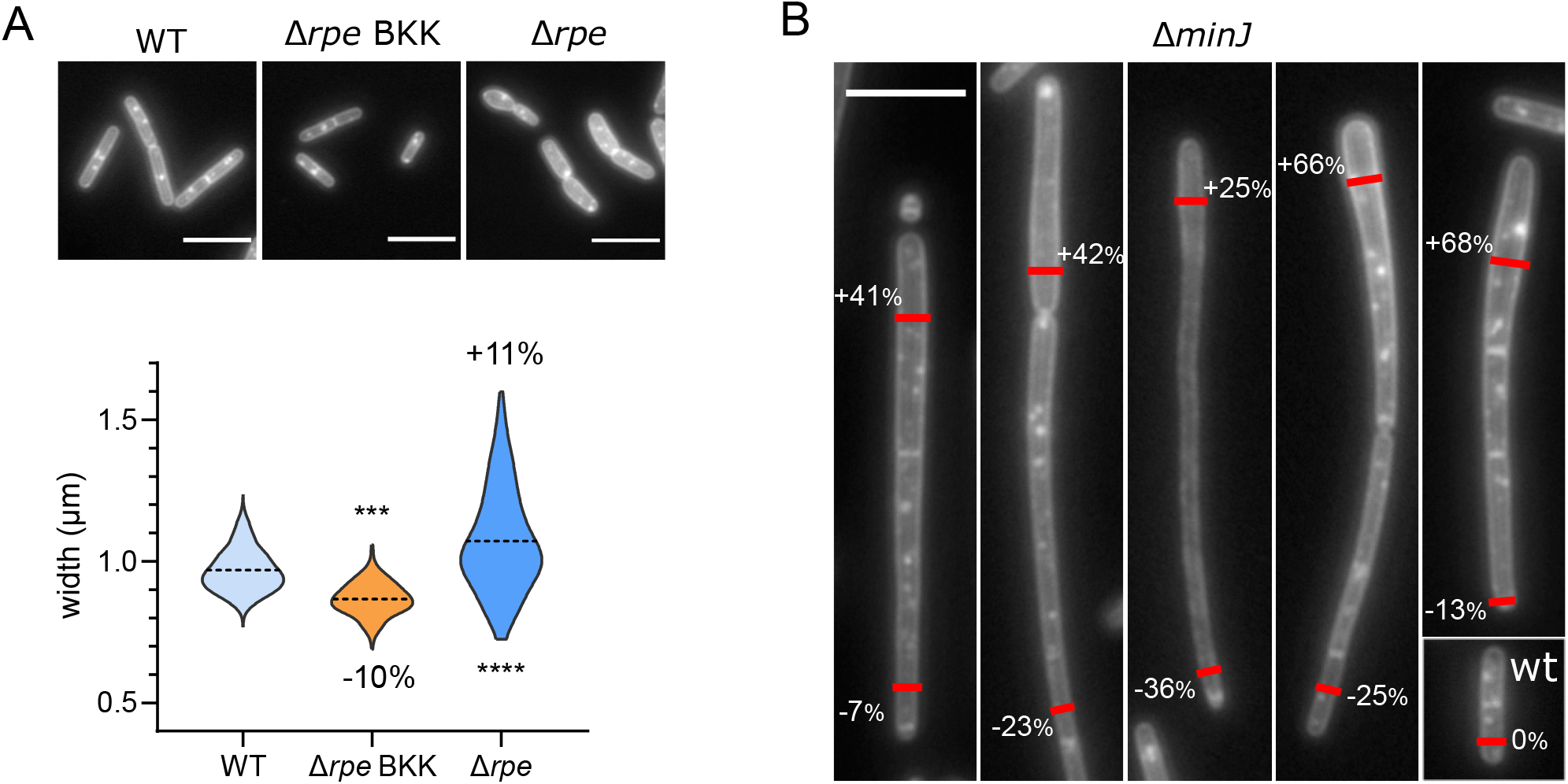
Δ*minJ* and Δ*rpe* mutants exhibit an uncontrolled diameter phenotype. **A**. Images of membrane-labelled strains and corresponding distribution of cell widths for the wild type strain, Δ*rpe* from the BKK collection and Δ*rpe* backcrossed into 168 wild type background. Broken line: mean. The differences between the means of the mutants and the wild type are indicated, in %. Data are compilations of two independent experiments and statistical significance between mutants and the wild type strains was calculated with a nested t-test. **B**. Images of membrane-labelled Δ*minJ* mutant, presenting varying width along single cells or chain of cells. This phenotype is observed in approximately 19 % of the population. Percentages indicate the difference of width compared to that of the wild type strain, at the position of the red marks. Scale bars, 5µm.

The second strain with a variable diameter was Δ*minJ* (Fig. 3B and Table S5). This mutant of the “Min” system involved in division site selection, displays reduced septation, leading to long filamentous cells (Fig. 4B) (52, 61). Although the average width of Δ*minJ* cells was marginally affected (−1%), the SD was unusually large, with widths ranging from 0.7 to 1.4 µm (Fig. 3B and Fig. 4B). Furthermore, uneven diameters were observed along the lengths of individual Δ*minJ* filamentous cells (Fig. 4B). This phenotype displayed a polarity, with cells progressively widening or thinning from a cell pole, and affected only a fraction of the population (19.16 %, n> 428), which could explain why it was not previously reported.

### Mutants of metabolism and cell wall homeostasis are differently affected in growth and S/V ratio

During the phenotypic characterization of the 13 width-deficient mutants selected in our screen, we noticed a variability of cell length too, with cells of the *guaA* mutant unambiguously being the shortest and cells of the *minJ* mutant forming very long cells (Fig. 3C). We quantified the average length of all mutants and found that to the exception of *cwlO*, they were all significantly longer or shorter than the wild type (Fig. 5A ‘Length’, Table 1). It should be noted that Δ*minJ* and Δ*rodZ* mutants form minicells (Fig. S6)(22, 67) that were not taken into account in our length quantification. Interestingly, shorter and wider mutants were all related to CW homeostasis while metabolism mutants were shorter and thinner than the wild type, again to the exception of Δ*rpe* (Fig. 3B, 3C, Fig. 5A and Table 1). However, no direct correlation between cell width and length was observed across the strains (Fig. S7). Because cell length - but not width-of *B. subtilis* usually correlates with growth rate (the “growth law”) (26, 28), we wondered if differences of cell length between the mutants would mirror differences in growth rate. The generation time (GT), determined during mid exponential growth (see ‘Materials and methods), showed no significant difference with the GT of the wild type, for CW homeostasis mutants (Fig. 5A ‘GT’). However, the metabolism mutants displayed an increased GT of > 63 % relative to the wild type (Fig. 5A). For these strains, the GT strongly correlated with the average cell length (R^2^=0.837) indicating that in such mutants the “growth law” is conserved (Fig. 5B). In contrast, no correlation was observed between their GT and their cell width (Fig. 5B), further indicating that these two parameters are not connected.

**Fig. 5.**
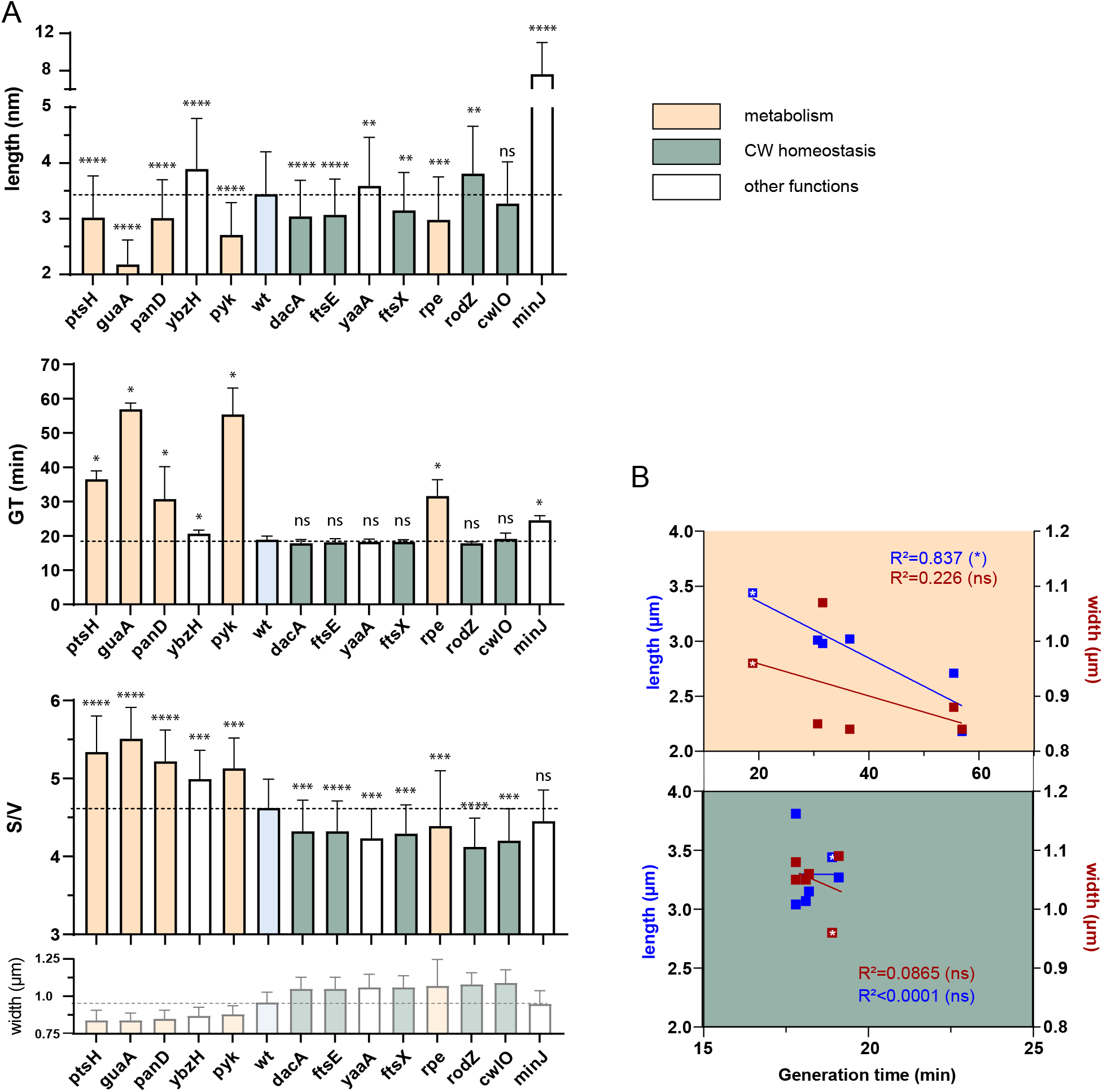
Relationship between generation time, length, width and surface to volume ratio in the selected mutants. **A**. Average length, generation time (GT) and surface to volume ratio (S/V) of cell width deficient (backcrossed) mutants compared to the wild type strain. Width of each strain is recalled (from Fig. 3) for comparison. The dotted line marks the level of the average wild type value. GT are calculated on populations (see the corresponding method section) and are the average of 4 independent experiments. Length and S/V ratio data are calculated per cell and compiled from three independent experiments. Statistical significance were determined as described in the method section ‘statistical analysis’, and displayed with * for *P*-values. Error bars represent SD. **B**. Average length and width as a function of the generation time (GT). Upper panel: metabolism-related mutant; lower panel: CW homeostasis-related mutant. R^2^ of the linear regressions (lines) are indicated on the panels. White stars indicate the wild type values.

Finally, we calculated the surface area (S) to volume (V) ratio of the mutants. This parameter was proposed to be maintained constant in a given condition, as a key determinant of cell shape (68). The S/V was significantly altered in all our mutants excepting Δ*minJ*. All CW mutants displayed a reduced S/V ratio (Fig. 5A ‘S/V’), a consequence of the increased width and the subsequent increased cell volume (the length having a limited contribution to it (Fig. 5A ‘length’, 5B). This S/V reduction is reminiscent of the effect previously reported for fosfomycin-treated bacteria, an antibiotic inhibiting PG biosynthesis (68), and is consistent with the proposed model that reduction of the rate of S growth (i.e when CW synthesis is reduced) increases cell width and reduces S/V (68). In contrast, all metabolism mutants but Δ*rpe* displayed a larger S/V as a consequence of the important drop of both width and length affecting the surface and the volume (Fig. 5A and B).

Taken together, our results discriminate between two main groups of width-deficient mutants with specific phenotypes. Mutants of metabolic genes (to the exception of the Δ*rpe*) display a reduced width and increased S/V, and are strongly impaired in growth while, conversely, mutants affected in CW homeostasis display an increased width and reduced S/V, but their GT is unaffected relative to the wild type.

## Discussion

Cell width is probably one of the most tightly regulated physiological parameters in *B. subtilis* (35). However, the mechanisms allowing its fine control remains unclear. Our approach aimed at revealing in a systematic way non-essential genes involved in this process. We confirmed several of the previously reported non-essential genes acting on *B. subtilis* width control (Table S1) and identified 9 new genes whose deletion strongly affects *B. subtilis* diameter. Since we arbitrarily set up a cutoff to select the most drastically affected mutants (top 1%), it is likely that additional genes contribute to width control, along with essential genes or genes acting synthetically. A quick survey of our screen data with a less stringent cutoff (top 10 % most affected mutants, Table S4) shows a few dozens of genes involved in CW (e.g. *walH, pbpG, lytG, yocH, murQ, murE*…), lipid metabolism (*fabI, lipL, araM, fadE*…) and central carbon metabolism (*tkt, ywjH, coaA*…). This list should nevertheless be taken with caution because the two-step verification performed on our top 1% selection revealed a significant number of false positives, and because the high-throughput-constructed BKK library may contains suppressors, as exemplified in this work with the *rpe* mutant.

To our knowledge, most genes previously described to affect cell width are directly involved in CW homeostasis (Table S1). The remaining genes (~1/4) are involved in a variety of pathways, but they were shown to affect the levels or localization of CW synthetic proteins or the levels of PG precursors (Table S1). In agreement with this, many mutants identified in our screen are related to CW homeostasis as well. In addition to *cwlO, rodZ, ftsX* and *ftsE*, whose mutants were known to display width defects, we identified *dacA*, encoding the major vegetative DD-carboxypeptidase PBP5, responsible for the maturation of the PG by trimming the terminal D-Ala of the pentapeptide (69).

Unexpectedly, we also identified several genes involved in width control that belong to other functional categories, including five metabolic genes: *ptsH, guaA, rpe, pyk* and *panD*. So far, the only metabolic gene described to affect cell width was *glmR*, which encodes a regulator controlling the carbon flux that stimulates the PG precursor synthetic pathway in neoglucogenic conditions (70, 71). Other studies have linked the cell metabolic status with cell size, although in these cases the mutants were affected in cell length (reviewed in (28)). Out of these, only *pyk*, encoding PykA, which produces pyruvate in the final step of glycolysis (Fig. S4), was identified in our screen, which may suggest a central role for this protein to coordinate the cell metabolic status with the control of length/division and width/elongation.

Another salient point of the study is that the two main groups of width-deficient mutants are discriminated by their phenotypes. Genes involved in CW homeostasis display an increased width and a reduced S/V but unaffected growth rate, while metabolism mutants display a reduced width and an increased S/V, and their growth rate is strongly affected. Based on these observations, it is tempting to speculate that *yaaA* is somehow involved in CW homeostasis while *ybzH* function may be directly or indirectly related to a metabolic pathway. In this dichotomy, *rpe* is stepping out, sharing characteristics of both groups and thus suggesting that its phenotypes might reflect a defect in both pathways. This hypothesis is strengthen by the presence of genes connected with cell shape control in the same operon than *rpe*: *prpC, prkC* and *cpgA*. PrkC is a Ser/Thr kinase –and PrpC its cognate Ser/Thr phosphatase-regulating many proteins including some reported to affect cell width in *B. subtilis* as LtaS, YfnI, YqgS (72), CpgA (73), GlmR (YvcK) (74), RodZ (75) and GpsB (76) (Table S1). CpgA was recently shown to moonlight as a detoxifying enzyme of erythronate-4P, whose accumulation induces a depletion of fructose-6P, the entry of the PG precursor pathway (77). Thus, the *rpe* operon may be at the crossroad between the metabolic and CW homeostasis pathways.

Noteworthy, the phenotypes of the CW mutants are consistent with the model proposed by Harris and coworkers (68). They proposed a ‘relative rate’ model in which the rates of S and V growth are both functions of V (and not functions of S and V, respectively) and that S/V is the key parameter maintained constant in a given condition rather that the respective rates of S and V expansion (68). A consequence of their model is that a diminishing rate of S growth, for example when reducing the CW synthesis, both increases cell width and reduces S/V, even for a constant growth rate. Thus, one could hypothesizes that the increase of width observed in the mutants identified in our screen may be a direct consequence of a crippled cell surface synthesis.

This could also be interpreted in the light of the Rod/aPBP balance model from Dion and coworkers in which the unbalance activity between the two CW synthetic machineries leads to thinner or larger cells (7). According to this model, the CW mutants selected in this study (*dacA, ftsE, ftsX, cwlO* and *rodZ*) would present an unbalanced PG-synthesizing activity in favor of the aPBPs. Following the same line of thought, the metabolism mutants identified in this study, slender (to the exception of *rpe*), should present the opposite imbalance, with increased activity of the Rod complex or decreased activity of aPBPs.

In summary, cell with control appears as a very tightly regulated process in which different cellular circuits are at play. Our results indicate that metabolism is a major contributor to the control of cell width, suggesting the presence of unsuspected regulators or moonlighters affecting the synthesis of the CW. Among the genes identified here, 3 are stepping out and are of particular interest. On the one hand, RodZ acts on both cell division and elongation, and its activity depends on the medium composition (Fig. S3), strengthening a possible link between metabolism/width. On the other hand, *minJ* and *rpe* mutant cells display unique uncontrolled width suggesting that the ‘check and balance’ of width control is lost. Deciphering how these genes affect the control of cell width of *B. subtilis* will be a challenge for future research.

## Materials and methods

### General methods and bacterial growth conditions

Methods for growth of *B. subtilis*, transformation, selection of transformants and so on have been described extensively elsewhere (78). DNA manipulations were carried out by standard methods. *B. subtilis* strains used in this study are listed in Table S6. *B. subtilis* strains were grown at 30 °C or 37 °C in rich lysogeny broth medium (LB), except for assaying growth in poor media, where strains were grown in Modified Salt Medium (MSM) supplemented with 10 mM MgSO_4_ (48) and S medium (33) with the corrected 1.2 µg/ml of MnSO_4_. For precultures, media supplements were added at the following final concentrations: MgSO_4_ 20 mM, neomycin 15 µg.ml^-1^, spectinomycin 100 µg.ml^-1^ or chloramphenicol 5 µg.ml^-1^ (Table S6). Transformants were selected on LB agar plates supplemented with MgSO_4_ and neomycin. For the determination of generation time (GT), cells from overnight cultures were diluted to a fixed starting OD_600nm_ of 0.01 in fresh LB medium supplemented with MgSO_4_ in 96-well cell culture plates (CellStar) and grown in a microplate reader Synergy2 (BioTek Instruments, Vt, USA) at maximum rpm at 37°C. GT was calculated using a Matlab script available at the following link: https://github.com/CyrilleBillaudeau/GenerationTime_ofBacteria_withOD.

### General screening procedure

Our screening was performed on the BKK library (24) using HCSm setups (see ‘High content screening microscopy’), leading to the selection of 3 974 out of the 3 983 mutants from this collection (9 clones were absent from the published library or failed to re-grow). Images were processed and the cell diameter was measured (see ‘Image processing and cell size quantification’). The 1% most affected strains (40 mutants) were selected and their phenotype confirmed using an epifluorescence microscope (see ‘Low throughput epifluorescence microscopy’ and ‘Image processing and cell size quantification’). Deletions in the selected clones were verified by PCR, revealing that 2 mutants (*yoqC, yorP*) were wild type for the expected locus and thus they were discarded for further analysis. The remaining 38 mutants were back-crossed into the wild type 168 strain to be analyzed over triplicate experiments using low throughput microscopy. An arbitrary cutoff of 8% Δwidth, obtained by comparison to the wild type strain width, was chosen and 12 genes were finally selected.

### High content screening microscopy

Cells from overnight cultures, grown in the presence of neomycin and MgSO_4_, were diluted at 1/600 in fresh LB medium supplemented with MgSO_4_ in 96-well cell culture plates (CellStar) and grown on an orbital shaker at 250 rpm at 37°C until mid-exponential phase (OD_600_ ~ 0.2). To fix the cells, 150 µL of culture were mixed with 50 µL of fixation solution (0.5 M KPO_4_ pH 7, 8 % paraformaldehyde, 0.08 % glutaraldehyde) in 96-well PCR plates and incubated 15 min at room temperature followed by 15 min on ice. The cells were pelleted by a 5 min centrifugation at 450 g, and the supernatant carefully removed by pipetting. The pellets were washed with 200 µL of washing buffer (KPO_4_ 0.1 M pH 7), centrifuged again, resuspended in 20 µL of water containing 3.3 µg/mL FM™1-43FX (ThermoFisher, F35355) and incubated 5 min at room temperature. 180 µL of washing buffer were added and the cells were centrifuged a last time to be concentrated 3.75 × in 40 µL washing buffer. 96-well (Fisher) or 384-well (Greiner) microscopy plates were treated with 60 µL of poly-L-lysine 0,01 % and washed with 60 µL of deionized water. 40 µL of cells were put in each well and discarded after a 1 min incubation. Finally, 40 or 120 µL of deionized water were added into each well of 96-well or 384-well plates. Imaging was performed either on an ImageXpress micro Confocal (Molecular Devices) or an IN Cell 6000 Analyzer (GE Healthcare) used in non-confocal mode. The ImageXpress HCS microscope was equipped with a 60 × Nikon air objective (NA 0.95), a FITC filter (Ex.488/Em.536) and a Zyla 4.2 Andor sCMOS camera with a final pixel size of 115 nm and controlled by MetaXpress software package. The INCell 6000 analyzer was equipped with a 60 × water objective (NA 0.95), a FITC filter (Ex.488/Em.525) and a sCMOS 5.5 Mpixels camera with a final pixel size of 108 nm and controlled by the INCell 6000 Analyzer - Acquisition v.7.1 Software. Images from 4 fields of view were acquired for each strain.

### Low throughput epifluorescence microscopy

Cultures were performed as for HCS microscopy but in shaking tubes instead of microplates. For live cell imaging, 300 µL of cultures were directly mixed with FM™ 1-43FX (ThermoFisher) to reach the concentration of 3.3 µg/mL and concentrated 3.75 × before 1 µL of the preparation was spotted onto a thin 2 % agarose-LB pad, topped by a coverslip and immersion oil, and mounted immediately in the temperature-controlled microscope stage. For the imaging of fixed cells, cells were fixed as described for HCS microscopy except that 300 µL of culture were mixed with 100 µL of fixation solution and subsequently washed with 300 µL of buffer. Cells were spotted on a 2 % agarose-LB pad or on poly-L-lysine-treated 96 well microscopy plate. For the latter, the wells were washed then filled with deionized water. Epifluorescence images of the membrane-stained cells were acquired on a previously described setup equipped with a 100 × objective (36).

### Image processing and cell size quantification

The post-acquisition treatment of the images was done with the Fiji software and the measures (mean cell diameter and length) with the MicrobeJ plugin (44–46). In MicrobeJ, the cell width was calculated as the mean value along the medial axis of the cell. The parameters used for the MicrobeJ module are listed in Table S2. Cells aggregates were excluded and segmentation was manually corrected when necessary.

During high-throughput screening, the cell width of each strain was calculated as the mean of 225 cells (in average). When the four-image set contained less than 30 measurable cells, a new acquisition was performed. Because the library is devoid of a wild type reference, and to prevent putative plate-to-plate variability, the mean cell width of each mutant was compared to the average width of all measured cells of its 96-well plate, the ADP (<u>a</u>verage cell <u>d</u>iameter per <u>p</u>late) index (17-19.10^3^ cells/plate) (Fig. S2A). Each strain’s diameter deviation relatively to this index was calculated, as:

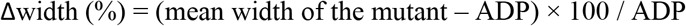

From these differences, the 0.5^st^ and 99.5^th^ percentile were calculated and the 99 % of the mutants between these two values eliminated.

During low-throughput microscopy (for the verification of the BKK candidates and for clones resulting from the backcross into the 168 strain) the cell width and length of each strain was calculated as the mean of 245 cells (in average). The calculated Δwidth was expressed by comparison with the wild type cell size.

### Alternative methods for cell width measurement

Cell widths were measured either with the ChainTracer plugin of the Fiji software, or by determining “manually” the width at maximum height on intensity profiles (79) (Fig. S1). For ChainTracer, we used a stack of phase contrast and epifluorescence images of membrane-stained cells, and only analyzed isolated chains of cells to prevent segmentation issues. For the measurement using intensity profiles, a line was manually drawn perpendicularly to the cell’s long axis, on epifluorescence images of stained membranes, and a profile plot of the fluorescence intensity was generated. The cell diameter was determined by measuring the distance between the two maxima.

### Statistical analysis

All statistical analyses were performed with Prism 9 (GraphPad Software, LLC). To analyze the variance between replicates, a multiple (pairwise) comparison was performed using one-way ANOVA (Fig. 1A). For pairwise comparison between means of a control and its tested sample with 2 or more replicates, we performed nested t-tests (Fig. 1B, 1C, 3B, 4A, 5A_’length’, 5A_’S/V’). Note that the plots show the pooled values of the replicates. When t-tests were not possible (e.g. if N<30), pairwise comparisons were done with a non-parametric Mann-Whitney test (Fig. 5A_’GT’). *P*-values are displayed as follows: **** = P<0.0001; *** = 0.0001<P<0.001; ** = 0.001<P<0.01; * = 0.01<P<0.05; ns = P>0.05.

## Supporting information

all supplementary informations, tables, and figures

## Acknowledgments

This project has received funding from the European Research Council (ERC) under the European Union’s Horizon 2020 research and innovation program (grant agreement No 311231 and grant agreement No 772178, to R.C.-L.). We thank Alexandre Vandeputte and the BioImaging Center Lille-Nord de France (BICeL) facility (Lille, France) for HCSm acquisitions. Financial support for the HCS equipment was provided by the FEDER (12001407 (D-AL) Equipex Imaginex BioMed).

## List of supplementary materials

### Supplementary information

Comparison of width measurements obtained with different methods

RodZ, a non-essential protein involved in cell shape control

List (names and sequences) of oligonucleotides

### Supplementary tables

Table S1. Genes reported to affect cell width in *B. subtilis*

Table S2. Settings used for MicrobeJ plugin

Table S3. Average width differences (%) across replicates

Table S4. Cell width of mutants of the BKK collection

Table S5: Width of the 0.5% largest and thinnest selected strains

Table S6. *B. subtilis* strains used in this study

### Supplementary figures

Fig. S1. Comparative cell width distributions of wild type *B. subtilis* cells.

Fig. S2. ADPs are constant across plates and equal to the width of the wild type strain.

Fig. S3. Growth and cell shape of *B. subtilis rodZ* mutants vary depending on the growth media and the genetic background.

Fig. S4. Carbon metabolic pathways involving the selected mutants deficient for cell width control.

Fig. S5. Backcross of the *rpe* deletion reveals a strong growth defect. Fig. S6. The *rodZ* and *minJ* mutants form minicells.

Fig. S7. Average cell lengths as a function of average cell widths for each mutant.

### Sup. Figures legends

**Fig. Sup. 1. Comparative cell width distributions of wild type *B. subtilis* cells. A**. Cell width of live and fixed cells were measured using the Fiji plugins MicrobeJ and Chain tracer, or by manual measurements (see ‘Materials and methods’). The differences between the means of live and fixed cells (in %) is specified for each method. **B**. Comparative cell width distribution of fixed wild type *B. subtilis* cells grown in LB with and without 20 mM magnesium supplementation. Broken line: mean. Differences between the means, expressed as a percent, are indicated on the plots. Statistical analysis were performed using nested t-tests. Data (A, B) are compilations of at least two independent experiments.

**Fig. Sup. 2. ADPs are constant across plates and equal to the width of the wild type strain. A**. ADP of the 48 96-well plates containing the BKK library. Each ADP is the mean of all measured cell widths (~20 000) on a plate, and error bars are the standard deviations (SD). For the 48^th^ plate, the acquisition was performed on the epifluorescence microscope (with 100x magnification) and not on the HCSm, which explains the reduced values (as in Fig. 1C). **B**. Comparison of the average of the ADPs of all 48 plates and the average width of a wild type cell population measured with the HCSm. There is no significant difference between the two values according to the Mann-Whitney non-parametric test, indicating ADPs are similar to the wild type diameter (1.160 µm vs 1.153 µm, respectively). Similarly, the ADP calculated for the 48^th^ plate is close to the wild type strain measured in the corresponding microscope (0.975 µm vs 0.964 µm, respectively). Error bars are SD.

**Fig. Sup. 3. Growth and cell shape of *B. subtilis rodZ* mutants vary depending on the growth medium and the genetic background. A-B**. Typical growth curves of wild type and Δ*rodZ* mutants of *B. subtilis* in rich LB (blue), poor MSM (green) and S (orange) media. Strains are derivative of the 168 (A) or the PY79 (B) wild type parental strains, and either wild type (plain), or deleted for *rodZ* (dashed; CcBs351 or CcBs628). Panel A displays the growth of two additional Δ*rodZ* mutants, the one from the BKK library (circles; BKK16910) and the BKK Δ*rodZ* mutant backcrossed into the 168 wild type strain (dotted; RCL828).

**C-D**. images of *B. subtilis* cells grown to mid exponential growth phase, stained with the FM1-43FX membrane dye. Strains are derivative of the 168 (**C**) or the PY79 (**D**) wild type strains, and either wild type (‘wt’), carrying our *rodZ* deletion (CcBs351; CcBs628), *rodZ*-from the BKK library (BKK16910), or *rodZ*-from the BKK backcrossed into 168 wild type (RCL828). Scale bars: 1 µm.

**E-F**. Cell width distribution of the *rodZ* mutants (black) and their parental wild type 168 (E) or PY79 (F) strains (blue), grown to mid exponential phase in LB and MSM media. Data are compilations of two independent experiments. Statistical significance of the comparison between each mutants and the wild type strain was estimated using the Mann-Whitney non-parametric test (**** = *P*-value <0.0001).

**Fig. Sup. 4. Carbon metabolic pathways involving the selected mutants deficient for cell width control**. The four genes selected in our screen that are involved in carbon metabolism, *panD, ptsH, pyk* and *rpe*, encode enzymes required for the pentothenate, glycolysis and pentose pathways, respectively. PanD converts L-asp into β-alanine, the first step of the pathway leading to Coenzyme A (Co-A) synthesis. Co-A is used in glycolysis as a substrate for Pyk to form acetyl-CoA. HPr (encoded by *ptsH*) is a bi-functional protein acting also in glycolysis as a part of the PTS (phosphotransferase system) required for the import/phosphorylation of sugars, and in the regulation of the carbon catabolite control, as an allosteric regulator with CcpA. Rpe is predicted to produce xylulose-5P in the pentose pathway.

**Fig. Sup. 5. Backcross of the *rpe* deletion reveals a strong growth defect**. Chromosomal DNA of strain BKK15790, knockout for *rpe*, was transformed into the wild type *B. subtilis* 168 strain (wt) to generate strain RCL856. The RCL856 strain displays a ‘small-colony’ phenotype indicative of a growth defect. Isolated colonies of each strain were streaked on an LB plate and grown for 24h at 37°C.

**Fig. Sup. 6. The *rodZ* and *minJ* mutants form minicells**. Display are images of the Δ*rodZ* (strain RCL828) and Δ*minJ* (RLC834) mutants grown to mid exponential phase in LB medium, stained with FM1-43FX and fixed, imaged by epifluorescence microscopy. Arrowheads point to minicells. Scale bar: 2 µm.

**Fig. Sup. 7. Average cell length as a function of average cell width for each mutant**. The wild type strain is labelled in light blue. R^2^ of the linear regressions (lines) are indicated on the panel. Data are compilations of at least three independent experiments.

## Notes

### Competing Interest Statement

The authors have declared no competing interest.

### Summary of Updates

The last 3 sections of the results have been reorganized as well as the corresponding figures 5 and S3 (/!\ Sup figure order modified). There is also a new discussion plus many typo and small rewriting all over the text. Figures are now colorblind friendly.

