## Supplementary material for "A high content microscopy screening identifies new genes involved in cell width control in *Bacillus subtilis*": all supplementary informations, tables, and figures

**Comparison of width measurements obtained with different methods.**

Because MicrobeJ was designed to determine cell width based on phase contrast images (an option not available on the HCS microscopes at our disposal) and not on fluorescent images of membrane-stained cells, we first tested the feasibility of using the plugin to our needs. Using the settings reported in Table S2, we compared the estimation of width on a population of wild type *B. subtilis* cells obtained with MicrobeJ (1) with another Fiji/ImageJ plugin, ChainTracer (2), and with a manual measurement of mid-cell widths. The measures were made on both live and fixed cells stained with the FM1-43FX membrane dye, on a Nikon N-Ti epifluorescence microscope. Additional phase contrast images were acquired as they are mandatory for the ChainTracer process (the main reason why this plugin could not be used in our screen). As seen on Fig. S1A, the distribution of cell width differs between methods, ChainTracer slightly minimizing and MicrobeJ maximizing the estimates. Although the average width measured with ChainTracer is closer to that obtained with a manual measurement than the width measured with MicrobeJ, we noticed that a large majority of cells were excluded from the automatic measurement with ChainTracer (~60 %), and that the width distribution was unexpectedly bimodal (Fig. S1A). Conversely, both MicrobeJ and the manual measures gave similar Gaussian distribution of cell widths. We also noticed a slight reduction of widths when measuring fixed cells, this time with all three methods. We concluded that, despite a larger estimation of the width, MicrobeJ is an efficient method for a relative estimation of the width, able to discriminate between average widths varying of only a few percent.

**RodZ, a non-essential protein involved in cell shape control** The *rodZ* gene was originally reported to be essential in *B. subtilis* (3). However, a more recent report indicated that its inactivation leads to robust growth and only mild shape defects (4). Furthermore, three available *B. subtilis* knockout libraries include a Δ*rodZ* mutant (5, 6). To address this discrepancy, we decided to construct by homologous recombination a new, independent *rodZ* deletion mutant (strain CcBs351).

In this new *rodZ* knockout mutant most of the open reading frame (863/914 bp) was replaced, by double cross-over recombination, with a chloramphenicol resistance cassette (*cat*). For this, the upstream and downstream flanking regions of *rodZ* were PCR amplified using chromosomal DNA of *B. subtilis* 168 as a template and oligonucleotides cc295/cc292 and cc293/cc296, respectively. The *cat* cassette was PCR amplified using cc291/cc294 as primers and pAH328 as DNA template (7). The three generated DNA fragments were combined by isothermal "Gibson" assembly (8) and transformed into wild type 168 *B. subtilis* strain, competent for natural genetic transformation. To prevent the potential appearance of suppressor mutations, transformants were selected on LB medium supplemented with 20 mM magnesium, in addition to the selection pressure. Isolated clones were subsequently checked, by sequencing the complete area that was subjected to PCR amplification. This new mutant was readily constructed. Similarly, the backcross of the Δ*rodZ* from the BKK library (BKK16910) into the wild type 168 strain was performed and gave countless positive transformants (strain RCL828), further suggesting the non-essentiality of the *rodZ* gene in *B. subtilis* in the 168 genetic background.

We then assayed for growth and shape in different conditions the three *rodZ* mutants to our disposal: the newly constructed strain (CcBs351), the original mutant from the BKK library (BKK16910) and its backcross (RCL828). In rich LB medium, the Δ*rodZ* strains displayed no difference of growth (Fig. S3A) or cell shape (Fig. S3C), but were slightly wider than their parental wild type strain (Fig. S3E). We also confirmed the presence of minicells (Fig. S6), indicative of the perturbation of the division process as previously reported (9). When grown on the poorer MSM medium, Δ*rodZ* cells were significantly wider and frequently divided asymmetrically (Fig. S3C, E) and displayed a solid but slightly reduced growth rate compared with the wild type strain (Fig. S3A). This reduction of growth in MSM might not be directly linked to the richness of the medium since all strains grew almost identically in the much poorer citrate/glucose-based S medium (Fig. S3A). Notably, the three *rodZ null* mutants did not grow identically. The strain BKK16910 displayed the highest reduction of maximum cell density and highest growth lag compared to the two other mutant strains. Since the backcrossed deletion of the BKK16910 (RCL828) and our independently constructed deletion mutant CcBs351 did not display this growth defect, we inferred that some unknown genetic differences in the BKK16910 strain rather than the *rodZ* deletion itself could be at play.

The mutants previously published in the literature were constructed in different genetic backgrounds, namely the 168 (4-6) and PY79 (3) wild type laboratory strains. The 168 and PY79 strains share a common origin, the ancestral 3610 "Marburg" wild isolate of *B. subtilis*, but are the product of different histories (mutagenesis and selection cycles) that drove to significant genomic differences (mutations, deletions and rearrangements) (10). We thus wondered if the impossibility to obtain a *rodZ* null mutant originally reported could have been be due to the use of the PY79 (3) instead of the 168 wild type used by us and others. We then transferred our newly constructed *rodZ* deletion into the PY79 background (strain CcBs628). The transformation caused no difficulties and the cell shape defects of the resulting CcBs628 mutant appeared minimal in rich medium, with only a slight increase of cell width (Fig. S3D, F). Again, the increased width was more pronounced when the cells were grown in poorer media (Fig. S3D, F), but overall the shapes of the Δ*rodZ* mutants were similar in both the 168 and PY79 background. The only notable difference between the 2 genetic backgrounds was a strong growth lag of the *rodZ* null in PY79, in the poor S medium (Fig. S3B).

Taken together, these results indicate that (i) *rodZ* is not essential for growth in *B. subtilis*, (ii) cells lacking *rodZ* display division defects and limited width and growth alterations, which are accentuated in poor growth medium, and (iii) that the genetic parental background influences this sensitivity to the gr owth medium.

<u>List (names and sequences) of oligonucleotides:</u>

**cc291** CAAAGAAGCCAGAGAGGAAAAAGCAATGAACTTTAATAAAATTGATTTAGACAATTGG
**cc292** CCAATTGTCTAAATCAATTTTATTAAAGTTCATTGCTTTTTCCTCTCTGGCTTCTTT
**cc293** TAGGCCTAATGACTGGCTTTTATAATTACCAGATGACTTTTCTTCACG
**cc294** GTGAAGAAAAGTCATCTGGTAATTATAAAAGCCAGTCATTAGGC
**cc295** GCACTCACTAGGAAGAGAGGG
**cc296** CACGTCAGAGCCTTCGATCAC

Fig. Sup. 1.

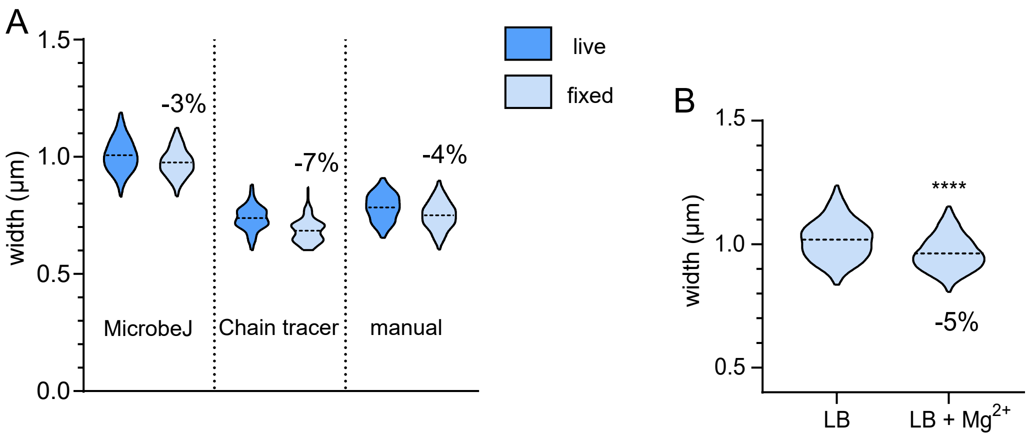


Fig. Sup. 1. Comparative cell width distributions of wild type *B. subtilis* cells. **A**. Cell width of live and fixed cells were measured using the Fiji plugins MicrobeJ and Chain tracer, or by manual measurements (see 'Materials and methods'). The differences between the means of live and fixed cells (in %) is specified for each method. **B**. Comparative cell width distribution of fixed wild type *B. subtilis* cells grown in LB with and without 20 mM magnesium supplementation. Broken line: mean. Differences between the means, expressed as a percent, are indicated on the plots. Statistical analysis were performed using nested t-tests. Data (A, B) are compilations of at least two independent experiments.

Fig. Sup. 2.

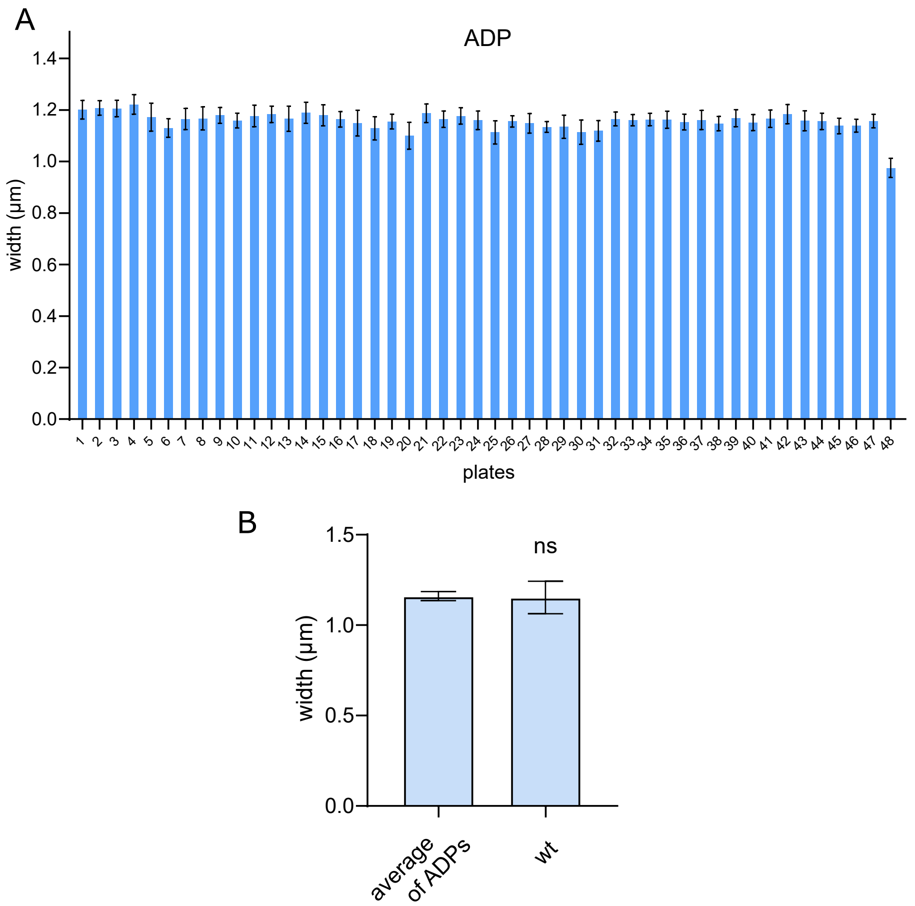


Fig. Sup. 2. ADPs are constant across plates and equal to the width of the wild type strain. **A**. ADP of the 48 96-well plates containing the BKK library. Each ADP is the mean of all measured cell widths (~20 000) on a plate, and error bars are the standard deviations (SD). For the 48th plate, the acquisition was performed on the epifluorescence microscope (with 100x magnification) and not on the HCSm, which explains the reduced values (as in Fig. 1C). **B**. Comparison of the average of the ADPs of all 48 plates and the average width of a wild type cell population measured with the HCSm. There is no significant difference between the two values according to the Mann-Whitney non-parametric test, indicating ADPs are similar to the wild type diameter (1.160 µm vs 1.153 µm, respectively). Similarly, the ADP calculated for the 48th plate is close to the wild type strain measured in the corresponding microscope (0.975 µm vs 0.964 µm, respectively). Error bars are SD.

Fig. Sup. 3.

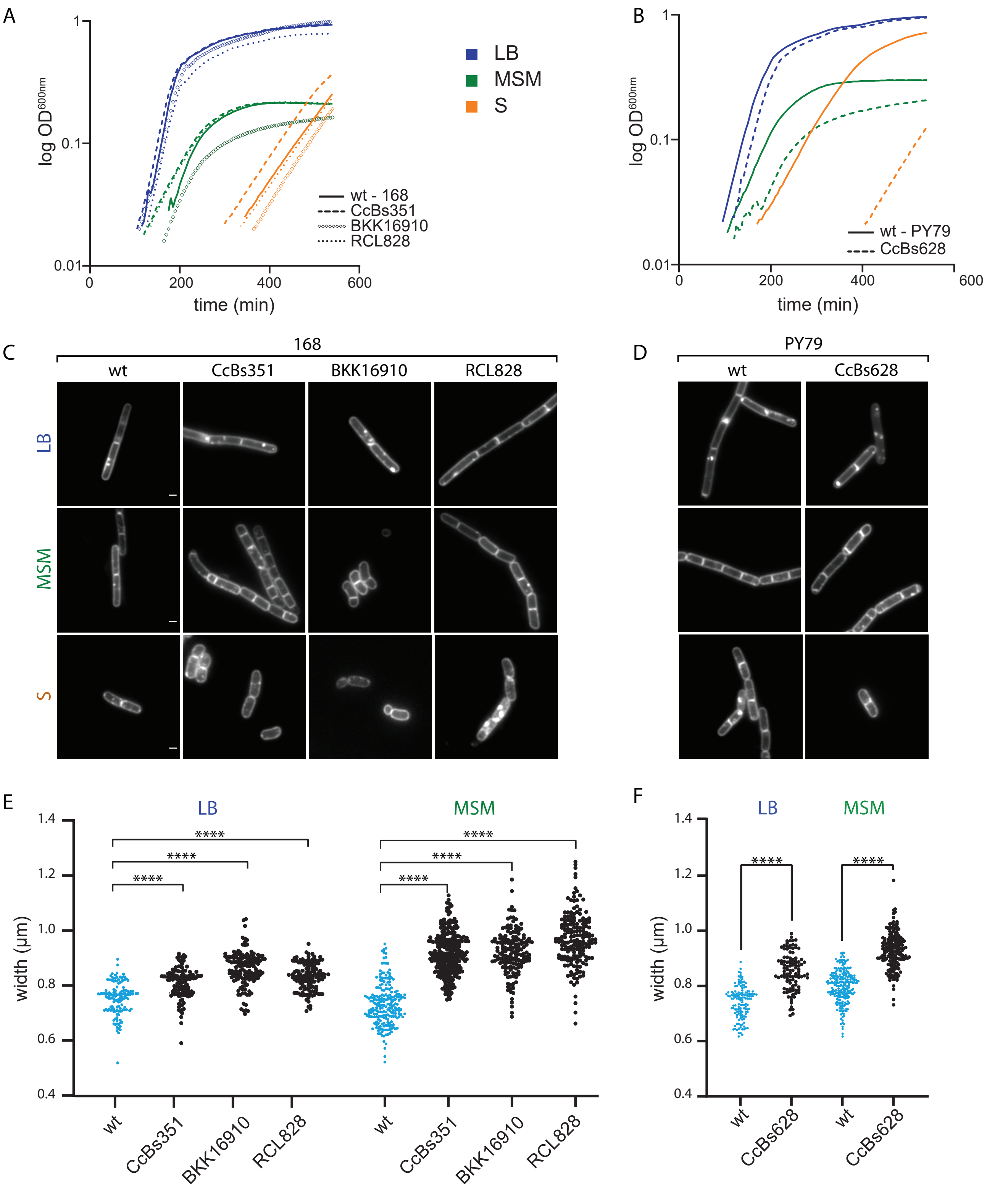


**Fig. Sup. 3.** Growth and cell shape of *B. subtilis rodZ* mutants vary depending on the growth media and the genetic background. **A-B**. Typical growth curves of wild type and Δ*rodZ* mutants of *B. subtilis* in rich LB (blue), poor MSM (green) and S (orange) media. Strains are derivative of the 168 (A) or PY79 (B) wild type strains, and either wild type (plain), or deleted for *rodZ* (dashed; CcBs351 or CcBs628). Panel A displays the growth of two additional Δ*rodZ,* the one from the BKK library (circles; BKK16910) and the BKK Δ*rodZ* mutant backcrossed into the 168 wild type strain (dotted; RCL828).
**C-D**. Epifluorescence images of *B. subtilis* cells grown to mid exponential growth phase, stained with the FM1-43FX membrane dye. Strains are derivative of the 168 (C) or PY79 (D) wild type strains, and either wild type ('wt'), carrying our *rodZ* deletion (CcBs351; CcBs628), *rodZ* - from the BKK library (BKK16910), or *rodZ* - from the BKK backcrossed into 168 wild type (RCL828). Scale bars: 1 µm.
**E-F**. Cell width distributions of the *rodZ* mutant (black) and its parental wild type 168 (E) or PY79 (F) strains (blue), grown to mid exponential phase in LB and MSM media. Data are compilations of two independent experiments. Statistical significance of the comparison between each mutants and the wild type strain was estimated using the Mann-Whitney non-parametric test (**** = P-value <0.0001).

Fig. Sup. 4

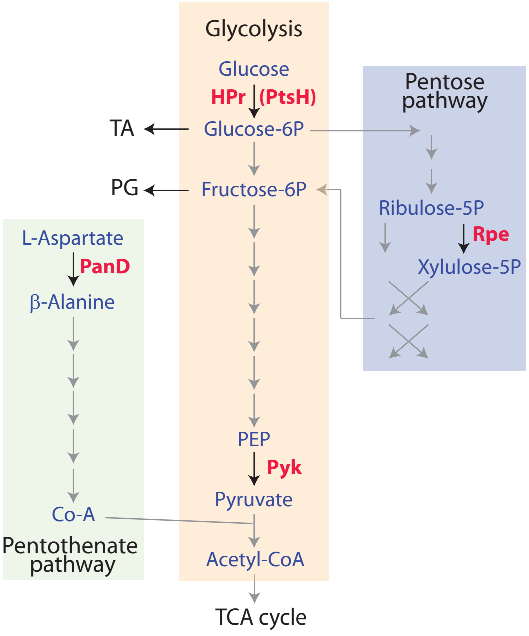


Fig. Sup. 4. Carbon metabolic pathways involving the selected mutants deficient for cell width control. The four genes selected in our screen that are involved in carbon metabolism, *panD, ptsH, pyk* and *rpe*, encode enzymes required for the pentothenate, glycolysis and pentose pathway respectively. PanD converts L-asp into β-alanine, the first step of the pathway leading to Coenzyme A (Co-A) synthesis. Co-A is used in glycolysis as a substrate for Pyk to form acetyl-CoA. HPr (encoded by *ptsH*) is a bi-functional protein acting also in glycolysis as a part of the PTS (phosphotransferase system) required for the import/phosphorylation of sugars, and in the regulation of the carbon catabolite control, as an allosteric regulator with CcpA. Rpe is predicted to produce xylulose-5P in the pentose pathway.

Fig. Sup. 5.

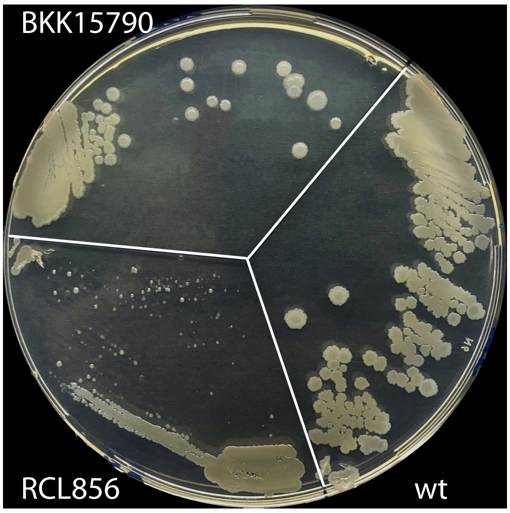


Fig. Sup. 5. Backcross of the *rpe* deletion reveals a strong growth defect.
Chromosomal DNA of strain BKK15790, knockout for *rpe*, was transformed into the wild type *B. subtilis* 168 strain (wt) to generate strain RCL856. The RCL856 strain displays a 'small-colony' phenotype indicative of a growth defect. Isolated colonies of each strain were streaked on an LB plate and grown for 24h at 37°C.

Fig. Sup. 6.

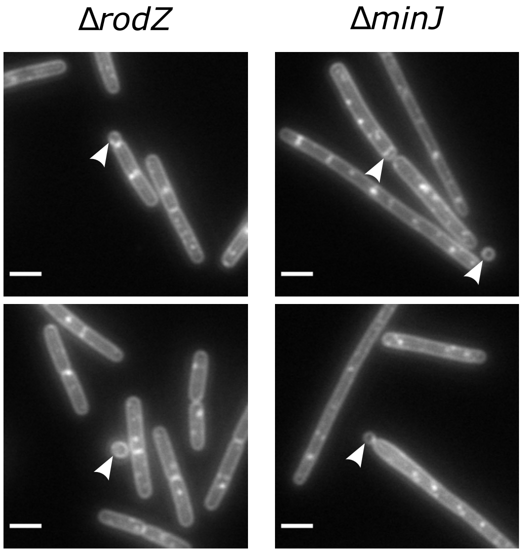


Fig. Sup. 6. The *rodZ* and *minJ* mutants form minicells. Display are images of the Δ*rodZ* (strain RCL828) and Δ*minJ* (RLC834) mutants grown to mid exponential phase in LB medium, stained with FM1-43FX and fixed, imaged by epifluorescence microscopy. Arrowheads point to minicells. Scale bar: 2 µm.

Fig. Sup. 7.

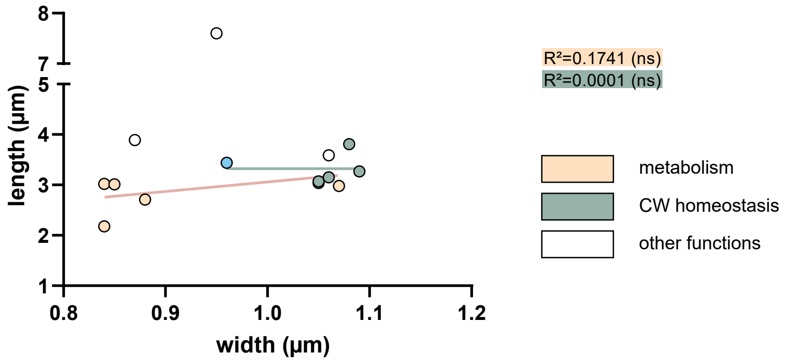


Fig. Sup. 7. Average cell length as a function of average cell width for each mutant. The wild type strain is labelled in light blue. $R^2$ of the linear regressions (lines) are indicated on the panel. Data are compilations of at least three independent experiments.

Sup. Table 1. Genes reported to affect cell width in *B. subtilis*

| gene | fonction | link with CW | essentiality | effect[1] | references [2] | this study[3] |
|---|---|---|---|---|---|---|
| CW homeostasis | | | | | | |
| *mreB* | rod complex regulation | | no | + | Jones *et al.,* 01 | na |
| *mbl* | rod complex regulation | | no | + | Abhayawardhane *et al.,* 95, Jones *et al.,* 01 | na |
| *mreBH* | rod complex regulation | | no | - | Carballido-López *et al.* , 06; Sassine *et al.* , 20 | no |
| *mreC* | rod complex transpeptidase regulator | | no | R | Lee *et al.,* 03; Leaver *et al.,* 05 | na |
| *mreD* | rod complex, unknown function | | yes | R | Leaver *et al.,* 05 | na |
| *pbpA+pbpH* | rod complex transpeptidases (bPBPs) | | yes[4] | R | Wei *et al.,* 03 | na |
| *rodA* | rod complex transglycosylase | | yes | R | Henriques *et al.,* 98 | na |
| *rodZ* | rod complex, regulator | | no[5] | + | Muchova *et al.,* 13; Van Beilen *et al.* , 16 | yes |
| *murB* | PG precursor synthetic pathway | | yes | + | Peters *et al.,* 16 | na |
| *ponA* | transglycosylase/transpeptidase (aPBP) | | no | - | Popham *et al.,* 96; Claessen *et al.,* 08 | yes |
| *lytE* | PG-hydrolase | | no | ?[6] | Dominguez-Cuevas *et al.,* 13; Carballido-López *et al.* , 06; Sassine *et al.* 20 | no |
| *cwlO* | PG-hydrolase | | no | + | Dominguez-Cuevas *et al.,* 13; Meisner *et al.,* 13 | yes |
| *ftsE* | ABC transporter, activator of CwlO | | no | + | Dominguez-Cuevas *et al.,* 13; Meisner *et al.,* 13 | yes |
| *ftsX* | ABC transporter, activator of CwlO | | no | + | Dominguez-Cuevas *et al.,* 13; Meisner *et al.,* 13 | yes |
| *tagT+TagV* | Teichoic acid synthesis | | no | + | Kawai *et al* ., 11 | na |
| *tagO* | TA synthesis (first step) | | no[8] | R | D'Elia *et al.,* 06 | na |
| *tagA* | TA synthesis (first commited step) | | no[8] | R | D'Elia *et al.,* 09 | na |
| *ltaS (yflE)* | lipoteichoic acid synthase | | no | -[9] | Sassine *et al.* , 20 | na |
| *yqgS* | lipoteichoic acid synthase | | no | -[9] | Sassine *et al.* , 20 | na |
| *yfnI (ltaSA)* | lipoteichoic acid synthase | | no | -[9] | Sassine *et al.* , 20 | na |
| Other processes | | link with CW | | | | |
| *glmR (yvcK)* | regulation of C flux | stimulates PG precursor synthetic pathway | yes[7] | +[7] | Foulquier *et al.,* 11 | na |
| *rny (ymdA)* | Rnase Y | affect *rodA* , *mreBCD* and *mreBH* expression | no | - | Figaro *et al.,* 13 | no |
| *rnJA* | Rnase J1 | affect *rodA* , *mreBCD* and *mreBH* expression | no | + | Figaro *et al.,* 13 | na |
| *cpgA* | ribosome assembly & detoxification of erythronate-4P | detoxification prevents downstream depletion of CW precursors | no | +[9] | Cladiere *et al.* , 06 | na |
| *ezrA+gpsB* | FtsZ inhibitor and late divisome protein | *ponA* localization | no | - | Claessen *et al.,* 08 | na |

1: + or - impact on cell width upon gene deletion or depletion; R stands for round cell, a consequence of the absence of elongation
2: references for measurement of width
3: mutants selected in the present screen; na (not available) indicates that the mutant could not be be find in this screen du to its absence from the BKK library, its essentiality or because the phenotype is synthetic
4: synthetically lethal
5: essential in Muchova *et al.,* 13 but inactivated in *B* . *subtilis* in Van Beilen *et al.,* 16, Koo *et al.* , 17, Kobayashi *et al.,* 03, and this study
6: thinner in Dominguez *et al., 20* 13, Carballido *et al., 20* 06, and Sassine, 2020 but wild type width in Meisner *et al.* , 13
7: only during neoglucogenic condition
8: erroneously annotated as essential in the subtiwiki database (and therefore absent from the BKK library)

Sup. Table 2. Settings used for the MicrobeJ plugin

| | |
|---|---|
| Particle selection parameters | |
| Area [µm²]: | 3.7-20 |
| Length [µm]: | 0-max |
| Width [µm]: | 0.5-2 |
| - Range [µm]: | 0-max |
| - Variation: | 0-0.15 |
| Circularity: | 0.3-0.7 |
| Curvature: | 0-max |
| Sinuosity: | 0-max |
| Angularity [rad]: | 0-0.25 |
| Solidity: | 0.85-max |
| Intensity: | 0-max |
| Segmentation method | |
| Dark, Otsu, median axis | |
| Treatment | |
| Thresholding: | Use ROI (enabled) |
| Resampling: | Resolution: 0.5, Method: Bilinear |
| Other: | Include Holes (enabled) |
| Pre-Processing (enabled): | Subtract Background, rolling=12 |
| Threshold Calculator (enabled): | Binary, Dark, Otsu, offset: 0, area: 1000-max |

### Sup. Table 3. Average width differences (%) across replicates[1]

| | WT 1 | WT 2 | WT 3 | WT 4 | WT 5 | WT 6 |
|---|---|---|---|---|---|---|
| WT 1 | - | +1.7 % | +0.8 % | +0.5 % | +0.2 % | +0.1 % |
| WT 2 | - | - | -0.9 % | -1.2 % | -1.5 % | -1.6 % |
| WT 3 | - | - | - | -0.3 % | -0.6 % | -0.7 % |
| WT 4 | - | - | - | - | -0.4 % | -0.4 % |
| WT 5 | - | - | - | - | - | -0.1 % |
| width (µm) | 1.01 | 1.028 | 1.012 | 1.016 | 1.013 | 1.012 |
| +/- | 0.076 | 0.080 | 0.071 | 0.076 | 0.089 | 0.071 |

1: ordinary one-way ANOVA  test between replicates conclude to no significant differences

### Sup. Table 4. Cell width of mutants of the BKK collection

| BKK name [1] | gene | screening delta$^2$ (%) | erage width (µ | +/- | nb | ADP |
|---|---|---|---|---|---|---|
| **BKK34800** | *cwlO* | 23,357 | 1,418 | 0,136 | 301 | 1,149 |
| **BKK12760** | *xkdW* | 21,842 | 1,421 | 0,125 | 150 | 1,166 |
| **BKK35260** | *ftsE* | 19,51 | 1,383 | 0,153 | 109 | 1,158 |
| **BKK22849** | *ypzH* | 15,994 | 1,276 | 0,133 | 401 | 1,100 |
| **BKK00100** | *dacA* | 15,94 | 1,342 | 0,149 | 107 | 1,158 |
| **BKK35250** | *ftsX* | 15,586 | 1,329 | 0,183 | 286 | 1,149 |
| **BKK20680** | *yoqC* | 13,238 | 1,323 | 0,180 | 411 | 1,168 |
| **BKK16825** | *ymfD* | 12,668 | 1,309 | 0,172 | 82 | 1,162 |
| **BKK27320** | *greA* | 12,654 | 1,098 | 0,100 | 40 | 0,975 |
| **BKK35450** | *comFC* | 11,474 | 1,242 | 0,102 | 90 | 1,114 |
| **BKK16910** | *rodZ* | 11,399 | 1,294 | 0,108 | 287 | 1,162 |
| **BKK31070** | *yuaC* | 11,327 | 1,293 | 0,130 | 238 | 1,161 |
| **BKK12770** | *xkdX* | 10,489 | 1,289 | 0,121 | 135 | 1,166 |
| **BKK00030** | *yaaA* | 9,941 | 1,283 | 0,122 | 189 | 1,167 |
| **BKK17000** | *kbl* | 9,825 | 1,276 | 0,164 | 152 | 1,162 |
| **BKK35440** | *yvyF* | 9,578 | 1,221 | 0,111 | 161 | 1,114 |
| **BKK23990** | *yqiW* | 9,495 | 1,263 | 0,141 | 142 | 1,153 |
| **BKK35220** | *minJ* | 9,381 | 1,242 | 0,131 | 104 | 1,135 |
| **BKK13030** | *ykhA* | 9,169 | 1,273 | 0,183 | 109 | 1,166 |
| **BKK02750** | *natA* | 9,096 | 1,311 | 0,124 | 127 | 1,201 |
| BKK19550 | *yodC* | 9,083 | 1,232 | 0,110 | 425 | 1,129 |
| BKK18230 | *yngG* | 8,914 | 1,276 | 0,084 | 255 | 1,172 |
| BKK36280 | *ywqA* | 8,891 | 1,213 | 0,146 | 347 | 1,114 |
| BKK15450 | *lspA* | 8,741 | 1,210 | 0,108 | 210 | 1,113 |
| BKK28800 | *araA* | 8,593 | 1,209 | 0,100 | 31 | 1,113 |
| BKK15730 | *fmt* | 8,544 | 1,058 | 0,086 | 72 | 0,975 |
| BKK08690 | *ygaD* | 8,54 | 1,312 | 0,134 | 34 | 1,208 |
| BKK05344 | *ydzS* | 8,47 | 1,266 | 0,099 | 161 | 1,167 |
| BKK35520 | *yvhJ* | 8,434 | 1,246 | 0,130 | 196 | 1,149 |
| BKK16660 | *truB* | 8,397 | 1,261 | 0,121 | 75 | 1,164 |
| BKK34110 | *rsbP* | 8,384 | 1,230 | 0,109 | 124 | 1,135 |
| BKK36130 | *ywrA* | 8,371 | 1,057 | 0,089 | 33 | 0,975 |
| BKK19440 | *norM* | 8,352 | 1,224 | 0,130 | 613 | 1,129 |
| BKK23520 | *fur* | 8,293 | 1,249 | 0,140 | 93 | 1,153 |
| BKK36269 | *ywzD* | 8,241 | 1,252 | 0,112 | 101 | 1,157 |
| BKK29280 | *ytnM* | 8,103 | 1,203 | 0,118 | 155 | 1,113 |
| BKK12619 | *ykzL* | 7,983 | 1,282 | 0,140 | 81 | 1,187 |
| BKK14750 | *ylaE* | 7,862 | 1,272 | 0,183 | 72 | 1,179 |
| BKK09840 | *hemZ* | 7,816 | 1,269 | 0,127 | 140 | 1,177 |
| BKK27290 | *yrzA* | 7,775 | 1,268 | 0,128 | 45 | 1,177 |
| BKK34760 | *yvcK* | 7,743 | 1,223 | 0,105 | 271 | 1,135 |
| BKK12850 | *ykaA* | 7,694 | 1,252 | 0,130 | 271 | 1,163 |
| BKK19370 | *odhA* | 7,657 | 1,216 | 0,124 | 315 | 1,129 |
| BKK31580 | *maeN* | 7,642 | 1,236 | 0,099 | 103 | 1,148 |
| BKK13280 | *ykoJ* | 7,628 | 1,255 | 0,166 | 168 | 1,166 |
| BKK31570 | *yufQ* | 7,616 | 1,315 | 0,129 | 119 | 1,222 |
| BKK38229 | *slrA* | 7,495 | 1,203 | 0,112 | 144 | 1,119 |
| BKK23250 | *ribH* | 7,444 | 1,275 | 0,102 | 339 | 1,187 |
| BKK18320 | *ppsC* | 7,406 | 1,258 | 0,113 | 91 | 1,172 |
| BKK04410 | *ydbB* | 7,404 | 1,251 | 0,109 | 372 | 1,165 |
| BKK01950 | *skfE* | 7,358 | 1,222 | 0,097 | 31 | 1,138 |
| BKK00330 | *yabA* | 7,321 | 1,265 | 0,127 | 430 | 1,179 |
| BKK18150 | *xynC* | 7,264 | 1,257 | 0,112 | 231 | 1,172 |
| BKK18130 | *eglS* | 7,223 | 1,256 | 0,116 | 231 | 1,172 |
| BKK33710 | *opuBC* | 7,139 | 1,232 | 0,108 | 486 | 1,149 |
| BKK39020 | *yxjA* | 7,131 | 1,293 | 0,161 | 103 | 1,206 |
| BKK14570 | *ykyA* | 7,126 | 1,263 | 0,180 | 32 | 1,179 |
| BKK29730 | *ytxD* | 7,116 | 1,192 | 0,093 | 30 | 1,113 |
| BKK30050 | *ytgP* | 7,092 | 1,309 | 0,213 | 112 | 1,222 |
| BKK22770 | *mtrB* | 7,077 | 1,235 | 0,143 | 232 | 1,153 |
| BKK24930 | *yqzD* | 7,065 | 1,246 | 0,104 | 415 | 1,164 |
| BKK01700 | *murQ* | 7,045 | 1,218 | 0,146 | 34 | 1,138 |
| BKK12672 | *xkzB* | 7,034 | 1,273 | 0,176 | 40 | 1,189 |
| BKK26720 | *azlB* | 6,983 | 1,259 | 0,109 | 392 | 1,177 |
| BKK22620 | *hisC* | 6,963 | 1,177 | 0,121 | 304 | 1,100 |
| BKK05330 | *aseR* | 6,954 | 1,248 | 0,068 | 122 | 1,167 |
| BKK19510 | *yojB* | 6,876 | 1,242 | 0,141 | 243 | 1,162 |
| BKK03040 | *amyE* | 6,864 | 1,208 | 0,137 | 60 | 1,130 |
| BKK03651 | *bsdD* | 6,852 | 1,245 | 0,102 | 150 | 1,165 |
| BKK34130 | *ganA* | 6,791 | 1,212 | 0,115 | 30 | 1,135 |
| BKK02610 | *ycbR* | 6,782 | 1,207 | 0,128 | 56 | 1,130 |
| BKK04770 | *ydcH* | 6,741 | 1,243 | 0,138 | 298 | 1,165 |
| BKK22530 | *ypjA* | 6,735 | 1,216 | 0,103 | 76 | 1,139 |
| BKK28450 | *sdhC* | 6,734 | 1,238 | 0,097 | 32 | 1,160 |
| BKK31370 | *yugJ* | 6,723 | 1,225 | 0,095 | 618 | 1,148 |
| BKK34560 | *malL* | 6,715 | 1,211 | 0,142 | 116 | 1,135 |
| BKK39360 | *hutU* | 6,694 | 1,244 | 0,152 | 105 | 1,166 |
| BKK04060 | *ycsG* | 6,653 | 1,242 | 0,082 | 146 | 1,165 |
| BKK35150 | *yvzB* | 6,63 | 1,210 | 0,141 | 62 | 1,135 |

| BKK name [1] | gene | screening delta$^2$ (%) | erage width (µ | +/- | nb | ADP |
|---|---|---|---|---|---|---|
| BKK19390 | *yojN* | 6,61 | 1,204 | 0,116 | 324 | 1,129 |
| BKK36380 | *rapD* | 6,61 | 1,188 | 0,116 | 769 | 1,114 |
| BKK10100 | *yhgC* | 6,591 | 1,255 | 0,115 | 206 | 1,177 |
| BKK37680 | *ywfH* | 6,58 | 1,192 | 0,099 | 116 | 1,119 |
| BKK25250 | *ccpN* | 6,574 | 1,240 | 0,106 | 98 | 1,164 |
| BKK19530 | *yodA* | 6,542 | 1,203 | 0,155 | 502 | 1,129 |
| BKK36390 | *flhP* | 6,537 | 1,187 | 0,123 | 160 | 1,114 |
| BKK36860 | *atpE* | 6,487 | 1,222 | 0,128 | 253 | 1,147 |
| BKK25840 | *phrE* | 6,454 | 1,038 | 0,095 | 43 | 0,975 |
| BKK23160 | *rluB* | 6,451 | 1,264 | 0,105 | 212 | 1,187 |
| BKK13840 | *stoA* | 6,443 | 1,242 | 0,123 | 39 | 1,166 |
| BKK31150 | *uppP* | 6,429 | 1,236 | 0,110 | 286 | 1,161 |
| BKK11590 | *yjbL* | 6,402 | 1,265 | 0,169 | 516 | 1,189 |
| BKK26590 | *blt* | 6,37 | 1,300 | 0,088 | 72 | 1,222 |
| BKK14090 | *ykuI* | 6,306 | 1,240 | 0,133 | 387 | 1,166 |
| BKK29400 | *ytlI* | 6,306 | 1,235 | 0,108 | 257 | 1,161 |
| BKK10210 | *yhfF* | 6,304 | 1,251 | 0,109 | 116 | 1,177 |
| BKK05450 | *ydfK* | 6,302 | 1,241 | 0,116 | 146 | 1,167 |
| BKK21750 | *scuA* | 6,3 | 1,226 | 0,142 | 215 | 1,153 |
| BKK15570 | *cysH* | 6,219 | 1,219 | 0,099 | 35 | 1,148 |
| BKK15270 | *sbp* | 6,206 | 1,252 | 0,143 | 140 | 1,179 |
| BKK32080 | *yuiB* | 6,197 | 1,219 | 0,079 | 152 | 1,148 |
| BKK00610 | *yabQ* | 6,179 | 1,244 | 0,108 | 94 | 1,172 |
| BKK13140 | *ohrA* | 6,162 | 1,234 | 0,146 | 212 | 1,163 |
| BKK37350 | *sboA* | 6,153 | 1,188 | 0,110 | 116 | 1,119 |
| BKK40290 | *yycN* | 6,108 | 1,221 | 0,122 | 299 | 1,151 |
| BKK24220 | *spo0A* | 6,107 | 1,257 | 0,101 | 161 | 1,184 |
| BKK15590 | *sat* | 6,105 | 1,239 | 0,130 | 241 | 1,168 |
| BKK09490 | *yhdJ* | 6,083 | 1,249 | 0,121 | 86 | 1,177 |
| BKK23030 | *ypbB* | 6,074 | 1,221 | 0,124 | 294 | 1,151 |
| BKK09620 | *yhdW* | 6,056 | 1,248 | 0,117 | 148 | 1,177 |
| BKK22990 | *ypbF* | 6,034 | 1,167 | 0,099 | 570 | 1,100 |
| BKK05650 | *ydgH* | 6,024 | 1,238 | 0,126 | 149 | 1,167 |
| BKK18400 | *yoeD* | 6,008 | 1,242 | 0,087 | 327 | 1,172 |
| BKK18250 | *yngI* | 5,923 | 1,241 | 0,117 | 178 | 1,172 |
| BKK38480 | *ywaC* | 5,912 | 1,185 | 0,127 | 30 | 1,119 |
| BKK05390 | *ydfF* | 5,893 | 1,236 | 0,090 | 391 | 1,167 |
| BKK04460 | *dctR* | 5,887 | 1,233 | 0,128 | 392 | 1,165 |
| BKK24720 | *comGB* | 5,883 | 1,232 | 0,124 | 251 | 1,164 |
| BKK23280 | *ribD* | 5,871 | 1,257 | 0,113 | 218 | 1,187 |
| BKK15780 | *cpgA* | 5,86 | 1,032 | 0,156 | 41 | 0,975 |
| BKK32100 | *yumB* | 5,847 | 1,215 | 0,095 | 63 | 1,148 |
| BKK17890 | *tkt* | 5,816 | 1,229 | 0,120 | 550 | 1,162 |
| BKK34730 | *yvcN* | 5,799 | 1,201 | 0,124 | 592 | 1,135 |
| BKK25700 | *yqeF* | 5,798 | 1,220 | 0,125 | 367 | 1,153 |
| BKK40980 | *yyaB* | 5,775 | 1,253 | 0,116 | 97 | 1,184 |
| BKK22660 | *trpC* | 5,766 | 1,217 | 0,127 | 348 | 1,151 |
| BKK20650 | *yoqF* | 5,748 | 1,221 | 0,130 | 194 | 1,155 |
| BKK03820 | *yclP* | 5,726 | 1,203 | 0,122 | 89 | 1,138 |
| BKK15620 | *sirB* | 5,71 | 1,213 | 0,092 | 100 | 1,148 |
| BKK06220 | *ydjJ* | 5,698 | 1,246 | 0,111 | 285 | 1,179 |
| BKK23460 | *spoIIAB* | 5,696 | 1,275 | 0,161 | 93 | 1,206 |
| BKK13810 | *ykvS* | 5,695 | 1,229 | 0,136 | 221 | 1,163 |
| BKK23120 | *resD* | 5,674 | 1,163 | 0,099 | 350 | 1,100 |
| BKK23510 | *ripX* | 5,671 | 1,254 | 0,095 | 148 | 1,187 |
| BKK05490 | *ydfO* | 5,658 | 1,233 | 0,115 | 151 | 1,167 |
| BKK29870 | *ytzB* | 5,644 | 1,030 | 0,079 | 100 | 0,975 |
| BKK35820 | *gerBC* | 5,632 | 1,214 | 0,107 | 408 | 1,149 |
| BKK31500 | *yuxK* | 5,625 | 1,212 | 0,147 | 56 | 1,148 |
| BKK40810 | *yyaM* | 5,597 | 1,233 | 0,148 | 144 | 1,167 |
| BKK00050 | *yaaB* | 5,574 | 1,245 | 0,126 | 178 | 1,179 |
| BKK25650 | *yqeI* | 5,559 | 1,228 | 0,099 | 346 | 1,164 |
| BKK08820 | *katA* | 5,529 | 1,241 | 0,124 | 162 | 1,177 |
| BKK11300 | *med* | 5,512 | 1,249 | 0,132 | 67 | 1,183 |
| BKK30740 | *mntD* | 5,498 | 1,214 | 0,123 | 218 | 1,151 |
| BKK35670 | *gtaB* | 5,489 | 1,175 | 0,131 | 218 | 1,114 |
| BKK26060 | *yqbM* | 5,483 | 1,241 | 0,104 | 292 | 1,177 |
| BKK34910 | *hisD* | 5,475 | 1,197 | 0,092 | 377 | 1,135 |
| BKK35570 | *tuaE* | 5,475 | 1,212 | 0,132 | 504 | 1,149 |
| BKK37460 | *rapF* | 5,469 | 1,180 | 0,131 | 258 | 1,119 |
| BKK10300 | *aprE* | 5,445 | 1,248 | 0,141 | 117 | 1,183 |
| BKK03490 | *srfAB* | 5,444 | 1,028 | 0,085 | 80 | 0,975 |
| BKK03290 | *nasE* | 5,431 | 1,191 | 0,145 | 37 | 1,130 |
| BKK22700 | *aroB* | 5,422 | 1,249 | 0,114 | 132 | 1,184 |
| BKK02670 | *lmrB* | 5,417 | 1,200 | 0,124 | 52 | 1,138 |
| BKK21030 | *yonP* | 5,411 | 1,216 | 0,117 | 398 | 1,153 |
| BKK27480 | *yrrC* | 5,403 | 1,222 | 0,105 | 146 | 1,160 |
| BKK04130 | *yczJ* | 5,388 | 1,228 | 0,130 | 143 | 1,165 |
| BKK22630 | *trpA* | 5,379 | 1,159 | 0,107 | 118 | 1,100 |

| BKK name [1] | gene | screening delta$^2$ (%) | erage width (µ | +/- | nb | ADP |
|---|---|---|---|---|---|---|
| BKK39260 | *bglH* | 5,364 | 1,227 | 0,120 | 79 | 1,165 |
| BKK34870 | *hisF* | 5,362 | 1,196 | 0,100 | 246 | 1,135 |
| BKK40579 | *yyzL* | 5,346 | 1,229 | 0,128 | 85 | 1,166 |
| BKK29410 | *ytkL* | 5,34 | 1,223 | 0,128 | 410 | 1,161 |
| BKK40260 | *yycQ* | 5,338 | 1,213 | 0,115 | 203 | 1,151 |
| BKK09340 | *yhdA* | 5,327 | 1,240 | 0,101 | 102 | 1,177 |
| BKK33630 | *secG* | 5,322 | 1,211 | 0,090 | 354 | 1,149 |
| BKK04070 | *ycsI* | 5,301 | 1,227 | 0,100 | 92 | 1,165 |
| BKK02960 | *yceJ* | 5,29 | 1,265 | 0,148 | 47 | 1,201 |
| BKK11580 | *yjbK* | 5,282 | 1,252 | 0,132 | 921 | 1,189 |
| BKK10950 | *yitD* | 5,271 | 1,246 | 0,123 | 70 | 1,183 |
| BKK09990 | *scoC* | 5,267 | 1,239 | 0,105 | 67 | 1,177 |
| BKK19540 | *yodB* | 5,241 | 1,188 | 0,117 | 565 | 1,129 |
| BKK04450 | *dctS* | 5,231 | 1,264 | 0,094 | 65 | 1,201 |
| BKK25380 | *yqfA* | 5,211 | 1,224 | 0,105 | 148 | 1,164 |
| BKK32820 | *fadE* | 5,205 | 1,227 | 0,114 | 96 | 1,166 |
| BKK14810 | *ylaK* | 5,201 | 1,240 | 0,071 | 86 | 1,179 |
| BKK12671 | *xkdN* | 5,181 | 1,251 | 0,174 | 77 | 1,189 |
| BKK33130 | *liaI* | 5,172 | 1,193 | 0,118 | 62 | 1,134 |
| BKK03410 | *bglC* | 5,146 | 1,188 | 0,133 | 38 | 1,130 |
| BKK37110 | *ywjH* | 5,144 | 1,176 | 0,141 | 115 | 1,119 |
| BKK19669 | *yoyF* | 5,14 | 1,187 | 0,133 | 315 | 1,129 |
| BKK02880 | *yceB* | 5,132 | 1,188 | 0,130 | 67 | 1,130 |
| BKK04930 | *yddD* | 5,113 | 1,217 | 0,092 | 96 | 1,158 |
| BKK27490 | *yrrB* | 5,109 | 1,219 | 0,110 | 148 | 1,160 |
| BKK39810 | *csbC* | 5,096 | 1,208 | 0,114 | 422 | 1,149 |
| BKK01830 | *ndhF* | 5,086 | 1,219 | 0,107 | 274 | 1,160 |
| BKK05720 | *ydhE* | 5,085 | 1,227 | 0,081 | 366 | 1,167 |
| BKK14510 | *ktrC* | 5,078 | 1,239 | 0,097 | 359 | 1,179 |
| BKK19520 | *yojA* | 5,07 | 1,187 | 0,116 | 686 | 1,129 |
| BKK00170 | *yaaH* | 5,06 | 1,218 | 0,124 | 606 | 1,160 |
| BKK18160 | *xynD* | 5,052 | 1,231 | 0,115 | 249 | 1,172 |
| BKK05350 | *ydfB* | 5,036 | 1,226 | 0,075 | 179 | 1,167 |
| BKK16990 | *tdh* | 5,036 | 1,222 | 0,105 | 110 | 1,164 |
| BKK38440 | *ywaF* | 5,015 | 1,175 | 0,125 | 32 | 1,119 |
| BKK29890 | *ytnP* | 5,002 | 1,169 | 0,121 | 39 | 1,113 |
| BKK11630 | *prpE* | 4,999 | 1,248 | 0,140 | 803 | 1,189 |
| BKK23402 | *spoVAEB* | 4,99 | 1,267 | 0,187 | 36 | 1,206 |
| BKK14500 | *ykqA* | 4,976 | 1,238 | 0,116 | 301 | 1,179 |
| BKK16860 | *ymfH* | 4,967 | 1,221 | 0,127 | 82 | 1,164 |
| BKK22740 | *hepT* | 4,961 | 1,155 | 0,101 | 110 | 1,100 |
| BKK37170 | *acdA* | 4,938 | 1,224 | 0,116 | 89 | 1,166 |
| BKK01800 | *alkA* | 4,929 | 1,217 | 0,104 | 324 | 1,160 |
| BKK09370 | *lytF* | 4,921 | 1,268 | 0,136 | 47 | 1,208 |
| BKK17440 | *ynbB* | 4,909 | 1,221 | 0,114 | 160 | 1,164 |
| BKK06630 | *yerH* | 4,908 | 1,260 | 0,073 | 89 | 1,201 |
| BKK05130 | *ydeB* | 4,893 | 1,224 | 0,112 | 366 | 1,167 |
| BKK28320 | *ysnD* | 4,891 | 1,216 | 0,103 | 447 | 1,160 |
| BKK30170 | *ytcP* | 4,889 | 1,282 | 0,140 | 33 | 1,222 |
| BKK26700 | *azlD* | 4,888 | 1,282 | 0,078 | 201 | 1,222 |
| BKK34880 | *hisA* | 4,887 | 1,191 | 0,086 | 270 | 1,135 |
| BKK24450 | *efp* | 4,884 | 1,245 | 0,108 | 402 | 1,187 |
| BKK29510 | *ytfI* | 4,882 | 1,167 | 0,110 | 102 | 1,113 |
| BKK31740 | *yuxH* | 4,882 | 1,204 | 0,092 | 121 | 1,148 |
| BKK35020 | *nagBA* | 4,879 | 1,190 | 0,123 | 126 | 1,135 |
| BKK35679 | *yvzH* | 4,867 | 1,205 | 0,122 | 239 | 1,149 |
| BKK11700 | *thiF* | 4,86 | 1,247 | 0,145 | 929 | 1,189 |
| BKK11990 | *yjdB* | 4,847 | 1,247 | 0,180 | 70 | 1,189 |
| BKK28750 | *araN* | 4,846 | 1,281 | 0,078 | 69 | 1,222 |
| BKK03810 | *yclO* | 4,844 | 1,260 | 0,073 | 113 | 1,201 |
| BKK22040 | *ypbQ* | 4,843 | 1,154 | 0,147 | 35 | 1,100 |
| BKK21040 | *yonO* | 4,833 | 1,209 | 0,108 | 308 | 1,153 |
| BKK18430 | *yogA* | 4,827 | 1,228 | 0,086 | 84 | 1,172 |
| BKK09900 | *natB* | 4,82 | 1,234 | 0,091 | 98 | 1,177 |
| BKK08920 | *yhbB* | 4,802 | 1,218 | 0,120 | 207 | 1,163 |
| BKK09965 | *yhaJ* | 4,797 | 1,233 | 0,131 | 93 | 1,177 |
| BKK35390 | *yviE* | 4,79 | 1,189 | 0,123 | 430 | 1,135 |
| BKK24510 | *yqhO* | 4,783 | 1,206 | 0,112 | 311 | 1,151 |
| BKK40000 | *yxnA* | 4,779 | 1,202 | 0,121 | 256 | 1,147 |
| BKK00320 | *yaaT* | 4,777 | 1,235 | 0,114 | 214 | 1,179 |
| BKK29500 | *ytfJ* | 4,769 | 1,166 | 0,100 | 181 | 1,113 |
| BKK16120 | *topA* | 4,768 | 1,241 | 0,126 | 110 | 1,184 |
| BKK34400 | *padC* | 4,765 | 1,189 | 0,096 | 68 | 1,135 |
| BKK24680 | *comGF* | 4,754 | 1,208 | 0,140 | 178 | 1,153 |
| BKK12090 | *cotT* | 4,749 | 1,245 | 0,203 | 112 | 1,189 |
| BKK24500 | *yqhP* | 4,746 | 1,206 | 0,106 | 404 | 1,151 |
| BKK33670 | *rghRB* | 4,743 | 1,188 | 0,139 | 33 | 1,134 |
| BKK03050 | *ldh* | 4,742 | 1,184 | 0,127 | 46 | 1,130 |
| BKK21720 | *ypmT* | 4,739 | 1,206 | 0,132 | 237 | 1,151 |

| BKK name [1] | gene | screening delta$^2$ (%) | erage width (µ | +/- | nb | ADP |
|---|---|---|---|---|---|---|
| BKK10690 | *gerPD* | 4,734 | 1,239 | 0,107 | 93 | 1,183 |
| BKK27580 | *yrvJ* | 4,725 | 1,280 | 0,064 | 102 | 1,222 |
| BKK27710 | *tgt* | 4,718 | 1,214 | 0,101 | 352 | 1,160 |
| BKK29920 | *ytmP* | 4,71 | 1,166 | 0,128 | 41 | 1,113 |
| BKK20690 | *yoqB* | 4,7 | 1,209 | 0,112 | 317 | 1,155 |
| BKK26580 | *bltR* | 4,698 | 1,279 | 0,110 | 56 | 1,222 |
| BKK06076 | *ydzW* | 4,693 | 1,222 | 0,146 | 449 | 1,167 |
| BKK05408 | *ydzP* | 4,67 | 1,193 | 0,138 | 32 | 1,139 |
| BKK12380 | *uxaB* | 4,666 | 1,244 | 0,170 | 666 | 1,189 |
| BKK06200 | *ydjH* | 4,662 | 1,234 | 0,129 | 191 | 1,179 |
| BKK05640 | *ydgG* | 4,653 | 1,222 | 0,092 | 289 | 1,167 |
| BKK27930 | *spo0B* | 4,648 | 1,207 | 0,145 | 126 | 1,153 |
| BKK12600 | *xkdF* | 4,635 | 1,244 | 0,125 | 181 | 1,189 |
| BKK27400 | *yrzL* | 4,631 | 1,213 | 0,118 | 464 | 1,160 |
| BKK14830 | *glsB* | 4,626 | 1,234 | 0,082 | 201 | 1,179 |
| BKK34180 | *yvfI* | 4,619 | 1,188 | 0,101 | 230 | 1,135 |
| BKK09020 | *yhcB* | 4,618 | 1,231 | 0,109 | 97 | 1,177 |
| BKK33180 | *yvrC* | 4,613 | 1,278 | 0,109 | 77 | 1,222 |
| BKK27600 | *relA* | 4,606 | 1,213 | 0,098 | 421 | 1,160 |
| BKK38530 | *dltD* | 4,605 | 1,202 | 0,166 | 142 | 1,149 |
| BKK01845 | *ybcC* | 4,599 | 1,213 | 0,104 | 268 | 1,160 |
| BKK37320 | *narK* | 4,597 | 1,200 | 0,108 | 349 | 1,147 |
| BKK28650 | *ysgA* | 4,59 | 1,164 | 0,117 | 121 | 1,113 |
| BKK18019 | *ynzL* | 4,588 | 1,217 | 0,119 | 105 | 1,164 |
| BKK07480 | *yfmG* | 4,582 | 1,212 | 0,114 | 333 | 1,159 |
| BKK24950 | *pstBB* | 4,578 | 1,262 | 0,148 | 150 | 1,206 |
| BKK34380 | *slrR* | 4,578 | 1,187 | 0,087 | 234 | 1,135 |
| BKK04480 | *ydbI* | 4,563 | 1,256 | 0,091 | 43 | 1,201 |
| BKK28620 | *rnhC* | 4,561 | 1,164 | 0,099 | 170 | 1,113 |
| BKK30390 | *bceS* | 4,561 | 1,278 | 0,090 | 74 | 1,222 |
| BKK13450 | *sigI* | 4,549 | 1,219 | 0,136 | 43 | 1,166 |
| BKK02760 | *natB* | 4,547 | 1,256 | 0,177 | 90 | 1,201 |
| BKK37960 | *ywdH* | 4,547 | 1,200 | 0,114 | 436 | 1,147 |
| BKK20310 | *yorO* | 4,516 | 1,214 | 0,109 | 191 | 1,162 |
| BKK23260 | *ribAB* | 4,514 | 1,241 | 0,121 | 254 | 1,187 |
| BKK17430 | *ynbA* | 4,513 | 1,216 | 0,109 | 71 | 1,164 |
| BKK35410 | *flgK* | 4,507 | 1,165 | 0,107 | 1066 | 1,114 |
| BKK02980 | *opuAA* | 4,495 | 1,255 | 0,132 | 69 | 1,201 |
| BKK06180 | *pspA* | 4,493 | 1,232 | 0,117 | 192 | 1,179 |
| BKK40560 | *yybP* | 4,489 | 1,219 | 0,123 | 132 | 1,166 |
| BKK03840 | *ycnB* | 4,477 | 1,217 | 0,105 | 221 | 1,165 |
| BKK33660 | *rghRA* | 4,473 | 1,019 | 0,076 | 55 | 0,975 |
| BKK09230 | *yhcV* | 4,472 | 1,230 | 0,096 | 153 | 1,177 |
| BKK16000 | *ylqC* | 4,461 | 1,169 | 0,120 | 241 | 1,119 |
| BKK22380 | *ypmB* | 4,45 | 1,149 | 0,105 | 218 | 1,100 |
| BKK14270 | *moeB* | 4,448 | 1,232 | 0,109 | 344 | 1,179 |
| BKK31900 | *yukD* | 4,448 | 1,199 | 0,111 | 148 | 1,148 |
| BKK23680 | *yqjZ* | 4,446 | 1,202 | 0,112 | 340 | 1,151 |
| BKK20700 | *yoqA* | 4,444 | 1,206 | 0,116 | 398 | 1,155 |
| BKK31700 | *comX* | 4,443 | 1,218 | 0,102 | 223 | 1,166 |
| BKK04830 | *sacV* | 4,439 | 1,217 | 0,091 | 355 | 1,165 |
| BKK02230 | *purT* | 4,437 | 1,180 | 0,123 | 85 | 1,130 |
| BKK11670 | *thiO* | 4,437 | 1,242 | 0,153 | 883 | 1,189 |
| BKK04810 | *immA* | 4,435 | 1,217 | 0,123 | 156 | 1,165 |
| BKK40210 | *yydC* | 4,434 | 1,218 | 0,193 | 47 | 1,166 |
| BKK12080 | *ctaO* | 4,427 | 1,242 | 0,183 | 156 | 1,189 |
| BKK17010 | *ymcB* | 4,416 | 1,215 | 0,120 | 146 | 1,164 |
| BKK06340 | *yeaD* | 4,414 | 1,198 | 0,134 | 301 | 1,147 |
| BKK23270 | *ribE* | 4,409 | 1,239 | 0,103 | 363 | 1,187 |
| BKK27370 | *yrrL* | 4,406 | 1,211 | 0,105 | 221 | 1,160 |
| BKK05290 | *ydeP* | 4,398 | 1,219 | 0,084 | 214 | 1,167 |
| BKK03380 | *yckB* | 4,396 | 1,254 | 0,130 | 282 | 1,201 |
| BKK25460 | *dnaJ* | 4,382 | 1,215 | 0,126 | 211 | 1,164 |
| BKK01900 | *ybcM* | 4,373 | 1,162 | 0,112 | 35 | 1,113 |
| BKK06380 | *yebC* | 4,371 | 1,230 | 0,144 | 366 | 1,179 |
| BKK26259 | *yqzO* | 4,358 | 1,228 | 0,100 | 128 | 1,177 |
| BKK35210 | *yvkA* | 4,356 | 1,200 | 0,117 | 376 | 1,149 |
| BKK37000 | *prmC* | 4,352 | 1,163 | 0,130 | 78 | 1,114 |
| BKK38018 | *ywzG* | 4,346 | 1,217 | 0,127 | 71 | 1,166 |
| BKK28700 | *ysfE* | 4,331 | 1,161 | 0,078 | 32 | 1,113 |
| BKK38850 | *yxkC* | 4,329 | 1,215 | 0,125 | 58 | 1,165 |
| BKK10390 | *yhfW* | 4,319 | 1,234 | 0,091 | 176 | 1,183 |
| BKK03788 | *yczM* | 4,309 | 1,207 | 0,138 | 43 | 1,158 |
| BKK14590 | *pdhB* | 4,304 | 1,230 | 0,148 | 301 | 1,179 |
| BKK02780 | *ycdA* | 4,3 | 1,253 | 0,212 | 33 | 1,201 |
| BKK08860 | *ssuD* | 4,292 | 1,227 | 0,109 | 234 | 1,177 |
| BKK16740 | *spoVFB* | 4,285 | 1,215 | 0,122 | 89 | 1,164 |
| BKK20670 | *yoqD* | 4,284 | 1,204 | 0,124 | 286 | 1,155 |
| BKK34150 | *ganP* | 4,283 | 1,184 | 0,115 | 62 | 1,135 |

### Sup. Table 4: Cell width of mutants of the BKK collection (continued)

| BKK name [1] | gene | screening delta[2] (%) | erage width (µ | +/- | nb | ADP |
|---|---|---|---|---|---|---|
| BKK00340 | *yabB* | 4,272 | 1,230 | 0,152 | 79 | 1,179 |
| BKK26150 | *yqbD* | 4,272 | 1,017 | 0,091 | 96 | 0,975 |
| BKK29040 | *ytbD* | 4,268 | 1,274 | 0,076 | 114 | 1,222 |
| BKK39070 | *bglS* | 4,253 | 1,214 | 0,137 | 96 | 1,165 |
| BKK01990 | *ybdG* | 4,252 | 1,178 | 0,132 | 69 | 1,130 |
| BKK35940 | *rbsA* | 4,231 | 1,198 | 0,130 | 336 | 1,149 |
| BKK29050 | *ytbE* | 4,217 | 1,160 | 0,111 | 70 | 1,113 |
| BKK14370 | *yknZ* | 4,216 | 1,229 | 0,120 | 77 | 1,179 |
| BKK27785 | *yrzF* | 4,209 | 1,208 | 0,099 | 405 | 1,160 |
| BKK25900 | *cwlA* | 4,196 | 1,273 | 0,073 | 192 | 1,222 |
| BKK20250 | *mtbP* | 4,195 | 1,211 | 0,127 | 199 | 1,162 |
| BKK13680 | *motB* | 4,193 | 1,217 | 0,122 | 296 | 1,168 |
| BKK31230 | *tlpB* | 4,193 | 1,204 | 0,131 | 144 | 1,156 |
| BKK17330 | *miaA* | 4,187 | 1,212 | 0,128 | 109 | 1,164 |
| BKK34729 | *yvzJ* | 4,186 | 1,198 | 0,122 | 289 | 1,149 |
| BKK22000 | *sspL* | 4,185 | 1,203 | 0,115 | 238 | 1,155 |
| BKK09130 | *tcyP* | 4,177 | 1,226 | 0,117 | 77 | 1,177 |
| BKK10060 | *ecsC* | 4,17 | 1,217 | 0,120 | 300 | 1,168 |
| BKK38400 | *epr* | 4,17 | 1,195 | 0,112 | 399 | 1,147 |
| BKK15370 | *ylmD* | 4,169 | 1,187 | 0,122 | 53 | 1,139 |
| BKK25760 | *spoIVCB* | 4,165 | 1,273 | 0,102 | 232 | 1,222 |
| BKK00810 | *dusB* | 4,162 | 1,208 | 0,126 | 361 | 1,160 |
| BKK31350 | *pgi* | 4,162 | 1,196 | 0,120 | 103 | 1,148 |
| BKK12000 | *manR* | 4,158 | 1,238 | 0,171 | 444 | 1,189 |
| BKK11180 | *yitZ* | 4,147 | 1,232 | 0,116 | 118 | 1,183 |
| BKK24620 | *tasA* | 4,144 | 1,015 | 0,117 | 64 | 0,975 |
| BKK05340 | *ydfA* | 4,14 | 1,015 | 0,085 | 47 | 0,975 |
| BKK19400 | *sodC* | 4,136 | 1,176 | 0,131 | 489 | 1,129 |
| BKK05140 | *ydzE* | 4,135 | 1,208 | 0,127 | 633 | 1,160 |
| BKK20210 | *yorY* | 4,135 | 1,176 | 0,134 | 88 | 1,129 |
| BKK36010 | *alsS* | 4,13 | 1,160 | 0,114 | 47 | 1,114 |
| BKK37430 | *albG* | 4,125 | 1,197 | 0,114 | 403 | 1,149 |
| BKK12400 | *yjnA* | 4,11 | 1,238 | 0,150 | 551 | 1,189 |
| BKK19570 | *yodE* | 4,104 | 1,176 | 0,098 | 440 | 1,129 |
| BKK39600 | *yxeC* | 4,103 | 1,195 | 0,107 | 270 | 1,147 |
| BKK40529 | *yyzH* | 4,103 | 1,214 | 0,118 | 105 | 1,166 |
| BKK30000 | *ythQ* | 4,095 | 1,272 | 0,174 | 141 | 1,222 |
| BKK25110 | *yqfU* | 4,094 | 1,211 | 0,103 | 647 | 1,164 |
| BKK07770 | *yfkS* | 4,086 | 1,216 | 0,129 | 357 | 1,168 |
| BKK12190 | *yjhB* | 4,08 | 1,238 | 0,095 | 267 | 1,189 |
| BKK07860 | *yfkL* | 4,073 | 1,258 | 0,161 | 100 | 1,208 |
| BKK31100 | *ktrB* | 4,058 | 1,208 | 0,137 | 223 | 1,161 |
| BKK02350 | *nagP* | 4,056 | 1,176 | 0,128 | 139 | 1,130 |
| BKK08880 | *rpsNB* | 4,054 | 1,225 | 0,099 | 151 | 1,177 |
| BKK33210 | *yvrG* | 4,052 | 1,214 | 0,105 | 121 | 1,166 |
| BKK29070 | *ytaF* | 4,045 | 1,203 | 0,099 | 79 | 1,156 |
| BKK13819 | *ykzS* | 4,043 | 1,214 | 0,108 | 259 | 1,166 |
| BKK24970 | *pstA* | 4,033 | 1,255 | 0,156 | 51 | 1,206 |
| BKK23210 | *scpB* | 4,03 | 1,232 | 0,111 | 79 | 1,184 |
| BKK02010 | *ybdK* | 4,027 | 1,184 | 0,128 | 137 | 1,138 |
| BKK28310 | *ilvB* | 4,024 | 1,206 | 0,119 | 295 | 1,160 |
| BKK34210 | *yvfG* | 4,022 | 1,181 | 0,108 | 85 | 1,135 |
| BKK27910 | *yszB* | 4,011 | 1,206 | 0,121 | 200 | 1,160 |
| BKK12980 | *ykfB* | 4,008 | 1,257 | 0,101 | 35 | 1,208 |
| BKK23320 | *ypzC* | 4,001 | 1,235 | 0,111 | 107 | 1,187 |
| BKK26890 | *csn* | 3,998 | 1,224 | 0,108 | 318 | 1,177 |
| BKK24890 | *yqgN* | 3,985 | 1,210 | 0,099 | 280 | 1,164 |
| BKK40970 | *parA* | 3,973 | 1,232 | 0,104 | 130 | 1,184 |
| BKK32120 | *yuzG* | 3,965 | 1,193 | 0,110 | 30 | 1,148 |
| BKK10009 | *yhzF* | 3,952 | 1,223 | 0,105 | 148 | 1,177 |
| BKK07680 | *yflH* | 3,95 | 1,205 | 0,119 | 206 | 1,159 |
| BKK33830 | *opuCA* | 3,95 | 1,195 | 0,110 | 332 | 1,149 |
| BKK24560 | *gcvPA* | 3,945 | 1,234 | 0,099 | 271 | 1,187 |
| BKK30430 | *ytrD* | 3,931 | 1,207 | 0,113 | 280 | 1,161 |
| BKK30340 | *ytvA* | 3,929 | 1,207 | 0,113 | 277 | 1,161 |
| BKK07750 | *yflA* | 3,917 | 1,214 | 0,129 | 253 | 1,168 |
| BKK09889 | *yhzD* | 3,905 | 1,223 | 0,126 | 134 | 1,177 |
| BKK37560 | *thrZ* | 3,9 | 1,162 | 0,101 | 154 | 1,119 |
| BKK08800 | *ygaK* | 3,887 | 1,204 | 0,127 | 394 | 1,159 |
| BKK14260 | *mobA* | 3,883 | 1,225 | 0,121 | 282 | 1,179 |
| BKK11540 | *pepF* | 3,879 | 1,229 | 0,113 | 89 | 1,183 |
| BKK25289 | *yqzL* | 3,878 | 1,209 | 0,094 | 332 | 1,164 |
| BKK07280 | *yfnG* | 3,874 | 1,225 | 0,149 | 108 | 1,179 |
| BKK10620 | *addB* | 3,874 | 1,229 | 0,102 | 114 | 1,183 |
| BKK30490 | *ytqB* | 3,873 | 1,201 | 0,104 | 140 | 1,156 |
| BKK08210 | *malQ* | 3,87 | 1,255 | 0,111 | 31 | 1,208 |
| BKK20460 | *yoqZ* | 3,87 | 1,173 | 0,157 | 67 | 1,129 |
| BKK18610 | *yoaH* | 3,867 | 1,217 | 0,113 | 138 | 1,172 |
| BKK33430 | *cysI* | 3,864 | 1,013 | 0,078 | 41 | 0,975 |

| BKK name [1] | gene | screening delta[2] (%) | erage width (µ | +/- | nb | ADP |
|---|---|---|---|---|---|---|
| BKK06160 | *gutP* | 3,856 | 1,224 | 0,143 | 116 | 1,179 |
| BKK03220 | *ycgO* | 3,843 | 1,173 | 0,104 | 114 | 1,130 |
| BKK35460 | *comFB* | 3,826 | 1,157 | 0,090 | 103 | 1,114 |
| BKK18239 | *yngHB* | 3,824 | 1,216 | 0,086 | 342 | 1,172 |
| BKK15390 | *sepF* | 3,82 | 1,213 | 0,119 | 153 | 1,168 |
| BKK08050 | *yfjL* | 3,818 | 1,254 | 0,139 | 74 | 1,208 |
| BKK26260 | *yqaM* | 3,811 | 1,222 | 0,111 | 255 | 1,177 |
| BKK07250 | *yetO* | 3,808 | 1,202 | 0,092 | 78 | 1,158 |
| BKK08740 | *ygzB* | 3,808 | 1,203 | 0,110 | 507 | 1,159 |
| BKK19670 | *yodN* | 3,804 | 1,172 | 0,145 | 185 | 1,129 |
| BKK04359 | *ydzK* | 3,799 | 1,209 | 0,091 | 552 | 1,165 |
| BKK05630 | *dinB* | 3,797 | 1,212 | 0,075 | 188 | 1,167 |
| BKK04720 | *rsbW* | 3,788 | 1,209 | 0,106 | 403 | 1,165 |
| BKK10310 | *yhfO* | 3,786 | 1,228 | 0,112 | 141 | 1,183 |
| BKK18220 | *yngF* | 3,784 | 1,216 | 0,099 | 132 | 1,172 |
| BKK34840 | *yvcB* | 3,784 | 1,178 | 0,113 | 132 | 1,135 |
| BKK37180 | *fadF* | 3,778 | 1,161 | 0,130 | 401 | 1,119 |
| BKK14550 | *ykrA* | 3,775 | 1,224 | 0,154 | 69 | 1,179 |
| BKK38310 | *ywbI* | 3,775 | 1,161 | 0,102 | 43 | 1,119 |
| BKK06130 | *ydjC* | 3,769 | 1,223 | 0,141 | 198 | 1,179 |
| BKK12230 | *yjjA* | 3,761 | 1,234 | 0,090 | 127 | 1,189 |
| BKK29590 | *iscS* | 3,761 | 1,155 | 0,127 | 70 | 1,113 |
| BKK02990 | *opuAB* | 3,756 | 1,247 | 0,101 | 134 | 1,201 |
| BKK06370 | *pbuG* | 3,753 | 1,223 | 0,136 | 384 | 1,179 |
| BKK28950 | *thrS* | 3,749 | 1,155 | 0,097 | 32 | 1,113 |
| BKK03640 | *ubiD* | 3,748 | 1,209 | 0,112 | 170 | 1,165 |
| BKK13610 | *mtnB* | 3,744 | 1,210 | 0,145 | 74 | 1,166 |
| BKK32660 | *yurT* | 3,744 | 1,177 | 0,088 | 90 | 1,134 |
| BKK24150 | *mmgC* | 3,741 | 1,231 | 0,103 | 395 | 1,187 |
| BKK09760 | *yheE* | 3,733 | 1,221 | 0,118 | 133 | 1,177 |
| BKK12260 | *yjlA* | 3,732 | 1,233 | 0,065 | 115 | 1,189 |
| BKK15970 | *ylxM* | 3,729 | 1,160 | 0,132 | 73 | 1,119 |
| BKK11560 | *ykbI* | 3,718 | 1,211 | 0,114 | 200 | 1,168 |
| BKK13750 | *queF* | 3,704 | 1,210 | 0,133 | 258 | 1,166 |
| BKK07760 | *yfkT* | 3,698 | 1,202 | 0,123 | 371 | 1,159 |
| BKK18620 | *yoaI* | 3,698 | 1,215 | 0,081 | 250 | 1,172 |
| BKK26920 | *yraJ* | 3,693 | 1,220 | 0,110 | 292 | 1,177 |
| BKK15260 | *ylxX* | 3,681 | 1,223 | 0,110 | 622 | 1,179 |
| BKK29650 | *ytrP* | 3,678 | 1,154 | 0,082 | 59 | 1,113 |
| BKK40350 | *rocR* | 3,667 | 1,193 | 0,121 | 257 | 1,151 |
| BKK38090 | *vpr* | 3,665 | 1,160 | 0,125 | 302 | 1,119 |
| BKK23010 | *ypbD* | 3,663 | 1,141 | 0,114 | 591 | 1,100 |
| BKK23560 | *mleN* | 3,653 | 1,230 | 0,104 | 191 | 1,187 |
| BKK29820 | *ytpR* | 3,65 | 1,154 | 0,108 | 88 | 1,113 |
| BKK11650 | *tenA* | 3,647 | 1,211 | 0,129 | 253 | 1,168 |
| BKK09720 | *yheH* | 3,638 | 1,252 | 0,163 | 44 | 1,208 |
| BKK33000 | *htrB* | 3,629 | 1,203 | 0,107 | 475 | 1,161 |
| BKK06410 | *yebG* | 3,618 | 1,189 | 0,111 | 296 | 1,147 |
| BKK13509 | *ykzP* | 3,604 | 1,208 | 0,138 | 159 | 1,166 |
| BKK21520 | *yolC* | 3,601 | 1,196 | 0,115 | 408 | 1,155 |
| BKK40790 | *yyaO* | 3,598 | 1,189 | 0,121 | 369 | 1,147 |
| BKK09480 | *yhdI* | 3,596 | 1,219 | 0,123 | 129 | 1,177 |
| BKK09010 | *yhcA* | 3,591 | 1,204 | 0,112 | 245 | 1,163 |
| BKK16210 | *fliF* | 3,591 | 1,204 | 0,111 | 224 | 1,163 |
| BKK05160 | *ydeD* | 3,58 | 1,209 | 0,116 | 65 | 1,167 |
| BKK39520 | *yxeK* | 3,577 | 1,206 | 0,115 | 67 | 1,165 |
| BKK07900 | *yfkH* | 3,576 | 1,200 | 0,124 | 95 | 1,159 |
| BKK18270 | *ynzE* | 3,559 | 1,213 | 0,099 | 191 | 1,172 |
| BKK26980 | *yraE* | 3,559 | 1,219 | 0,106 | 284 | 1,177 |
| BKK12060 | *yjdI* | 3,557 | 1,231 | 0,133 | 340 | 1,189 |
| BKK10640 | *sbcD* | 3,556 | 1,225 | 0,100 | 164 | 1,183 |
| BKK20910 | *yopF* | 3,553 | 1,196 | 0,106 | 336 | 1,155 |
| BKK11640 | *yjbQ* | 3,547 | 1,231 | 0,137 | 824 | 1,189 |
| BKK07720 | *yflD* | 3,544 | 1,200 | 0,116 | 327 | 1,159 |
| BKK36270 | *ywqB* | 3,538 | 1,208 | 0,103 | 95 | 1,166 |
| BKK38770 | *cimH* | 3,533 | 1,178 | 0,118 | 53 | 1,138 |
| BKK09520 | *sigM* | 3,531 | 1,219 | 0,150 | 65 | 1,177 |
| BKK27390 | *yrrK* | 3,521 | 1,200 | 0,099 | 448 | 1,160 |
| BKK40650 | *yybG* | 3,521 | 1,226 | 0,116 | 154 | 1,184 |
| BKK40990 | *noc* | 3,518 | 1,226 | 0,100 | 70 | 1,184 |
| BKK07630 | *nosA* | 3,514 | 1,200 | 0,111 | 433 | 1,159 |
| BKK06320 | *yeaB* | 3,513 | 1,200 | 0,114 | 382 | 1,160 |
| BKK29780 | *ytxG* | 3,513 | 1,179 | 0,086 | 114 | 1,139 |
| BKK15380 | *ylmE* | 3,511 | 1,221 | 0,117 | 47 | 1,179 |
| BKK31321 | *mstX* | 3,508 | 1,197 | 0,108 | 92 | 1,156 |
| BKK01740 | *rsiW* | 3,506 | 1,244 | 0,073 | 33 | 1,201 |
| BKK08660 | *sspE* | 3,503 | 1,199 | 0,113 | 358 | 1,159 |
| BKK38410 | *sacX* | 3,502 | 1,197 | 0,106 | 134 | 1,157 |
| BKK34420 | *yveF* | 3,499 | 1,175 | 0,106 | 150 | 1,135 |

| BKK name [1] | gene | screening delta[2] (%) | erage width (µ | +/- | nb | ADP |
|---|---|---|---|---|---|---|
| BKK09740 | *yheF* | 3,49 | 1,218 | 0,113 | 105 | 1,177 |
| BKK16640 | *ylxP* | 3,487 | 1,204 | 0,112 | 110 | 1,164 |
| BKK37990 | *ywdE* | 3,475 | 1,187 | 0,110 | 391 | 1,147 |
| BKK39480 | *yxeO* | 3,474 | 1,264 | 0,091 | 72 | 1,222 |
| BKK01650 | *ybbC* | 3,467 | 1,243 | 0,105 | 34 | 1,201 |
| BKK12450 | *yjpA* | 3,466 | 1,179 | 0,128 | 64 | 1,139 |
| BKK39320 | *yxiB* | 3,458 | 1,205 | 0,113 | 144 | 1,165 |
| BKK21530 | *yolB* | 3,455 | 1,195 | 0,128 | 462 | 1,155 |
| BKK03280 | *nasF* | 3,452 | 1,169 | 0,134 | 64 | 1,130 |
| BKK08425 | *mprF* | 3,446 | 1,208 | 0,116 | 230 | 1,168 |
| BKK21598 | *yoyK* | 3,446 | 1,191 | 0,117 | 243 | 1,151 |
| BKK23190 | *dacB* | 3,446 | 1,248 | 0,168 | 48 | 1,206 |
| BKK38600 | *licR* | 3,442 | 1,205 | 0,112 | 189 | 1,165 |
| BKK18340 | *ppsA* | 3,441 | 1,212 | 0,085 | 43 | 1,172 |
| BKK24830 | *yqgT* | 3,441 | 1,248 | 0,165 | 154 | 1,206 |
| BKK21160 | *yonA* | 3,435 | 1,195 | 0,114 | 404 | 1,155 |
| BKK07350 | *yfmT* | 3,429 | 1,208 | 0,124 | 257 | 1,168 |
| BKK38380 | *ywbB* | 3,425 | 1,196 | 0,120 | 64 | 1,157 |
| BKK15410 | *ylmH* | 3,412 | 1,219 | 0,153 | 72 | 1,179 |
| BKK00490 | *spoVG* | 3,402 | 1,177 | 0,124 | 51 | 1,138 |
| BKK15820 | *rpmB* | 3,397 | 1,174 | 0,102 | 30 | 1,135 |
| BKK04990 | *yddJ* | 3,396 | 1,199 | 0,104 | 98 | 1,160 |
| BKK06269 | *ydzJ* | 3,393 | 1,219 | 0,125 | 242 | 1,179 |
| BKK31180 | *yulE* | 3,387 | 1,195 | 0,129 | 89 | 1,156 |
| BKK04220 | *ydaG* | 3,382 | 1,204 | 0,115 | 274 | 1,165 |
| BKK06270 | *ydjO* | 3,377 | 1,219 | 0,115 | 143 | 1,179 |
| BKK06730 | *yefA* | 3,372 | 1,219 | 0,132 | 263 | 1,179 |
| BKK22760 | *hepS* | 3,371 | 1,192 | 0,109 | 161 | 1,153 |
| BKK25300 | *cdd* | 3,365 | 1,203 | 0,102 | 299 | 1,164 |
| BKK25470 | *dnaK* | 3,365 | 1,203 | 0,103 | 324 | 1,164 |
| BKK14009 | *ykzT* | 3,364 | 1,229 | 0,158 | 168 | 1,189 |
| BKK20800 | *yopQ* | 3,364 | 1,201 | 0,110 | 349 | 1,162 |
| BKK40140 | *yydJ* | 3,363 | 1,206 | 0,140 | 78 | 1,166 |
| BKK26040 | *yqbN* | 3,362 | 1,216 | 0,094 | 148 | 1,177 |
| BKK00700 | *coaX* | 3,358 | 1,178 | 0,097 | 106 | 1,139 |
| BKK01620 | *feuB* | 3,358 | 1,216 | 0,107 | 517 | 1,177 |
| BKK22830 | *gpsA* | 3,352 | 1,137 | 0,115 | 44 | 1,100 |
| BKK20440 | *yorB* | 3,336 | 1,167 | 0,145 | 88 | 1,129 |
| BKK24330 | *yqhY* | 3,333 | 1,189 | 0,123 | 307 | 1,151 |
| BKK30750 | *mntC* | 3,328 | 1,263 | 0,088 | 47 | 1,222 |
| BKK13060 | *ykjA* | 3,323 | 1,205 | 0,098 | 44 | 1,166 |
| BKK09730 | *yheG* | 3,32 | 1,216 | 0,123 | 146 | 1,177 |
| BKK26050 | *yqdB* | 3,32 | 1,192 | 0,124 | 242 | 1,153 |
| BKK11450 | *oppC* | 3,318 | 1,207 | 0,122 | 237 | 1,168 |
| BKK40280 | *yycO* | 3,318 | 1,189 | 0,114 | 311 | 1,151 |
| BKK26180 | *yqbA* | 3,317 | 1,216 | 0,117 | 139 | 1,177 |
| BKK09200 | *yhcS* | 3,316 | 1,176 | 0,111 | 59 | 1,138 |
| BKK08970 | *prkA* | 3,298 | 1,216 | 0,121 | 163 | 1,177 |
| BKK28460 | *yslB* | 3,296 | 1,196 | 0,097 | 87 | 1,158 |
| BKK16200 | *fliE* | 3,292 | 1,201 | 0,109 | 347 | 1,163 |
| BKK05730 | *ydhF* | 3,291 | 1,198 | 0,105 | 346 | 1,160 |
| BKK29480 | *ytxK* | 3,29 | 1,150 | 0,132 | 82 | 1,113 |
| BKK37830 | *spsJ* | 3,286 | 1,156 | 0,146 | 53 | 1,119 |
| BKK05740 | *phoB* | 3,278 | 1,205 | 0,083 | 202 | 1,167 |
| BKK14450 | *ampS* | 3,274 | 1,218 | 0,103 | 66 | 1,179 |
| BKK39030 | *yxiT* | 3,271 | 1,203 | 0,133 | 117 | 1,165 |
| BKK13799 | *ykzR* | 3,256 | 1,204 | 0,120 | 177 | 1,166 |
| BKK21610 | *yokF* | 3,25 | 1,188 | 0,114 | 341 | 1,151 |
| BKK08270 | *yfiH* | 3,247 | 1,196 | 0,117 | 410 | 1,159 |
| BKK21970 | *ypeP* | 3,241 | 1,192 | 0,108 | 486 | 1,155 |
| BKK17080 | *pksA* | 3,24 | 1,201 | 0,119 | 62 | 1,164 |
| BKK34430 | *racX* | 3,238 | 1,187 | 0,114 | 381 | 1,149 |
| BKK34320 | *epsF* | 3,232 | 1,187 | 0,109 | 371 | 1,149 |
| BKK29270 | *ytoI* | 3,23 | 1,149 | 0,099 | 186 | 1,113 |
| BKK34570 | *malK* | 3,223 | 1,172 | 0,123 | 31 | 1,135 |
| BKK22680 | *trpE* | 3,222 | 1,188 | 0,114 | 242 | 1,151 |
| BKK22700 | *aroB* | 3,222 | 1,136 | 0,094 | 42 | 1,100 |
| BKK35470 | *comFA* | 3,222 | 1,199 | 0,118 | 398 | 1,161 |
| BKK07880 | *yfkJ* | 3,209 | 1,206 | 0,116 | 272 | 1,168 |
| BKK10630 | *addA* | 3,208 | 1,221 | 0,105 | 125 | 1,183 |
| BKK14140 | *ccpC* | 3,208 | 1,204 | 0,117 | 884 | 1,166 |
| BKK30700 | *rpmEB* | 3,206 | 1,195 | 0,104 | 119 | 1,158 |
| BKK13970 | *ykwD* | 3,198 | 1,200 | 0,128 | 215 | 1,163 |
| BKK32490 | *pucC* | 3,195 | 1,185 | 0,122 | 40 | 1,148 |
| BKK30950 | *glgA* | 3,188 | 1,193 | 0,098 | 122 | 1,156 |
| BKK37290 | *arfM* | 3,17 | 1,154 | 0,106 | 59 | 1,119 |
| BKK04290 | *ydaL* | 3,166 | 1,202 | 0,099 | 156 | 1,165 |
| BKK26760 | *yrdC* | 3,156 | 1,214 | 0,110 | 172 | 1,177 |
| BKK11410 | *yjbA* | 3,155 | 1,221 | 0,134 | 94 | 1,183 |

| BKK name [1] | gene | screening delta[2] (%) | erage width (µ | +/- | nb | ADP |
|---|---|---|---|---|---|---|
| BKK12680 | *xkdO* | 3,154 | 1,227 | 0,162 | 83 | 1,189 |
| BKK35320 | *fliT* | 3,153 | 1,171 | 0,124 | 449 | 1,135 |
| BKK36160 | *ywqM* | 3,153 | 1,149 | 0,126 | 92 | 1,114 |
| BKK15350 | *thiQ* | 3,152 | 1,216 | 0,152 | 168 | 1,179 |
| BKK09330 | *yhcZ* | 3,15 | 1,205 | 0,118 | 286 | 1,168 |
| BKK11010 | *yitJ* | 3,134 | 1,220 | 0,094 | 130 | 1,183 |
| BKK39129 | *yxzI* | 3,13 | 1,201 | 0,116 | 114 | 1,165 |
| BKK14072 | *ykzU* | 3,128 | 1,203 | 0,153 | 401 | 1,166 |
| BKK21070 | *yonJ* | 3,124 | 1,191 | 0,126 | 185 | 1,155 |
| BKK40460 | *yyzB* | 3,116 | 1,203 | 0,105 | 128 | 1,166 |
| BKK06210 | *ydjI* | 3,115 | 1,196 | 0,108 | 899 | 1,160 |
| BKK32669 | *yuzN* | 3,114 | 1,170 | 0,103 | 160 | 1,134 |
| BKK24790 | *yqgX* | 3,11 | 1,200 | 0,100 | 262 | 1,164 |
| BKK09560 | *cueR* | 3,108 | 1,246 | 0,119 | 96 | 1,208 |
| BKK07830 | *yfkO* | 3,104 | 1,195 | 0,120 | 398 | 1,159 |
| BKK27620 | *recJ* | 3,104 | 1,196 | 0,104 | 151 | 1,160 |
| BKK35130 | *yvlA* | 3,101 | 1,170 | 0,104 | 377 | 1,135 |
| BKK39830 | *yxcA* | 3,097 | 1,193 | 0,102 | 115 | 1,158 |
| BKK26630 | *yrdQ* | 3,093 | 1,189 | 0,119 | 303 | 1,153 |
| BKK03570 | *sfp* | 3,091 | 1,193 | 0,107 | 80 | 1,158 |
| BKK07840 | *yfkN* | 3,09 | 1,195 | 0,112 | 454 | 1,159 |
| BKK25660 | *aroD* | 3,09 | 1,200 | 0,101 | 462 | 1,164 |
| BKK38360 | *ywbD* | 3,085 | 1,153 | 0,135 | 37 | 1,119 |
| BKK05610 | *vmlR* | 3,078 | 1,195 | 0,125 | 425 | 1,160 |
| BKK40960 | *parB* | 3,071 | 1,221 | 0,101 | 118 | 1,184 |
| BKK28560 | *lcfA* | 3,07 | 1,192 | 0,110 | 102 | 1,156 |
| BKK29000 | *nrdR* | 3,063 | 1,193 | 0,094 | 114 | 1,158 |
| BKK08870 | *ygaN* | 3,062 | 1,213 | 0,108 | 128 | 1,177 |
| BKK10150 | *yhgD* | 3,062 | 1,213 | 0,103 | 138 | 1,177 |
| BKK01550 | *gerD* | 3,061 | 1,173 | 0,112 | 35 | 1,138 |
| BKK06600 | *pcrB* | 3,061 | 1,215 | 0,113 | 261 | 1,179 |
| BKK27420 | *yrrI* | 3,06 | 1,189 | 0,112 | 290 | 1,153 |
| BKK14890 | *ctaC* | 3,059 | 1,215 | 0,109 | 327 | 1,179 |
| BKK14770 | *bipA* | 3,055 | 1,215 | 0,097 | 259 | 1,179 |
| BKK09980 | *yhaI* | 3,054 | 1,245 | 0,158 | 82 | 1,208 |
| BKK08250 | *yfiF* | 3,052 | 1,194 | 0,129 | 108 | 1,159 |
| BKK28540 | *fadB* | 3,049 | 1,005 | 0,094 | 159 | 0,975 |
| BKK11110 | *yitS* | 3,048 | 1,219 | 0,099 | 76 | 1,183 |
| BKK27809 | *yrzT* | 3,048 | 1,195 | 0,103 | 164 | 1,160 |
| BKK39010 | *yxjB* | 3,048 | 1,200 | 0,116 | 101 | 1,165 |
| BKK27130 | *rsiV* | 3,047 | 1,259 | 0,067 | 80 | 1,222 |
| BKK16180 | *flgB* | 3,045 | 1,204 | 0,101 | 348 | 1,168 |
| BKK37620 | *rsfA* | 3,044 | 1,153 | 0,107 | 78 | 1,119 |
| BKK31020 | *yuaF* | 3,043 | 1,197 | 0,120 | 371 | 1,161 |
| BKK00750 | *pabA* | 3,041 | 1,134 | 0,116 | 53 | 1,100 |
| BKK02920 | *yceF* | 3,039 | 1,238 | 0,152 | 67 | 1,201 |
| BKK38590 | *licB* | 3,037 | 1,200 | 0,114 | 83 | 1,165 |
| BKK08290 | *yfjJ* | 3,034 | 1,245 | 0,172 | 39 | 1,208 |
| BKK13550 | *mtnA* | 3,034 | 1,202 | 0,126 | 162 | 1,166 |
| BKK19450 | *rsbRC* | 3,027 | 1,163 | 0,118 | 459 | 1,129 |
| BKK31710 | *comQ* | 3,02 | 1,183 | 0,123 | 52 | 1,148 |
| BKK22870 | *fni* | 3,018 | 1,133 | 0,100 | 251 | 1,100 |
| BKK07590 | *citT* | 3,013 | 1,203 | 0,103 | 261 | 1,168 |
| BKK17320 | *ymaF* | 3,007 | 1,199 | 0,096 | 117 | 1,164 |
| BKK33680 | *yvaP* | 2,985 | 1,168 | 0,109 | 71 | 1,134 |
| BKK23640 | *yqkD* | 2,984 | 1,222 | 0,109 | 241 | 1,187 |
| BKK00950 | *mrnC* | 2,982 | 1,222 | 0,099 | 150 | 1,187 |
| BKK27670 | *spoVB* | 2,966 | 1,258 | 0,069 | 80 | 1,222 |
| BKK24070 | *buk* | 2,959 | 1,222 | 0,113 | 356 | 1,187 |
| BKK09210 | *yhcT* | 2,958 | 1,212 | 0,098 | 147 | 1,177 |
| BKK20010 | *yosT* | 2,957 | 1,163 | 0,123 | 351 | 1,129 |
| BKK26170 | *yqbB* | 2,955 | 1,211 | 0,100 | 159 | 1,177 |
| BKK05360 | *ydfC* | 2,947 | 1,202 | 0,089 | 163 | 1,167 |
| BKK23930 | *yqjC* | 2,942 | 1,222 | 0,097 | 343 | 1,187 |
| BKK03630 | *ubiX* | 2,94 | 1,199 | 0,109 | 77 | 1,165 |
| BKK10510 | *yhjH* | 2,939 | 1,218 | 0,093 | 161 | 1,183 |
| BKK25520 | *yqxA* | 2,937 | 1,185 | 0,103 | 262 | 1,151 |
| BKK09630 | *yhdX* | 2,931 | 1,211 | 0,120 | 85 | 1,177 |
| BKK35350 | *yvyC* | 2,928 | 1,168 | 0,106 | 233 | 1,135 |
| BKK06520 | *purH* | 2,922 | 1,213 | 0,122 | 487 | 1,179 |
| BKK31959 | *ybdZ* | 2,92 | 1,181 | 0,107 | 230 | 1,148 |
| BKK14200 | *ykuS* | 2,919 | 1,214 | 0,147 | 767 | 1,179 |
| BKK38550 | *ilvK* | 2,917 | 1,191 | 0,135 | 85 | 1,157 |
| BKK20930 | *yopD* | 2,914 | 1,189 | 0,122 | 528 | 1,155 |
| BKK37920 | *ywdL* | 2,914 | 1,191 | 0,112 | 67 | 1,157 |
| BKK06590 | *yerD* | 2,912 | 1,213 | 0,137 | 180 | 1,179 |
| BKK23080 | *aroC* | 2,909 | 1,132 | 0,097 | 247 | 1,100 |
| BKK05500 | *ydfP* | 2,908 | 1,201 | 0,092 | 309 | 1,167 |
| BKK23440 | *spoVAA* | 2,906 | 1,242 | 0,157 | 38 | 1,206 |

### Sup. Table 4: Cell width of mutants of the BKK collection (continued)

| BKK name [1] | gene | screening delta[2] (%) | erage width (µ | +/- | nb | ADP |
|---|---|---|---|---|---|---|
| BKK09360 | *yhdC* | 2,9 | 1,211 | 0,092 | 94 | 1,177 |
| BKK29080 | *mutM* | 2,9 | 1,195 | 0,120 | 488 | 1,161 |
| BKK08730 | *perR* | 2,899 | 1,191 | 0,103 | 116 | 1,158 |
| BKK35080 | *yvmB* | 2,895 | 1,168 | 0,125 | 76 | 1,135 |
| BKK03950 | *ycnJ* | 2,894 | 1,199 | 0,085 | 135 | 1,165 |
| BKK39550 | *yxeH* | 2,888 | 1,198 | 0,101 | 161 | 1,165 |
| BKK09460 | *ctrA* | 2,887 | 1,196 | 0,111 | 268 | 1,163 |
| BKK18390 | *yoeC* | 2,885 | 1,195 | 0,110 | 289 | 1,162 |
| BKK18680 | *yoaO* | 2,883 | 1,205 | 0,101 | 61 | 1,172 |
| BKK00080 | *yaaC* | 2,872 | 1,193 | 0,126 | 254 | 1,160 |
| BKK06560 | *yerA* | 2,872 | 1,213 | 0,104 | 402 | 1,179 |
| BKK38640 | *yxlH* | 2,869 | 1,191 | 0,108 | 155 | 1,158 |
| BKK24660 | *yqzE* | 2,868 | 1,197 | 0,106 | 173 | 1,164 |
| BKK06033 | *ydzT* | 2,867 | 1,201 | 0,088 | 410 | 1,167 |
| BKK06880 | *yesF* | 2,866 | 1,213 | 0,142 | 279 | 1,179 |
| BKK32190 | *yuzB* | 2,864 | 1,181 | 0,138 | 110 | 1,148 |
| BKK13950 | *mcpC* | 2,857 | 1,200 | 0,144 | 76 | 1,166 |
| BKK23620 | *yqkF* | 2,844 | 1,186 | 0,125 | 296 | 1,153 |
| BKK32600 | *frlO* | 2,843 | 1,257 | 0,129 | 176 | 1,222 |
| BKK27210 | *yrhF* | 2,841 | 1,210 | 0,105 | 358 | 1,177 |
| BKK06720 | *dagK* | 2,84 | 1,218 | 0,125 | 80 | 1,184 |
| BKK05790 | *ydhK* | 2,839 | 1,200 | 0,092 | 148 | 1,167 |
| BKK13920 | *splA* | 2,839 | 1,199 | 0,124 | 135 | 1,166 |
| BKK28330 | *ysnE* | 2,838 | 1,186 | 0,124 | 521 | 1,153 |
| BKK40820 | *yyaL* | 2,836 | 1,218 | 0,102 | 168 | 1,184 |
| BKK27720 | *queA* | 2,831 | 1,192 | 0,120 | 201 | 1,160 |
| BKK14160 | *ykuO* | 2,83 | 1,213 | 0,135 | 1011 | 1,179 |
| BKK14670 | *suhB* | 2,829 | 1,213 | 0,125 | 249 | 1,179 |
| BKK33040 | *fumC* | 2,827 | 1,166 | 0,087 | 48 | 1,134 |
| BKK10530 | *ntdC* | 2,821 | 1,201 | 0,107 | 252 | 1,168 |
| BKK07000 | *yesR* | 2,82 | 1,212 | 0,125 | 227 | 1,179 |
| BKK01460 | *ybaE* | 2,819 | 1,170 | 0,116 | 62 | 1,138 |
| BKK18830 | *pps* | 2,819 | 1,170 | 0,090 | 121 | 1,138 |
| BKK39500 | *yxeM* | 2,818 | 1,256 | 0,074 | 65 | 1,222 |
| BKK21790 | *yplQ* | 2,809 | 1,186 | 0,110 | 344 | 1,153 |
| BKK06490 | *purF* | 2,803 | 1,212 | 0,121 | 358 | 1,179 |
| BKK18260 | *yngJ* | 2,798 | 1,204 | 0,116 | 59 | 1,172 |
| BKK33950 | *cggR* | 2,798 | 1,166 | 0,099 | 107 | 1,134 |
| BKK19240 | *yocK* | 2,796 | 1,194 | 0,121 | 410 | 1,162 |
| BKK20760 | *yopU* | 2,796 | 1,187 | 0,105 | 503 | 1,155 |
| BKK12300 | *uxaC* | 2,795 | 1,222 | 0,142 | 347 | 1,189 |
| BKK20720 | *yopY* | 2,791 | 1,201 | 0,124 | 192 | 1,168 |
| BKK26730 | *yrdF* | 2,79 | 1,210 | 0,094 | 77 | 1,177 |
| BKK38930 | *yxjJ* | 2,787 | 1,197 | 0,130 | 46 | 1,165 |
| BKK18290 | *yngL* | 2,786 | 1,194 | 0,118 | 340 | 1,162 |
| BKK26010 | *yqbQ* | 2,786 | 1,196 | 0,115 | 72 | 1,164 |
| BKK38300 | *thiM* | 2,781 | 1,199 | 0,110 | 115 | 1,166 |
| BKK18420 | *ftsR* | 2,779 | 1,204 | 0,083 | 219 | 1,172 |
| BKK25569 | *yqzM* | 2,779 | 1,196 | 0,102 | 333 | 1,164 |
| BKK38720 | *yxkO* | 2,777 | 1,179 | 0,110 | 397 | 1,147 |
| BKK34990 | *lgt* | 2,773 | 1,181 | 0,108 | 535 | 1,149 |
| BKK01730 | *sigW* | 2,77 | 1,192 | 0,087 | 103 | 1,160 |
| BKK23370 | *ypuA* | 2,77 | 1,185 | 0,124 | 247 | 1,153 |
| BKK08770 | *ygzA* | 2,769 | 1,191 | 0,122 | 552 | 1,159 |
| BKK25950 | *yqcC* | 2,766 | 1,002 | 0,100 | 88 | 0,975 |
| BKK29150 | *ytwI* | 2,765 | 1,193 | 0,116 | 466 | 1,161 |
| BKK22400 | *dinG* | 2,757 | 1,131 | 0,117 | 87 | 1,100 |
| BKK40720 | *yyaT* | 2,754 | 1,217 | 0,096 | 113 | 1,184 |
| BKK09165 | *yhcO* | 2,751 | 1,200 | 0,117 | 336 | 1,168 |
| BKK40710 | *yybA* | 2,751 | 1,217 | 0,096 | 171 | 1,184 |
| BKK04590 | *ydbS* | 2,75 | 1,197 | 0,091 | 424 | 1,165 |
| BKK18669 | *yozS* | 2,75 | 1,194 | 0,120 | 196 | 1,162 |
| BKK20530 | *yoqS* | 2,749 | 1,200 | 0,105 | 292 | 1,168 |
| BKK15830 | *yloU* | 2,745 | 1,145 | 0,096 | 102 | 1,114 |
| BKK16730 | *spoVFA* | 2,74 | 1,240 | 0,125 | 96 | 1,206 |
| BKK24940 | *yqzC* | 2,732 | 1,195 | 0,114 | 222 | 1,164 |
| BKK28200 | *lonA* | 2,732 | 1,183 | 0,120 | 278 | 1,151 |
| BKK07930 | *yfkD* | 2,729 | 1,190 | 0,109 | 450 | 1,159 |
| BKK12040 | *yjdG* | 2,727 | 1,221 | 0,157 | 285 | 1,189 |
| BKK12130 | *yjfC* | 2,727 | 1,221 | 0,157 | 285 | 1,189 |
| BKK03020 | *ycgA* | 2,726 | 1,234 | 0,075 | 101 | 1,201 |
| BKK34100 | *rsbQ* | 2,726 | 1,166 | 0,153 | 96 | 1,135 |
| BKK22540 | *qcrC* | 2,721 | 1,130 | 0,119 | 419 | 1,100 |
| BKK15530 | *pyrK* | 2,72 | 1,187 | 0,103 | 87 | 1,156 |
| BKK34010 | *yvbW* | 2,719 | 1,189 | 0,097 | 77 | 1,158 |
| BKK16440 | *cheW* | 2,714 | 1,193 | 0,106 | 399 | 1,161 |
| BKK19930 | *yotC* | 2,704 | 1,193 | 0,131 | 428 | 1,162 |
| BKK13180 | *metE* | 2,696 | 1,170 | 0,130 | 39 | 1,139 |
| BKK07800 | *treP* | 2,69 | 1,190 | 0,115 | 388 | 1,159 |

| BKK name [1] | gene | screening delta[2] (%) | erage width (µ | +/- | nb | ADP |
|---|---|---|---|---|---|---|
| BKK30560 | *pckA* | 2,685 | 1,187 | 0,120 | 102 | 1,156 |
| BKK30460 | *ytrA* | 2,676 | 1,187 | 0,130 | 108 | 1,156 |
| BKK03170 | *ycgK* | 2,668 | 1,191 | 0,111 | 325 | 1,160 |
| BKK03700 | *gerKA* | 2,667 | 1,196 | 0,090 | 126 | 1,165 |
| BKK26039 | *yqbN* | 2,667 | 1,208 | 0,103 | 545 | 1,177 |
| BKK13640 | *spo0E* | 2,664 | 1,241 | 0,155 | 58 | 1,208 |
| BKK13860 | *ykvY* | 2,664 | 1,197 | 0,151 | 329 | 1,166 |
| BKK01140 | *ybaC* | 2,661 | 1,195 | 0,112 | 228 | 1,164 |
| BKK32290 | *yutF* | 2,652 | 1,192 | 0,116 | 377 | 1,161 |
| BKK08440 | *yfiY* | 2,651 | 1,240 | 0,172 | 67 | 1,208 |
| BKK09880 | *yhaR* | 2,641 | 1,208 | 0,103 | 153 | 1,177 |
| BKK21300 | *yomM* | 2,633 | 1,181 | 0,105 | 141 | 1,151 |
| BKK09100 | *cspB* | 2,629 | 1,208 | 0,098 | 129 | 1,177 |
| BKK27650 | *secDF* | 2,629 | 1,254 | 0,084 | 135 | 1,222 |
| BKK00180 | *tadA* | 2,627 | 1,208 | 0,098 | 98 | 1,177 |
| BKK31310 | *yugP* | 2,625 | 1,186 | 0,112 | 75 | 1,156 |
| BKK21460 | *bdbA* | 2,622 | 1,181 | 0,120 | 280 | 1,151 |
| BKK30010 | *ythP* | 2,621 | 1,186 | 0,096 | 92 | 1,156 |
| BKK08120 | *yfjF* | 2,617 | 1,189 | 0,110 | 158 | 1,159 |
| BKK04540 | *ydbO* | 2,616 | 1,233 | 0,097 | 318 | 1,201 |
| BKK15120 | *yllA* | 2,614 | 1,210 | 0,126 | 188 | 1,179 |
| BKK03880 | *yczG* | 2,606 | 1,195 | 0,104 | 478 | 1,165 |
| BKK28070 | *comC* | 2,606 | 1,190 | 0,116 | 242 | 1,160 |
| BKK31040 | *yuaD* | 2,603 | 1,186 | 0,117 | 118 | 1,156 |
| BKK07120 | *lplC* | 2,602 | 1,240 | 0,154 | 538 | 1,208 |
| BKK18380 | *iseA* | 2,601 | 1,238 | 0,146 | 84 | 1,206 |
| BKK40570 | *yybO* | 2,599 | 1,197 | 0,115 | 141 | 1,166 |
| BKK22060 | *pbuX* | 2,598 | 1,169 | 0,122 | 97 | 1,139 |
| BKK27200 | *yrhG* | 2,596 | 1,207 | 0,097 | 296 | 1,177 |
| BKK18140 | *ynfE* | 2,595 | 1,202 | 0,124 | 213 | 1,172 |
| BKK40250 | *yycR* | 2,594 | 1,177 | 0,110 | 255 | 1,147 |
| BKK21060 | *yonK* | 2,593 | 1,185 | 0,113 | 539 | 1,155 |
| BKK25180 | *trmK* | 2,58 | 1,181 | 0,113 | 344 | 1,151 |
| BKK10110 | *pbpF* | 2,579 | 1,207 | 0,110 | 107 | 1,177 |
| BKK04320 | *ydaO* | 2,576 | 1,232 | 0,070 | 42 | 1,201 |
| BKK29420 | *ytkK* | 2,576 | 1,000 | 0,061 | 39 | 0,975 |
| BKK39380 | *hutG* | 2,576 | 1,195 | 0,136 | 88 | 1,165 |
| BKK28240 | *ysoA* | 2,573 | 1,189 | 0,103 | 185 | 1,160 |
| BKK29190 | *pfkA* | 2,573 | 1,142 | 0,099 | 199 | 1,113 |
| BKK15850 | *sdaAB* | 2,564 | 1,143 | 0,077 | 79 | 1,114 |
| BKK01870 | *ybcH* | 2,559 | 1,000 | 0,090 | 59 | 0,975 |
| BKK26450 | *yrkN* | 2,559 | 1,183 | 0,119 | 334 | 1,153 |
| BKK04610 | *ydcA* | 2,55 | 1,195 | 0,101 | 135 | 1,165 |
| BKK02380 | *ybgB* | 2,546 | 1,159 | 0,086 | 63 | 1,130 |
| BKK11549 | *yizD* | 2,543 | 1,213 | 0,106 | 84 | 1,183 |
| BKK04100 | *kipR* | 2,536 | 1,194 | 0,080 | 126 | 1,165 |
| BKK17100 | *pksC* | 2,535 | 1,193 | 0,106 | 145 | 1,164 |
| BKK28810 | *abnA* | 2,533 | 1,168 | 0,104 | 123 | 1,139 |
| BKK09240 | *yhcW* | 2,532 | 1,207 | 0,107 | 99 | 1,177 |
| BKK03190 | *ycgL* | 2,53 | 1,159 | 0,128 | 72 | 1,130 |
| BKK24410 | *spoIIIAC* | 2,53 | 1,237 | 0,122 | 84 | 1,206 |
| BKK19710 | *yodQ* | 2,527 | 1,158 | 0,134 | 190 | 1,129 |
| BKK26190 | *yqaT* | 2,527 | 1,180 | 0,135 | 334 | 1,151 |
| BKK03800 | *yclN* | 2,52 | 1,232 | 0,152 | 603 | 1,201 |
| BKK33700 | *opuBD* | 2,515 | 1,163 | 0,094 | 94 | 1,134 |
| BKK13490 | *htpX* | 2,507 | 1,196 | 0,129 | 69 | 1,166 |
| BKK08400 | *yfiU* | 2,499 | 1,197 | 0,121 | 249 | 1,168 |
| BKK23401 | *spoVAEA* | 2,495 | 1,237 | 0,158 | 95 | 1,206 |
| BKK01540 | *salA* | 2,494 | 1,206 | 0,108 | 231 | 1,177 |
| BKK24060 | *lpdV* | 2,494 | 1,217 | 0,116 | 246 | 1,187 |
| BKK19680 | *yozE* | 2,486 | 1,157 | 0,124 | 673 | 1,129 |
| BKK07370 | *yfmR* | 2,485 | 1,238 | 0,174 | 111 | 1,208 |
| BKK28050 | *maf* | 2,476 | 1,188 | 0,095 | 158 | 1,160 |
| BKK25320 | *yqfG* | 2,474 | 1,194 | 0,111 | 400 | 1,165 |
| BKK29310 | *ytnJ* | 2,466 | 1,141 | 0,093 | 67 | 1,113 |
| BKK21210 | *yomV* | 2,463 | 1,183 | 0,121 | 218 | 1,155 |
| BKK09140 | *yhcM* | 2,46 | 1,206 | 0,108 | 110 | 1,177 |
| BKK14380 | *fruR* | 2,458 | 1,208 | 0,156 | 69 | 1,179 |
| BKK34230 | *epsN* | 2,455 | 1,163 | 0,114 | 198 | 1,135 |
| BKK39140 | *yxiK* | 2,453 | 1,193 | 0,123 | 112 | 1,165 |
| BKK00390 | *yabD* | 2,448 | 1,167 | 0,137 | 58 | 1,139 |
| BKK31990 | *dhbC* | 2,448 | 1,186 | 0,093 | 116 | 1,158 |
| BKK27729 | *yrzS* | 2,446 | 1,188 | 0,101 | 429 | 1,160 |
| BKK15840 | *yloV* | 2,437 | 1,141 | 0,096 | 124 | 1,114 |
| BKK30960 | *glgD* | 2,436 | 1,184 | 0,098 | 87 | 1,156 |
| BKK23890 | *oxaAB* | 2,434 | 1,236 | 0,193 | 31 | 1,206 |
| BKK04440 | *dctB* | 2,43 | 1,193 | 0,117 | 267 | 1,165 |
| BKK35190 | *yvkC* | 2,43 | 1,163 | 0,094 | 354 | 1,135 |
| BKK09750 | *sspB* | 2,429 | 1,206 | 0,094 | 76 | 1,177 |

| BKK name [1] | gene | screening delta[2] (%) | erage width (µ | +/- | nb | ADP |
|---|---|---|---|---|---|---|
| BKK11100 | *nprB* | 2,429 | 1,212 | 0,120 | 122 | 1,183 |
| BKK20330 | *yorM* | 2,428 | 1,190 | 0,102 | 219 | 1,162 |
| BKK01520 | *ybaK* | 2,427 | 0,999 | 0,056 | 30 | 0,975 |
| BKK36170 | *ywqL* | 2,427 | 1,141 | 0,139 | 31 | 1,114 |
| BKK36630 | *ywnA* | 2,427 | 1,175 | 0,113 | 275 | 1,147 |
| BKK34370 | *epsA* | 2,426 | 1,163 | 0,083 | 192 | 1,135 |
| BKK34480 | *yvdT* | 2,426 | 1,163 | 0,083 | 192 | 1,135 |
| BKK21229 | *youA* | 2,423 | 1,183 | 0,105 | 382 | 1,155 |
| BKK35580 | *tuaD* | 2,416 | 1,177 | 0,124 | 516 | 1,149 |
| BKK14350 | *yknX* | 2,415 | 1,165 | 0,092 | 154 | 1,138 |
| BKK16190 | *flgC* | 2,412 | 1,191 | 0,106 | 426 | 1,163 |
| BKK26770 | *yrdB* | 2,411 | 1,205 | 0,103 | 262 | 1,177 |
| BKK31810 | *yuzE* | 2,411 | 1,176 | 0,120 | 271 | 1,148 |
| BKK20929 | *yoyI* | 2,41 | 1,183 | 0,112 | 505 | 1,155 |
| BKK31460 | *kapB* | 2,409 | 1,176 | 0,127 | 31 | 1,148 |
| BKK21220 | *yomU* | 2,407 | 1,179 | 0,119 | 403 | 1,151 |
| BKK03350 | *yciB* | 2,389 | 1,230 | 0,104 | 61 | 1,201 |
| BKK13940 | *ykwB* | 2,385 | 1,190 | 0,112 | 280 | 1,163 |
| BKK38940 | *yxjI* | 2,385 | 1,192 | 0,124 | 88 | 1,165 |
| BKK00450 | *sspF* | 2,384 | 1,191 | 0,101 | 123 | 1,164 |
| BKK31920 | *yukF* | 2,384 | 1,175 | 0,126 | 89 | 1,148 |
| BKK10790 | *asnO* | 2,382 | 1,211 | 0,116 | 85 | 1,183 |
| BKK13290 | *ykzD* | 2,377 | 1,190 | 0,118 | 220 | 1,163 |
| BKK31430 | *yugE* | 2,375 | 1,175 | 0,128 | 61 | 1,148 |
| BKK26990 | *yraD* | 2,374 | 1,205 | 0,105 | 205 | 1,177 |
| BKK37160 | *rpoE* | 2,37 | 1,145 | 0,118 | 331 | 1,119 |
| BKK07320 | *yfnC* | 2,354 | 1,237 | 0,153 | 522 | 1,208 |
| BKK06980 | *yesP* | 2,349 | 1,165 | 0,101 | 30 | 1,138 |
| BKK08930 | *cspR* | 2,349 | 1,190 | 0,114 | 213 | 1,163 |
| BKK18810 | *yobA* | 2,347 | 1,235 | 0,138 | 54 | 1,206 |
| BKK03440 | *tlpC* | 2,344 | 1,187 | 0,127 | 413 | 1,160 |
| BKK09220 | *yhcU* | 2,344 | 1,205 | 0,111 | 165 | 1,177 |
| BKK37670 | *ywfI* | 2,339 | 1,212 | 0,113 | 98 | 1,184 |
| BKK02660 | *ycbU* | 2,334 | 0,998 | 0,098 | 50 | 0,975 |
| BKK02410 | *ybgG* | 2,326 | 1,156 | 0,088 | 126 | 1,130 |
| BKK38990 | *scoA* | 2,326 | 1,192 | 0,107 | 74 | 1,165 |
| BKK36980 | *ywlA* | 2,315 | 1,184 | 0,114 | 126 | 1,158 |
| BKK07100 | *lplA* | 2,314 | 1,236 | 0,119 | 105 | 1,208 |
| BKK05520 | *ydzH* | 2,313 | 1,194 | 0,083 | 333 | 1,167 |
| BKK38970 | *yxjF* | 2,312 | 1,193 | 0,107 | 82 | 1,166 |
| BKK05470 | *ydfM* | 2,31 | 1,229 | 0,099 | 95 | 1,201 |
| BKK33800 | *opuCD* | 2,309 | 1,176 | 0,115 | 336 | 1,149 |
| BKK20830 | *yopN* | 2,307 | 1,182 | 0,122 | 343 | 1,155 |
| BKK31279 | *yuzH* | 2,303 | 1,183 | 0,112 | 152 | 1,156 |
| BKK00410 | *rnmV* | 2,3 | 1,190 | 0,117 | 75 | 1,164 |
| BKK33490 | *copB* | 2,298 | 1,188 | 0,112 | 291 | 1,161 |
| BKK37420 | *albF* | 2,296 | 1,193 | 0,109 | 119 | 1,166 |
| BKK17280 | *ymaD* | 2,295 | 1,190 | 0,117 | 96 | 1,164 |
| BKK12650 | *xkdK* | 2,292 | 1,216 | 0,152 | 131 | 1,189 |
| BKK03300 | *nasD* | 2,29 | 1,156 | 0,132 | 58 | 1,130 |
| BKK25360 | *yqfC* | 2,29 | 1,190 | 0,103 | 151 | 1,164 |
| BKK08110 | *sspH* | 2,289 | 1,185 | 0,122 | 149 | 1,159 |
| BKK24980 | *pstC* | 2,289 | 1,177 | 0,124 | 413 | 1,151 |
| BKK40580 | *yybN* | 2,288 | 1,193 | 0,136 | 109 | 1,166 |
| BKK36510 | *amtB* | 2,286 | 1,176 | 0,108 | 331 | 1,149 |
| BKK12290 | *ndh* | 2,285 | 1,216 | 0,127 | 932 | 1,189 |
| BKK28230 | *tig* | 2,284 | 1,186 | 0,101 | 191 | 1,160 |
| BKK02240 | *mpr* | 2,283 | 1,156 | 0,113 | 144 | 1,130 |
| BKK35230 | *swrAA* | 2,282 | 1,161 | 0,116 | 321 | 1,135 |
| BKK09080 | *yhcH* | 2,281 | 1,195 | 0,121 | 420 | 1,168 |
| BKK00590 | *yabO* | 2,271 | 1,198 | 0,128 | 138 | 1,172 |
| BKK36440 | *ywoH* | 2,271 | 1,184 | 0,092 | 98 | 1,158 |
| BKK37520 | *ywhD* | 2,271 | 1,193 | 0,109 | 91 | 1,166 |
| BKK28400 | *ysmB* | 2,268 | 1,186 | 0,091 | 267 | 1,160 |
| BKK40850 | *maa* | 2,264 | 1,211 | 0,105 | 118 | 1,184 |
| BKK10230 | *yhfH* | 2,262 | 1,204 | 0,116 | 112 | 1,177 |
| BKK38890 | *yxjN* | 2,261 | 1,191 | 0,111 | 166 | 1,165 |
| BKK20610 | *yoqJ* | 2,255 | 1,181 | 0,135 | 124 | 1,155 |
| BKK26650 | *czcD* | 2,254 | 1,203 | 0,105 | 277 | 1,177 |
| BKK29805 | *ytpS* | 2,254 | 1,250 | 0,055 | 36 | 1,222 |
| BKK36050 | *cotB* | 2,252 | 1,184 | 0,094 | 106 | 1,158 |
| BKK31970 | *dhbB* | 2,25 | 1,174 | 0,110 | 112 | 1,148 |
| BKK06540 | *yezC* | 2,248 | 1,205 | 0,130 | 198 | 1,179 |
| BKK17830 | *yndM* | 2,243 | 1,190 | 0,110 | 80 | 1,164 |
| BKK25780 | *arsC* | 2,241 | 1,177 | 0,122 | 361 | 1,151 |
| BKK07780 | *yfkR* | 2,238 | 1,185 | 0,122 | 258 | 1,159 |
| BKK12140 | *yjgA* | 2,234 | 1,216 | 0,159 | 160 | 1,189 |
| BKK23530 | *spoIIM* | 2,231 | 1,179 | 0,115 | 310 | 1,153 |
| BKK35780 | *lytD* | 2,227 | 1,175 | 0,123 | 251 | 1,149 |

| BKK name [1] | gene | screening delta[2] (%) | erage width (µ | +/- | nb | ADP |
|---|---|---|---|---|---|---|
| BKK40700 | *yybB* | 2,224 | 1,211 | 0,092 | 101 | 1,184 |
| BKK34470 | *yveA* | 2,223 | 1,187 | 0,108 | 461 | 1,161 |
| BKK04020 | *yczH* | 2,219 | 1,191 | 0,079 | 160 | 1,165 |
| BKK24600 | *sinI* | 2,218 | 0,997 | 0,082 | 103 | 0,975 |
| BKK07660 | *yflJ* | 2,217 | 1,184 | 0,131 | 307 | 1,159 |
| BKK34930 | *hisZ* | 2,217 | 1,192 | 0,109 | 191 | 1,166 |
| BKK04190 | *ydaD* | 2,216 | 1,191 | 0,106 | 383 | 1,165 |
| BKK10990 | *yitH* | 2,214 | 1,209 | 0,122 | 132 | 1,183 |
| BKK01510 | *ybaJ* | 2,211 | 0,997 | 0,066 | 53 | 0,975 |
| BKK12590 | *xkdE* | 2,21 | 1,215 | 0,188 | 155 | 1,189 |
| BKK15320 | *sigE* | 2,207 | 1,205 | 0,122 | 164 | 1,179 |
| BKK05430 | *ydfJ* | 2,205 | 1,228 | 0,096 | 75 | 1,201 |
| BKK04400 | *gsiB* | 2,204 | 1,191 | 0,094 | 188 | 1,165 |
| BKK32620 | *yurQ* | 2,203 | 1,159 | 0,109 | 110 | 1,134 |
| BKK19380 | *yojO* | 2,202 | 1,154 | 0,125 | 293 | 1,129 |
| BKK22100 | *kdgA* | 2,202 | 1,125 | 0,123 | 117 | 1,100 |
| BKK34330 | *epsE* | 2,202 | 1,192 | 0,111 | 214 | 1,166 |
| BKK17110 | *pksD* | 2,2 | 1,189 | 0,106 | 174 | 1,164 |
| BKK04030 | *ycsD* | 2,19 | 1,185 | 0,117 | 345 | 1,160 |
| BKK31980 | *dhbE* | 2,188 | 1,187 | 0,105 | 422 | 1,161 |
| BKK25770 | *spoIVCA* | 2,185 | 1,249 | 0,090 | 124 | 1,222 |
| BKK37980 | *ywdF* | 2,183 | 1,143 | 0,114 | 141 | 1,119 |
| BKK31510 | *yufK* | 2,182 | 1,164 | 0,093 | 67 | 1,139 |
| BKK12890 | *ykcC* | 2,176 | 1,235 | 0,151 | 56 | 1,208 |
| BKK06580 | *yerC* | 2,169 | 1,204 | 0,111 | 334 | 1,179 |
| BKK13000 | *ykfD* | 2,169 | 1,188 | 0,117 | 283 | 1,163 |
| BKK40574 | *yyzJ* | 2,168 | 1,181 | 0,098 | 139 | 1,156 |
| BKK13880 | *glcT* | 2,165 | 1,192 | 0,127 | 241 | 1,166 |
| BKK38610 | *yxzF* | 2,163 | 1,190 | 0,099 | 56 | 1,165 |
| BKK13480 | *ykrK* | 2,162 | 1,192 | 0,148 | 66 | 1,166 |
| BKK39200 | *yxzC* | 2,157 | 1,190 | 0,131 | 96 | 1,165 |
| BKK37450 | *ywhK* | 2,156 | 1,172 | 0,113 | 303 | 1,147 |
| BKK01450 | *ybxA* | 2,154 | 1,189 | 0,096 | 175 | 1,164 |
| BKK08540 | *yfhI* | 2,153 | 1,234 | 0,155 | 165 | 1,208 |
| BKK08899 | *ygzD* | 2,146 | 1,202 | 0,086 | 107 | 1,177 |
| BKK34940 | *yvpB* | 2,146 | 1,159 | 0,105 | 429 | 1,135 |
| BKK39150 | *yxiJ* | 2,144 | 1,190 | 0,132 | 66 | 1,165 |
| BKK11799 | *yjzK* | 2,142 | 1,214 | 0,086 | 338 | 1,189 |
| BKK37550 | *ywhA* | 2,139 | 1,143 | 0,103 | 168 | 1,119 |
| BKK08160 | *yfjB* | 2,138 | 1,184 | 0,112 | 284 | 1,159 |
| BKK37210 | *ywjC* | 2,138 | 1,143 | 0,110 | 111 | 1,119 |
| BKK23950 | *yqjA* | 2,137 | 1,178 | 0,120 | 335 | 1,153 |
| BKK28490 | *uvrC* | 2,136 | 0,996 | 0,066 | 43 | 0,975 |
| BKK24570 | *gcvT* | 2,135 | 1,212 | 0,095 | 102 | 1,187 |
| BKK31410 | *yugG* | 2,135 | 1,172 | 0,097 | 63 | 1,148 |
| BKK10720 | *gerPA* | 2,132 | 1,209 | 0,099 | 170 | 1,183 |
| BKK05180 | *ydeF* | 2,123 | 1,192 | 0,100 | 251 | 1,167 |
| BKK07980 | *pdaA* | 2,122 | 1,183 | 0,103 | 419 | 1,159 |
| BKK20990 | *yonU* | 2,12 | 1,179 | 0,111 | 447 | 1,155 |
| BKK07550 | *yflT* | 2,115 | 1,183 | 0,114 | 217 | 1,159 |
| BKK20390 | *yorG* | 2,115 | 1,186 | 0,102 | 481 | 1,162 |
| BKK18010 | *yneN* | 2,113 | 1,188 | 0,137 | 83 | 1,164 |
| BKK20030 | *yosR* | 2,113 | 1,153 | 0,160 | 73 | 1,129 |
| BKK18970 | *yobI* | 2,109 | 1,186 | 0,103 | 478 | 1,162 |
| BKK04390 | *ydbA* | 2,105 | 1,184 | 0,114 | 397 | 1,160 |
| BKK15520 | *pyrAB* | 2,104 | 1,180 | 0,106 | 63 | 1,156 |
| BKK26160 | *yqbC* | 2,1 | 1,201 | 0,123 | 146 | 1,177 |
| BKK07220 | *yetL* | 2,097 | 0,995 | 0,086 | 60 | 0,975 |
| BKK24730 | *comGA* | 2,096 | 1,178 | 0,121 | 248 | 1,153 |
| BKK06860 | *yezE* | 2,092 | 1,204 | 0,127 | 476 | 1,179 |
| BKK38020 | *pdxK* | 2,092 | 1,181 | 0,112 | 101 | 1,157 |
| BKK23150 | *resA* | 2,088 | 1,232 | 0,182 | 124 | 1,206 |
| BKK37480 | *ywhH* | 2,088 | 1,142 | 0,096 | 370 | 1,119 |
| BKK16220 | *fliG* | 2,081 | 1,187 | 0,100 | 283 | 1,163 |
| BKK38560 | *licH* | 2,081 | 1,189 | 0,137 | 35 | 1,165 |
| BKK40160 | *yydH* | 2,079 | 1,191 | 0,145 | 85 | 1,166 |
| BKK06839 | *phrH* | 2,077 | 1,203 | 0,113 | 285 | 1,179 |
| BKK35480 | *yviA* | 2,077 | 1,137 | 0,139 | 71 | 1,114 |
| BKK23610 | *nudF* | 2,075 | 1,212 | 0,090 | 174 | 1,187 |
| BKK15960 | *ylqB* | 2,074 | 1,142 | 0,113 | 31 | 1,119 |
| BKK31550 | *yufO* | 2,072 | 1,247 | 0,071 | 88 | 1,222 |
| BKK08230 | *catD* | 2,071 | 1,233 | 0,130 | 100 | 1,208 |
| BKK13340 | *ykoM* | 2,065 | 1,190 | 0,094 | 172 | 1,166 |
| BKK02500 | *ycbG* | 2,062 | 1,184 | 0,113 | 369 | 1,160 |
| BKK02160 | *ybfA* | 2,059 | 1,153 | 0,108 | 167 | 1,130 |
| BKK09570 | *yhdR* | 2,057 | 1,201 | 0,114 | 78 | 1,177 |
| BKK00440 | *veg* | 2,054 | 1,187 | 0,102 | 171 | 1,164 |
| BKK40730 | *yyaS* | 2,054 | 1,209 | 0,101 | 132 | 1,184 |
| BKK02470 | *ycbD* | 2,053 | 1,153 | 0,135 | 95 | 1,130 |

Sup. Table 4: Cell width of mutants of the BKK collection (continued)

| BKK name [1] | gene | screening delta$^2$ (%) | erage width (µ | +/- | nb | ADP |
|---|---|---|---|---|---|---|
| BKK39160 | *yxiI* | 2,052 | 1,189 | 0,110 | 101 | 1,165 |
| BKK17120 | *pksE* | 2,051 | 1,231 | 0,150 | 206 | 1,206 |
| BKK16390 | *flhA* | 2,049 | 1,177 | 0,132 | 387 | 1,153 |
| BKK06970 | *yesO* | 2,048 | 1,226 | 0,081 | 234 | 1,201 |
| BKK25820 | *yqcI* | 2,046 | 1,177 | 0,119 | 230 | 1,153 |
| BKK04940 | *yddE* | 2,045 | 1,191 | 0,118 | 503 | 1,167 |
| BKK29530 | *sppA* | 2,041 | 1,163 | 0,097 | 191 | 1,139 |
| BKK12229 | *yjzI* | 2,039 | 1,213 | 0,089 | 1324 | 1,189 |
| BKK13040 | *hmp* | 2,038 | 1,190 | 0,117 | 451 | 1,166 |
| BKK17230 | *pksS* | 2,036 | 1,187 | 0,115 | 151 | 1,164 |
| BKK27070 | *levD* | 2,036 | 1,201 | 0,093 | 261 | 1,177 |
| BKK11680 | *thiS* | 2,032 | 1,213 | 0,150 | 755 | 1,189 |
| BKK35650 | *lytR* | 2,031 | 1,173 | 0,106 | 529 | 1,149 |
| BKK03000 | *opuAC* | 2,03 | 1,226 | 0,065 | 146 | 1,201 |
| BKK05590 | *ydgD* | 2,027 | 1,191 | 0,077 | 159 | 1,167 |
| BKK16080 | *ylqH* | 2,026 | 1,183 | 0,126 | 259 | 1,160 |
| BKK27310 | *pbpI* | 2,024 | 0,995 | 0,106 | 123 | 0,975 |
| BKK30450 | *ytrB* | 2,023 | 1,174 | 0,117 | 246 | 1,151 |
| BKK26910 | *yraK* | 2,022 | 1,200 | 0,106 | 204 | 1,177 |
| BKK04660 | *ndoA* | 2,02 | 1,188 | 0,092 | 72 | 1,165 |
| BKK36110 | *ywrC* | 2,013 | 1,190 | 0,131 | 93 | 1,166 |
| BKK39470 | *yxeP* | 2,013 | 1,188 | 0,107 | 95 | 1,165 |
| BKK29910 | *ytzH* | 2,011 | 1,135 | 0,135 | 46 | 1,113 |
| BKK26250 | *yqaN* | 2,007 | 1,200 | 0,103 | 195 | 1,177 |
| BKK40540 | *yybR* | 2,006 | 1,179 | 0,105 | 120 | 1,156 |
| BKK12710 | *xkdR* | 2,005 | 1,213 | 0,154 | 142 | 1,189 |
| BKK23470 | *spoIIAA* | 2,001 | 1,231 | 0,157 | 184 | 1,206 |
| BKK26790 | *aadK* | 1,996 | 1,200 | 0,101 | 286 | 1,177 |
| BKK39440 | *yxxB* | 1,993 | 1,231 | 0,109 | 192 | 1,206 |
| BKK40660 | *yybF* | 1,993 | 1,190 | 0,100 | 407 | 1,167 |
| BKK20928 | *yoyH* | 1,992 | 1,178 | 0,110 | 326 | 1,155 |
| BKK14440 | *panE* | 1,99 | 1,203 | 0,116 | 210 | 1,179 |
| BKK22030 | *ypbR* | 1,99 | 1,176 | 0,114 | 377 | 1,153 |
| BKK22720 | *cheR* | 1,99 | 1,176 | 0,120 | 252 | 1,153 |
| BKK27810 | *yrbD* | 1,98 | 1,183 | 0,092 | 212 | 1,160 |
| BKK33221 | *yvrH* | 1,976 | 0,994 | 0,081 | 41 | 0,975 |
| BKK33990 | *yvbU* | 1,976 | 1,157 | 0,126 | 72 | 1,134 |
| BKK03590 | *tcyC* | 1,974 | 1,225 | 0,085 | 153 | 1,201 |
| BKK07130 | *lplD* | 1,974 | 1,202 | 0,118 | 423 | 1,179 |
| BKK17580 | *xynB* | 1,972 | 1,186 | 0,112 | 205 | 1,164 |
| BKK21780 | *yplP* | 1,97 | 1,178 | 0,130 | 274 | 1,155 |
| BKK11190 | *argC* | 1,966 | 1,207 | 0,106 | 73 | 1,183 |
| BKK37750 | *ywfA* | 1,965 | 1,172 | 0,112 | 349 | 1,149 |
| BKK31520 | *yufL* | 1,964 | 1,170 | 0,114 | 329 | 1,148 |
| BKK22880 | *ypfD* | 1,962 | 1,176 | 0,114 | 318 | 1,153 |
| BKK15800 | *thiN* | 1,96 | 1,157 | 0,113 | 318 | 1,135 |
| BKK34780 | *yvcI* | 1,96 | 1,157 | 0,145 | 339 | 1,135 |
| BKK30630 | *ytkD* | 1,959 | 1,179 | 0,090 | 123 | 1,156 |
| BKK24000 | *bmrU* | 1,958 | 1,210 | 0,104 | 414 | 1,187 |
| BKK29220 | *ytsJ* | 1,958 | 1,162 | 0,099 | 103 | 1,139 |
| BKK34070 | *yvfT* | 1,955 | 1,172 | 0,120 | 289 | 1,149 |
| BKK34580 | *yvdJ* | 1,954 | 1,172 | 0,102 | 502 | 1,149 |
| BKK08850 | *ssuC* | 1,95 | 1,232 | 0,132 | 128 | 1,208 |
| BKK03180 | *cah* | 1,948 | 1,182 | 0,124 | 632 | 1,160 |
| BKK03930 | *gdh* | 1,944 | 1,188 | 0,086 | 175 | 1,165 |
| BKK40830 | *yyaK* | 1,942 | 1,207 | 0,103 | 172 | 1,184 |
| BKK33470 | *bdbC* | 1,941 | 1,189 | 0,104 | 160 | 1,166 |
| BKK17490 | *ynzG* | 1,939 | 1,184 | 0,107 | 433 | 1,162 |
| BKK19110 | *yobW* | 1,939 | 1,194 | 0,116 | 89 | 1,172 |
| BKK10130 | *hemH* | 1,938 | 1,178 | 0,105 | 92 | 1,156 |
| BKK36490 | *ywoC* | 1,938 | 1,189 | 0,095 | 81 | 1,166 |
| BKK14130 | *ykuL* | 1,937 | 1,189 | 0,142 | 774 | 1,166 |
| BKK28360 | *ysnA* | 1,937 | 1,173 | 0,118 | 288 | 1,151 |
| BKK40500 | *rplI* | 1,935 | 1,207 | 0,099 | 105 | 1,184 |
| BKK10020 | *serC* | 1,934 | 1,200 | 0,119 | 157 | 1,177 |
| BKK19030 | *yobO* | 1,934 | 1,194 | 0,139 | 98 | 1,172 |
| BKK30680 | *ytjA* | 1,932 | 1,178 | 0,113 | 78 | 1,156 |
| BKK11120 | *yitT* | 1,926 | 1,185 | 0,112 | 223 | 1,163 |
| BKK21170 | *yomZ* | 1,923 | 1,177 | 0,104 | 316 | 1,155 |
| BKK33580 | *yvaF* | 1,922 | 1,156 | 0,125 | 114 | 1,134 |
| BKK22640 | *trpB* | 1,921 | 1,121 | 0,096 | 196 | 1,100 |
| BKK13770 | *ykvO* | 1,919 | 1,189 | 0,130 | 68 | 1,166 |
| BKK21420 | *bhlA* | 1,919 | 1,173 | 0,116 | 216 | 1,151 |
| BKK21800 | *ypkP* | 1,918 | 1,177 | 0,117 | 566 | 1,155 |
| BKK10010 | *trpP* | 1,917 | 1,232 | 0,127 | 46 | 1,208 |
| BKK23810 | *yqjN* | 1,912 | 1,210 | 0,102 | 350 | 1,187 |
| BKK20590 | *yoqL* | 1,91 | 1,177 | 0,122 | 249 | 1,155 |
| BKK00600 | *yabP* | 1,909 | 1,169 | 0,107 | 550 | 1,147 |
| BKK13690 | *motA* | 1,898 | 1,189 | 0,102 | 117 | 1,166 |

| BKK name [1] | gene | screening delta$^2$ (%) | erage width (µ | +/- | nb | ADP |
|---|---|---|---|---|---|---|
| BKK23540 | *yqkK* | 1,898 | 1,210 | 0,110 | 283 | 1,187 |
| BKK03200 | *ycgM* | 1,892 | 1,151 | 0,119 | 36 | 1,130 |
| BKK10360 | *yhfT* | 1,891 | 1,206 | 0,101 | 217 | 1,183 |
| BKK32630 | *yurR* | 1,889 | 1,156 | 0,092 | 150 | 1,134 |
| BKK18490 | *rtp* | 1,888 | 1,184 | 0,126 | 266 | 1,162 |
| BKK11900 | *yjcL* | 1,885 | 1,184 | 0,120 | 287 | 1,163 |
| BKK19340 | *yocR* | 1,884 | 1,179 | 0,095 | 155 | 1,158 |
| BKK27730 | *ruvB* | 1,884 | 1,181 | 0,107 | 314 | 1,160 |
| BKK14990 | *ylbF* | 1,883 | 1,201 | 0,131 | 615 | 1,179 |
| BKK28250 | *leuD* | 1,88 | 1,181 | 0,099 | 224 | 1,160 |
| BKK35170 | *uvrB* | 1,876 | 1,156 | 0,117 | 87 | 1,135 |
| BKK32990 | *mrgA* | 1,873 | 1,156 | 0,091 | 90 | 1,134 |
| BKK10780 | *yisN* | 1,871 | 1,205 | 0,130 | 152 | 1,183 |
| BKK13780 | *ykvP* | 1,871 | 1,188 | 0,121 | 416 | 1,166 |
| BKK35950 | *rbsC* | 1,867 | 1,188 | 0,105 | 116 | 1,166 |
| BKK24140 | *prpC* | 1,865 | 1,209 | 0,102 | 659 | 1,187 |
| BKK26660 | *yrdN* | 1,857 | 1,199 | 0,117 | 403 | 1,177 |
| BKK38630 | *katX* | 1,855 | 1,186 | 0,131 | 34 | 1,165 |
| BKK04860 | *ydcQ* | 1,852 | 1,186 | 0,121 | 95 | 1,165 |
| BKK29540 | *ppnKB* | 1,851 | 1,161 | 0,089 | 140 | 1,139 |
| BKK06440 | *purB* | 1,849 | 1,201 | 0,124 | 179 | 1,179 |
| BKK19470 | *yojF* | 1,847 | 1,190 | 0,118 | 412 | 1,168 |
| BKK27220 | *yrhE* | 1,843 | 1,198 | 0,100 | 228 | 1,177 |
| BKK14300 | *moaE* | 1,838 | 1,201 | 0,107 | 880 | 1,179 |
| BKK17760 | *yndE* | 1,836 | 1,185 | 0,113 | 206 | 1,164 |
| BKK39270 | *bglP* | 1,835 | 1,186 | 0,113 | 79 | 1,165 |
| BKK01090 | *rplGB* | 1,832 | 1,185 | 0,109 | 249 | 1,164 |
| BKK05420 | *ydfI* | 1,824 | 1,223 | 0,116 | 68 | 1,201 |
| BKK07330 | *yfnB* | 1,818 | 1,200 | 0,112 | 548 | 1,179 |
| BKK13830 | *ykvU* | 1,816 | 1,184 | 0,115 | 326 | 1,163 |
| BKK31390 | *yugI* | 1,816 | 1,169 | 0,105 | 232 | 1,148 |
| BKK00970 | *yacP* | 1,815 | 1,181 | 0,123 | 307 | 1,160 |
| BKK10740 | *yisJ* | 1,814 | 1,205 | 0,113 | 75 | 1,183 |
| BKK02480 | *gudP* | 1,81 | 1,223 | 0,069 | 197 | 1,201 |
| BKK32790 | *yusG* | 1,81 | 1,155 | 0,095 | 171 | 1,134 |
| BKK13800 | *ykvR* | 1,804 | 1,187 | 0,132 | 58 | 1,166 |
| BKK05770 | *ydhI* | 1,798 | 1,188 | 0,110 | 439 | 1,167 |
| BKK02490 | *gudD* | 1,797 | 1,150 | 0,123 | 43 | 1,130 |
| BKK25930 | *yqcE* | 1,796 | 1,172 | 0,104 | 256 | 1,151 |
| BKK31050 | *gbsB* | 1,796 | 1,177 | 0,126 | 106 | 1,156 |
| BKK33480 | *bdbD* | 1,796 | 1,155 | 0,095 | 157 | 1,134 |
| BKK17590 | *xylR* | 1,79 | 1,184 | 0,113 | 81 | 1,164 |
| BKK04520 | *ydbM* | 1,788 | 1,180 | 0,115 | 696 | 1,160 |
| BKK21599 | *yoyK* | 1,788 | 1,176 | 0,107 | 459 | 1,155 |
| BKK40370 | *yycJ* | 1,785 | 1,150 | 0,098 | 131 | 1,130 |
| BKK40610 | *yybK* | 1,784 | 1,187 | 0,121 | 101 | 1,166 |
| BKK15560 | *pyrE* | 1,783 | 1,160 | 0,092 | 111 | 1,139 |
| BKK38830 | *aldY* | 1,781 | 1,185 | 0,118 | 77 | 1,165 |
| BKK38950 | *yxjH* | 1,78 | 1,178 | 0,094 | 70 | 1,158 |
| BKK03610 | *tcyA* | 1,775 | 1,168 | 0,116 | 422 | 1,147 |
| BKK16140 | *codV* | 1,77 | 1,205 | 0,125 | 65 | 1,184 |
| BKK19140 | *yozB* | 1,768 | 1,182 | 0,116 | 362 | 1,162 |
| BKK08070 | *acoB* | 1,767 | 1,179 | 0,118 | 417 | 1,159 |
| BKK09310 | *pgcA* | 1,767 | 1,230 | 0,127 | 38 | 1,208 |
| BKK02710 | *yczC* | 1,765 | 1,223 | 0,067 | 82 | 1,201 |
| BKK36150 | *ywqN* | 1,764 | 1,134 | 0,087 | 54 | 1,114 |
| BKK40840 | *yyaJ* | 1,764 | 1,188 | 0,117 | 192 | 1,167 |
| BKK04420 | *ydbC* | 1,763 | 1,185 | 0,100 | 263 | 1,165 |
| BKK02170 | *ybfB* | 1,758 | 1,168 | 0,120 | 442 | 1,147 |
| BKK39280 | *yxxE* | 1,757 | 1,185 | 0,122 | 80 | 1,165 |
| BKK14010 | *cheV* | 1,754 | 1,183 | 0,123 | 333 | 1,163 |
| BKK02310 | *ybfO* | 1,752 | 1,150 | 0,097 | 86 | 1,130 |
| BKK08830 | *ssuB* | 1,745 | 1,229 | 0,167 | 96 | 1,208 |
| BKK13530 | *kinE* | 1,743 | 1,188 | 0,103 | 359 | 1,168 |
| BKK18690 | *yoaP* | 1,743 | 1,182 | 0,114 | 392 | 1,162 |
| BKK10610 | *yhjR* | 1,74 | 1,204 | 0,108 | 150 | 1,183 |
| BKK21240 | *yomS* | 1,739 | 1,175 | 0,123 | 411 | 1,155 |
| BKK29430 | *ytzD* | 1,739 | 1,159 | 0,092 | 208 | 1,139 |
| BKK29610 | *ezrA* | 1,739 | 1,132 | 0,116 | 65 | 1,113 |
| BKK03100 | *ycgG* | 1,737 | 1,150 | 0,107 | 110 | 1,130 |
| BKK02000 | *ybdJ* | 1,735 | 1,150 | 0,097 | 72 | 1,130 |
| BKK04490 | *ydbJ* | 1,734 | 1,222 | 0,098 | 109 | 1,201 |
| BKK07490 | *yfmF* | 1,733 | 1,179 | 0,094 | 296 | 1,159 |
| BKK10890 | *yisX* | 1,732 | 1,204 | 0,120 | 111 | 1,183 |
| BKK05320 | *ydeS* | 1,726 | 1,187 | 0,122 | 143 | 1,167 |
| BKK06700 | *yerO* | 1,725 | 1,199 | 0,126 | 373 | 1,179 |
| BKK06073 | *ydzW* | 1,722 | 1,167 | 0,098 | 121 | 1,147 |
| BKK29700 | *acuB* | 1,721 | 1,178 | 0,105 | 142 | 1,158 |
| BKK24210 | *yqiG* | 1,719 | 1,207 | 0,145 | 583 | 1,187 |

| BKK name [1] | gene | screening delta$^2$ (%) | erage width (µ | +/- | nb | ADP |
|---|---|---|---|---|---|---|
| BKK13120 | *proB* | 1,714 | 1,186 | 0,123 | 109 | 1,166 |
| BKK28740 | *araP* | 1,713 | 1,159 | 0,102 | 107 | 1,139 |
| BKK04700 | *rsbU* | 1,71 | 1,185 | 0,115 | 93 | 1,165 |
| BKK10500 | *yhjG* | 1,71 | 1,204 | 0,102 | 100 | 1,183 |
| BKK27350 | *yrrN* | 1,698 | 1,171 | 0,123 | 232 | 1,151 |
| BKK29490 | *tpx* | 1,695 | 1,132 | 0,120 | 128 | 1,113 |
| BKK40578 | *yyzK* | 1,692 | 1,186 | 0,114 | 161 | 1,166 |
| BKK04170 | *ydaB* | 1,687 | 1,185 | 0,097 | 343 | 1,165 |
| BKK22160 | *yptA* | 1,68 | 1,119 | 0,097 | 30 | 1,100 |
| BKK03060 | *lctP* | 1,674 | 1,149 | 0,121 | 51 | 1,130 |
| BKK19689 | *yokU* | 1,674 | 1,188 | 0,106 | 330 | 1,168 |
| BKK24290 | *xseB* | 1,672 | 1,207 | 0,112 | 260 | 1,187 |
| BKK10600 | *yhjQ* | 1,665 | 1,203 | 0,120 | 85 | 1,183 |
| BKK30940 | *glgP* | 1,665 | 1,175 | 0,103 | 132 | 1,156 |
| BKK09400 | *spoVR* | 1,663 | 1,228 | 0,112 | 201 | 1,208 |
| BKK16950 | *pbpX* | 1,661 | 1,227 | 0,160 | 62 | 1,206 |
| BKK21380 | *yomF* | 1,661 | 1,174 | 0,125 | 384 | 1,155 |
| BKK18410 | *ggt* | 1,66 | 1,157 | 0,097 | 89 | 1,138 |
| BKK20090 | *yosK* | 1,656 | 1,148 | 0,113 | 70 | 1,129 |
| BKK28300 | *ilvH* | 1,656 | 1,179 | 0,096 | 313 | 1,160 |
| BKK04180 | *ydaC* | 1,655 | 1,184 | 0,087 | 146 | 1,165 |
| BKK16170 | *codY* | 1,649 | 1,182 | 0,138 | 230 | 1,163 |
| BKK39350 | *hutH* | 1,647 | 1,184 | 0,115 | 102 | 1,165 |
| BKK09260 | *yhxA* | 1,643 | 1,182 | 0,104 | 262 | 1,163 |
| BKK09120 | *yhcK* | 1,636 | 1,182 | 0,108 | 295 | 1,163 |
| BKK30840 | *yteA* | 1,632 | 1,175 | 0,116 | 117 | 1,156 |
| BKK06890 | *cotJA* | 1,631 | 1,198 | 0,113 | 433 | 1,179 |
| BKK25190 | *cccA* | 1,63 | 1,172 | 0,111 | 473 | 1,153 |
| BKK39680 | *iolI* | 1,621 | 1,166 | 0,106 | 353 | 1,147 |
| BKK06500 | *purM* | 1,62 | 1,198 | 0,121 | 369 | 1,179 |
| BKK36060 | *cotH* | 1,619 | 1,168 | 0,118 | 334 | 1,149 |
| BKK36550 | *spoIIQ* | 1,619 | 1,168 | 0,118 | 334 | 1,149 |
| BKK35690 | *ggaA* | 1,617 | 1,176 | 0,105 | 68 | 1,158 |
| BKK04370 | *ydaS* | 1,61 | 1,184 | 0,094 | 466 | 1,165 |
| BKK08550 | *sspK* | 1,601 | 1,177 | 0,114 | 318 | 1,159 |
| BKK10340 | *phoE* | 1,601 | 1,202 | 0,115 | 140 | 1,183 |
| BKK17460 | *glnA* | 1,601 | 1,182 | 0,115 | 59 | 1,164 |
| BKK24540 | *yqhL* | 1,6 | 1,206 | 0,110 | 223 | 1,187 |
| BKK06330 | *yeaC* | 1,596 | 1,198 | 0,120 | 155 | 1,179 |
| BKK27140 | *oatA* | 1,596 | 1,242 | 0,068 | 82 | 1,222 |
| BKK32860 | *yusN* | 1,596 | 1,152 | 0,102 | 110 | 1,134 |
| BKK10970 | *yitF* | 1,591 | 1,202 | 0,119 | 107 | 1,183 |
| BKK20180 | *yosB* | 1,589 | 1,180 | 0,114 | 343 | 1,162 |
| BKK35840 | *ywtF* | 1,589 | 1,132 | 0,100 | 277 | 1,114 |
| BKK00220 | *yaaL* | 1,586 | 1,202 | 0,124 | 98 | 1,183 |
| BKK17040 | *mutS* | 1,584 | 1,182 | 0,112 | 258 | 1,164 |
| BKK39060 | *citH* | 1,584 | 1,183 | 0,114 | 87 | 1,165 |
| BKK39290 | *yxxD* | 1,584 | 1,185 | 0,121 | 66 | 1,166 |
| BKK17980 | *sspP* | 1,582 | 1,182 | 0,111 | 159 | 1,164 |
| BKK05870 | *gmuF* | 1,581 | 1,186 | 0,116 | 431 | 1,167 |
| BKK22180 | *gpsB* | 1,577 | 1,118 | 0,115 | 76 | 1,100 |
| BKK06048 | *ydzU* | 1,574 | 1,186 | 0,113 | 280 | 1,167 |
| BKK30580 | *ytmA* | 1,574 | 1,174 | 0,085 | 99 | 1,156 |
| BKK31540 | *yufN* | 1,571 | 1,241 | 0,082 | 57 | 1,222 |
| BKK32980 | *yusZ* | 1,569 | 1,180 | 0,115 | 395 | 1,161 |
| BKK09350 | *yhdB* | 1,568 | 1,195 | 0,105 | 280 | 1,177 |
| BKK05700 | *ydhC* | 1,558 | 1,185 | 0,075 | 162 | 1,167 |
| BKK03920 | *glcU* | 1,555 | 1,220 | 0,082 | 32 | 1,201 |
| BKK34680 | *yvcT* | 1,554 | 1,153 | 0,107 | 60 | 1,135 |
| BKK19050 | *yobQ* | 1,551 | 1,190 | 0,106 | 142 | 1,172 |
| BKK29330 | *ytmO* | 1,551 | 1,130 | 0,108 | 791 | 1,113 |
| BKK12340 | *uxuA* | 1,549 | 1,207 | 0,092 | 1324 | 1,189 |
| BKK11170 | *yitY* | 1,547 | 1,202 | 0,098 | 110 | 1,183 |
| BKK30330 | *ytvB* | 1,547 | 1,174 | 0,089 | 111 | 1,156 |
| BKK23900 | *yqjF* | 1,546 | 1,225 | 0,164 | 94 | 1,206 |
| BKK09650 | *yhdZ* | 1,545 | 1,195 | 0,110 | 122 | 1,177 |
| BKK17990 | *sspO* | 1,543 | 1,180 | 0,124 | 356 | 1,162 |
| BKK39740 | *iolC* | 1,542 | 0,990 | 0,076 | 90 | 0,975 |
| BKK28670 | *ysfB* | 1,541 | 1,157 | 0,096 | 170 | 1,139 |
| BKK34250 | *epsL* | 1,538 | 1,153 | 0,093 | 142 | 1,135 |
| BKK07870 | *yfkK* | 1,537 | 1,180 | 0,120 | 342 | 1,163 |
| BKK09420 | *lytE* | 1,537 | 1,227 | 0,115 | 39 | 1,208 |
| BKK16590 | *ylxS* | 1,535 | 1,179 | 0,127 | 326 | 1,161 |
| BKK22869 | *ypzI* | 1,535 | 1,117 | 0,088 | 110 | 1,100 |
| BKK08650 | *fabL* | 1,533 | 1,177 | 0,155 | 135 | 1,159 |
| BKK40130 | *yydK* | 1,531 | 1,165 | 0,095 | 388 | 1,147 |
| BKK12960 | *dppE* | 1,529 | 1,155 | 0,093 | 140 | 1,138 |
| BKK09380 | *nsrR* | 1,528 | 1,195 | 0,104 | 120 | 1,177 |
| BKK08029 | *yfzA* | 1,524 | 1,176 | 0,110 | 516 | 1,159 |

| BKK name [1] | gene | screening delta$^2$ (%) | erage width (µ | +/- | nb | ADP |
|---|---|---|---|---|---|---|
| BKK05230 | *ydeK* | 1,523 | 1,157 | 0,085 | 95 | 1,139 |
| BKK27610 | *apt* | 1,518 | 1,177 | 0,098 | 142 | 1,160 |
| BKK02090 | *ybxI* | 1,516 | 1,147 | 0,129 | 111 | 1,130 |
| BKK08760 | *spo0M* | 1,514 | 1,227 | 0,147 | 248 | 1,208 |
| BKK37590 | *ywgA* | 1,514 | 1,136 | 0,131 | 72 | 1,119 |
| BKK25010 | *yqgE* | 1,512 | 1,225 | 0,116 | 49 | 1,206 |
| BKK29840 | *ytpP* | 1,511 | 1,157 | 0,103 | 98 | 1,139 |
| BKK23050 | *fmnP* | 1,509 | 1,117 | 0,103 | 148 | 1,100 |
| BKK00210 | *recR* | 1,502 | 1,165 | 0,163 | 503 | 1,147 |
| BKK21990 | *ypdQ* | 1,502 | 1,172 | 0,103 | 484 | 1,155 |
| BKK06910 | *cotJC* | 1,5 | 1,197 | 0,115 | 261 | 1,179 |
| BKK35420 | *yvyG* | 1,5 | 1,131 | 0,108 | 862 | 1,114 |
| BKK40950 | *yyaC* | 1,498 | 1,202 | 0,096 | 139 | 1,184 |
| BKK04840 | *ydcO* | 1,495 | 1,177 | 0,131 | 359 | 1,160 |
| BKK24610 | *sinR* | 1,492 | 1,202 | 0,088 | 162 | 1,184 |
| BKK11660 | *tenI* | 1,49 | 1,207 | 0,118 | 264 | 1,189 |
| BKK30660 | *ytkA* | 1,486 | 1,240 | 0,055 | 32 | 1,222 |
| BKK12970 | *ykfA* | 1,484 | 1,226 | 0,139 | 121 | 1,208 |
| BKK33450 | *helD* | 1,476 | 1,151 | 0,085 | 97 | 1,134 |
| BKK05620 | *ydgF* | 1,475 | 1,177 | 0,113 | 690 | 1,160 |
| BKK19630 | *deoD* | 1,472 | 1,146 | 0,119 | 548 | 1,129 |
| BKK19430 | *yojJ* | 1,469 | 1,146 | 0,132 | 269 | 1,129 |
| BKK20820 | *yopO* | 1,465 | 1,172 | 0,106 | 450 | 1,155 |
| BKK13010 | *ykgB* | 1,462 | 1,183 | 0,126 | 555 | 1,166 |
| BKK10710 | *gerPB* | 1,461 | 1,201 | 0,117 | 127 | 1,183 |
| BKK16800 | *spoIIIE* | 1,461 | 1,224 | 0,175 | 62 | 1,206 |
| BKK26810 | *yrpC* | 1,454 | 1,240 | 0,084 | 165 | 1,222 |
| BKK01660 | *ybbD* | 1,451 | 1,219 | 0,083 | 66 | 1,201 |
| BKK11390 | *appB* | 1,451 | 1,226 | 0,162 | 190 | 1,208 |
| BKK06150 | *gutB* | 1,447 | 1,196 | 0,126 | 82 | 1,179 |
| BKK32160 | *yutM* | 1,446 | 1,178 | 0,110 | 434 | 1,161 |
| BKK33720 | *opuBB* | 1,445 | 1,183 | 0,114 | 181 | 1,166 |
| BKK32500 | *pucB* | 1,444 | 1,183 | 0,114 | 156 | 1,166 |
| BKK32300 | *yutE* | 1,441 | 1,178 | 0,113 | 384 | 1,161 |
| BKK35810 | *gerBB* | 1,435 | 1,130 | 0,116 | 31 | 1,114 |
| BKK25860 | *yqcG* | 1,431 | 1,168 | 0,107 | 300 | 1,151 |
| BKK27280 | *yrrT* | 1,431 | 0,989 | 0,085 | 47 | 0,975 |
| BKK33590 | *yvaG* | 1,43 | 1,151 | 0,107 | 178 | 1,134 |
| BKK26020 | *yqbP* | 1,427 | 1,193 | 0,122 | 119 | 1,177 |
| BKK19120 | *czrA* | 1,426 | 1,188 | 0,109 | 157 | 1,172 |
| BKK40630 | *yybI* | 1,426 | 1,201 | 0,109 | 194 | 1,184 |
| BKK07570 | *yflS* | 1,422 | 1,226 | 0,140 | 73 | 1,208 |
| BKK34240 | *epsM* | 1,421 | 1,151 | 0,108 | 130 | 1,135 |
| BKK08530 | *yfhH* | 1,417 | 1,175 | 0,106 | 257 | 1,159 |
| BKK17220 | *pksR* | 1,415 | 1,180 | 0,108 | 157 | 1,164 |
| BKK28910 | *lrgA* | 1,404 | 1,178 | 0,108 | 203 | 1,161 |
| BKK34080 | *yvfS* | 1,404 | 1,166 | 0,097 | 241 | 1,149 |
| BKK40120 | *yyzE* | 1,398 | 0,989 | 0,063 | 46 | 0,975 |
| BKK06740 | *yefB* | 1,394 | 1,195 | 0,113 | 251 | 1,179 |
| BKK28040 | *radC* | 1,394 | 1,169 | 0,118 | 183 | 1,153 |
| BKK35860 | *pgdS* | 1,393 | 1,165 | 0,103 | 527 | 1,149 |
| BKK34120 | *ganB* | 1,391 | 1,177 | 0,106 | 416 | 1,161 |
| BKK14310 | *moaD* | 1,386 | 1,195 | 0,107 | 574 | 1,179 |
| BKK24430 | *spoIIIAA* | 1,386 | 1,167 | 0,113 | 452 | 1,151 |
| BKK40590 | *yybM* | 1,383 | 1,182 | 0,107 | 186 | 1,166 |
| BKK08950 | *yhbE* | 1,382 | 1,225 | 0,165 | 85 | 1,208 |
| BKK14009 | *ykzT* | 1,382 | 1,182 | 0,130 | 124 | 1,166 |
| BKK09000 | *yhbJ* | 1,381 | 1,193 | 0,123 | 131 | 1,177 |
| BKK27820 | *yrbC* | 1,38 | 1,169 | 0,114 | 470 | 1,153 |
| BKK25370 | *yqfB* | 1,378 | 0,988 | 0,083 | 49 | 0,975 |
| BKK07440 | *yfmK* | 1,372 | 1,175 | 0,106 | 308 | 1,159 |
| BKK32810 | *yusI* | 1,366 | 1,150 | 0,103 | 65 | 1,134 |
| BKK37030 | *racA* | 1,362 | 1,173 | 0,121 | 116 | 1,157 |
| BKK24820 | *yqgU* | 1,361 | 1,179 | 0,106 | 322 | 1,164 |
| BKK03830 | *yclQ* | 1,36 | 1,218 | 0,091 | 69 | 1,201 |
| BKK25050 | *yqgA* | 1,36 | 1,169 | 0,115 | 206 | 1,153 |
| BKK36560 | *ywnH* | 1,36 | 1,182 | 0,103 | 162 | 1,166 |
| BKK05370 | *ydfD* | 1,355 | 1,175 | 0,128 | 409 | 1,160 |
| BKK01590 | *ybaS* | 1,353 | 1,153 | 0,107 | 123 | 1,138 |
| BKK22070 | *xpt* | 1,352 | 1,155 | 0,112 | 112 | 1,139 |
| BKK02520 | *ycbJ* | 1,348 | 1,145 | 0,089 | 215 | 1,130 |
| BKK08980 | *yhbH* | 1,347 | 1,193 | 0,106 | 183 | 1,177 |
| BKK29290 | *ytnL* | 1,346 | 1,155 | 0,092 | 133 | 1,139 |
| BKK05100 | *yddT* | 1,34 | 1,183 | 0,112 | 351 | 1,167 |
| BKK32440 | *pucK* | 1,339 | 1,163 | 0,110 | 68 | 1,148 |
| BKK17600 | *xylA* | 1,333 | 1,179 | 0,138 | 65 | 1,164 |
| BKK26089 | *yqzN* | 1,329 | 1,193 | 0,118 | 191 | 1,177 |
| BKK31830 | *yueE* | 1,328 | 1,163 | 0,111 | 251 | 1,148 |
| BKK19650 | *yodM* | 1,325 | 1,144 | 0,118 | 1093 | 1,129 |

### Sup. Table 4: Cell width of mutants of the BKK collection (continued)

| BKK name [1] | gene | screening delta[2] (%) | erage width (µ | +/- | nb | ADP |
|---|---|---|---|---|---|---|
| BKK02630 | *tatAD* | 1,323 | 1,217 | 0,079 | 43 | 1,201 |
| BKK21920 | *ugtP* | 1,322 | 1,222 | 0,162 | 73 | 1,206 |
| BKK13230 | *thiV* | 1,317 | 1,224 | 0,144 | 59 | 1,208 |
| BKK11710 | *thiD* | 1,316 | 1,205 | 0,097 | 800 | 1,189 |
| BKK33980 | *yvbT* | 1,311 | 1,177 | 0,117 | 449 | 1,161 |
| BKK35400 | *flgL* | 1,309 | 1,150 | 0,127 | 456 | 1,135 |
| BKK35990 | *ywrO* | 1,308 | 1,129 | 0,093 | 92 | 1,114 |
| BKK26370 | *yqaC* | 1,306 | 1,192 | 0,098 | 415 | 1,177 |
| BKK07190 | *yezD* | 1,304 | 1,194 | 0,120 | 279 | 1,179 |
| BKK14290 | *mobB* | 1,304 | 1,195 | 0,098 | 513 | 1,179 |
| BKK41020 | *trmE* | 1,303 | 1,175 | 0,113 | 189 | 1,160 |
| BKK29860 | *ytoP* | 1,301 | 1,128 | 0,120 | 71 | 1,113 |
| BKK10320 | *yhfP* | 1,298 | 1,178 | 0,106 | 390 | 1,163 |
| BKK22020 | *ypbS* | 1,298 | 1,115 | 0,131 | 151 | 1,100 |
| BKK15440 | *ylyA* | 1,297 | 1,154 | 0,088 | 127 | 1,139 |
| BKK24960 | *pstBA* | 1,293 | 1,222 | 0,123 | 132 | 1,206 |
| BKK05150 | *ydeC* | 1,292 | 1,173 | 0,117 | 96 | 1,158 |
| BKK24530 | *yqhM* | 1,29 | 1,202 | 0,120 | 251 | 1,187 |
| BKK19360 | *odhB* | 1,287 | 1,144 | 0,121 | 549 | 1,129 |
| BKK37090 | *ywjI* | 1,282 | 1,162 | 0,108 | 254 | 1,147 |
| BKK38260 | *efeN* | 1,277 | 1,181 | 0,110 | 63 | 1,166 |
| BKK02250 | *ybfJ* | 1,275 | 1,217 | 0,068 | 77 | 1,201 |
| BKK35310 | *yvyD* | 1,272 | 1,150 | 0,124 | 178 | 1,135 |
| BKK22090 | *kdgT* | 1,27 | 1,222 | 0,126 | 196 | 1,206 |
| BKK30620 | *ytlD* | 1,269 | 1,238 | 0,081 | 34 | 1,222 |
| BKK03530 | *ycxA* | 1,267 | 1,217 | 0,115 | 381 | 1,201 |
| BKK18180 | *yngB* | 1,266 | 1,222 | 0,167 | 172 | 1,206 |
| BKK04040 | *ycsE* | 1,264 | 1,174 | 0,120 | 687 | 1,160 |
| BKK35970 | *ywsB* | 1,263 | 1,128 | 0,143 | 32 | 1,114 |
| BKK24520 | *mntR* | 1,262 | 1,168 | 0,118 | 145 | 1,153 |
| BKK05529 | *ydzR* | 1,26 | 0,987 | 0,076 | 37 | 0,975 |
| BKK15110 | *ylbQ* | 1,26 | 1,194 | 0,129 | 310 | 1,179 |
| BKK15100 | *ylbP* | 1,259 | 1,171 | 0,107 | 130 | 1,156 |
| BKK19060 | *yobR* | 1,258 | 1,199 | 0,139 | 119 | 1,184 |
| BKK05570 | *ydgB* | 1,25 | 1,182 | 0,127 | 169 | 1,167 |
| BKK14960 | *ylbC* | 1,249 | 1,194 | 0,116 | 99 | 1,179 |
| BKK11090 | *yitR* | 1,244 | 1,177 | 0,102 | 427 | 1,163 |
| BKK29620 | *hisJ* | 1,242 | 1,127 | 0,106 | 116 | 1,113 |
| BKK30160 | *ytcQ* | 1,242 | 1,237 | 0,132 | 33 | 1,222 |
| BKK20200 | *yorZ* | 1,238 | 1,176 | 0,118 | 242 | 1,162 |
| BKK21360 | *yomH* | 1,238 | 1,169 | 0,108 | 660 | 1,155 |
| BKK30190 | *bioI* | 1,237 | 1,170 | 0,111 | 146 | 1,156 |
| BKK36940 | *ywlD* | 1,237 | 0,987 | 0,067 | 60 | 0,975 |
| BKK18470 | *proJ* | 1,233 | 1,186 | 0,107 | 362 | 1,172 |
| BKK21560 | *yokK* | 1,232 | 1,168 | 0,113 | 374 | 1,153 |
| BKK06319 | *ydzX* | 1,231 | 1,193 | 0,111 | 263 | 1,179 |
| BKK11740 | *cotZ* | 1,229 | 1,177 | 0,114 | 369 | 1,163 |
| BKK32540 | *bsn* | 1,223 | 1,153 | 0,085 | 138 | 1,139 |
| BKK06120 | *ydjB* | 1,222 | 1,153 | 0,103 | 162 | 1,139 |
| BKK29690 | *acuA* | 1,222 | 1,153 | 0,091 | 97 | 1,139 |
| BKK03070 | *mdr* | 1,221 | 1,216 | 0,071 | 60 | 1,201 |
| BKK32570 | *frlD* | 1,218 | 0,987 | 0,085 | 108 | 0,975 |
| BKK18330 | *ppsB* | 1,217 | 1,186 | 0,104 | 265 | 1,172 |
| BKK03660 | *yclE* | 1,216 | 1,152 | 0,084 | 77 | 1,138 |
| BKK10580 | *yhjO* | 1,213 | 1,223 | 0,167 | 120 | 1,208 |
| BKK35960 | *rbsB* | 1,21 | 1,175 | 0,108 | 324 | 1,161 |
| BKK07390 | *yfmP* | 1,206 | 1,173 | 0,122 | 115 | 1,159 |
| BKK03210 | *ycgN* | 1,205 | 1,144 | 0,117 | 102 | 1,130 |
| BKK12830 | *spoIISA* | 1,205 | 1,182 | 0,112 | 386 | 1,168 |
| BKK23670 | *yqkA* | 1,203 | 1,165 | 0,111 | 373 | 1,151 |
| BKK40780 | *tetL* | 1,202 | 1,161 | 0,100 | 434 | 1,147 |
| BKK37400 | *albD* | 1,199 | 1,180 | 0,104 | 91 | 1,166 |
| BKK02770 | *yccK* | 1,193 | 1,173 | 0,111 | 306 | 1,160 |
| BKK27090 | *aapA* | 1,193 | 1,191 | 0,095 | 644 | 1,177 |
| BKK08450 | *yfiZ* | 1,191 | 0,987 | 0,077 | 110 | 0,975 |
| BKK32390 | *yunF* | 1,19 | 1,162 | 0,104 | 91 | 1,148 |
| BKK33880 | *yvbJ* | 1,19 | 0,987 | 0,073 | 46 | 0,975 |
| BKK23330 | *ypuC* | 1,187 | 1,201 | 0,097 | 299 | 1,187 |
| BKK11550 | *yjbH* | 1,185 | 1,197 | 0,107 | 164 | 1,183 |
| BKK18890 | *yobF* | 1,181 | 1,151 | 0,087 | 139 | 1,138 |
| BKK24110 | *yqzF* | 1,181 | 1,167 | 0,120 | 340 | 1,153 |
| BKK30480 | *ytqA* | 1,179 | 1,170 | 0,119 | 97 | 1,156 |
| BKK31800 | *yueF* | 1,174 | 1,161 | 0,113 | 30 | 1,148 |
| BKK07580 | *citS* | 1,172 | 1,223 | 0,135 | 97 | 1,208 |
| BKK23340 | *ypuB* | 1,172 | 1,201 | 0,096 | 492 | 1,187 |
| BKK19200 | *desR* | 1,17 | 1,185 | 0,081 | 187 | 1,172 |
| BKK13700 | *clpE* | 1,166 | 1,180 | 0,171 | 37 | 1,166 |
| BKK02210 | *ybfH* | 1,165 | 1,143 | 0,101 | 173 | 1,130 |
| BKK39670 | *iolJ* | 1,165 | 1,178 | 0,100 | 128 | 1,165 |

| BKK name [1] | gene | screening delta[2] (%) | erage width (µ | +/- | nb | ADP |
|---|---|---|---|---|---|---|
| BKK19690 | *kamA* | 1,164 | 1,142 | 0,121 | 516 | 1,129 |
| BKK30690 | *ytiB* | 1,163 | 1,169 | 0,101 | 118 | 1,156 |
| BKK34450 | *sacB* | 1,163 | 1,148 | 0,118 | 344 | 1,135 |
| BKK18310 | *ppsD* | 1,159 | 1,151 | 0,074 | 48 | 1,138 |
| BKK24420 | *spoIIIAB* | 1,155 | 1,220 | 0,143 | 44 | 1,206 |
| BKK20999 | *yoyJ* | 1,154 | 1,168 | 0,101 | 696 | 1,155 |
| BKK37790 | *rocG* | 1,141 | 1,132 | 0,104 | 77 | 1,119 |
| BKK22580 | *ypiB* | 1,14 | 1,113 | 0,101 | 432 | 1,100 |
| BKK20840 | *yopM* | 1,138 | 1,167 | 0,109 | 463 | 1,153 |
| BKK35120 | *yvlB* | 1,138 | 1,148 | 0,095 | 221 | 1,135 |
| BKK11600 | *yjbM* | 1,13 | 1,202 | 0,123 | 874 | 1,189 |
| BKK08500 | *yfhE* | 1,128 | 1,172 | 0,107 | 303 | 1,159 |
| BKK25610 | *yqeM* | 1,125 | 1,177 | 0,086 | 116 | 1,164 |
| BKK09860 | *khtT* | 1,124 | 1,222 | 0,149 | 81 | 1,208 |
| BKK21230 | *yomT* | 1,118 | 1,168 | 0,121 | 384 | 1,155 |
| BKK02400 | *ybgF* | 1,117 | 1,143 | 0,092 | 116 | 1,130 |
| BKK06077 | *ydzW* | 1,114 | 1,171 | 0,098 | 123 | 1,158 |
| BKK34530 | *cotR* | 1,114 | 1,148 | 0,101 | 139 | 1,135 |
| BKK06300 | *cotA* | 1,113 | 1,192 | 0,137 | 231 | 1,179 |
| BKK39100 | *yxiO* | 1,113 | 1,220 | 0,122 | 240 | 1,206 |
| BKK19040 | *csaA* | 1,103 | 1,185 | 0,105 | 258 | 1,172 |
| BKK29300 | *rbfK* | 1,1 | 1,152 | 0,102 | 129 | 1,139 |
| BKK13090 | *ykkC* | 1,099 | 1,222 | 0,147 | 54 | 1,208 |
| BKK15870 | *recG* | 1,094 | 1,170 | 0,106 | 87 | 1,157 |
| BKK33070 | *gerAC* | 1,092 | 1,147 | 0,090 | 128 | 1,134 |
| BKK19900 | *yotF* | 1,09 | 1,170 | 0,105 | 101 | 1,158 |
| BKK12270 | *yjlB* | 1,086 | 1,202 | 0,139 | 1310 | 1,189 |
| BKK03320 | *nasB* | 1,083 | 1,172 | 0,122 | 628 | 1,160 |
| BKK14080 | *ykuH* | 1,083 | 1,179 | 0,121 | 496 | 1,166 |
| BKK31530 | *yufM* | 1,082 | 1,164 | 0,123 | 167 | 1,151 |
| BKK20160 | *yosD* | 1,078 | 1,141 | 0,145 | 94 | 1,129 |
| BKK18770 | *cyeA* | 1,073 | 1,150 | 0,093 | 165 | 1,138 |
| BKK23230 | *ypuF* | 1,071 | 1,200 | 0,109 | 87 | 1,187 |
| BKK12620 | *xkdH* | 1,07 | 1,202 | 0,146 | 325 | 1,189 |
| BKK32420 | *pucR* | 1,068 | 1,174 | 0,104 | 189 | 1,161 |
| BKK08130 | *yfjE* | 1,06 | 0,985 | 0,086 | 35 | 0,975 |
| BKK27340 | *yrrO* | 1,056 | 1,163 | 0,119 | 280 | 1,151 |
| BKK34090 | *yvfR* | 1,056 | 1,162 | 0,114 | 355 | 1,149 |
| BKK26800 | *yrpB* | 1,054 | 1,189 | 0,104 | 219 | 1,177 |
| BKK30640 | *ytkC* | 1,053 | 1,150 | 0,100 | 144 | 1,138 |
| BKK39930 | *yxaM* | 1,051 | 1,162 | 0,102 | 406 | 1,149 |
| BKK32200 | *yutJ* | 1,05 | 1,151 | 0,091 | 155 | 1,139 |
| BKK03160 | *ycgJ* | 1,049 | 1,172 | 0,103 | 409 | 1,160 |
| BKK21290 | *yomN* | 1,047 | 1,167 | 0,102 | 284 | 1,155 |
| BKK30230 | *bioK* | 1,047 | 1,163 | 0,122 | 231 | 1,151 |
| BKK23730 | *yqjV* | 1,044 | 1,199 | 0,107 | 514 | 1,187 |
| BKK09250 | *yhcX* | 1,04 | 1,189 | 0,106 | 161 | 1,177 |
| BKK18820 | *yobB* | 1,038 | 1,174 | 0,114 | 363 | 1,162 |
| BKK39340 | *hutP* | 1,031 | 1,177 | 0,110 | 128 | 1,165 |
| BKK26210 | *yqaR* | 1,027 | 1,165 | 0,113 | 220 | 1,153 |
| BKK06037 | *ydzT* | 1,026 | 1,179 | 0,126 | 384 | 1,167 |
| BKK39330 | *abnB* | 1,022 | 1,159 | 0,097 | 385 | 1,147 |
| BKK37780 | *rocA* | 1,02 | 1,159 | 0,119 | 283 | 1,147 |
| BKK14520 | *adeC* | 1,018 | 1,191 | 0,134 | 117 | 1,179 |
| BKK21250 | *yomR* | 1,016 | 1,167 | 0,109 | 564 | 1,155 |
| BKK21050 | *yonN* | 1,015 | 1,167 | 0,111 | 523 | 1,155 |
| BKK02120 | *ybeC* | 1,014 | 1,141 | 0,091 | 47 | 1,130 |
| BKK08330 | *yfiN* | 1,011 | 1,171 | 0,101 | 487 | 1,159 |
| BKK17410 | *cwlC* | 1,01 | 1,219 | 0,172 | 262 | 1,206 |
| BKK27080 | *levR* | 1,009 | 1,189 | 0,102 | 272 | 1,177 |
| BKK26380 | *yqaB* | 1,008 | 1,189 | 0,105 | 524 | 1,177 |
| BKK05710 | *ydhD* | 1,005 | 1,214 | 0,085 | 97 | 1,201 |
| BKK39690 | *iolH* | 1,004 | 1,176 | 0,107 | 125 | 1,165 |
| BKK32720 | *yurZ* | 1,003 | 1,146 | 0,109 | 150 | 1,134 |
| BKK02830 | *ycdI* | 1,001 | 1,171 | 0,097 | 372 | 1,160 |
| BKK18670 | *yoaN* | 1,001 | 1,149 | 0,087 | 138 | 1,138 |
| BKK28340 | *ysnF* | 1,001 | 1,171 | 0,102 | 308 | 1,160 |
| BKK31260 | *mcpB* | 1 | 1,173 | 0,113 | 392 | 1,161 |
| BKK27440 | *glnH* | 0,995 | 1,171 | 0,111 | 306 | 1,160 |
| BKK16330 | *cheY* | 0,994 | 1,173 | 0,112 | 242 | 1,162 |
| BKK29390 | *ytmI* | 0,992 | 1,173 | 0,112 | 387 | 1,161 |
| BKK04800 | *ydcL* | 0,989 | 1,159 | 0,119 | 400 | 1,147 |
| BKK27840 | *safA* | 0,988 | 1,165 | 0,122 | 303 | 1,153 |
| BKK11210 | *argB* | 0,985 | 1,195 | 0,107 | 73 | 1,183 |
| BKK34190 | *yvfH* | 0,985 | 1,146 | 0,112 | 621 | 1,135 |
| BKK39220 | *yxxG* | 0,984 | 1,176 | 0,119 | 107 | 1,165 |
| BKK27880 | *nifS* | 0,982 | 1,162 | 0,113 | 333 | 1,151 |
| BKK14880 | *ctaB* | 0,981 | 1,191 | 0,144 | 304 | 1,179 |
| BKK18980 | *yobJ* | 0,981 | 1,173 | 0,107 | 322 | 1,162 |

| BKK name [1] | gene | screening delta[2] (%) | erage width (µ | +/- | nb | ADP |
|---|---|---|---|---|---|---|
| BKK39210 | *yxiF* | 0,98 | 1,176 | 0,087 | 93 | 1,165 |
| BKK04120 | *yczI* | 0,979 | 1,176 | 0,071 | 157 | 1,165 |
| BKK17530 | *ynaE* | 0,978 | 1,173 | 0,109 | 333 | 1,162 |
| BKK34410 | *yveG* | 0,977 | 1,146 | 0,127 | 133 | 1,135 |
| BKK40060 | *gntK* | 0,976 | 1,179 | 0,123 | 198 | 1,168 |
| BKK05170 | *ydeE* | 0,975 | 0,985 | 0,090 | 172 | 0,975 |
| BKK15470 | *pyrR* | 0,975 | 1,151 | 0,074 | 139 | 1,139 |
| BKK27060 | *levE* | 0,973 | 1,162 | 0,119 | 322 | 1,151 |
| BKK11880 | *metC* | 0,972 | 1,201 | 0,109 | 76 | 1,189 |
| BKK33170 | *yvrB* | 0,971 | 1,145 | 0,072 | 109 | 1,134 |
| BKK05300 | *ydeQ* | 0,969 | 1,171 | 0,111 | 442 | 1,160 |
| BKK24750 | *yqhB* | 0,966 | 1,175 | 0,101 | 199 | 1,164 |
| BKK22150 | *ypvA* | 0,965 | 1,111 | 0,129 | 31 | 1,100 |
| BKK30890 | *ytxO* | 0,965 | 1,167 | 0,102 | 119 | 1,156 |
| BKK37410 | *albE* | 0,963 | 1,161 | 0,113 | 302 | 1,149 |
| BKK09280 | *glpF* | 0,962 | 1,179 | 0,100 | 204 | 1,168 |
| BKK11860 | *yjcH* | 0,962 | 1,200 | 0,130 | 1045 | 1,189 |
| BKK13390 | *ykoT* | 0,962 | 1,178 | 0,160 | 133 | 1,166 |
| BKK07050 | *yesW* | 0,956 | 1,171 | 0,118 | 723 | 1,160 |
| BKK25910 | *yqxH* | 0,951 | 1,169 | 0,093 | 161 | 1,158 |
| BKK30510 | *ytpA* | 0,951 | 1,167 | 0,088 | 123 | 1,156 |
| BKK14100 | *ykuJ* | 0,949 | 1,150 | 0,096 | 157 | 1,139 |
| BKK33050 | *gerAA* | 0,948 | 1,145 | 0,108 | 98 | 1,134 |
| BKK27460 | *glnP* | 0,944 | 1,234 | 0,069 | 98 | 1,222 |
| BKK04140 | *pbpC* | 0,941 | 1,213 | 0,074 | 38 | 1,201 |
| BKK35930 | *rbsD* | 0,94 | 1,160 | 0,109 | 441 | 1,149 |
| BKK27540 | *yrvM* | 0,937 | 1,164 | 0,118 | 194 | 1,153 |
| BKK18510 | *yoxC* | 0,936 | 1,183 | 0,106 | 142 | 1,172 |
| BKK34340 | *epsD* | 0,935 | 1,177 | 0,099 | 149 | 1,166 |
| BKK32840 | *fadN* | 0,933 | 1,145 | 0,109 | 126 | 1,134 |
| BKK29990 | *pbuO* | 0,932 | 1,233 | 0,098 | 43 | 1,222 |
| BKK40080 | *gntZ* | 0,93 | 1,179 | 0,112 | 275 | 1,168 |
| BKK40470 | *yycC* | 0,929 | 1,177 | 0,122 | 74 | 1,166 |
| BKK36540 | *ywnJ* | 0,925 | 1,125 | 0,114 | 142 | 1,114 |
| BKK20410 | *yorE* | 0,921 | 1,173 | 0,108 | 274 | 1,162 |
| BKK30500 | *ytpB* | 0,921 | 1,167 | 0,117 | 135 | 1,156 |
| BKK28060 | *spoIIB* | 0,918 | 1,233 | 0,069 | 47 | 1,222 |
| BKK02580 | *ycbO* | 0,912 | 1,140 | 0,092 | 111 | 1,130 |
| BKK08260 | *yfiG* | 0,912 | 1,219 | 0,123 | 61 | 1,208 |
| BKK08460 | *yfhA* | 0,912 | 1,179 | 0,110 | 271 | 1,168 |
| BKK11929 | *yjzG* | 0,91 | 1,173 | 0,124 | 238 | 1,163 |
| BKK15690 | *rpoZ* | 0,907 | 1,145 | 0,113 | 146 | 1,134 |
| BKK35500 | *degS* | 0,907 | 1,195 | 0,096 | 368 | 1,184 |
| BKK21080 | *yonI* | 0,894 | 1,165 | 0,111 | 263 | 1,155 |
| BKK38170 | *qoxA* | 0,892 | 1,167 | 0,111 | 118 | 1,157 |
| BKK06230 | *iolT* | 0,891 | 1,212 | 0,099 | 51 | 1,201 |
| BKK30350 | *yttB* | 0,891 | 1,233 | 0,104 | 53 | 1,222 |
| BKK29600 | *braB* | 0,888 | 1,150 | 0,097 | 141 | 1,139 |
| BKK32220 | *yutI* | 0,888 | 1,158 | 0,116 | 96 | 1,148 |
| BKK07910 | *yfkF* | 0,887 | 1,173 | 0,111 | 307 | 1,163 |
| BKK18720 | *yoaR* | 0,887 | 1,172 | 0,117 | 408 | 1,162 |
| BKK27430 | *glnQ* | 0,885 | 1,233 | 0,073 | 63 | 1,222 |
| BKK33270 | *yvrO* | 0,885 | 1,177 | 0,092 | 100 | 1,166 |
| BKK02850 | *adcA* | 0,882 | 1,212 | 0,185 | 62 | 1,201 |
| BKK28530 | *etfB* | 0,88 | 1,170 | 0,104 | 54 | 1,160 |
| BKK15810 | *spoVM* | 0,875 | 1,217 | 0,143 | 136 | 1,206 |
| BKK18849 | *yozV* | 0,872 | 1,148 | 0,096 | 136 | 1,138 |
| BKK12500 | *xkdA* | 0,87 | 1,199 | 0,131 | 581 | 1,189 |
| BKK21180 | *yomY* | 0,87 | 1,165 | 0,105 | 121 | 1,155 |
| BKK30710 | *ythA* | 0,869 | 1,171 | 0,110 | 517 | 1,161 |
| BKK32380 | *yunE* | 0,869 | 1,233 | 0,088 | 187 | 1,222 |
| BKK30659 | *ytzI* | 0,864 | 1,166 | 0,099 | 112 | 1,156 |
| BKK29630 | *yttP* | 0,863 | 1,161 | 0,103 | 266 | 1,151 |
| BKK40022 | *yxaC* | 0,862 | 1,159 | 0,118 | 436 | 1,149 |
| BKK29940 | *ytlR* | 0,859 | 1,166 | 0,091 | 104 | 1,156 |
| BKK13620 | *mtnD* | 0,858 | 1,173 | 0,106 | 247 | 1,163 |
| BKK05670 | *ydgJ* | 0,856 | 1,177 | 0,083 | 202 | 1,167 |
| BKK03680 | *yclG* | 0,855 | 1,148 | 0,083 | 150 | 1,138 |
| BKK39540 | *yxeI* | 0,853 | 1,176 | 0,161 | 61 | 1,166 |
| BKK17680 | *thyA* | 0,843 | 1,172 | 0,102 | 391 | 1,162 |
| BKK07735 | *yflB* | 0,842 | 1,169 | 0,116 | 352 | 1,159 |
| BKK06170 | *ydjE* | 0,841 | 1,169 | 0,109 | 242 | 1,160 |
| BKK11490 | *yjbC* | 0,84 | 1,193 | 0,108 | 104 | 1,183 |
| BKK30990 | *thiT* | 0,839 | 1,232 | 0,144 | 449 | 1,222 |
| BKK18110 | *ynfC* | 0,838 | 1,181 | 0,097 | 86 | 1,172 |
| BKK26490 | *yrkJ* | 0,836 | 1,163 | 0,118 | 313 | 1,153 |
| BKK37280 | *narG* | 0,836 | 1,128 | 0,095 | 66 | 1,119 |
| BKK06480 | *purL* | 0,835 | 1,157 | 0,105 | 230 | 1,147 |
| BKK30135 | *yteP* | 0,834 | 1,167 | 0,105 | 100 | 1,158 |

| BKK name [1] | gene | screening delta[2] (%) | erage width (µ | +/- | nb | ADP |
|---|---|---|---|---|---|---|
| BKK32770 | *yusE* | 0,831 | 1,144 | 0,103 | 127 | 1,134 |
| BKK15250 | *ylxW* | 0,829 | 1,178 | 0,106 | 230 | 1,168 |
| BKK29160 | *ytvI* | 0,829 | 0,983 | 0,092 | 59 | 0,975 |
| BKK35050 | *yvnA* | 0,828 | 1,171 | 0,104 | 484 | 1,161 |
| BKK38330 | *lrgB* | 0,825 | 1,176 | 0,115 | 146 | 1,166 |
| BKK17020 | *ymcA* | 0,823 | 1,173 | 0,121 | 89 | 1,164 |
| BKK06240 | *bdhA* | 0,822 | 1,189 | 0,118 | 208 | 1,179 |
| BKK14560 | *defB* | 0,822 | 1,189 | 0,111 | 157 | 1,179 |
| BKK23180 | *spmA* | 0,821 | 1,161 | 0,110 | 214 | 1,151 |
| BKK33310 | *fhuB* | 0,821 | 1,176 | 0,089 | 132 | 1,166 |
| BKK16260 | *ylxF* | 0,819 | 1,172 | 0,105 | 385 | 1,163 |
| BKK19000 | *yobL* | 0,818 | 1,171 | 0,109 | 228 | 1,162 |
| BKK05540 | *ydfS* | 0,817 | 1,157 | 0,108 | 443 | 1,147 |
| BKK16870 | *ymfI* | 0,815 | 1,173 | 0,127 | 152 | 1,164 |
| BKK23960 | *artR* | 0,815 | 1,147 | 0,126 | 64 | 1,138 |
| BKK34440 | *pbpE* | 0,815 | 1,159 | 0,104 | 531 | 1,149 |
| BKK17180 | *pksJ* | 0,813 | 1,173 | 0,138 | 53 | 1,164 |
| BKK03940 | *ycnI* | 0,808 | 1,167 | 0,103 | 145 | 1,158 |
| BKK16940 | *recA* | 0,808 | 1,171 | 0,136 | 314 | 1,162 |
| BKK02440 | *glnJ* | 0,805 | 1,139 | 0,095 | 145 | 1,130 |
| BKK36820 | *atpG* | 0,805 | 1,157 | 0,108 | 161 | 1,147 |
| BKK15490 | *pyrB* | 0,801 | 1,165 | 0,095 | 92 | 1,156 |
| BKK16890 | *ymfK* | 0,801 | 1,173 | 0,116 | 110 | 1,164 |
| BKK38160 | *qoxB* | 0,8 | 1,157 | 0,106 | 300 | 1,147 |
| BKK20940 | *yopC* | 0,797 | 1,164 | 0,106 | 347 | 1,155 |
| BKK23740 | *yqjU* | 0,796 | 1,197 | 0,101 | 476 | 1,187 |
| BKK04260 | *topB* | 0,794 | 1,169 | 0,124 | 161 | 1,160 |
| BKK13070 | *ykkA* | 0,794 | 1,176 | 0,091 | 75 | 1,166 |
| BKK36140 | *ywqO* | 0,792 | 1,123 | 0,129 | 34 | 1,114 |
| BKK38670 | *yxlE* | 0,791 | 1,174 | 0,100 | 71 | 1,165 |
| BKK04970 | *yddH* | 0,788 | 1,167 | 0,103 | 187 | 1,158 |
| BKK15040 | *ylbK* | 0,786 | 1,188 | 0,099 | 57 | 1,179 |
| BKK35980 | *ywsA* | 0,785 | 1,166 | 0,104 | 214 | 1,157 |
| BKK12170 | *yjgD* | 0,782 | 1,198 | 0,171 | 663 | 1,189 |
| BKK05810 | *gmuB* | 0,781 | 1,176 | 0,070 | 262 | 1,167 |
| BKK04550 | *ydbP* | 0,78 | 1,174 | 0,115 | 38 | 1,165 |
| BKK16980 | *spoVS* | 0,78 | 1,171 | 0,114 | 489 | 1,162 |
| BKK39110 | *deaD* | 0,78 | 1,174 | 0,108 | 71 | 1,165 |
| BKK14930 | *ctaG* | 0,778 | 1,188 | 0,084 | 340 | 1,179 |
| BKK38250 | *ywbO* | 0,775 | 1,166 | 0,128 | 73 | 1,157 |
| BKK20570 | *yoqN* | 0,773 | 1,138 | 0,129 | 216 | 1,129 |
| BKK22430 | *panB* | 0,77 | 1,148 | 0,093 | 106 | 1,139 |
| BKK05030 | *yddM* | 0,763 | 1,168 | 0,114 | 341 | 1,160 |
| BKK07210 | *yetK* | 0,763 | 1,177 | 0,114 | 305 | 1,168 |
| BKK21270 | *yomP* | 0,759 | 1,164 | 0,106 | 512 | 1,155 |
| BKK34350 | *epsC* | 0,759 | 1,158 | 0,105 | 493 | 1,149 |
| BKK18000 | *citB* | 0,758 | 1,171 | 0,116 | 333 | 1,162 |
| BKK23200 | *ypuI* | 0,758 | 1,196 | 0,113 | 175 | 1,187 |
| BKK14690 | *yktD* | 0,757 | 1,188 | 0,123 | 159 | 1,179 |
| BKK38220 | *ywcC* | 0,755 | 1,127 | 0,129 | 194 | 1,119 |
| BKK39910 | *yxnB* | 0,754 | 1,173 | 0,116 | 114 | 1,165 |
| BKK02650 | *pcp* | 0,751 | 1,156 | 0,099 | 407 | 1,147 |
| BKK04110 | *lipC* | 0,749 | 1,168 | 0,123 | 363 | 1,160 |
| BKK33150 | *yvqK* | 0,749 | 1,143 | 0,105 | 183 | 1,134 |
| BKK04690 | *rsbT* | 0,747 | 1,174 | 0,125 | 153 | 1,165 |
| BKK17480 | *ynzF* | 0,747 | 1,171 | 0,107 | 549 | 1,162 |
| BKK38760 | *cydA* | 0,746 | 1,173 | 0,112 | 80 | 1,165 |
| BKK28150 | *hemC* | 0,744 | 1,168 | 0,122 | 118 | 1,160 |
| BKK12810 | *xlyA* | 0,738 | 1,171 | 0,124 | 192 | 1,163 |
| BKK37730 | *bacB* | 0,738 | 1,158 | 0,110 | 383 | 1,149 |
| BKK16880 | *ymfJ* | 0,737 | 1,172 | 0,115 | 163 | 1,164 |
| BKK25800 | *yqcK* | 0,735 | 1,162 | 0,119 | 297 | 1,153 |
| BKK20920 | *yopE* | 0,734 | 1,163 | 0,093 | 358 | 1,155 |
| BKK32340 | *lytH* | 0,734 | 1,231 | 0,068 | 103 | 1,222 |
| BKK22670 | *trpD* | 0,727 | 1,108 | 0,100 | 210 | 1,100 |
| BKK34860 | *hisI* | 0,727 | 1,143 | 0,118 | 365 | 1,135 |
| BKK08750 | *ygxA* | 0,722 | 1,171 | 0,117 | 280 | 1,163 |
| BKK07300 | *yfnE* | 0,721 | 0,982 | 0,107 | 90 | 0,975 |
| BKK17310 | *ymaG* | 0,72 | 1,172 | 0,103 | 209 | 1,164 |
| BKK08220 | *yfiC* | 0,719 | 1,217 | 0,149 | 146 | 1,208 |
| BKK20790 | *yopR* | 0,719 | 1,163 | 0,110 | 540 | 1,155 |
| BKK10330 | *yhfQ* | 0,717 | 1,171 | 0,101 | 412 | 1,163 |
| BKK04839 | *ydzL* | 0,716 | 1,173 | 0,101 | 120 | 1,165 |
| BKK21370 | *yomG* | 0,713 | 1,163 | 0,115 | 493 | 1,155 |
| BKK24190 | *yqiI* | 0,713 | 1,215 | 0,141 | 36 | 1,206 |
| BKK39139 | *yxzJ* | 0,713 | 1,173 | 0,106 | 60 | 1,165 |
| BKK10810 | *yisP* | 0,711 | 1,192 | 0,109 | 100 | 1,183 |
| BKK38980 | *scoB* | 0,71 | 1,175 | 0,111 | 95 | 1,166 |
| BKK07310 | *yfnD* | 0,706 | 1,187 | 0,117 | 483 | 1,179 |

### Sup. Table 4: Cell width of mutants of the BKK collection (continued)

| BKK name [1] | gene | screening delta$^2$ (%) | erage width (µı | +/- | nb | ADP |
|---|---|---|---|---|---|---|
| BKK22950 | *ypdA* | 0,704 | 1,108 | 0,122 | 68 | 1,100 |
| BKK16280 | *flgD* | 0,7 | 1,176 | 0,110 | 238 | 1,168 |
| BKK19980 | *yosW* | 0,699 | 1,170 | 0,113 | 242 | 1,162 |
| BKK12010 | *manP* | 0,698 | 1,197 | 0,110 | 207 | 1,189 |
| BKK24400 | *spoIIIAD* | 0,697 | 1,215 | 0,156 | 73 | 1,206 |
| BKK00760 | *pabC* | 0,694 | 1,166 | 0,108 | 91 | 1,158 |
| BKK13760 | *ykvN* | 0,694 | 1,171 | 0,104 | 317 | 1,163 |
| BKK19940 | *yotB* | 0,694 | 1,170 | 0,112 | 401 | 1,162 |
| BKK39310 | *yxiC* | 0,693 | 1,173 | 0,095 | 154 | 1,165 |
| BKK20620 | *yoqI* | 0,69 | 1,170 | 0,123 | 243 | 1,162 |
| BKK30060 | *ytfP* | 0,688 | 1,164 | 0,097 | 179 | 1,156 |
| BKK07410 | *yfmN* | 0,686 | 1,167 | 0,117 | 156 | 1,159 |
| BKK12539 | *ykzK* | 0,685 | 1,157 | 0,127 | 328 | 1,149 |
| BKK12600 | *xkdF* | 0,684 | 1,174 | 0,139 | 219 | 1,166 |
| BKK38620 | *aag* | 0,684 | 1,173 | 0,118 | 86 | 1,165 |
| BKK12030 | *yjdF* | 0,683 | 1,217 | 0,121 | 62 | 1,208 |
| BKK27110 | *yrhO* | 0,681 | 1,185 | 0,110 | 374 | 1,177 |
| BKK27230 | *yrhD* | 0,681 | 1,185 | 0,110 | 374 | 1,177 |
| BKK33460 | *yvgT* | 0,681 | 1,142 | 0,102 | 219 | 1,134 |
| BKK31670 | *yuxO* | 0,677 | 1,156 | 0,143 | 105 | 1,148 |
| BKK38910 | *yxjL* | 0,677 | 1,215 | 0,133 | 72 | 1,206 |
| BKK02260 | *ybfK* | 0,676 | 1,167 | 0,107 | 479 | 1,160 |
| BKK02940 | *yceH* | 0,672 | 1,138 | 0,090 | 112 | 1,130 |
| BKK05000 | *yddK* | 0,668 | 1,167 | 0,120 | 413 | 1,160 |
| BKK18050 | *yneQ* | 0,668 | 1,170 | 0,120 | 382 | 1,162 |
| BKK34000 | *cyeB* | 0,665 | 1,157 | 0,102 | 296 | 1,149 |
| BKK21120 | *yonE* | 0,662 | 1,163 | 0,103 | 431 | 1,155 |
| BKK17750 | *yndD* | 0,657 | 1,171 | 0,099 | 143 | 1,164 |
| BKK24020 | *bmrR* | 0,655 | 1,161 | 0,113 | 349 | 1,153 |
| BKK24230 | *spoIVB* | 0,651 | 1,214 | 0,151 | 171 | 1,206 |
| BKK28140 | *hemD* | 0,649 | 1,167 | 0,115 | 32 | 1,160 |
| BKK30670 | *luxS* | 0,648 | 1,164 | 0,092 | 146 | 1,156 |
| BKK12630 | *xkdI* | 0,643 | 1,197 | 0,115 | 243 | 1,189 |
| BKK37850 | *spsG* | 0,643 | 1,126 | 0,106 | 156 | 1,119 |
| BKK01480 | *truA* | 0,642 | 1,167 | 0,109 | 987 | 1,160 |
| BKK03970 | *ycnL* | 0,64 | 1,172 | 0,075 | 138 | 1,165 |
| BKK02700 | *estA* | 0,637 | 1,209 | 0,069 | 46 | 1,201 |
| BKK40690 | *yybC* | 0,637 | 1,192 | 0,082 | 150 | 1,184 |
| BKK25530 | *spoIIP* | 0,634 | 1,230 | 0,122 | 228 | 1,222 |
| BKK39640 | *yxdL* | 0,634 | 1,230 | 0,123 | 41 | 1,222 |
| BKK18460 | *gltC* | 0,633 | 1,145 | 0,104 | 86 | 1,138 |
| BKK09070 | *yhcG* | 0,632 | 1,184 | 0,119 | 349 | 1,177 |
| BKK39570 | *yxeF* | 0,632 | 1,230 | 0,119 | 41 | 1,222 |
| BKK37970 | *ung* | 0,628 | 1,126 | 0,117 | 234 | 1,119 |
| BKK05400 | *ydfG* | 0,627 | 1,175 | 0,090 | 283 | 1,167 |
| BKK10800 | *yizA* | 0,627 | 1,191 | 0,116 | 78 | 1,183 |
| BKK24250 | *ahrC* | 0,626 | 1,194 | 0,115 | 224 | 1,187 |
| BKK06074 | *ydzW* | 0,624 | 1,175 | 0,131 | 184 | 1,167 |
| BKK15650 | *yloB* | 0,624 | 1,141 | 0,087 | 176 | 1,134 |
| BKK22570 | *ypiF* | 0,623 | 1,107 | 0,115 | 586 | 1,100 |
| BKK08190 | *malR* | 0,622 | 1,166 | 0,120 | 268 | 1,159 |
| BKK29570 | *sspA* | 0,62 | 1,120 | 0,105 | 74 | 1,113 |
| BKK14650 | *yktB* | 0,618 | 1,186 | 0,131 | 229 | 1,179 |
| BKK38200 | *galK* | 0,615 | 1,155 | 0,101 | 480 | 1,147 |
| BKK17699 | *ynzK* | 0,614 | 1,171 | 0,101 | 59 | 1,164 |
| BKK21450 | *yolJ* | 0,614 | 1,158 | 0,108 | 327 | 1,151 |
| BKK40860 | *yyaH* | 0,611 | 1,192 | 0,115 | 101 | 1,184 |
| BKK07450 | *yfmJ* | 0,604 | 1,166 | 0,110 | 189 | 1,159 |
| BKK23390 | *spoVAF* | 0,602 | 1,214 | 0,144 | 33 | 1,206 |
| BKK39490 | *yxeN* | 0,597 | 1,145 | 0,103 | 70 | 1,138 |
| BKK02450 | *glnL* | 0,596 | 1,167 | 0,118 | 302 | 1,160 |
| BKK12050 | *yjdH* | 0,596 | 1,196 | 0,132 | 53 | 1,189 |
| BKK02360 | *nagBB* | 0,593 | 1,137 | 0,109 | 196 | 1,130 |
| BKK08140 | *yfjD* | 0,591 | 1,166 | 0,104 | 594 | 1,159 |
| BKK00420 | *ksgA* | 0,585 | 1,154 | 0,103 | 542 | 1,147 |
| BKK12610 | *xkdG* | 0,585 | 1,196 | 0,089 | 288 | 1,189 |
| BKK16020 | *rimM* | 0,584 | 1,163 | 0,115 | 58 | 1,156 |
| BKK11470 | *oppF* | 0,58 | 1,169 | 0,115 | 270 | 1,163 |
| BKK16299 | *ylzI* | 0,576 | 1,169 | 0,124 | 264 | 1,163 |
| BKK36870 | *atpB* | 0,568 | 1,173 | 0,101 | 96 | 1,166 |
| BKK38000 | *ywdD* | 0,568 | 1,125 | 0,083 | 111 | 1,119 |
| BKK40380 | *yycI* | 0,565 | 1,136 | 0,096 | 121 | 1,130 |
| BKK00890 | *yacL* | 0,563 | 1,154 | 0,107 | 520 | 1,147 |
| BKK30310 | *ytwF* | 0,563 | 1,163 | 0,099 | 107 | 1,156 |
| BKK15880 | *fapR* | 0,562 | 1,125 | 0,126 | 861 | 1,119 |
| BKK38030 | *ywdA* | 0,562 | 1,125 | 0,095 | 98 | 1,119 |
| BKK21400 | *yomD* | 0,558 | 1,161 | 0,101 | 255 | 1,155 |
| BKK30530 | *ytnA* | 0,558 | 1,162 | 0,103 | 83 | 1,156 |
| BKK31340 | *yugM* | 0,553 | 1,144 | 0,094 | 147 | 1,138 |

| BKK name [1] | gene | screening delta$^2$ (%) | erage width (µı | +/- | nb | ADP |
|---|---|---|---|---|---|---|
| BKK14790 | *ylaI* | 0,552 | 1,186 | 0,101 | 798 | 1,179 |
| BKK36930 | *ywlE* | 0,551 | 1,163 | 0,095 | 139 | 1,157 |
| BKK19060 | *yobR* | 0,546 | 1,144 | 0,096 | 95 | 1,138 |
| BKK32550 | *yurJ* | 0,544 | 1,146 | 0,091 | 112 | 1,139 |
| BKK00880 | *disA* | 0,541 | 1,144 | 0,128 | 65 | 1,138 |
| BKK19090 | *yobU* | 0,541 | 1,168 | 0,116 | 346 | 1,162 |
| BKK29950 | *ytlQ* | 0,541 | 1,162 | 0,090 | 85 | 1,156 |
| BKK16780 | *rnjB* | 0,54 | 1,170 | 0,103 | 100 | 1,164 |
| BKK05910 | *ydiB* | 0,539 | 1,174 | 0,094 | 138 | 1,167 |
| BKK17200 | *pksM* | 0,538 | 1,170 | 0,123 | 99 | 1,164 |
| BKK33440 | *cysJ* | 0,538 | 1,140 | 0,123 | 146 | 1,134 |
| BKK22310 | *recU* | 0,533 | 1,106 | 0,126 | 173 | 1,100 |
| BKK06640 | *yerI* | 0,531 | 1,185 | 0,138 | 221 | 1,179 |
| BKK14030 | *ykuC* | 0,53 | 1,213 | 0,164 | 91 | 1,206 |
| BKK35590 | *tuaC* | 0,53 | 1,172 | 0,112 | 139 | 1,166 |
| BKK13250 | *ykoG* | 0,529 | 1,173 | 0,098 | 350 | 1,166 |
| BKK12100 | *yjeA* | 0,526 | 1,169 | 0,111 | 403 | 1,163 |
| BKK24550 | *gcvPB* | 0,522 | 1,193 | 0,116 | 252 | 1,187 |
| BKK04880 | *ydcS* | 0,517 | 1,171 | 0,133 | 55 | 1,165 |
| BKK03750 | *yclJ* | 0,515 | 1,171 | 0,090 | 172 | 1,165 |
| BKK31380 | *yuzA* | 0,512 | 1,154 | 0,097 | 152 | 1,148 |
| BKK36620 | *ywnB* | 0,508 | 1,153 | 0,102 | 370 | 1,147 |
| BKK15069 | *ylzH* | 0,506 | 1,145 | 0,123 | 68 | 1,139 |
| BKK30470 | *ytzC* | 0,505 | 1,162 | 0,102 | 111 | 1,156 |
| BKK39960 | *yxaI* | 0,505 | 1,171 | 0,101 | 113 | 1,165 |
| BKK38050 | *sacP* | 0,503 | 1,124 | 0,113 | 151 | 1,119 |
| BKK11350 | *yjaZ* | 0,502 | 1,168 | 0,108 | 254 | 1,163 |
| BKK13820 | *ykvT* | 0,501 | 1,172 | 0,104 | 198 | 1,166 |
| BKK32760 | *yusD* | 0,498 | 1,167 | 0,114 | 491 | 1,161 |
| BKK36990 | *ywkF* | 0,497 | 1,163 | 0,124 | 81 | 1,157 |
| BKK18908 | *yozY* | 0,495 | 1,144 | 0,094 | 152 | 1,138 |
| BKK26510 | *yrkH* | 0,495 | 1,183 | 0,103 | 265 | 1,177 |
| BKK22560 | *qcrA* | 0,492 | 1,106 | 0,116 | 509 | 1,100 |
| BKK39780 | *iolS* | 0,491 | 1,170 | 0,099 | 67 | 1,165 |
| BKK09640 | *yhdY* | 0,488 | 1,183 | 0,113 | 140 | 1,177 |
| BKK37720 | *bacC* | 0,487 | 1,124 | 0,097 | 60 | 1,119 |
| BKK16370 | *fliR* | 0,486 | 1,167 | 0,104 | 307 | 1,162 |
| BKK37900 | *spsB* | 0,485 | 1,155 | 0,109 | 325 | 1,149 |
| BKK19620 | *yodJ* | 0,484 | 1,212 | 0,137 | 277 | 1,206 |
| BKK16090 | *sucC* | 0,482 | 1,165 | 0,106 | 324 | 1,160 |
| BKK40870 | *ccpB* | 0,48 | 1,153 | 0,106 | 296 | 1,147 |
| BKK18920 | *phrK* | 0,479 | 1,174 | 0,123 | 361 | 1,168 |
| BKK22940 | *prsW* | 0,479 | 1,212 | 0,113 | 177 | 1,206 |
| BKK08700 | *ygaE* | 0,475 | 1,164 | 0,116 | 366 | 1,159 |
| BKK03720 | *gerKB* | 0,474 | 1,170 | 0,119 | 187 | 1,165 |
| BKK31330 | *yugN* | 0,474 | 1,161 | 0,101 | 90 | 1,156 |
| BKK26140 | *yqbE* | 0,472 | 1,157 | 0,107 | 509 | 1,151 |
| BKK23519 | *yqzK* | 0,469 | 1,193 | 0,103 | 228 | 1,187 |
| BKK20050 | *yosQ* | 0,467 | 1,135 | 0,128 | 244 | 1,129 |
| BKK24240 | *recN* | 0,462 | 1,193 | 0,098 | 405 | 1,187 |
| BKK17290 | *ebrB* | 0,459 | 1,212 | 0,123 | 103 | 1,206 |
| BKK12490 | *yjqC* | 0,456 | 1,194 | 0,132 | 479 | 1,189 |
| BKK23710 | *polYB* | 0,455 | 1,192 | 0,105 | 309 | 1,187 |
| BKK04680 | *rsbS* | 0,454 | 1,170 | 0,106 | 190 | 1,165 |
| BKK13720 | *queC* | 0,453 | 1,172 | 0,115 | 305 | 1,166 |
| BKK04920 | *yddC* | 0,451 | 1,153 | 0,104 | 382 | 1,147 |
| BKK18500 | *yoxD* | 0,441 | 1,167 | 0,116 | 316 | 1,162 |
| BKK28680 | *glcD* | 0,441 | 1,144 | 0,102 | 258 | 1,139 |
| BKK14210 | *ykuT* | 0,44 | 1,168 | 0,122 | 316 | 1,163 |
| BKK32350 | *yunB* | 0,439 | 1,144 | 0,103 | 309 | 1,139 |
| BKK39760 | *mmsA* | 0,439 | 1,163 | 0,085 | 196 | 1,158 |
| BKK40030 | *yxaB* | 0,439 | 1,161 | 0,102 | 150 | 1,156 |
| BKK16690 | *pnpA* | 0,437 | 1,167 | 0,103 | 296 | 1,162 |
| BKK13160 | *ohrB* | 0,436 | 1,171 | 0,144 | 251 | 1,166 |
| BKK13200 | *rsbRB* | 0,432 | 1,168 | 0,113 | 293 | 1,163 |
| BKK40110 | *bglA* | 0,432 | 1,173 | 0,112 | 233 | 1,168 |
| BKK02220 | *ybfI* | 0,43 | 1,135 | 0,092 | 170 | 1,130 |
| BKK28510 | *xsa* | 0,428 | 1,165 | 0,117 | 121 | 1,160 |
| BKK32960 | *yusY* | 0,423 | 1,171 | 0,100 | 174 | 1,166 |
| BKK05270 | *ydzF* | 0,422 | 1,165 | 0,117 | 427 | 1,160 |
| BKK23360 | *ppiB* | 0,418 | 1,158 | 0,109 | 475 | 1,153 |
| BKK38320 | *cidA* | 0,418 | 1,154 | 0,114 | 449 | 1,149 |
| BKK23380 | *lysA* | 0,417 | 1,192 | 0,107 | 296 | 1,187 |
| BKK32910 | *yusS* | 0,417 | 1,139 | 0,099 | 102 | 1,134 |
| BKK23850 | *zwf* | 0,414 | 1,158 | 0,119 | 278 | 1,153 |
| BKK39450 | *yxeR* | 0,414 | 1,211 | 0,143 | 161 | 1,206 |
| BKK17170 | *pksI* | 0,412 | 1,168 | 0,122 | 55 | 1,164 |
| BKK19490 | *gerT* | 0,408 | 1,134 | 0,110 | 855 | 1,129 |
| BKK16290 | *flgE* | 0,406 | 1,167 | 0,105 | 264 | 1,163 |

| BKK name [1] | gene | screening delta$^2$ (%) | erage width (µı | +/- | nb | ADP |
|---|---|---|---|---|---|---|
| BKK39410 | *nupC* | 0,406 | 1,211 | 0,121 | 169 | 1,206 |
| BKK39940 | *yxaL* | 0,406 | 1,169 | 0,120 | 149 | 1,165 |
| BKK12320 | *yjmC* | 0,402 | 1,194 | 0,112 | 487 | 1,189 |
| BKK26870 | *yraN* | 0,401 | 1,227 | 0,077 | 226 | 1,222 |
| BKK00540 | *yabK* | 0,399 | 1,168 | 0,116 | 119 | 1,164 |
| BKK23720 | *yqzH* | 0,399 | 1,192 | 0,111 | 149 | 1,187 |
| BKK17640 | *alrB* | 0,398 | 1,211 | 0,140 | 126 | 1,206 |
| BKK20900 | *yopG* | 0,397 | 1,159 | 0,111 | 293 | 1,155 |
| BKK04900 | *yddA* | 0,394 | 1,170 | 0,097 | 174 | 1,165 |
| BKK12069 | *yjzH* | 0,394 | 1,167 | 0,121 | 380 | 1,163 |
| BKK05580 | *ydgC* | 0,389 | 1,172 | 0,078 | 186 | 1,167 |
| BKK27120 | *sigV* | 0,389 | 1,181 | 0,111 | 106 | 1,177 |
| BKK15500 | *pyrC* | 0,388 | 1,162 | 0,099 | 114 | 1,158 |
| BKK33049 | *yvzF* | 0,388 | 1,139 | 0,107 | 137 | 1,134 |
| BKK24100 | *bkdR* | 0,386 | 1,162 | 0,101 | 76 | 1,158 |
| BKK36800 | *atpC* | 0,386 | 1,119 | 0,100 | 114 | 1,114 |
| BKK20870 | *yopJ* | 0,385 | 1,156 | 0,121 | 241 | 1,151 |
| BKK29250 | *nrnA* | 0,383 | 1,117 | 0,091 | 182 | 1,113 |
| BKK06010 | *ydiL* | 0,381 | 1,172 | 0,093 | 276 | 1,167 |
| BKK14970 | *ylbD* | 0,376 | 1,172 | 0,101 | 330 | 1,168 |
| BKK28720 | *abfA* | 0,375 | 1,144 | 0,093 | 167 | 1,139 |
| BKK34020 | *yvbX* | 0,375 | 1,139 | 0,089 | 166 | 1,134 |
| BKK31360 | *yugK* | 0,372 | 1,152 | 0,115 | 238 | 1,148 |
| BKK17630 | *yncC* | 0,371 | 1,211 | 0,179 | 185 | 1,206 |
| BKK35340 | *fliD* | 0,37 | 1,139 | 0,112 | 342 | 1,135 |
| BKK01600 | *ybbA* | 0,369 | 1,181 | 0,096 | 187 | 1,177 |
| BKK27980 | *spoIVFA* | 0,369 | 1,227 | 0,103 | 37 | 1,222 |
| BKK08470 | *yfhB* | 0,363 | 1,163 | 0,106 | 647 | 1,159 |
| BKK04280 | *ydaK* | 0,359 | 1,169 | 0,102 | 90 | 1,165 |
| BKK17140 | *pksF* | 0,358 | 1,168 | 0,102 | 125 | 1,164 |
| BKK22170 | *ypsC* | 0,356 | 1,143 | 0,091 | 185 | 1,139 |
| BKK22140 | *kduD* | 0,355 | 1,104 | 0,136 | 162 | 1,100 |
| BKK34950 | *pelC* | 0,355 | 1,170 | 0,118 | 121 | 1,166 |
| BKK03520 | *srfAD* | 0,349 | 1,206 | 0,110 | 358 | 1,201 |
| BKK33190 | *yvrD* | 0,349 | 1,138 | 0,096 | 120 | 1,134 |
| BKK39770 | *iolR* | 0,349 | 1,151 | 0,099 | 347 | 1,147 |
| BKK21620 | *yokE* | 0,345 | 1,157 | 0,112 | 268 | 1,153 |
| BKK06400 | *yebE* | 0,338 | 1,162 | 0,108 | 155 | 1,158 |
| BKK14730 | *ylaC* | 0,337 | 1,183 | 0,164 | 47 | 1,179 |
| BKK08350 | *estB* | 0,336 | 1,166 | 0,107 | 343 | 1,163 |
| BKK31060 | *gbsA* | 0,336 | 1,162 | 0,101 | 87 | 1,158 |
| BKK14120 | *ykzF* | 0,333 | 1,170 | 0,118 | 709 | 1,166 |
| BKK03740 | *yclI* | 0,327 | 1,163 | 0,114 | 365 | 1,160 |
| BKK33840 | *yvbF* | 0,322 | 1,138 | 0,093 | 94 | 1,134 |
| BKK02290 | *psd* | 0,318 | 1,151 | 0,109 | 486 | 1,147 |
| BKK04960 | *yddG* | 0,314 | 1,205 | 0,144 | 424 | 1,201 |
| BKK15140 | *mraW* | 0,311 | 1,210 | 0,144 | 38 | 1,206 |
| BKK20000 | *yosU* | 0,309 | 1,133 | 0,144 | 350 | 1,129 |
| BKK23140 | *resB* | 0,309 | 1,191 | 0,104 | 848 | 1,187 |
| BKK36840 | *atpH* | 0,309 | 1,118 | 0,117 | 116 | 1,114 |
| BKK10490 | *sipV* | 0,308 | 1,166 | 0,109 | 259 | 1,163 |
| BKK08610 | *yfhO* | 0,306 | 1,172 | 0,112 | 328 | 1,168 |
| BKK18640 | *yoaK* | 0,304 | 1,141 | 0,090 | 121 | 1,138 |
| BKK20020 | *yosS* | 0,303 | 1,133 | 0,130 | 174 | 1,129 |
| BKK36720 | *ywmE* | 0,302 | 1,151 | 0,112 | 230 | 1,147 |
| BKK08010 | *yfjP* | 0,296 | 1,162 | 0,113 | 362 | 1,159 |
| BKK28550 | *fadR* | 0,294 | 1,141 | 0,088 | 128 | 1,138 |
| BKK16270 | *fliK* | 0,293 | 1,171 | 0,120 | 278 | 1,168 |
| BKK27330 | *udk* | 0,291 | 0,978 | 0,120 | 35 | 0,975 |
| BKK04850 | *ydcP* | 0,29 | 1,163 | 0,107 | 520 | 1,160 |
| BKK37260 | *narJ* | 0,29 | 1,122 | 0,102 | 110 | 1,119 |
| BKK00840 | *mcsA* | 0,287 | 1,161 | 0,106 | 73 | 1,158 |
| BKK23790 | *yqjP* | 0,277 | 1,190 | 0,109 | 238 | 1,187 |
| BKK08840 | *ssuA* | 0,273 | 1,171 | 0,111 | 284 | 1,168 |
| BKK17420 | *spoVK* | 0,273 | 1,210 | 0,144 | 106 | 1,206 |
| BKK03340 | *folEB* | 0,272 | 1,133 | 0,087 | 153 | 1,130 |
| BKK40359 | *yyzO* | 0,271 | 1,153 | 0,114 | 356 | 1,149 |
| BKK32450 | *pucL* | 0,269 | 1,143 | 0,082 | 149 | 1,139 |
| BKK23550 | *mleA* | 0,268 | 1,190 | 0,103 | 198 | 1,187 |
| BKK08000 | *yfjQ* | 0,266 | 1,162 | 0,101 | 373 | 1,159 |
| BKK33820 | *opuCB* | 0,266 | 1,137 | 0,091 | 98 | 1,134 |
| BKK39610 | *yxeB* | 0,266 | 1,151 | 0,104 | 590 | 1,147 |
| BKK00570 | *yabM* | 0,265 | 1,141 | 0,131 | 38 | 1,138 |
| BKK27830 | *coxA* | 0,264 | 1,163 | 0,101 | 115 | 1,160 |
| BKK28220 | *clpX* | 0,263 | 1,163 | 0,112 | 147 | 1,160 |
| BKK15400 | *ylmG* | 0,258 | 1,182 | 0,156 | 135 | 1,179 |
| BKK32780 | *yusF* | 0,257 | 1,137 | 0,100 | 178 | 1,134 |
| BKK25940 | *yqcD* | 0,252 | 1,154 | 0,104 | 313 | 1,151 |
| BKK24310 | *folD* | 0,25 | 1,154 | 0,120 | 106 | 1,151 |

| BKK name [1] | gene | screening delta$^2$ (%) | erage width (µı | +/- | nb | ADP |
|---|---|---|---|---|---|---|
| BKK23750 | *yqjT* | 0,249 | 0,977 | 0,082 | 55 | 0,975 |
| BKK33390 | *yvgM* | 0,249 | 1,141 | 0,099 | 123 | 1,138 |
| BKK07600 | *yflP* | 0,247 | 1,162 | 0,103 | 326 | 1,159 |
| BKK19010 | *yobM* | 0,246 | 1,174 | 0,088 | 124 | 1,172 |
| BKK10730 | *yisI* | 0,243 | 1,186 | 0,121 | 118 | 1,183 |
| BKK30020 | *ytzE* | 0,243 | 1,159 | 0,101 | 138 | 1,156 |
| BKK22610 | *tyrA* | 0,242 | 1,103 | 0,142 | 69 | 1,100 |
| BKK33400 | *yvgN* | 0,236 | 1,137 | 0,097 | 121 | 1,134 |
| BKK02370 | *ybgA* | 0,235 | 1,133 | 0,097 | 68 | 1,130 |
| BKK18240 | *yngHA* | 0,235 | 1,174 | 0,104 | 136 | 1,172 |
| BKK05070 | *yddQ* | 0,232 | 1,162 | 0,107 | 412 | 1,160 |
| BKK25340 | *phoH* | 0,231 | 1,166 | 0,098 | 211 | 1,164 |
| BKK09500 | *yhdK* | 0,23 | 1,211 | 0,198 | 77 | 1,208 |
| BKK10030 | *hinT* | 0,23 | 1,180 | 0,108 | 124 | 1,177 |
| BKK31010 | *yuaG* | 0,228 | 1,159 | 0,103 | 99 | 1,156 |
| BKK13410 | *ykoV* | 0,227 | 1,169 | 0,121 | 65 | 1,166 |
| BKK14780 | *ylaH* | 0,226 | 1,182 | 0,104 | 302 | 1,179 |
| BKK17570 | *xynP* | 0,226 | 1,166 | 0,124 | 87 | 1,164 |
| BKK05480 | *ydfN* | 0,225 | 1,170 | 0,100 | 142 | 1,167 |
| BKK26120 | *yqbG* | 0,225 | 1,179 | 0,090 | 222 | 1,177 |
| BKK03790 | *yclM* | 0,224 | 1,168 | 0,101 | 695 | 1,165 |
| BKK13169 | *ykzN* | 0,223 | 1,211 | 0,118 | 207 | 1,208 |
| BKK20350 | *yorK* | 0,222 | 1,142 | 0,109 | 41 | 1,139 |
| BKK09810 | *yhaZ* | 0,22 | 1,165 | 0,110 | 260 | 1,163 |
| BKK04730 | *sigB* | 0,219 | 1,167 | 0,175 | 164 | 1,165 |
| BKK31960 | *dhbF* | 0,219 | 1,169 | 0,112 | 184 | 1,166 |
| BKK37070 | *rpmEA* | 0,217 | 1,160 | 0,098 | 107 | 1,158 |
| BKK02550 | *ycbL* | 0,215 | 1,132 | 0,088 | 130 | 1,130 |
| BKK26560 | *yrkC* | 0,215 | 1,156 | 0,102 | 583 | 1,153 |
| BKK31910 | *yukE* | 0,215 | 1,142 | 0,101 | 140 | 1,139 |
| BKK23650 | *yqkC* | 0,214 | 1,190 | 0,111 | 197 | 1,187 |
| BKK14860 | *pycA* | 0,213 | 1,165 | 0,110 | 228 | 1,163 |
| BKK30910 | *cotSA* | 0,213 | 1,158 | 0,095 | 140 | 1,156 |
| BKK18860 | *yozH* | 0,212 | 1,164 | 0,110 | 370 | 1,162 |
| BKK09550 | *yhdP* | 0,211 | 1,211 | 0,169 | 39 | 1,208 |
| BKK10430 | *yhxD* | 0,211 | 1,186 | 0,119 | 104 | 1,183 |
| BKK05750 | *ydhG* | 0,203 | 1,170 | 0,075 | 265 | 1,167 |
| BKK36920 | *ywlF* | 0,203 | 1,159 | 0,145 | 30 | 1,157 |
| BKK39880 | *yxbC* | 0,202 | 1,150 | 0,112 | 397 | 1,147 |
| BKK40300 | *rapG* | 0,202 | 1,169 | 0,110 | 107 | 1,166 |
| BKK30650 | *dps* | 0,201 | 1,158 | 0,115 | 113 | 1,156 |
| BKK23170 | *spmB* | 0,197 | 1,189 | 0,090 | 186 | 1,187 |
| BKK07700 | *nagP* | 0,196 | 1,161 | 0,113 | 434 | 1,159 |
| BKK13260 | *ykoH* | 0,196 | 1,140 | 0,115 | 113 | 1,138 |
| BKK08560 | *yfhJ* | 0,195 | 1,161 | 0,112 | 591 | 1,159 |
| BKK32090 | *yuiA* | 0,195 | 1,150 | 0,121 | 401 | 1,148 |
| BKK32510 | *pucA* | 0,192 | 1,150 | 0,098 | 85 | 1,148 |
| BKK25030 | *yqgC* | 0,189 | 1,156 | 0,121 | 260 | 1,153 |
| BKK39170 | *yxzG* | 0,184 | 1,167 | 0,111 | 78 | 1,165 |
| BKK11870 | *metI* | 0,179 | 1,191 | 0,133 | 619 | 1,189 |
| BKK28570 | *yshE* | 0,178 | 1,162 | 0,097 | 70 | 1,160 |
| BKK34390 | *pnbA* | 0,177 | 1,137 | 0,108 | 195 | 1,135 |
| BKK32320 | *yutC* | 0,175 | 1,168 | 0,103 | 241 | 1,166 |
| BKK37040 | *ywkB* | 0,174 | 1,151 | 0,115 | 428 | 1,149 |
| BKK33620 | *yvaK* | 0,167 | 1,136 | 0,096 | 148 | 1,134 |
| BKK15080 | *rpmF* | 0,166 | 1,181 | 0,154 | 33 | 1,179 |
| BKK00290 | *yaaQ* | 0,16 | 1,181 | 0,143 | 721 | 1,179 |
| BKK11950 | *yjcQ* | 0,159 | 1,164 | 0,116 | 221 | 1,163 |
| BKK10370 | *bioY* | 0,156 | 1,210 | 0,136 | 251 | 1,208 |
| BKK33730 | *opuBA* | 0,155 | 1,163 | 0,114 | 603 | 1,161 |
| BKK01389 | *ybzG* | 0,148 | 1,165 | 0,103 | 312 | 1,164 |
| BKK03310 | *nasC* | 0,147 | 1,132 | 0,102 | 91 | 1,130 |
| BKK01880 | *ybcI* | 0,145 | 1,140 | 0,108 | 157 | 1,138 |
| BKK38800 | *yxkH* | 0,145 | 1,210 | 0,131 | 126 | 1,208 |
| BKK37390 | *albC* | 0,144 | 1,151 | 0,103 | 476 | 1,149 |
| BKK02050 | *ybdO* | 0,142 | 1,132 | 0,095 | 130 | 1,130 |
| BKK17710 | *tatAC* | 0,142 | 1,165 | 0,110 | 135 | 1,164 |
| BKK04890 | *ydcT* | 0,14 | 1,167 | 0,107 | 124 | 1,165 |
| BKK24180 | *yqiK* | 0,138 | 1,155 | 0,117 | 264 | 1,153 |
| BKK22169 | *ypzG* | 0,134 | 1,141 | 0,090 | 131 | 1,139 |
| BKK16420 | *cheB* | 0,132 | 1,163 | 0,107 | 233 | 1,161 |
| BKK33010 | *cssR* | 0,128 | 1,136 | 0,093 | 170 | 1,134 |
| BKK30520 | *ytoA* | 0,126 | 1,157 | 0,103 | 133 | 1,156 |
| BKK25450 | *prmA* | 0,125 | 1,165 | 0,086 | 229 | 1,164 |
| BKK01970 | *skfG* | 0,123 | 1,141 | 0,092 | 231 | 1,139 |
| BKK13220 | *thiW* | 0,123 | 1,210 | 0,185 | 60 | 1,208 |
| BKK32970 | *yusY* | 0,121 | 1,168 | 0,104 | 140 | 1,166 |
| BKK14740 | *ylaD* | 0,12 | 1,169 | 0,109 | 459 | 1,168 |
| BKK11940 | *yjcP* | 0,117 | 1,190 | 0,102 | 573 | 1,189 |

### Sup. Table 4: Cell width of mutants of the BKK collection (continued)

| BKK name [1] | gene | screening delta$^2$ (%) | erage width (µ | +/- | nb | ADP |
|---|---|---|---|---|---|---|
| BKK22750 | *menH* | 0,117 | 1,102 | 0,111 | 167 | 1,100 |
| BKK01960 | *skfF* | 0,116 | 1,159 | 0,110 | 97 | 1,158 |
| BKK06260 | *ydjN* | 0,115 | 1,180 | 0,126 | 106 | 1,179 |
| BKK02019 | *ybzI* | 0,113 | 1,131 | 0,081 | 88 | 1,130 |
| BKK38350 | *ywbE* | 0,113 | 1,120 | 0,108 | 196 | 1,119 |
| BKK36320 | *ywpG* | 0,112 | 1,158 | 0,106 | 104 | 1,157 |
| BKK27100 | *yrhP* | 0,111 | 1,223 | 0,067 | 74 | 1,222 |
| BKK35330 | *fliS* | 0,109 | 1,136 | 0,116 | 72 | 1,135 |
| BKK14600 | *pdhC* | 0,108 | 0,976 | 0,067 | 62 | 0,975 |
| BKK08940 | *yhbD* | 0,101 | 1,169 | 0,111 | 312 | 1,168 |
| BKK33370 | *yvgK* | 0,1 | 1,162 | 0,112 | 358 | 1,161 |
| BKK32900 | *yusR* | 0,099 | 1,135 | 0,098 | 156 | 1,134 |
| BKK31890 | *yukC* | 0,098 | 1,149 | 0,122 | 33 | 1,148 |
| BKK18660 | *yoaM* | 0,094 | 1,163 | 0,113 | 539 | 1,162 |
| BKK10450 | *yhjB* | 0,092 | 1,169 | 0,100 | 396 | 1,168 |
| BKK21510 | *yolD* | 0,092 | 1,156 | 0,106 | 445 | 1,155 |
| BKK14720 | *ylaB* | 0,088 | 1,169 | 0,114 | 205 | 1,168 |
| BKK30250 | *ytaP* | 0,088 | 1,157 | 0,101 | 182 | 1,156 |
| BKK35890 | *pgsC* | 0,088 | 1,150 | 0,114 | 508 | 1,149 |
| BKK06470 | *purQ* | 0,087 | 1,180 | 0,117 | 328 | 1,179 |
| BKK17090 | *pksB* | 0,087 | 1,208 | 0,142 | 95 | 1,206 |
| BKK08780 | *ygaJ* | 0,084 | 1,160 | 0,108 | 183 | 1,159 |
| BKK05040 | *yddN* | 0,08 | 1,161 | 0,110 | 508 | 1,160 |
| BKK11840 | *yjcF* | 0,08 | 1,190 | 0,102 | 239 | 1,189 |
| BKK21630 | *yokD* | 0,078 | 1,156 | 0,112 | 537 | 1,155 |
| BKK31875 | *yukB* | 0,077 | 1,223 | 0,085 | 142 | 1,222 |
| BKK16230 | *fliH* | 0,074 | 1,163 | 0,109 | 309 | 1,163 |
| BKK09390 | *ygxB* | 0,073 | 1,178 | 0,105 | 122 | 1,177 |
| BKK10870 | *yisU* | 0,071 | 1,184 | 0,122 | 89 | 1,183 |
| BKK08779 | *ygzC* | 0,069 | 0,976 | 0,068 | 78 | 0,975 |
| BKK31220 | *yuxG* | 0,068 | 1,157 | 0,090 | 121 | 1,156 |
| BKK07360 | *yfmS* | 0,067 | 1,209 | 0,162 | 277 | 1,208 |
| BKK20500 | *ligB* | 0,065 | 1,140 | 0,102 | 119 | 1,139 |
| BKK20540 | *yoqR* | 0,065 | 1,140 | 0,096 | 140 | 1,139 |
| BKK35040 | *yvnB* | 0,064 | 1,136 | 0,099 | 258 | 1,135 |
| BKK21940 | *degR* | 0,062 | 1,156 | 0,106 | 328 | 1,155 |
| BKK15090 | *ylbO* | 0,061 | 1,180 | 0,128 | 295 | 1,179 |
| BKK22120 | *kdgR* | 0,061 | 1,154 | 0,105 | 309 | 1,153 |
| BKK24800 | *yqgW* | 0,061 | 1,164 | 0,100 | 306 | 1,164 |
| BKK11260 | *yjzC* | 0,059 | 1,184 | 0,108 | 83 | 1,183 |
| BKK14240 | *rok* | 0,057 | 1,163 | 0,108 | 254 | 1,163 |
| BKK03870 | *ycnE* | 0,054 | 1,166 | 0,109 | 252 | 1,165 |
| BKK34060 | *yvfU* | 0,053 | 1,135 | 0,093 | 115 | 1,134 |
| BKK18850 | *yobD* | 0,045 | 1,139 | 0,083 | 33 | 1,138 |
| BKK38040 | *sacA* | 0,045 | 1,167 | 0,119 | 82 | 1,166 |
| BKK14910 | *ctaE* | 0,044 | 1,180 | 0,124 | 362 | 1,179 |
| BKK30570 | *ytmB* | 0,036 | 1,156 | 0,095 | 100 | 1,156 |
| BKK11270 | *yjzD* | 0,034 | 1,184 | 0,137 | 84 | 1,183 |
| BKK31170 | *yulF* | 0,033 | 1,156 | 0,105 | 133 | 1,156 |
| BKK17920 | *ynzD* | 0,032 | 1,164 | 0,122 | 83 | 1,164 |
| BKK19270 | *yozN* | 0,031 | 1,172 | 0,105 | 99 | 1,172 |
| BKK13110 | *purU* | 0,025 | 1,163 | 0,122 | 377 | 1,163 |
| BKK31190 | *yulD* | 0,025 | 1,151 | 0,116 | 329 | 1,151 |
| BKK39090 | *yxiP* | 0,023 | 1,165 | 0,111 | 167 | 1,165 |
| BKK24170 | *mmgA* | 0,022 | 0,975 | 0,086 | 46 | 0,975 |
| BKK16410 | *ylxH* | 0,021 | 1,156 | 0,093 | 246 | 1,156 |
| BKK28710 | *cstA* | 0,021 | 1,156 | 0,093 | 246 | 1,156 |
| BKK29960 | *ytlP* | 0,021 | 1,156 | 0,106 | 75 | 1,156 |
| BKK12780 | *xepA* | 0,015 | 1,209 | 0,116 | 125 | 1,208 |
| BKK05600 | *ydgE* | 0,014 | 1,167 | 0,078 | 333 | 1,167 |
| BKK11370 | *appF* | 0,012 | 1,168 | 0,112 | 350 | 1,168 |
| BKK10900 | *yisY* | 0,01 | 1,183 | 0,109 | 90 | 1,183 |
| BKK11240 | *carB* | 0,01 | 1,183 | 0,099 | 77 | 1,183 |
| BKK03260 | *ycgS* | 0,008 | 1,130 | 0,082 | 74 | 1,130 |
| BKK16240 | *fliI* | 0,008 | 1,163 | 0,102 | 402 | 1,163 |
| BKK20170 | *yosC* | 0,006 | 1,162 | 0,103 | 302 | 1,162 |
| BKK02200 | *ybfG* | 0,005 | 1,160 | 0,111 | 334 | 1,160 |
| BKK31280 | *yugU* | 0,005 | 1,158 | 0,098 | 182 | 1,158 |
| BKK35510 | *yvyE* | 0,004 | 1,114 | 0,115 | 94 | 1,114 |
| BKK33570 | *yvaE* | 0,003 | 1,222 | 0,096 | 64 | 1,222 |
| BKK05380 | *ydfE* | 0 | 1,167 | 0,119 | 117 | 1,167 |
| BKK38060 | *ywcJ* | -0,001 | 1,158 | 0,104 | 159 | 1,158 |
| BKK07380 | *yfmQ* | -0,002 | 1,159 | 0,114 | 290 | 1,159 |
| BKK04870 | *ydcR* | -0,003 | 1,147 | 0,113 | 421 | 1,147 |
| BKK13310 | *tnrA* | -0,003 | 1,163 | 0,107 | 316 | 1,163 |
| BKK11079 | *yizB* | -0,005 | 1,183 | 0,106 | 129 | 1,183 |
| BKK23940 | *yqjB* | -0,007 | 1,187 | 0,107 | 578 | 1,187 |
| BKK29460 | *moaB* | -0,008 | 1,113 | 0,112 | 56 | 1,113 |
| BKK35070 | *yvmC* | -0,008 | 1,158 | 0,118 | 102 | 1,158 |

| BKK name [1] | gene | screening delta$^2$ (%) | erage width (µ | +/- | nb | ADP |
|---|---|---|---|---|---|---|
| BKK33750 | *sdpA* | -0,01 | 1,166 | 0,114 | 98 | 1,166 |
| BKK03270 | *ycgT* | -0,013 | 1,130 | 0,087 | 70 | 1,130 |
| BKK19810 | *ypqP* | -0,013 | 1,162 | 0,108 | 490 | 1,162 |
| BKK03900 | *gabT* | -0,016 | 1,159 | 0,116 | 439 | 1,160 |
| BKK08280 | *yfiI* | -0,016 | 1,159 | 0,109 | 485 | 1,159 |
| BKK13020 | *ykgA* | -0,017 | 1,166 | 0,113 | 222 | 1,166 |
| BKK21980 | *ypdP* | -0,017 | 1,155 | 0,104 | 410 | 1,155 |
| BKK19080 | *yobT* | -0,019 | 1,138 | 0,096 | 153 | 1,138 |
| BKK11400 | *appC* | -0,021 | 1,162 | 0,113 | 319 | 1,163 |
| BKK09060 | *yhcF* | -0,022 | 1,177 | 0,109 | 213 | 1,177 |
| BKK40940 | *yyaD* | -0,028 | 1,167 | 0,086 | 1044 | 1,167 |
| BKK02680 | *lmrA* | -0,031 | 1,130 | 0,091 | 136 | 1,130 |
| BKK17470 | *ynxB* | -0,038 | 1,161 | 0,104 | 322 | 1,162 |
| BKK06140 | *gutR* | -0,039 | 1,178 | 0,110 | 197 | 1,179 |
| BKK10700 | *gerPC* | -0,039 | 1,183 | 0,099 | 144 | 1,183 |
| BKK13580 | *mtnE* | -0,039 | 1,166 | 0,100 | 307 | 1,166 |
| BKK29930 | *amyX* | -0,044 | 1,156 | 0,096 | 114 | 1,156 |
| BKK22010 | *exoA* | -0,046 | 1,157 | 0,118 | 101 | 1,158 |
| BKK17840 | *fosB* | -0,048 | 1,163 | 0,101 | 79 | 1,164 |
| BKK25420 | *yqeW* | -0,049 | 1,137 | 0,093 | 161 | 1,138 |
| BKK07610 | *citM* | -0,052 | 1,208 | 0,162 | 70 | 1,208 |
| BKK23970 | *artQ* | -0,052 | 1,186 | 0,092 | 375 | 1,187 |
| BKK29679 | *ytzK* | -0,053 | 1,150 | 0,096 | 183 | 1,151 |
| BKK06038 | *ydzT* | -0,055 | 1,157 | 0,097 | 121 | 1,158 |
| BKK10480 | *yhjE* | -0,059 | 1,208 | 0,125 | 156 | 1,208 |
| BKK39240 | *yxxF* | -0,059 | 1,206 | 0,133 | 120 | 1,206 |
| BKK02280 | *ybfM* | -0,061 | 1,129 | 0,088 | 66 | 1,130 |
| BKK08020 | *yfjO* | -0,061 | 1,158 | 0,110 | 459 | 1,159 |
| BKK21150 | *yonB* | -0,067 | 1,154 | 0,116 | 258 | 1,155 |
| BKK11020 | *yitK* | -0,069 | 1,162 | 0,105 | 400 | 1,163 |
| BKK36040 | *ywrJ* | -0,072 | 1,160 | 0,105 | 395 | 1,161 |
| BKK33160 | *yvrA* | -0,076 | 1,133 | 0,087 | 67 | 1,134 |
| BKK28280 | *leuA* | -0,077 | 1,159 | 0,100 | 263 | 1,160 |
| BKK22270 | *yppE* | -0,081 | 1,099 | 0,127 | 308 | 1,100 |
| BKK40420 | *purA* | -0,082 | 1,147 | 0,103 | 301 | 1,147 |
| BKK40530 | *cotF* | -0,084 | 1,164 | 0,100 | 110 | 1,165 |
| BKK23570 | *ansB* | -0,086 | 1,186 | 0,099 | 322 | 1,187 |
| BKK26110 | *yqbH* | -0,088 | 1,176 | 0,101 | 82 | 1,177 |
| BKK19230 | *yocJ* | -0,09 | 1,161 | 0,098 | 389 | 1,162 |
| BKK05190 | *ydeG* | -0,094 | 1,200 | 0,122 | 600 | 1,201 |
| BKK12790 | *xhlA* | -0,096 | 1,165 | 0,126 | 299 | 1,166 |
| BKK18300 | *ppsE* | -0,096 | 1,137 | 0,098 | 126 | 1,138 |
| BKK21280 | *yomO* | -0,096 | 1,154 | 0,102 | 187 | 1,155 |
| BKK18930 | *yobH* | -0,097 | 1,161 | 0,108 | 456 | 1,162 |
| BKK23240 | *ypzK* | -0,097 | 1,152 | 0,111 | 266 | 1,153 |
| BKK12900 | *htrA* | -0,103 | 1,161 | 0,108 | 201 | 1,163 |
| BKK33530 | *yvaA* | -0,104 | 1,133 | 0,107 | 126 | 1,134 |
| BKK00270 | *yaaO* | -0,105 | 1,188 | 0,154 | 255 | 1,189 |
| BKK30760 | *mntB* | -0,107 | 1,155 | 0,104 | 87 | 1,156 |
| BKK17510 | *ynaC* | -0,112 | 1,162 | 0,108 | 108 | 1,164 |
| BKK02040 | *ybdN* | -0,113 | 1,158 | 0,112 | 519 | 1,160 |
| BKK12990 | *ykfC* | -0,113 | 1,207 | 0,136 | 89 | 1,208 |
| BKK25060 | *yqfZ* | -0,113 | 1,152 | 0,115 | 437 | 1,153 |
| BKK00930 | *cysE* | -0,114 | 1,099 | 0,119 | 344 | 1,100 |
| BKK30830 | *menF* | -0,114 | 1,155 | 0,097 | 132 | 1,156 |
| BKK24690 | *comGE* | -0,115 | 1,150 | 0,117 | 411 | 1,151 |
| BKK28430 | *sdhB* | -0,117 | 0,974 | 0,088 | 37 | 0,975 |
| BKK11930 | *yjcO* | -0,119 | 1,188 | 0,115 | 502 | 1,189 |
| BKK33030 | *yuxN* | -0,119 | 1,133 | 0,092 | 114 | 1,134 |
| BKK16460 | *cheD* | -0,12 | 1,160 | 0,105 | 594 | 1,161 |
| BKK25100 | *zur* | -0,121 | 1,162 | 0,099 | 222 | 1,164 |
| BKK03670 | *dtpT* | -0,122 | 1,137 | 0,098 | 191 | 1,138 |
| BKK11200 | *argJ* | -0,122 | 1,167 | 0,101 | 274 | 1,168 |
| BKK06770 | *yeeB* | -0,123 | 1,177 | 0,128 | 196 | 1,179 |
| BKK24870 | *glpG* | -0,124 | 1,162 | 0,106 | 221 | 1,164 |
| BKK04150 | *ycsN* | -0,126 | 1,156 | 0,110 | 186 | 1,158 |
| BKK20360 | *yorJ* | -0,126 | 1,138 | 0,109 | 54 | 1,139 |
| BKK24090 | *ptb* | -0,127 | 1,186 | 0,113 | 204 | 1,187 |
| BKK03910 | *gabD* | -0,129 | 1,137 | 0,094 | 125 | 1,138 |
| BKK10550 | *ntdA* | -0,13 | 1,167 | 0,107 | 302 | 1,168 |
| BKK02900 | *yceD* | -0,131 | 1,128 | 0,092 | 151 | 1,130 |
| BKK04340 | *ydaP* | -0,131 | 1,163 | 0,102 | 180 | 1,165 |
| BKK19250 | *yocL* | -0,131 | 1,170 | 0,087 | 154 | 1,172 |
| BKK35699 | *yvzE* | -0,132 | 1,160 | 0,104 | 471 | 1,161 |
| BKK22650 | *trpF* | -0,133 | 1,099 | 0,099 | 205 | 1,100 |
| BKK23350 | *ypzD* | -0,133 | 1,185 | 0,095 | 555 | 1,187 |
| BKK02740 | *natR* | -0,134 | 1,128 | 0,081 | 33 | 1,130 |
| BKK13400 | *ykoU* | -0,134 | 1,165 | 0,100 | 214 | 1,166 |
| BKK21190 | *yomX* | -0,135 | 1,150 | 0,103 | 417 | 1,151 |

| BKK name [1] | gene | screening delta$^2$ (%) | erage width (µ | +/- | nb | ADP |
|---|---|---|---|---|---|---|
| BKK35680 | *ggaB* | -0,137 | 1,160 | 0,101 | 475 | 1,161 |
| BKK17500 | *ynaB* | -0,138 | 1,160 | 0,102 | 456 | 1,162 |
| BKK03500 | *comS* | -0,14 | 1,156 | 0,117 | 101 | 1,158 |
| BKK18200 | *yngD* | -0,144 | 1,170 | 0,116 | 40 | 1,172 |
| BKK08410 | *yfiV* | -0,147 | 1,157 | 0,110 | 450 | 1,159 |
| BKK25590 | *comEA* | -0,147 | 1,162 | 0,108 | 120 | 1,164 |
| BKK19600 | *yodH* | -0,149 | 0,974 | 0,082 | 95 | 0,975 |
| BKK22290 | *sspM* | -0,157 | 1,099 | 0,129 | 223 | 1,100 |
| BKK11530 | *coiA* | -0,158 | 1,181 | 0,118 | 141 | 1,183 |
| BKK11160 | *yitW* | -0,163 | 1,181 | 0,102 | 106 | 1,183 |
| BKK23800 | *proI* | -0,163 | 1,152 | 0,110 | 315 | 1,153 |
| BKK40800 | *yyaN* | -0,163 | 1,183 | 0,110 | 194 | 1,184 |
| BKK17660 | *yncF* | -0,165 | 1,160 | 0,106 | 345 | 1,162 |
| BKK39660 | *yxdJ* | -0,165 | 1,163 | 0,121 | 126 | 1,165 |
| BKK29750 | *aroA* | -0,167 | 1,138 | 0,091 | 202 | 1,139 |
| BKK16380 | *flhB* | -0,168 | 1,151 | 0,108 | 434 | 1,153 |
| BKK38900 | *yxjM* | -0,169 | 1,206 | 0,165 | 87 | 1,208 |
| BKK10240 | *yhfI* | -0,177 | 1,175 | 0,097 | 124 | 1,177 |
| BKK28290 | *ilvC* | -0,179 | 1,158 | 0,093 | 161 | 1,160 |
| BKK36790 | *ywmA* | -0,179 | 1,155 | 0,107 | 122 | 1,157 |
| BKK12560 | *xpf* | -0,185 | 1,187 | 0,140 | 193 | 1,189 |
| BKK33520 | *csoR* | -0,19 | 1,132 | 0,095 | 118 | 1,134 |
| BKK14820 | *ylaL* | -0,191 | 1,204 | 0,122 | 65 | 1,206 |
| BKK15000 | *ylbG* | -0,192 | 1,177 | 0,122 | 43 | 1,179 |
| BKK28160 | *hemX* | -0,197 | 1,157 | 0,121 | 145 | 1,160 |
| BKK39980 | *qodI* | -0,198 | 1,145 | 0,110 | 477 | 1,147 |
| BKK36340 | *ywpE* | -0,202 | 1,155 | 0,133 | 66 | 1,157 |
| BKK18570 | *yoaE* | -0,206 | 1,136 | 0,092 | 106 | 1,138 |
| BKK19610 | *yodI* | -0,208 | 1,127 | 0,127 | 787 | 1,129 |
| BKK02860 | *adcC* | -0,209 | 1,145 | 0,105 | 129 | 1,147 |
| BKK33640 | *yvaA* | -0,209 | 1,132 | 0,083 | 92 | 1,134 |
| BKK20480 | *yoqX* | -0,21 | 1,159 | 0,112 | 334 | 1,162 |
| BKK32859 | *yuzM* | -0,211 | 1,132 | 0,098 | 229 | 1,134 |
| BKK40070 | *gntP* | -0,211 | 1,145 | 0,101 | 447 | 1,147 |
| BKK05950 | *ydiF* | -0,212 | 1,199 | 0,097 | 201 | 1,201 |
| BKK10880 | *yisV* | -0,212 | 1,181 | 0,120 | 113 | 1,183 |
| BKK17810 | *yndK* | -0,212 | 1,161 | 0,104 | 81 | 1,164 |
| BKK12700 | *xkdQ* | -0,214 | 1,137 | 0,122 | 51 | 1,139 |
| BKK33550 | *yvaC* | -0,219 | 1,155 | 0,093 | 106 | 1,158 |
| BKK19880 | *yotH* | -0,224 | 1,165 | 0,111 | 370 | 1,168 |
| BKK22080 | *ypwA* | -0,224 | 1,137 | 0,089 | 115 | 1,139 |
| BKK10350 | *yhfS* | -0,229 | 1,165 | 0,112 | 381 | 1,168 |
| BKK17670 | *cotU* | -0,229 | 1,161 | 0,102 | 60 | 1,164 |
| BKK40360 | *htrC* | -0,229 | 1,148 | 0,114 | 206 | 1,151 |
| BKK08150 | *yfjC* | -0,233 | 1,165 | 0,104 | 219 | 1,168 |
| BKK27300 | *yrrS* | -0,238 | 1,174 | 0,113 | 99 | 1,177 |
| BKK11080 | *yitQ* | -0,239 | 1,180 | 0,098 | 89 | 1,183 |
| BKK22970 | *mecB* | -0,239 | 1,098 | 0,119 | 380 | 1,100 |
| BKK02140 | *glpT* | -0,24 | 1,157 | 0,106 | 419 | 1,160 |
| BKK07080 | *yesZ* | -0,24 | 1,176 | 0,125 | 170 | 1,179 |
| BKK11750 | *cotY* | -0,24 | 1,160 | 0,105 | 250 | 1,163 |
| BKK12070 | *yjdJ* | -0,244 | 1,137 | 0,115 | 128 | 1,139 |
| BKK33650 | *yvzC* | -0,248 | 1,132 | 0,097 | 106 | 1,134 |
| BKK06850 | *yeeK* | -0,252 | 1,176 | 0,118 | 284 | 1,179 |
| BKK33690 | *yvaQ* | -0,252 | 1,131 | 0,089 | 144 | 1,134 |
| BKK13730 | *queD* | -0,253 | 1,165 | 0,113 | 294 | 1,168 |
| BKK29580 | *thiI* | -0,258 | 1,110 | 0,095 | 79 | 1,113 |
| BKK10460 | *yhjC* | -0,259 | 1,180 | 0,110 | 94 | 1,183 |
| BKK17650 | *yncE* | -0,259 | 1,161 | 0,121 | 77 | 1,164 |
| BKK12950 | *dppD* | -0,262 | 1,205 | 0,221 | 52 | 1,208 |
| BKK25570 | *comEC* | -0,262 | 1,148 | 0,113 | 195 | 1,151 |
| BKK16960 | *ymdA* | -0,267 | 1,160 | 0,118 | 66 | 1,164 |
| BKK03430 | *nucA* | -0,268 | 1,157 | 0,111 | 483 | 1,160 |
| BKK14700 | *nprE* | -0,272 | 1,176 | 0,130 | 125 | 1,179 |
| BKK33080 | *liaR* | -0,273 | 1,131 | 0,113 | 94 | 1,134 |
| BKK33890 | *yvbK* | -0,278 | 1,146 | 0,109 | 488 | 1,149 |
| BKK15060 | *ylbM* | -0,279 | 1,176 | 0,133 | 31 | 1,179 |
| BKK08310 | *yfiL* | -0,28 | 1,205 | 0,173 | 344 | 1,208 |
| BKK29340 | *tcyN* | -0,281 | 1,136 | 0,102 | 79 | 1,139 |
| BKK03580 | *yczE* | -0,282 | 1,135 | 0,083 | 161 | 1,138 |
| BKK21090 | *yonH* | -0,283 | 1,150 | 0,106 | 441 | 1,153 |
| BKK01720 | *ybbK* | -0,285 | 1,156 | 0,099 | 107 | 1,160 |
| BKK09470 | *yhdH* | -0,285 | 1,205 | 0,154 | 154 | 1,208 |
| BKK08180 | *malA* | -0,287 | 1,159 | 0,106 | 297 | 1,163 |
| BKK11500 | *spxA* | -0,288 | 1,154 | 0,108 | 160 | 1,158 |
| BKK26400 | *yrkS* | -0,289 | 1,150 | 0,104 | 160 | 1,153 |
| BKK07290 | *yfnF* | -0,294 | 0,972 | 0,101 | 55 | 0,975 |
| BKK18950 | *yozL* | -0,295 | 1,168 | 0,100 | 174 | 1,172 |
| BKK41040 | *oxaAA* | -0,295 | 1,163 | 0,156 | 108 | 1,166 |

| BKK name [1] | gene | screening delta$^2$ (%) | erage width (µ | +/- | nb | ADP |
|---|---|---|---|---|---|---|
| BKK05110 | *ydeA* | -0,297 | 1,156 | 0,100 | 417 | 1,160 |
| BKK09320 | *yhcY* | -0,299 | 1,135 | 0,095 | 61 | 1,138 |
| BKK02690 | *ansZ* | -0,3 | 1,198 | 0,073 | 70 | 1,201 |
| BKK12930 | *dppB* | -0,301 | 1,163 | 0,110 | 480 | 1,166 |
| BKK11382 | *appA* | -0,302 | 1,165 | 0,106 | 303 | 1,168 |
| BKK30970 | *glgC* | -0,303 | 1,153 | 0,122 | 89 | 1,156 |
| BKK14410 | *sipT* | -0,306 | 1,176 | 0,095 | 649 | 1,179 |
| BKK30980 | *glgB* | -0,307 | 1,152 | 0,086 | 152 | 1,156 |
| BKK38130 | *ywcE* | -0,307 | 1,153 | 0,109 | 126 | 1,157 |
| BKK18360 | *yoxA* | -0,308 | 1,168 | 0,118 | 122 | 1,172 |
| BKK25500 | *hemN* | -0,313 | 1,160 | 0,095 | 276 | 1,164 |
| BKK26430 | *yrkP* | -0,313 | 1,147 | 0,110 | 263 | 1,151 |
| BKK08910 | *yhbA* | -0,315 | 1,173 | 0,096 | 108 | 1,177 |
| BKK20400 | *yorF* | -0,318 | 1,158 | 0,109 | 391 | 1,162 |
| BKK38110 | *nfrA* | -0,321 | 1,144 | 0,114 | 420 | 1,147 |
| BKK18280 | *yngK* | -0,322 | 1,134 | 0,096 | 99 | 1,138 |
| BKK08990 | *yhbI* | -0,324 | 1,173 | 0,107 | 115 | 1,177 |
| BKK01630 | *feuA* | -0,329 | 1,198 | 0,064 | 38 | 1,201 |
| BKK05690 | *ydhB* | -0,329 | 1,163 | 0,080 | 303 | 1,167 |
| BKK10520 | *glcP* | -0,332 | 1,179 | 0,101 | 80 | 1,183 |
| BKK31030 | *yuaE* | -0,334 | 1,152 | 0,097 | 92 | 1,156 |
| BKK02800 | *ycdC* | -0,336 | 1,152 | 0,099 | 127 | 1,156 |
| BKK39650 | *yxdK* | -0,338 | 1,161 | 0,105 | 67 | 1,165 |
| BKK12220 | *yjiC* | -0,34 | 1,159 | 0,105 | 418 | 1,163 |
| BKK39850 | *yxbF* | -0,34 | 1,144 | 0,108 | 397 | 1,147 |
| BKK32800 | *yusH* | -0,341 | 1,154 | 0,096 | 112 | 1,158 |
| BKK25000 | *pbpA* | -0,343 | 0,972 | 0,075 | 76 | 0,975 |
| BKK02540 | *ycbK* | -0,344 | 1,197 | 0,064 | 120 | 1,201 |
| BKK37340 | *ywiB* | -0,344 | 1,162 | 0,113 | 66 | 1,166 |
| BKK16470 | *sigD* | -0,345 | 1,157 | 0,099 | 212 | 1,161 |
| BKK39620 | *yxeA* | -0,35 | 1,161 | 0,106 | 109 | 1,165 |
| BKK08590 | *yfhM* | -0,354 | 1,155 | 0,117 | 288 | 1,159 |
| BKK17270 | *ymaC* | -0,356 | 1,159 | 0,119 | 88 | 1,164 |
| BKK19160 | *yocC* | -0,356 | 1,153 | 0,102 | 125 | 1,158 |
| BKK34050 | *yvfV* | -0,359 | 1,130 | 0,093 | 74 | 1,134 |
| BKK17850 | *lexA* | -0,361 | 1,159 | 0,144 | 93 | 1,164 |
| BKK28600 | *yshB* | -0,362 | 1,156 | 0,094 | 108 | 1,160 |
| BKK19220 | *yocI* | -0,364 | 1,158 | 0,107 | 360 | 1,162 |
| BKK10440 | *yhjA* | -0,366 | 1,179 | 0,097 | 76 | 1,183 |
| BKK28099 | *yszA* | -0,366 | 1,155 | 0,096 | 293 | 1,160 |
| BKK31420 | *yugF* | -0,369 | 1,135 | 0,094 | 104 | 1,139 |
| BKK26620 | *yrdR* | -0,371 | 1,217 | 0,126 | 196 | 1,222 |
| BKK39530 | *yxeJ* | -0,374 | 1,160 | 0,101 | 76 | 1,165 |
| BKK16799 | *ylzJ* | -0,375 | 1,159 | 0,110 | 125 | 1,164 |
| BKK40330 | *rocE* | -0,38 | 1,147 | 0,095 | 426 | 1,151 |
| BKK25490 | *hrcA* | -0,381 | 1,159 | 0,103 | 336 | 1,164 |
| BKK33870 | *yvbI* | -0,386 | 1,157 | 0,112 | 359 | 1,161 |
| BKK31160 | *yubA* | -0,388 | 1,152 | 0,105 | 64 | 1,156 |
| BKK31110 | *yubF* | -0,389 | 1,152 | 0,092 | 86 | 1,156 |
| BKK36670 | *csbD* | -0,389 | 1,110 | 0,104 | 79 | 1,114 |
| BKK29850 | *ytoQ* | -0,39 | 1,157 | 0,109 | 367 | 1,161 |
| BKK30810 | *ytxM* | -0,393 | 1,151 | 0,113 | 71 | 1,156 |
| BKK39190 | *yxiG* | -0,394 | 1,143 | 0,104 | 454 | 1,147 |
| BKK20730 | *yopX* | -0,395 | 1,150 | 0,113 | 409 | 1,155 |
| BKK05880 | *gmuG* | -0,398 | 1,155 | 0,108 | 553 | 1,160 |
| BKK08640 | *yfhS* | -0,398 | 1,154 | 0,117 | 132 | 1,159 |
| BKK22250 | *yppG* | -0,4 | 1,096 | 0,138 | 56 | 1,100 |
| BKK10260 | *yhfK* | -0,405 | 1,172 | 0,093 | 116 | 1,177 |
| BKK14330 | *yknV* | -0,405 | 1,202 | 0,148 | 113 | 1,206 |
| BKK29360 | *tcyL* | -0,405 | 1,109 | 0,092 | 96 | 1,113 |
| BKK29260 | *ytpI* | -0,406 | 1,109 | 0,140 | 221 | 1,113 |
| BKK26540 | *yrkE* | -0,407 | 1,172 | 0,104 | 315 | 1,177 |
| BKK07170 | *yetI* | -0,408 | 1,174 | 0,123 | 200 | 1,179 |
| BKK36240 | *ywqE* | -0,41 | 1,157 | 0,119 | 478 | 1,161 |
| BKK07260 | *ltaSA* | -0,414 | 1,155 | 0,124 | 873 | 1,160 |
| BKK00730 | *cysK* | -0,415 | 1,150 | 0,157 | 186 | 1,155 |
| BKK05060 | *lrpB* | -0,415 | 1,155 | 0,117 | 452 | 1,160 |
| BKK10380 | *hemAT* | -0,416 | 1,203 | 0,179 | 117 | 1,208 |
| BKK12440 | *phrA* | -0,417 | 1,163 | 0,108 | 296 | 1,168 |
| BKK38650 | *yxlG* | -0,417 | 1,161 | 0,146 | 88 | 1,166 |
| BKK05250 | *ydeM* | -0,418 | 1,153 | 0,094 | 211 | 1,158 |
| BKK02180 | *ybfE* | -0,422 | 1,155 | 0,111 | 629 | 1,160 |
| BKK22920 | *ypeB* | -0,423 | 1,149 | 0,110 | 254 | 1,153 |
| BKK06830 | *rapH* | -0,426 | 1,174 | 0,112 | 301 | 1,179 |
| BKK00230 | *bofA* | -0,428 | 1,178 | 0,122 | 117 | 1,183 |
| BKK02190 | *ybfF* | -0,429 | 1,155 | 0,102 | 935 | 1,160 |
| BKK06820 | *yeeG* | -0,429 | 1,174 | 0,111 | 408 | 1,179 |
| BKK34280 | *epsJ* | -0,431 | 1,130 | 0,120 | 451 | 1,135 |
| BKK40600 | *yybL* | -0,431 | 1,161 | 0,112 | 83 | 1,166 |

### Sup. Table 4: Cell width of mutants of the BKK collection (continued)

| BKK name [1] | gene | screening delta[2] (%) | erage width (µ | +/- | nb | ADP |
|---|---|---|---|---|---|---|
| BKK31090 | *ktrA* | -0,432 | 1,151 | 0,092 | 124 | 1,156 |
| BKK23020 | *recQ* | -0,434 | 1,096 | 0,111 | 512 | 1,100 |
| BKK18840 | *xynA* | -0,435 | 1,152 | 0,108 | 127 | 1,157 |
| BKK36850 | *atpF* | -0,436 | 1,161 | 0,107 | 114 | 1,166 |
| BKK22730 | *ndk* | -0,437 | 1,153 | 0,094 | 91 | 1,158 |
| BKK02510 | *garD* | -0,438 | 1,125 | 0,076 | 32 | 1,130 |
| BKK27900 | *pheA* | -0,445 | 1,154 | 0,110 | 78 | 1,160 |
| BKK04010 | *sipU* | -0,449 | 1,160 | 0,119 | 114 | 1,165 |
| BKK29100 | *phoR* | -0,449 | 1,108 | 0,104 | 53 | 1,113 |
| BKK06460 | *purS* | -0,45 | 1,174 | 0,111 | 333 | 1,179 |
| BKK36810 | *atpD* | -0,453 | 1,109 | 0,119 | 259 | 1,114 |
| BKK06430 | *purK* | -0,455 | 1,173 | 0,148 | 134 | 1,179 |
| BKK07060 | *yesX* | -0,456 | 1,152 | 0,108 | 93 | 1,158 |
| BKK25810 | *arsR* | -0,459 | 0,971 | 0,081 | 89 | 0,975 |
| BKK02950 | *yceI* | -0,46 | 1,196 | 0,139 | 96 | 1,201 |
| BKK21870 | *ilvD* | -0,46 | 1,150 | 0,108 | 532 | 1,155 |
| BKK24050 | *bkdAA* | -0,46 | 1,182 | 0,112 | 134 | 1,187 |
| BKK38340 | *ywbF* | -0,461 | 1,114 | 0,120 | 229 | 1,119 |
| BKK29560 | *ytcI* | -0,462 | 1,108 | 0,120 | 58 | 1,113 |
| BKK04380 | *ydaT* | -0,463 | 1,154 | 0,106 | 433 | 1,160 |
| BKK14900 | *ctaD* | -0,465 | 1,174 | 0,101 | 488 | 1,179 |
| BKK06710 | *swrC* | -0,466 | 1,154 | 0,107 | 502 | 1,160 |
| BKK07140 | *yetF* | -0,469 | 1,173 | 0,112 | 281 | 1,179 |
| BKK21490 | *sunI* | -0,469 | 1,148 | 0,110 | 595 | 1,153 |
| BKK39590 | *yxeD* | -0,47 | 1,159 | 0,108 | 161 | 1,165 |
| BKK03359 | *yczL* | -0,471 | 1,125 | 0,081 | 91 | 1,130 |
| BKK19830 | *yotM* | -0,474 | 1,124 | 0,141 | 604 | 1,129 |
| BKK19640 | *yodL* | -0,475 | 1,124 | 0,101 | 1106 | 1,129 |
| BKK27150 | *yrhK* | -0,479 | 1,148 | 0,110 | 477 | 1,153 |
| BKK21690 | *msrA* | -0,48 | 1,148 | 0,107 | 495 | 1,153 |
| BKK24920 | *yqgL* | -0,48 | 1,158 | 0,089 | 136 | 1,164 |
| BKK28690 | *glcF* | -0,484 | 1,146 | 0,109 | 285 | 1,151 |
| BKK24840 | *yqgS* | -0,485 | 1,201 | 0,157 | 48 | 1,206 |
| BKK25090 | *yqfW* | -0,489 | 1,158 | 0,099 | 203 | 1,164 |
| BKK28960 | *ytxC* | -0,491 | 1,108 | 0,096 | 33 | 1,113 |
| BKK12910 | *proG* | -0,492 | 1,161 | 0,094 | 429 | 1,166 |
| BKK07160 | *yetH* | -0,495 | 1,152 | 0,086 | 159 | 1,158 |
| BKK31210 | *yulB* | -0,497 | 1,150 | 0,089 | 111 | 1,156 |
| BKK31240 | *mcpA* | -0,497 | 1,150 | 0,091 | 103 | 1,156 |
| BKK38730 | *cydD* | -0,497 | 1,159 | 0,099 | 176 | 1,165 |
| BKK21740 | *ypmR* | -0,498 | 1,200 | 0,116 | 101 | 1,206 |
| BKK33110 | *liaG* | -0,498 | 1,129 | 0,109 | 107 | 1,134 |
| BKK32849 | *yuzL* | -0,499 | 1,129 | 0,089 | 135 | 1,134 |
| BKK18630 | *exlX* | -0,501 | 1,166 | 0,095 | 178 | 1,172 |
| BKK13600 | *mtnX* | -0,502 | 1,161 | 0,101 | 269 | 1,166 |
| BKK33740 | *yvaV* | -0,502 | 1,155 | 0,104 | 423 | 1,161 |
| BKK39700 | *iolG* | -0,502 | 1,159 | 0,125 | 128 | 1,165 |
| BKK01680 | *ybbF* | -0,509 | 1,154 | 0,108 | 425 | 1,160 |
| BKK04039 | *yczO* | -0,51 | 1,154 | 0,112 | 617 | 1,160 |
| BKK13510 | *ykzE* | -0,511 | 1,157 | 0,110 | 245 | 1,163 |
| BKK19100 | *yobV* | -0,512 | 1,166 | 0,081 | 90 | 1,172 |
| BKK25150 | *yqfQ* | -0,513 | 1,158 | 0,103 | 256 | 1,164 |
| BKK40139 | *yyzN* | -0,515 | 1,142 | 0,101 | 412 | 1,147 |
| BKK23310 | *sipS* | -0,516 | 1,181 | 0,112 | 172 | 1,187 |
| BKK32930 | *yusU* | -0,516 | 1,128 | 0,085 | 121 | 1,134 |
| BKK33029 | *yuzO* | -0,517 | 1,128 | 0,100 | 133 | 1,134 |
| BKK12250 | *yjkB* | -0,52 | 1,202 | 0,179 | 419 | 1,208 |
| BKK20110 | *yosI* | -0,52 | 1,123 | 0,159 | 139 | 1,129 |
| BKK27000 | *yraB* | -0,524 | 1,171 | 0,097 | 137 | 1,177 |
| BKK04230 | *ydaH* | -0,525 | 1,152 | 0,118 | 155 | 1,158 |
| BKK07030 | *yesU* | -0,527 | 1,173 | 0,126 | 429 | 1,179 |
| BKK04160 | *mtlR* | -0,532 | 1,159 | 0,110 | 91 | 1,165 |
| BKK10180 | *yhfC* | -0,533 | 1,156 | 0,105 | 283 | 1,163 |
| BKK04650 | *endB* | -0,537 | 1,159 | 0,096 | 107 | 1,165 |
| BKK28610 | *yshA* | -0,539 | 1,151 | 0,108 | 129 | 1,158 |
| BKK28470 | *lysC* | -0,54 | 0,970 | 0,073 | 32 | 0,975 |
| BKK40640 | *yybH* | -0,546 | 1,178 | 0,108 | 186 | 1,184 |
| BKK07340 | *yfnA* | -0,547 | 1,151 | 0,107 | 68 | 1,158 |
| BKK02430 | *glsA* | -0,548 | 1,153 | 0,096 | 386 | 1,160 |
| BKK10830 | *yisR* | -0,548 | 1,177 | 0,106 | 71 | 1,183 |
| BKK08080 | *acoC* | -0,555 | 1,152 | 0,100 | 334 | 1,159 |
| BKK40100 | *ahpF* | -0,556 | 1,162 | 0,106 | 417 | 1,168 |
| BKK03360 | *yciC* | -0,559 | 1,124 | 0,099 | 94 | 1,130 |
| BKK36400 | *flhO* | -0,559 | 1,155 | 0,090 | 293 | 1,161 |
| BKK27700 | *yrbF* | -0,56 | 1,215 | 0,088 | 197 | 1,222 |
| BKK03240 | *ycgQ* | -0,561 | 1,141 | 0,110 | 538 | 1,147 |
| BKK17520 | *ynaD* | -0,564 | 1,155 | 0,105 | 431 | 1,162 |
| BKK19070 | *yobS* | -0,564 | 1,165 | 0,082 | 212 | 1,172 |
| BKK07270 | *yfnH* | -0,565 | 1,151 | 0,104 | 109 | 1,158 |

| BKK name [1] | gene | screening delta[2] (%) | erage width (µ | +/- | nb | ADP |
|---|---|---|---|---|---|---|
| BKK02300 | *ybfN* | -0,569 | 1,153 | 0,110 | 417 | 1,160 |
| BKK28660 | *sspI* | -0,573 | 1,133 | 0,103 | 135 | 1,139 |
| BKK25690 | *sda* | -0,574 | 1,147 | 0,122 | 141 | 1,153 |
| BKK11760 | *cotX* | -0,575 | 1,182 | 0,129 | 720 | 1,189 |
| BKK21900 | *bsaA* | -0,575 | 1,148 | 0,114 | 230 | 1,155 |
| BKK01890 | *ybcL* | -0,578 | 1,131 | 0,119 | 141 | 1,138 |
| BKK12930 | *dppB* | -0,581 | 1,182 | 0,131 | 1017 | 1,189 |
| BKK30920 | *cotI* | -0,585 | 1,149 | 0,092 | 124 | 1,156 |
| BKK33200 | *yvrE* | -0,585 | 1,128 | 0,099 | 106 | 1,134 |
| BKK06960 | *yesN* | -0,586 | 1,172 | 0,105 | 370 | 1,179 |
| BKK31250 | *tlpA* | -0,588 | 1,149 | 0,104 | 117 | 1,156 |
| BKK32590 | *frlN* | -0,6 | 1,133 | 0,085 | 266 | 1,139 |
| BKK40240 | *yycS* | -0,602 | 1,144 | 0,110 | 363 | 1,151 |
| BKK33510 | *copZ* | -0,609 | 1,127 | 0,106 | 100 | 1,134 |
| BKK27990 | *minD* | -0,61 | 1,131 | 0,093 | 76 | 1,138 |
| BKK23070 | *serA* | -0,611 | 1,151 | 0,112 | 147 | 1,158 |
| BKK03120 | *ycgI* | -0,613 | 1,153 | 0,108 | 731 | 1,160 |
| BKK29980 | *ytjP* | -0,614 | 1,149 | 0,107 | 92 | 1,156 |
| BKK01580 | *ybaR* | -0,616 | 1,194 | 0,074 | 45 | 1,201 |
| BKK04980 | *yddI* | -0,618 | 1,152 | 0,105 | 375 | 1,160 |
| BKK18030 | *tlp* | -0,621 | 1,156 | 0,103 | 91 | 1,164 |
| BKK33239 | *yvrJ* | -0,622 | 1,127 | 0,105 | 103 | 1,134 |
| BKK37240 | *ywiE* | -0,625 | 1,112 | 0,110 | 74 | 1,119 |
| BKK05200 | *ydeH* | -0,628 | 1,160 | 0,112 | 169 | 1,167 |
| BKK06034 | *ydzT* | -0,628 | 1,160 | 0,122 | 635 | 1,167 |
| BKK00990 | *rpmGB* | -0,635 | 0,969 | 0,092 | 74 | 0,975 |
| BKK31270 | *tgl* | -0,635 | 1,149 | 0,096 | 54 | 1,156 |
| BKK39920 | *asnH* | -0,638 | 1,157 | 0,112 | 88 | 1,165 |
| BKK39029 | *yxiT* | -0,642 | 1,157 | 0,124 | 56 | 1,165 |
| BKK17950 | *yneJ* | -0,648 | 1,154 | 0,113 | 407 | 1,162 |
| BKK30035 | *ytzG* | -0,65 | 1,149 | 0,104 | 177 | 1,156 |
| BKK22500 | *ypjD* | -0,651 | 1,093 | 0,110 | 315 | 1,100 |
| BKK07400 | *yfmO* | -0,654 | 1,200 | 0,185 | 537 | 1,208 |
| BKK00520 | *ctc* | -0,655 | 1,156 | 0,119 | 116 | 1,164 |
| BKK00720 | *yacD* | -0,655 | 1,131 | 0,104 | 111 | 1,138 |
| BKK22910 | *ypfA* | -0,656 | 1,199 | 0,165 | 62 | 1,206 |
| BKK03770 | *rapC* | -0,658 | 1,131 | 0,084 | 139 | 1,138 |
| BKK11070 | *yitP* | -0,658 | 1,176 | 0,130 | 112 | 1,183 |
| BKK22190 | *ypsA* | -0,658 | 1,093 | 0,115 | 234 | 1,100 |
| BKK25480 | *grpE* | -0,658 | 1,156 | 0,094 | 228 | 1,164 |
| BKK19590 | *ctpA* | -0,659 | 1,122 | 0,140 | 308 | 1,129 |
| BKK13430 | *ykoX* | -0,661 | 1,200 | 0,116 | 84 | 1,208 |
| BKK30180 | *ytbQ* | -0,663 | 1,148 | 0,087 | 125 | 1,156 |
| BKK07560 | *pel* | -0,667 | 1,151 | 0,108 | 301 | 1,159 |
| BKK09850 | *yhaU* | -0,671 | 1,200 | 0,142 | 65 | 1,208 |
| BKK32830 | *fadA* | -0,673 | 1,127 | 0,096 | 160 | 1,134 |
| BKK09440 | *citA* | -0,68 | 1,155 | 0,099 | 349 | 1,163 |
| BKK07810 | *treA* | -0,683 | 1,151 | 0,119 | 341 | 1,159 |
| BKK29240 | *ytrI* | -0,684 | 1,105 | 0,107 | 208 | 1,113 |
| BKK12640 | *xkdJ* | -0,686 | 1,160 | 0,117 | 308 | 1,168 |
| BKK38520 | *dltC* | -0,686 | 1,142 | 0,117 | 106 | 1,149 |
| BKK21410 | *blyA* | -0,688 | 1,145 | 0,108 | 318 | 1,153 |
| BKK15860 | *sdaAA* | -0,69 | 1,107 | 0,119 | 86 | 1,114 |
| BKK30110 | *yteS* | -0,691 | 1,130 | 0,098 | 172 | 1,138 |
| BKK35560 | *tuaF* | -0,692 | 0,968 | 0,082 | 46 | 0,975 |
| BKK24760 | *rsbRD* | -0,694 | 1,155 | 0,095 | 103 | 1,164 |
| BKK30420 | *ytrE* | -0,694 | 1,214 | 0,099 | 160 | 1,222 |
| BKK26640 | *yrdP* | -0,697 | 1,145 | 0,098 | 461 | 1,153 |
| BKK31820 | *yuzF* | -0,697 | 1,132 | 0,096 | 98 | 1,139 |
| BKK23580 | *ansA* | -0,698 | 1,179 | 0,099 | 267 | 1,187 |
| BKK38100 | *ywcH* | -0,7 | 1,149 | 0,098 | 103 | 1,157 |
| BKK12870 | *mhqA* | -0,703 | 1,158 | 0,122 | 71 | 1,166 |
| BKK36220 | *ywqG* | -0,704 | 1,149 | 0,105 | 146 | 1,157 |
| BKK36650 | *ureB* | -0,704 | 1,149 | 0,105 | 146 | 1,157 |
| BKK24460 | *yqhT* | -0,705 | 1,143 | 0,104 | 402 | 1,151 |
| BKK32920 | *yusT* | -0,705 | 0,968 | 0,072 | 110 | 0,975 |
| BKK40930 | *yyaE* | -0,705 | 1,148 | 0,106 | 143 | 1,156 |
| BKK36740 | *ywmC* | -0,708 | 1,106 | 0,103 | 101 | 1,114 |
| BKK13460 | *rsgI* | -0,71 | 1,160 | 0,110 | 284 | 1,168 |
| BKK15300 | *bpr* | -0,712 | 1,154 | 0,110 | 315 | 1,163 |
| BKK24440 | *yqhV* | -0,713 | 1,179 | 0,103 | 289 | 1,187 |
| BKK17266 | *ymzE* | -0,714 | 1,155 | 0,085 | 76 | 1,164 |
| BKK40880 | *exoAA* | -0,716 | 1,176 | 0,089 | 156 | 1,184 |
| BKK16700 | *ylxY* | -0,721 | 1,155 | 0,112 | 89 | 1,164 |
| BKK37220 | *ywjB* | -0,724 | 1,158 | 0,116 | 106 | 1,166 |
| BKK26840 | *sigZ* | -0,727 | 1,168 | 0,101 | 270 | 1,177 |
| BKK15480 | *pyrP* | -0,732 | 1,148 | 0,110 | 96 | 1,156 |
| BKK06040 | *ydiM* | -0,733 | 1,131 | 0,120 | 118 | 1,139 |
| BKK23630 | *yqkE* | -0,733 | 1,178 | 0,094 | 656 | 1,187 |

| BKK name [1] | gene | screening delta[2] (%) | erage width (µ | +/- | nb | ADP |
|---|---|---|---|---|---|---|
| BKK25830 | *rapE* | -0,734 | 1,155 | 0,093 | 64 | 1,164 |
| BKK17700 | *cotC* | -0,739 | 1,155 | 0,094 | 106 | 1,164 |
| BKK17740 | *ynzB* | -0,739 | 1,153 | 0,107 | 311 | 1,162 |
| BKK21820 | *thyB* | -0,743 | 1,143 | 0,106 | 188 | 1,151 |
| BKK09958 | *yhzE* | -0,744 | 1,168 | 0,092 | 134 | 1,177 |
| BKK35160 | *uvrA* | -0,745 | 1,127 | 0,119 | 693 | 1,135 |
| BKK24810 | *yqgV* | -0,747 | 1,155 | 0,092 | 280 | 1,164 |
| BKK17160 | *pksH* | -0,756 | 1,155 | 0,097 | 112 | 1,164 |
| BKK29740 | *ccpA* | -0,758 | 1,153 | 0,142 | 154 | 1,161 |
| BKK17240 | *ymzB* | -0,759 | 1,153 | 0,108 | 396 | 1,162 |
| BKK32890 | *yusQ* | -0,766 | 1,126 | 0,098 | 134 | 1,134 |
| BKK01810 | *adaA* | -0,767 | 1,151 | 0,096 | 148 | 1,160 |
| BKK19750 | *cgeE* | -0,771 | 1,121 | 0,138 | 159 | 1,129 |
| BKK04500 | *ydbK* | -0,773 | 1,151 | 0,117 | 466 | 1,160 |
| BKK19479 | *yoyC* | -0,781 | 1,120 | 0,107 | 532 | 1,129 |
| BKK21200 | *yomW* | -0,781 | 1,146 | 0,100 | 543 | 1,155 |
| BKK14340 | *yknW* | -0,782 | 1,170 | 0,122 | 261 | 1,179 |
| BKK12150 | *yjgB* | -0,783 | 1,180 | 0,151 | 191 | 1,189 |
| BKK31140 | *cdoA* | -0,788 | 1,147 | 0,111 | 87 | 1,156 |
| BKK11620 | *yjbO* | -0,789 | 1,130 | 0,105 | 59 | 1,139 |
| BKK22960 | *gudB* | -0,791 | 1,148 | 0,108 | 82 | 1,158 |
| BKK24300 | *xseA* | -0,791 | 1,178 | 0,104 | 258 | 1,187 |
| BKK27740 | *ruvA* | -0,791 | 1,150 | 0,119 | 126 | 1,160 |
| BKK05899 | *ydhU* | -0,792 | 1,130 | 0,098 | 190 | 1,139 |
| BKK36450 | *ywoG* | -0,792 | 1,140 | 0,085 | 265 | 1,149 |
| BKK22019 | *ypzF* | -0,794 | 1,092 | 0,147 | 61 | 1,100 |
| BKK31860 | *yueB* | -0,795 | 1,139 | 0,110 | 505 | 1,148 |
| BKK18990 | *yobK* | -0,797 | 1,129 | 0,089 | 101 | 1,138 |
| BKK17930 | *ccdA* | -0,799 | 1,154 | 0,133 | 71 | 1,164 |
| BKK25440 | *rsmE* | -0,801 | 1,154 | 0,099 | 258 | 1,164 |
| BKK33230 | *sigO* | -0,801 | 0,967 | 0,074 | 94 | 0,975 |
| BKK02130 | *glpQ* | -0,805 | 1,150 | 0,111 | 551 | 1,160 |
| BKK19580 | *yodF* | -0,807 | 1,120 | 0,106 | 819 | 1,129 |
| BKK29370 | *tcyK* | -0,807 | 1,129 | 0,094 | 64 | 1,138 |
| BKK12720 | *xkdS* | -0,811 | 1,130 | 0,101 | 226 | 1,139 |
| BKK30070 | *opuD* | -0,812 | 1,152 | 0,107 | 355 | 1,161 |
| BKK31080 | *yuaB* | -0,818 | 1,147 | 0,102 | 80 | 1,156 |
| BKK28100 | *ysxE* | -0,825 | 1,150 | 0,101 | 367 | 1,160 |
| BKK01020 | *rplK* | -0,826 | 1,154 | 0,099 | 453 | 1,164 |
| BKK25890 | *yqxI* | -0,828 | 1,144 | 0,104 | 437 | 1,153 |
| BKK14760 | *ylaF* | -0,829 | 1,169 | 0,131 | 403 | 1,179 |
| BKK18440 | *gltB* | -0,829 | 1,158 | 0,108 | 258 | 1,168 |
| BKK37760 | *rocC* | -0,83 | 1,109 | 0,140 | 54 | 1,119 |
| BKK38070 | *sacT* | -0,834 | 1,109 | 0,169 | 30 | 1,119 |
| BKK05410 | *ydfH* | -0,835 | 1,138 | 0,102 | 406 | 1,147 |
| BKK38870 | *yxkA* | -0,836 | 1,138 | 0,105 | 378 | 1,147 |
| BKK11980 | *yjdA* | -0,837 | 1,153 | 0,111 | 259 | 1,163 |
| BKK02110 | *ybyB* | -0,843 | 1,120 | 0,107 | 72 | 1,130 |
| BKK08890 | *ygaO* | -0,847 | 1,158 | 0,106 | 277 | 1,168 |
| BKK34290 | *epsI* | -0,849 | 1,156 | 0,109 | 145 | 1,166 |
| BKK38080 | *ywcI* | -0,858 | 1,109 | 0,093 | 102 | 1,119 |
| BKK26930 | *yraI* | -0,859 | 1,167 | 0,112 | 295 | 1,177 |
| BKK07820 | *treR* | -0,861 | 1,149 | 0,112 | 507 | 1,159 |
| BKK23100 | *sigX* | -0,863 | 1,091 | 0,118 | 364 | 1,100 |
| BKK24780 | *yqgY* | -0,863 | 1,154 | 0,104 | 231 | 1,164 |
| BKK00260 | *yaaN* | -0,864 | 1,156 | 0,136 | 719 | 1,166 |
| BKK16360 | *fliQ* | -0,865 | 1,152 | 0,118 | 488 | 1,162 |
| BKK03652 | *yclD* | -0,867 | 1,128 | 0,098 | 144 | 1,138 |
| BKK19420 | *yojK* | -0,867 | 1,152 | 0,106 | 434 | 1,162 |
| BKK23910 | *yqjE* | -0,867 | 1,141 | 0,127 | 264 | 1,151 |
| BKK32410 | *pucH* | -0,867 | 1,130 | 0,085 | 208 | 1,139 |
| BKK32950 | *yusW* | -0,869 | 1,128 | 0,090 | 114 | 1,138 |
| BKK10160 | *yhgE* | -0,875 | 1,167 | 0,076 | 97 | 1,177 |
| BKK10250 | *yhfJ* | -0,875 | 1,167 | 0,089 | 101 | 1,177 |
| BKK03789 | *yczN* | -0,88 | 1,155 | 0,104 | 309 | 1,165 |
| BKK20470 | *yoqY* | -0,88 | 1,152 | 0,113 | 368 | 1,162 |
| BKK18870 | *yozI* | -0,883 | 1,152 | 0,114 | 418 | 1,162 |
| BKK31200 | *yulC* | -0,889 | 1,147 | 0,102 | 142 | 1,158 |
| BKK33260 | *yvrN* | -0,889 | 1,147 | 0,119 | 95 | 1,158 |
| BKK01640 | *ybbB* | -0,89 | 1,149 | 0,118 | 376 | 1,160 |
| BKK05460 | *ydfL* | -0,892 | 1,149 | 0,105 | 224 | 1,160 |
| BKK03400 | *yckD* | -0,893 | 0,966 | 0,075 | 74 | 0,975 |
| BKK14060 | *fadH* | -0,893 | 1,156 | 0,119 | 643 | 1,166 |
| BKK04090 | *kipA* | -0,897 | 1,154 | 0,086 | 123 | 1,165 |
| BKK31470 | *kapD* | -0,899 | 1,138 | 0,118 | 264 | 1,148 |
| BKK06812 | *yeeF* | -0,909 | 1,149 | 0,105 | 430 | 1,160 |
| BKK27450 | *glnM* | -0,91 | 1,149 | 0,094 | 177 | 1,160 |
| BKK21260 | *yomQ* | -0,911 | 1,141 | 0,115 | 260 | 1,151 |
| BKK34650 | *yvdC* | -0,911 | 1,125 | 0,094 | 30 | 1,135 |

| BKK name [1] | gene | screening delta[2] (%) | erage width (µ | +/- | nb | ADP |
|---|---|---|---|---|---|---|
| BKK40340 | *rocD* | -0,911 | 1,141 | 0,112 | 191 | 1,151 |
| BKK35698 | *yvzI* | -0,914 | 1,104 | 0,094 | 225 | 1,114 |
| BKK35790 | *yvyI* | -0,914 | 1,104 | 0,120 | 98 | 1,114 |
| BKK39860 | *aldX* | -0,918 | 1,154 | 0,091 | 94 | 1,165 |
| BKK03030 | *ycgB* | -0,919 | 1,120 | 0,100 | 159 | 1,130 |
| BKK16300 | *fliL* | -0,921 | 1,157 | 0,104 | 169 | 1,168 |
| BKK19740 | *yodT* | -0,924 | 1,119 | 0,109 | 569 | 1,129 |
| BKK24010 | *bmr* | -0,924 | 1,147 | 0,097 | | 1,158 |
| BKK10750 | *yisK* | -0,926 | 1,152 | 0,113 | 418 | 1,163 |
| BKK34720 | *yvcP* | -0,928 | 1,139 | 0,095 | 317 | 1,149 |
| BKK27570 | *yrzK* | -0,931 | 1,149 | 0,089 | 55 | 1,160 |
| BKK35200 | *yvkB* | -0,934 | 1,124 | 0,109 | 345 | 1,135 |
| BKK12470 | *yjqA* | -0,935 | 1,178 | 0,098 | 823 | 1,189 |
| BKK15070 | *ylbN* | -0,935 | 1,152 | 0,105 | 238 | 1,163 |
| BKK18040 | *yneP* | -0,935 | 1,157 | 0,116 | 222 | 1,168 |
| BKK07020 | *rhgT* | -0,943 | 1,168 | 0,114 | 406 | 1,179 |
| BKK03390 | *yckC* | -0,946 | 1,147 | 0,111 | 106 | 1,158 |
| BKK03960 | *ycnK* | -0,946 | 1,149 | 0,105 | 386 | 1,160 |
| BKK29380 | *tcyJ* | -0,946 | 1,127 | 0,085 | 113 | 1,138 |
| BKK04200 | *ydaE* | -0,947 | 1,154 | 0,109 | 57 | 1,165 |
| BKK25350 | *yqfD* | -0,948 | 1,153 | 0,103 | 226 | 1,164 |
| BKK23328 | *ypzJ* | -0,949 | 1,140 | 0,105 | 342 | 1,151 |
| BKK40270 | *yycP* | -0,951 | 1,140 | 0,100 | 356 | 1,151 |
| BKK25990 | *yqbS* | -0,953 | 1,140 | 0,115 | 245 | 1,151 |
| BKK06750 | *yefC* | -0,956 | 1,168 | 0,108 | 131 | 1,179 |
| BKK12210 | *yjiB* | -0,956 | 1,178 | 0,101 | 1308 | 1,189 |
| BKK11850 | *yjcG* | -0,958 | 1,178 | 0,110 | 408 | 1,189 |
| BKK19700 | *yodP* | -0,961 | 1,195 | 0,135 | 101 | 1,206 |
| BKK20750 | *yopV* | -0,962 | 1,144 | 0,097 | 498 | 1,155 |
| BKK40190 | *fbp* | -0,966 | 1,157 | 0,108 | 254 | 1,168 |
| BKK20970 | *yonX* | -0,97 | 1,142 | 0,100 | 490 | 1,153 |
| BKK01690 | *ybbH* | -0,973 | 1,165 | 0,106 | 289 | 1,177 |
| BKK32180 | *yutK* | -0,978 | 1,210 | 0,067 | 54 | 1,222 |
| BKK02420 | *glnT* | -0,98 | 1,148 | 0,097 | 479 | 1,160 |
| BKK24630 | *sipW* | -0,981 | 0,965 | 0,084 | 50 | 0,975 |
| BKK25080 | *yqfX* | -0,982 | 1,152 | 0,114 | 249 | 1,164 |
| BKK17790 | *yndH* | -0,991 | 1,156 | 0,108 | 352 | 1,168 |
| BKK19780 | *cgeA* | -0,992 | 1,146 | 0,110 | 134 | 1,158 |
| BKK09660 | *yheN* | -0,993 | 1,165 | 0,104 | 115 | 1,177 |
| BKK17620 | *yncB* | -0,994 | 1,150 | 0,110 | 420 | 1,162 |
| BKK13470 | *sspD* | -0,995 | 1,155 | 0,112 | 115 | 1,166 |
| BKK09710 | *yheI* | -1 | 1,196 | 0,114 | 55 | 1,208 |
| BKK26670 | *gltR* | -1,003 | 1,142 | 0,099 | 508 | 1,153 |
| BKK37650 | *cysL* | -1,004 | 1,108 | 0,109 | 257 | 1,119 |
| BKK13650 | *eag* | -1,005 | 1,151 | 0,108 | 313 | 1,163 |
| BKK24670 | *comGG* | -1,009 | 1,152 | 0,098 | 163 | 1,164 |
| BKK24860 | *yqgQ* | -1,012 | 1,152 | 0,100 | 316 | 1,164 |
| BKK17870 | *yneB* | -1,014 | 1,150 | 0,113 | 307 | 1,162 |
| BKK25120 | *yqfT* | -1,014 | 1,142 | 0,104 | 482 | 1,153 |
| BKK14150 | *ykuN* | -1,018 | 1,167 | 0,123 | 907 | 1,179 |
| BKK36950 | *ywlC* | -1,02 | 0,965 | 0,112 | 41 | 0,975 |
| BKK23110 | *resE* | -1,029 | 1,194 | 0,094 | 126 | 1,206 |
| BKK26830 | *yrpE* | -1,029 | 1,209 | 0,067 | 94 | 1,222 |
| BKK26900 | *yraL* | -1,032 | 1,165 | 0,102 | 431 | 1,177 |
| BKK34220 | *epsO* | -1,033 | 1,123 | 0,111 | 96 | 1,135 |
| BKK40010 | *yxaD* | -1,036 | 1,136 | 0,107 | 433 | 1,147 |
| BKK35880 | *pgsA* | -1,037 | 1,138 | 0,095 | 548 | 1,149 |
| BKK36669 | *ywzF* | -1,038 | 1,136 | 0,100 | 159 | 1,147 |
| BKK27190 | *yrzI* | -1,039 | 1,146 | 0,112 | 86 | 1,158 |
| BKK02530 | *rtpA* | -1,04 | 1,136 | 0,101 | 428 | 1,147 |
| BKK34690 | *yvcS* | -1,041 | 1,137 | 0,105 | 563 | 1,149 |
| BKK33780 | *sdpI* | -1,043 | 1,122 | 0,096 | 107 | 1,134 |
| BKK06780 | *yeeC* | -1,045 | 1,167 | 0,102 | 601 | 1,179 |
| BKK37610 | *ywzC* | -1,045 | 1,107 | 0,104 | 320 | 1,119 |
| BKK38540 | *dltE* | -1,046 | 0,965 | 0,084 | 50 | 0,975 |
| BKK33060 | *gerAB* | -1,047 | 1,122 | 0,095 | 77 | 1,134 |
| BKK33300 | *fhuG* | -1,047 | 1,122 | 0,103 | 99 | 1,134 |
| BKK34710 | *yvcQ* | -1,047 | 1,137 | 0,106 | 440 | 1,149 |
| BKK03230 | *ycgP* | -1,048 | 1,118 | 0,091 | 41 | 1,130 |
| BKK26880 | *yraM* | -1,051 | 1,141 | 0,102 | 504 | 1,153 |
| BKK26420 | *yrkQ* | -1,054 | 1,209 | 0,092 | 161 | 1,222 |
| BKK35800 | *gerBA* | -1,054 | 1,103 | 0,087 | 38 | 1,114 |
| BKK23000 | *ypbE* | -1,057 | 1,089 | 0,113 | 607 | 1,100 |
| BKK18750 | *yooT* | -1,064 | 1,159 | 0,106 | 218 | 1,172 |
| BKK18900 | *yozJ* | -1,064 | 1,126 | 0,093 | 69 | 1,138 |
| BKK31590 | *yufS* | -1,064 | 1,127 | 0,110 | 61 | 1,139 |
| BKK05120 | *cspC* | -1,067 | 1,147 | 0,120 | 380 | 1,160 |
| BKK10910 | *yisZ* | -1,069 | 1,171 | 0,097 | 92 | 1,183 |
| BKK12820 | *spoIISB* | -1,069 | 1,195 | 0,160 | 87 | 1,208 |

### Sup. Table 4: Cell width of mutants of the BKK collection (continued)

| BKK name [1] | gene | screening delta [2] (%) | erage width (µ | +/- | nb | ADP |
|---|---|---|---|---|---|---|
| BKK33120 | *liaH* | -1,071 | 1,122 | 0,084 | 63 | 1,134 |
| BKK05260 | *ydeN* | -1,075 | 1,155 | 0,097 | 89 | 1,167 |
| BKK24160 | *mmgB* | -1,075 | 1,141 | 0,105 | 257 | 1,153 |
| BKK33320 | *fhuD* | -1,075 | 1,209 | 0,105 | 252 | 1,222 |
| BKK01710 | *ybbJ* | -1,077 | 1,164 | 0,102 | 133 | 1,177 |
| BKK27470 | *yrrD* | -1,077 | 1,147 | 0,099 | 194 | 1,160 |
| BKK18350 | *dacC* | -1,078 | 1,193 | 0,135 | 91 | 1,206 |
| BKK21550 | *yokL* | -1,079 | 1,142 | 0,104 | 393 | 1,155 |
| BKK39230 | *wapA* | -1,084 | 1,193 | 0,143 | 195 | 1,206 |
| BKK19870 | *yotI* | -1,086 | 1,117 | 0,135 | 332 | 1,129 |
| BKK09770 | *yheD* | -1,091 | 1,155 | 0,097 | 330 | 1,168 |
| BKK21020 | *yonR* | -1,092 | 1,141 | 0,099 | 548 | 1,153 |
| BKK36420 | *spoIIID* | -1,094 | 1,137 | 0,107 | 357 | 1,149 |
| BKK00250 | *xpaC* | -1,095 | 1,154 | 0,113 | 166 | 1,166 |
| BKK17300 | *ebrA* | -1,095 | 1,127 | 0,080 | 151 | 1,139 |
| BKK28970 | *ytxB* | -1,096 | 1,149 | 0,102 | 394 | 1,161 |
| BKK18960 | *yozM* | -1,103 | 1,149 | 0,111 | 410 | 1,162 |
| BKK40150 | *yydI* | -1,104 | 1,135 | 0,095 | 299 | 1,147 |
| BKK21320 | *yomL* | -1,106 | 1,141 | 0,119 | 521 | 1,153 |
| BKK23860 | *gndA* | -1,107 | 1,174 | 0,123 | 247 | 1,187 |
| BKK15760 | *prpC* | -1,108 | 1,122 | 0,105 | 108 | 1,134 |
| BKK02330 | *ybfQ* | -1,111 | 1,117 | 0,101 | 90 | 1,130 |
| BKK19300 | *yozC* | -1,114 | 1,149 | 0,115 | 374 | 1,162 |
| BKK38960 | *yxjG* | -1,116 | 1,152 | 0,112 | 148 | 1,165 |
| BKK07180 | *yetI* | -1,117 | 1,166 | 0,110 | 343 | 1,179 |
| BKK03090 | *ycgF* | -1,125 | 1,147 | 0,117 | 276 | 1,160 |
| BKK08600 | *csbB* | -1,128 | 1,149 | 0,111 | 343 | 1,163 |
| BKK22260 | *yppF* | -1,128 | 1,127 | 0,099 | 97 | 1,139 |
| BKK05440 | *nap* | -1,129 | 1,145 | 0,093 | 126 | 1,158 |
| BKK31680 | *comA* | -1,13 | 1,127 | 0,091 | 114 | 1,139 |
| BKK34200 | *sigL* | -1,131 | 1,122 | 0,150 | 50 | 1,135 |
| BKK07890 | *yfkI* | -1,132 | 1,146 | 0,112 | 300 | 1,159 |
| BKK10680 | *gerPE* | -1,132 | 1,155 | 0,094 | 302 | 1,168 |
| BKK34850 | *yvcA* | -1,135 | 1,136 | 0,117 | 519 | 1,149 |
| BKK06280 | *ydjP* | -1,136 | 1,134 | 0,107 | 302 | 1,147 |
| BKK10420 | *comK* | -1,136 | 1,170 | 0,094 | 96 | 1,183 |
| BKK11230 | *carA* | -1,136 | 1,170 | 0,126 | 87 | 1,183 |
| BKK32530 | *pucF* | -1,138 | 1,135 | 0,141 | 51 | 1,148 |
| BKK27680 | *yrbG* | -1,143 | 1,146 | 0,099 | 381 | 1,160 |
| BKK13170 | *guaD* | -1,145 | 1,153 | 0,132 | 208 | 1,166 |
| BKK06390 | *yebD* | -1,146 | 1,146 | 0,108 | 324 | 1,160 |
| BKK30200 | *bioB* | -1,148 | 1,143 | 0,097 | 92 | 1,156 |
| BKK30300 | *melA* | -1,149 | 1,143 | 0,096 | 118 | 1,156 |
| BKK35060 | *cypX* | -1,149 | 1,122 | 0,122 | 509 | 1,135 |
| BKK17820 | *yndL* | -1,152 | 1,150 | 0,117 | 119 | 1,164 |
| BKK04050 | *ycsF* | -1,156 | 1,134 | 0,098 | 508 | 1,147 |
| BKK13870 | *ykvZ* | -1,157 | 1,149 | 0,104 | 324 | 1,163 |
| BKK39460 | *yxeQ* | -1,157 | 1,151 | 0,090 | 123 | 1,165 |
| BKK38820 | *yxkF* | -1,158 | 1,151 | 0,100 | 179 | 1,165 |
| BKK31450 | *kinB* | -1,16 | 1,208 | 0,062 | 59 | 1,222 |
| BKK27010 | *adhA* | -1,166 | 1,163 | 0,099 | 387 | 1,177 |
| BKK20980 | *yonV* | -1,167 | 1,140 | 0,101 | 590 | 1,153 |
| BKK21660 | *yokA* | -1,168 | 1,141 | 0,094 | 236 | 1,155 |
| BKK18190 | *yngC* | -1,169 | 1,192 | 0,143 | 65 | 1,206 |
| BKK05080 | *yddR* | -1,171 | 1,154 | 0,114 | 251 | 1,167 |
| BKK25730 | *yqeC* | -1,173 | 1,150 | 0,116 | 153 | 1,164 |
| BKK00120 | *pdxT* | -1,176 | 1,144 | 0,111 | 91 | 1,158 |
| BKK22850 | *seaA* | -1,178 | 1,087 | 0,116 | 159 | 1,100 |
| BKK00430 | *yabG* | -1,179 | 1,150 | 0,094 | 141 | 1,164 |
| BKK13299 | *ykzO* | -1,179 | 1,087 | 0,112 | 133 | 1,100 |
| BKK32400 | *yunG* | -1,179 | 1,134 | 0,128 | 144 | 1,148 |
| BKK09530 | *yhdN* | -1,18 | 1,149 | 0,118 | 291 | 1,163 |
| BKK29970 | *ytkP* | -1,181 | 1,142 | 0,114 | 107 | 1,156 |
| BKK17350 | *ymzC* | -1,186 | 1,148 | 0,109 | 446 | 1,162 |
| BKK10170 | *fabHB* | -1,187 | 1,163 | 0,097 | 106 | 1,177 |
| BKK32030 | *bioYB* | -1,187 | 1,208 | 0,068 | 97 | 1,222 |
| BKK37310 | *fnr* | -1,187 | 1,134 | 0,101 | 308 | 1,147 |
| BKK37580 | *ywgB* | -1,188 | 1,105 | 0,120 | 802 | 1,119 |
| BKK12649 | *ykzM* | -1,189 | 1,154 | 0,111 | 241 | 1,168 |
| BKK02910 | *yceE* | -1,191 | 1,117 | 0,099 | 166 | 1,130 |
| BKK13670 | *mhqR* | -1,191 | 1,152 | 0,116 | 231 | 1,166 |
| BKK15660 | *yloC* | -1,191 | 1,121 | 0,102 | 102 | 1,134 |
| BKK08520 | *recX* | -1,193 | 1,145 | 0,103 | 280 | 1,159 |
| BKK33090 | *liaS* | -1,193 | 1,207 | 0,126 | 67 | 1,222 |
| BKK12180 | *yjhA* | -1,199 | 1,194 | 0,161 | 146 | 1,208 |
| BKK16160 | *clpY* | -1,2 | 1,149 | 0,105 | 311 | 1,163 |
| BKK04820 | *immR* | -1,201 | 1,144 | 0,095 | 123 | 1,158 |
| BKK30090 | *yteU* | -1,218 | 1,124 | 0,099 | 116 | 1,138 |
| BKK38500 | *dltA* | -1,218 | 1,135 | 0,124 | 115 | 1,149 |
| BKK08380 | *yfiS* | -1,219 | 1,145 | 0,110 | 285 | 1,159 |
| BKK35920 | *rbsK* | -1,219 | 1,101 | 0,093 | 72 | 1,114 |
| BKK08320 | *yfiM* | -1,221 | 1,145 | 0,107 | 270 | 1,159 |
| BKK30380 | *bceA* | -1,222 | 1,207 | 0,080 | 97 | 1,222 |
| BKK24650 | *yqzG* | -1,223 | 1,173 | 0,107 | 165 | 1,187 |
| BKK20850 | *yopL* | -1,224 | 1,137 | 0,101 | 257 | 1,151 |
| BKK36570 | *ywnG* | -1,224 | 1,101 | 0,107 | 208 | 1,114 |
| BKK20130 | *yosG* | -1,225 | 1,148 | 0,109 | 236 | 1,162 |
| BKK26780 | *yrdA* | -1,225 | 1,139 | 0,111 | 450 | 1,153 |
| BKK04630 | *ydcC* | -1,228 | 1,187 | 0,115 | 558 | 1,201 |
| BKK20740 | *yopW* | -1,228 | 1,141 | 0,097 | 412 | 1,155 |
| BKK38840 | *yxkD* | -1,233 | 1,193 | 0,123 | 83 | 1,208 |
| BKK38580 | *licC* | -1,234 | 1,152 | 0,127 | 75 | 1,166 |
| BKK14170 | *ykuP* | -1,241 | 1,148 | 0,098 | 319 | 1,163 |
| BKK07230 | *yetM* | -1,242 | 1,164 | 0,111 | 529 | 1,179 |
| BKK09090 | *yhcI* | -1,244 | 1,154 | 0,112 | 287 | 1,168 |
| BKK38880 | *yxjO* | -1,244 | 1,150 | 0,102 | 79 | 1,165 |
| BKK36700 | *moaA* | -1,245 | 1,100 | 0,111 | 86 | 1,114 |
| BKK06940 | *yesL* | -1,246 | 1,153 | 0,099 | 342 | 1,168 |
| BKK07500 | *yfmE* | -1,247 | 1,144 | 0,102 | 257 | 1,159 |
| BKK38710 | *yxlA* | -1,25 | 1,150 | 0,114 | 148 | 1,165 |
| BKK03330 | *nasA* | -1,255 | 1,186 | 0,074 | 69 | 1,201 |
| BKK02870 | *adcB* | -1,26 | 1,124 | 0,109 | 108 | 1,138 |
| BKK39420 | *deoC* | -1,262 | 1,150 | 0,108 | 126 | 1,165 |
| BKK35910 | *rbsR* | -1,264 | 1,152 | 0,108 | 135 | 1,166 |
| BKK20890 | *yopH* | -1,265 | 1,139 | 0,091 | 514 | 1,153 |
| BKK26680 | *yrdK* | -1,269 | 1,136 | 0,100 | 279 | 1,151 |
| BKK17450 | *glnR* | -1,272 | 1,147 | 0,101 | 632 | 1,162 |
| BKK11440 | *oppB* | -1,273 | 1,193 | 0,162 | 80 | 1,208 |
| BKK20070 | *nrdIB* | -1,276 | 1,115 | 0,120 | 112 | 1,129 |
| BKK00400 | *yabE* | -1,28 | 1,125 | 0,106 | 65 | 1,139 |
| BKK37130 | *spo0F* | -1,28 | 1,135 | 0,095 | 525 | 1,149 |
| BKK17130 | *acpK* | -1,293 | 1,148 | 0,120 | 90 | 1,164 |
| BKK34610 | *mdxE* | -1,296 | 1,146 | 0,117 | 352 | 1,161 |
| BKK25390 | *yqeZ* | -1,297 | 1,148 | 0,095 | 182 | 1,164 |
| BKK35280 | *yvjA* | -1,298 | 1,120 | 0,111 | 324 | 1,135 |
| BKK34040 | *yvfW* | -1,299 | 1,120 | 0,090 | 138 | 1,134 |
| BKK11000 | *yitI* | -1,302 | 1,147 | 0,117 | 303 | 1,163 |
| BKK21880 | *ypgR* | -1,302 | 1,140 | 0,097 | 477 | 1,155 |
| BKK00370 | *abrB* | -1,303 | 1,164 | 0,146 | 91 | 1,179 |
| BKK06450 | *purC* | -1,303 | 1,163 | 0,119 | 259 | 1,179 |
| BKK20280 | *yorR* | -1,303 | 1,153 | 0,106 | 244 | 1,168 |
| BKK38230 | *ywcB* | -1,303 | 1,104 | 0,117 | 69 | 1,119 |
| BKK17720 | *yndA* | -1,304 | 1,147 | 0,106 | 409 | 1,162 |
| BKK12730 | *xkdT* | -1,307 | 1,173 | 0,126 | 110 | 1,189 |
| BKK26390 | *spoIIIC* | -1,313 | 1,161 | 0,109 | 187 | 1,177 |
| BKK31400 | *yugH* | -1,315 | 1,133 | 0,116 | 120 | 1,148 |
| BKK12550 | *xtrA* | -1,317 | 1,153 | 0,120 | 208 | 1,168 |
| BKK34660 | *yvdB* | -1,318 | 1,134 | 0,106 | 357 | 1,149 |
| BKK06760 | *yeeA* | -1,321 | 1,163 | 0,105 | 420 | 1,179 |
| BKK07510 | *yfmD* | -1,321 | 1,192 | 0,147 | 53 | 1,208 |
| BKK20860 | *yopK* | -1,323 | 1,138 | 0,112 | 512 | 1,153 |
| BKK03860 | *ycnD* | -1,324 | 1,132 | 0,103 | 386 | 1,147 |
| BKK05660 | *ydgI* | -1,325 | 1,142 | 0,106 | 80 | 1,158 |
| BKK23430 | *spoVAB* | -1,33 | 1,190 | 0,130 | 100 | 1,206 |
| BKK07990 | *yfjR* | -1,331 | 1,143 | 0,109 | 324 | 1,159 |
| BKK04210 | *ydaF* | -1,333 | 1,149 | 0,096 | 136 | 1,165 |
| BKK37810 | *spsL* | -1,339 | 1,134 | 0,110 | 405 | 1,149 |
| BKK06920 | *yesJ* | -1,342 | 1,163 | 0,127 | 285 | 1,179 |
| BKK30410 | *ytrF* | -1,343 | 1,140 | 0,101 | 163 | 1,156 |
| BKK39510 | *yxeL* | -1,347 | 1,149 | 0,106 | 82 | 1,165 |
| BKK04780 | *ydcI* | -1,352 | 1,149 | 0,133 | 181 | 1,165 |
| BKK01980 | *skfH* | -1,353 | 1,144 | 0,101 | 436 | 1,160 |
| BKK08360 | *yfiQ* | -1,354 | 1,143 | 0,120 | 230 | 1,159 |
| BKK02620 | *phoD* | -1,357 | 1,132 | 0,109 | 508 | 1,147 |
| BKK37380 | *albB* | -1,359 | 1,104 | 0,094 | 61 | 1,119 |
| BKK24200 | *yqiH* | -1,361 | 1,190 | 0,163 | 31 | 1,206 |
| BKK38790 | *yxzE* | -1,361 | 1,149 | 0,108 | 112 | 1,165 |
| BKK18800 | *penP* | -1,364 | 0,962 | 0,101 | 39 | 0,975 |
| BKK27690 | *yrzE* | -1,364 | 1,144 | 0,103 | 394 | 1,160 |
| BKK31930 | *ald* | -1,364 | 1,132 | 0,120 | 51 | 1,148 |
| BKK10960 | *yitE* | -1,373 | 1,167 | 0,103 | 143 | 1,183 |
| BKK13100 | *ykkD* | -1,373 | 1,192 | 0,138 | 130 | 1,208 |
| BKK25430 | *rimO* | -1,374 | 1,148 | 0,103 | 365 | 1,164 |
| BKK34920 | *hisG* | -1,377 | 1,119 | 0,113 | 100 | 1,135 |
| BKK35430 | *flgM* | -1,377 | 1,141 | 0,106 | 89 | 1,157 |
| BKK16480 | *ylxL* | -1,38 | 1,145 | 0,117 | 375 | 1,161 |
| BKK11381 | *appA* | -1,386 | 1,167 | 0,107 | 111 | 1,183 |
| BKK10760 | *yisL* | -1,387 | 1,167 | 0,115 | 123 | 1,183 |
| BKK29760 | *ytxJ* | -1,39 | 1,124 | 0,095 | 87 | 1,139 |
| BKK18520 | *yoxB* | -1,391 | 1,141 | 0,105 | 122 | 1,157 |
| BKK34520 | *yvdP* | -1,391 | 1,119 | 0,104 | 77 | 1,135 |
| BKK25740 | *yqeB* | -1,396 | 1,147 | 0,112 | 122 | 1,164 |
| BKK28940 | *ysaA* | -1,398 | 1,098 | 0,120 | 292 | 1,113 |
| BKK40480 | *yycB* | -1,399 | 1,131 | 0,099 | 412 | 1,147 |
| BKK21390 | *yomE* | -1,401 | 1,137 | 0,109 | 434 | 1,153 |
| BKK39120 | *yxiM* | -1,404 | 1,148 | 0,116 | 30 | 1,165 |
| BKK12460 | *xlyB* | -1,41 | 1,172 | 0,109 | 914 | 1,189 |
| BKK33380 | *yvgL* | -1,41 | 1,205 | 0,089 | 94 | 1,222 |
| BKK06310 | *gabP* | -1,411 | 1,143 | 0,104 | 835 | 1,160 |
| BKK19920 | *yotD* | -1,414 | 1,113 | 0,142 | 144 | 1,129 |
| BKK03982 | *mtlF* | -1,419 | 1,148 | 0,084 | 173 | 1,165 |
| BKK17730 | *yndB* | -1,42 | 1,145 | 0,105 | 310 | 1,162 |
| BKK21540 | *yolA* | -1,423 | 1,138 | 0,114 | 236 | 1,155 |
| BKK36640 | *ureC* | -1,424 | 1,140 | 0,111 | 147 | 1,157 |
| BKK37190 | *clsB* | -1,424 | 1,103 | 0,104 | 64 | 1,119 |
| BKK06250 | *ydjM* | -1,425 | 1,162 | 0,107 | 316 | 1,179 |
| BKK30770 | *mntA* | -1,426 | 1,145 | 0,105 | 389 | 1,161 |
| BKK18540 | *yoaB* | -1,428 | 1,122 | 0,093 | 174 | 1,138 |
| BKK33240 | *oxdC* | -1,428 | 1,118 | 0,106 | 149 | 1,134 |
| BKK07940 | *yfkC* | -1,432 | 1,142 | 0,106 | 337 | 1,159 |
| BKK37060 | *tdk* | -1,432 | 1,098 | 0,123 | 30 | 1,114 |
| BKK30080 | *yteV* | -1,434 | 1,139 | 0,106 | 98 | 1,156 |
| BKK11130 | *ipi* | -1,437 | 1,166 | 0,095 | 131 | 1,183 |
| BKK23820 | *yqjM* | -1,437 | 1,137 | 0,105 | 489 | 1,153 |
| BKK15770 | *prkC* | -1,438 | 1,151 | 0,101 | 374 | 1,168 |
| BKK39339 | *yxzL* | -1,441 | 1,131 | 0,105 | 357 | 1,147 |
| BKK29680 | *acsA* | -1,445 | 1,097 | 0,115 | 54 | 1,113 |
| BKK16430 | *cheA* | -1,446 | 1,122 | 0,117 | 107 | 1,138 |
| BKK27469 | *yrzR* | -1,447 | 1,134 | 0,101 | 274 | 1,151 |
| BKK06990 | *yesQ* | -1,452 | 1,191 | 0,193 | 175 | 1,208 |
| BKK35140 | *yvkN* | -1,453 | 1,119 | 0,112 | 363 | 1,135 |
| BKK36880 | *atpI* | -1,454 | 1,098 | 0,135 | 177 | 1,114 |
| BKK02570 | *ycbN* | -1,461 | 1,184 | 0,071 | 153 | 1,201 |
| BKK08390 | *yfiT* | -1,468 | 1,142 | 0,113 | 277 | 1,159 |
| BKK07090 | *yetA* | -1,471 | 1,162 | 0,122 | 231 | 1,179 |
| BKK22590 | *ypiA* | -1,472 | 1,084 | 0,101 | 235 | 1,100 |
| BKK05090 | *yddS* | -1,475 | 1,143 | 0,116 | 439 | 1,160 |
| BKK25230 | *yqxD* | -1,475 | 1,146 | 0,092 | 192 | 1,164 |
| BKK38920 | *pepT* | -1,475 | 1,148 | 0,125 | 149 | 1,165 |
| BKK18580 | *yoaF* | -1,478 | 1,121 | 0,092 | 73 | 1,138 |
| BKK11510 | *yjbE* | -1,479 | 1,166 | 0,107 | 141 | 1,183 |
| BKK00240 | *csfB* | -1,48 | 1,166 | 0,114 | 178 | 1,183 |
| BKK02840 | *ycdG* | -1,48 | 1,113 | 0,095 | 53 | 1,130 |
| BKK17610 | *xylB* | -1,481 | 1,145 | 0,110 | 292 | 1,162 |
| BKK10270 | *lcfB* | -1,482 | 1,166 | 0,086 | 74 | 1,183 |
| BKK16110 | *dprA* | -1,482 | 1,142 | 0,122 | 183 | 1,160 |
| BKK08790 | *thiC* | -1,487 | 1,142 | 0,103 | 187 | 1,159 |
| BKK14430 | *ykpA* | -1,487 | 1,145 | 0,098 | 374 | 1,163 |
| BKK03710 | *gerKC* | -1,488 | 1,121 | 0,090 | 59 | 1,138 |
| BKK08480 | *yfhC* | -1,488 | 1,142 | 0,120 | 318 | 1,159 |
| BKK36100 | *ywrD* | -1,488 | 1,140 | 0,095 | 90 | 1,157 |
| BKK13630 | *ykvA* | -1,49 | 0,960 | 0,069 | 99 | 0,975 |
| BKK14680 | *ykzC* | -1,493 | 1,162 | 0,162 | 93 | 1,179 |
| BKK28790 | *araB* | -1,493 | 1,122 | 0,098 | 118 | 1,139 |
| BKK32070 | *yuiC* | -1,495 | 1,131 | 0,082 | 182 | 1,148 |
| BKK16310 | *fliM* | -1,501 | 1,145 | 0,113 | 212 | 1,163 |
| BKK08510 | *yfhF* | -1,503 | 1,141 | 0,104 | 306 | 1,159 |
| BKK16060 | *rnhB* | -1,503 | 1,147 | 0,096 | 66 | 1,165 |
| BKK19800 | *phy* | -1,503 | 1,144 | 0,115 | 426 | 1,162 |
| BKK35770 | *tagC* | -1,504 | 1,144 | 0,114 | 343 | 1,161 |
| BKK18020 | *sspN* | -1,506 | 1,146 | 0,113 | 69 | 1,164 |
| BKK32520 | *pucG* | -1,513 | 1,204 | 0,064 | 126 | 1,222 |
| BKK17190 | *pksL* | -1,514 | 1,150 | 0,101 | 460 | 1,168 |
| BKK23060 | *ypzE* | -1,514 | 1,140 | 0,088 | 95 | 1,158 |
| BKK11139 | *yizC* | -1,517 | 1,145 | 0,105 | 410 | 1,163 |
| BKK26950 | *yraG* | -1,518 | 1,159 | 0,109 | 169 | 1,177 |
| BKK18120 | *alsT* | -1,521 | 1,154 | 0,123 | 126 | 1,172 |
| BKK24380 | *spoIIIAF* | -1,521 | 1,188 | 0,140 | 54 | 1,206 |
| BKK23980 | *artP* | -1,522 | 1,136 | 0,113 | 432 | 1,153 |
| BKK02390 | *ilvE* | -1,528 | 1,142 | 0,099 | 480 | 1,160 |
| BKK16930 | *cinA* | -1,529 | 1,146 | 0,103 | 109 | 1,164 |
| BKK08300 | *yfiK* | -1,533 | 1,141 | 0,116 | 123 | 1,159 |
| BKK21860 | *yphP* | -1,535 | 1,137 | 0,102 | 356 | 1,155 |
| BKK06078 | *ydzW* | -1,536 | 1,130 | 0,102 | 158 | 1,147 |
| BKK26440 | *yrkO* | -1,536 | 1,159 | 0,108 | 235 | 1,177 |
| BKK39630 | *yxdM* | -1,539 | 1,203 | 0,082 | 74 | 1,222 |
| BKK21760 | *ypmP* | -1,542 | 1,137 | 0,112 | 541 | 1,155 |
| BKK03620 | *yclA* | -1,543 | 1,147 | 0,102 | 218 | 1,165 |
| BKK29900 | *trmB* | -1,543 | 1,143 | 0,115 | 339 | 1,161 |
| BKK30100 | *yteT* | -1,544 | 1,138 | 0,084 | 264 | 1,156 |
| BKK38450 | *ywaE* | -1,544 | 1,140 | 0,095 | 86 | 1,158 |
| BKK21930 | *cspD* | -1,545 | 1,136 | 0,115 | 292 | 1,153 |
| BKK37840 | *spsI* | -1,546 | 1,148 | 0,109 | 96 | 1,166 |
| BKK41060 | *rpmH* | -1,546 | 1,138 | 0,141 | 37 | 1,156 |
| BKK22550 | *qcrB* | -1,549 | 1,083 | 0,122 | 393 | 1,100 |
| BKK23600 | *yqxK* | -1,551 | 1,169 | 0,102 | 227 | 1,187 |
| BKK02460 | *ycbC* | -1,554 | 1,112 | 0,103 | 106 | 1,130 |
| BKK16100 | *sucD* | -1,556 | 1,144 | 0,103 | 270 | 1,163 |
| BKK25330 | *yqfF* | -1,559 | 1,145 | 0,095 | 287 | 1,164 |
| BKK02790 | *ycdB* | -1,56 | 1,112 | 0,101 | 71 | 1,130 |
| BKK38510 | *dltB* | -1,561 | 1,148 | 0,109 | 108 | 1,166 |
| BKK02340 | *gltP* | -1,571 | 1,112 | 0,086 | 113 | 1,130 |
| BKK13380 | *ykoS* | -1,571 | 1,150 | 0,104 | 463 | 1,168 |
| BKK31730 | *yuzC* | -1,571 | 1,122 | 0,085 | 67 | 1,139 |
| BKK11809 | *yjcZ* | -1,572 | 1,170 | 0,088 | 328 | 1,189 |
| BKK30720 | *ythB* | -1,572 | 1,138 | 0,101 | 112 | 1,156 |
| BKK40040 | *glxK* | -1,572 | 1,129 | 0,101 | 458 | 1,147 |
| BKK10120 | *hemE* | -1,574 | 1,158 | 0,118 | 98 | 1,177 |
| BKK29710 | *acuC* | -1,577 | 1,133 | 0,106 | 278 | 1,151 |
| BKK11910 | *yjcM* | -1,578 | 1,170 | 0,112 | 321 | 1,189 |
| BKK33222 | *rsoA* | -1,578 | 1,116 | 0,082 | 107 | 1,134 |
| BKK35360 | *hag* | -1,58 | 1,117 | 0,112 | 490 | 1,135 |
| BKK33610 | *rnr* | -1,581 | 1,116 | 0,098 | 95 | 1,134 |
| BKK17770 | *yndF* | -1,588 | 1,145 | 0,107 | 100 | 1,164 |
| BKK23090 | *rsiX* | -1,588 | 1,138 | 0,103 | 140 | 1,156 |
| BKK40680 | *yybD* | -1,588 | 1,166 | 0,095 | 235 | 1,184 |
| BKK08170 | *yfjA* | -1,59 | 1,140 | 0,107 | 392 | 1,159 |
| BKK16130 | *trmFO* | -1,592 | 1,141 | 0,096 | 555 | 1,160 |
| BKK28110 | *spoVID* | -1,593 | 1,141 | 0,104 | 89 | 1,160 |
| BKK29010 | *speD* | -1,593 | 1,095 | 0,102 | 167 | 1,113 |
| BKK20450 | *yorA* | -1,596 | 1,149 | 0,108 | 282 | 1,168 |
| BKK13050 | *ykzH* | -1,597 | 1,144 | 0,101 | 351 | 1,163 |
| BKK07670 | *yflI* | -1,6 | 1,141 | 0,099 | 508 | 1,160 |
| BKK19410 | *cwlS* | -1,603 | 1,143 | 0,119 | 379 | 1,162 |
| BKK03540 | *ycxB* | -1,605 | 1,112 | 0,097 | 128 | 1,130 |
| BKK20040 | *yosP* | -1,606 | 1,111 | 0,146 | 181 | 1,129 |
| BKK07850 | *yfkM* | -1,608 | 1,141 | 0,100 | 442 | 1,160 |
| BKK32640 | *sspG* | -1,609 | 1,116 | 0,080 | 162 | 1,134 |
| BKK36600 | *mta* | -1,609 | 1,096 | 0,102 | 68 | 1,114 |
| BKK10560 | *yhjM* | -1,611 | 1,164 | 0,102 | 93 | 1,183 |
| BKK17940 | *yneI* | -1,612 | 1,143 | 0,103 | 382 | 1,162 |
| BKK03510 | *srfAC* | -1,613 | 1,112 | 0,105 | 104 | 1,130 |
| BKK24850 | *glcK* | -1,614 | 1,145 | 0,097 | 126 | 1,164 |
| BKK11570 | *yjbJ* | -1,617 | 1,189 | 0,136 | 144 | 1,208 |
| BKK09600 | *crcBA* | -1,619 | 1,158 | 0,098 | 143 | 1,177 |
| BKK05890 | *ydhU* | -1,62 | 1,139 | 0,095 | 125 | 1,158 |
| BKK08900 | *yhzB* | -1,623 | 1,158 | 0,096 | 73 | 1,177 |
| BKK06840 | *yeeI* | -1,629 | 1,160 | 0,112 | 307 | 1,179 |
| BKK25880 | *yqxJ* | -1,632 | 0,959 | 0,070 | 80 | 0,975 |
| BKK01530 | *cwlD* | -1,634 | 0,959 | 0,076 | 49 | 0,975 |
| BKK25580 | *comEB* | -1,639 | 1,144 | 0,109 | 305 | 1,164 |
| BKK38270 | *efeM* | -1,641 | 1,131 | 0,107 | 296 | 1,149 |
| BKK08490 | *yfhD* | -1,643 | 1,140 | 0,101 | 197 | 1,159 |
| BKK20580 | *yoqM* | -1,644 | 1,136 | 0,098 | 448 | 1,155 |
| BKK34590 | *mdxG* | -1,644 | 1,131 | 0,110 | 557 | 1,149 |
| BKK30150 | *ytdP* | -1,647 | 1,137 | 0,087 | 99 | 1,156 |
| BKK37910 | *spsA* | -1,648 | 1,138 | 0,131 | 166 | 1,157 |
| BKK33860 | *yvbH* | -1,653 | 1,142 | 0,107 | 283 | 1,161 |
| BKK39890 | *yxbB* | -1,653 | 1,138 | 0,121 | 98 | 1,158 |
| BKK32050 | *pepA* | -1,655 | 1,129 | 0,097 | 189 | 1,148 |
| BKK07620 | *yflN* | -1,658 | 1,140 | 0,111 | 437 | 1,159 |
| BKK07955 | *yfkA* | -1,658 | 1,143 | 0,102 | 358 | 1,163 |
| BKK08340 | *padR* | -1,659 | 1,143 | 0,103 | 324 | 1,163 |
| BKK26550 | *yrkD* | -1,661 | 1,157 | 0,102 | 241 | 1,177 |
| BKK28590 | *polX* | -1,661 | 1,140 | 0,094 | 41 | 1,160 |
| BKK14280 | *moeA* | -1,667 | 1,159 | 0,105 | 186 | 1,179 |
| BKK12420 | *yjoB* | -1,668 | 1,169 | 0,115 | 635 | 1,189 |
| BKK02270 | *pssA* | -1,669 | 1,111 | 0,092 | 91 | 1,130 |
| BKK33340 | *sspJ* | -1,67 | 1,142 | 0,113 | 342 | 1,161 |
| BKK21890 | *ypgQ* | -1,672 | 1,136 | 0,106 | 342 | 1,155 |
| BKK07710 | *yflE* | -1,674 | 1,119 | 0,100 | 128 | 1,138 |
| BKK32010 | *besA* | -1,675 | 1,129 | 0,126 | 164 | 1,148 |
| BKK38680 | *yxlD* | -1,675 | 1,147 | 0,106 | 219 | 1,166 |
| BKK13520 | *ykrP* | -1,676 | 1,147 | 0,103 | 230 | 1,166 |
| BKK14470 | *mreBH* | -1,677 | 1,143 | 0,105 | 317 | 1,163 |
| BKK26480 | *yrkK* | -1,677 | 1,134 | 0,097 | 437 | 1,153 |

Sup. Table 4: Cell width of mutants of the BKK collection (continued)

| BKK name[1] | gene | screening delta[2] (%) | erage width (µ | +/- | nb | ADP |
|---|---|---|---|---|---|---|
| BKK17030 | *cotE* | -1,679 | 1,186 | 0,120 | 146 | 1,206 |
| BKK29320 | *ytnI* | -1,68 | 1,094 | 0,124 | 219 | 1,113 |
| BKK32240 | *thrB* | -1,681 | 1,129 | 0,128 | 97 | 1,148 |
| BKK10040 | *ecsA* | -1,685 | 1,119 | 0,106 | 117 | 1,138 |
| BKK00140 | *dck* | -1,686 | 1,159 | 0,111 | 494 | 1,179 |
| BKK21960 | *ypeQ* | -1,686 | 1,132 | 0,102 | 312 | 1,151 |
| BKK27270 | *mtnN* | -1,686 | 1,138 | 0,105 | 125 | 1,158 |
| BKK08060 | *acoA* | -1,69 | 1,148 | 0,100 | 355 | 1,168 |
| BKK29720 | *ytxE* | -1,693 | 1,120 | 0,098 | 134 | 1,139 |
| BKK00480 | *yabJ* | -1,696 | 1,144 | 0,104 | 133 | 1,164 |
| BKK16720 | *ymxH* | -1,696 | 1,144 | 0,110 | 151 | 1,164 |
| BKK09780 | *yheC* | -1,699 | 1,157 | 0,100 | 171 | 1,177 |
| BKK39990 | *yxaF* | -1,699 | 1,145 | 0,116 | 71 | 1,165 |
| BKK12280 | *rex* | -1,7 | 1,143 | 0,111 | 267 | 1,163 |
| BKK33960 | *araE* | -1,7 | 1,142 | 0,099 | 359 | 1,161 |
| BKK34970 | *ppaX* | -1,7 | 1,116 | 0,123 | 252 | 1,135 |
| BKK34600 | *mdxF* | -1,704 | 1,116 | 0,102 | 143 | 1,135 |
| BKK18710 | *yozF* | -1,707 | 1,152 | 0,106 | 89 | 1,172 |
| BKK02619 | *yczK* | -1,713 | 1,111 | 0,105 | 61 | 1,130 |
| BKK31560 | *yufP* | -1,713 | 1,146 | 0,104 | 163 | 1,166 |
| BKK05530 | *ydfR* | -1,715 | 1,140 | 0,104 | 319 | 1,160 |
| BKK39840 | *yxbG* | -1,716 | 0,958 | 0,091 | 42 | 0,975 |
| BKK03690 | *yczF* | -1,718 | 1,145 | 0,106 | 134 | 1,165 |
| BKK32560 | *frlR* | -1,72 | 1,120 | 0,092 | 204 | 1,139 |
| BKK30120 | *yteR* | -1,721 | 1,136 | 0,089 | 121 | 1,156 |
| BKK19890 | *yotG* | -1,724 | 1,110 | 0,125 | 487 | 1,129 |
| BKK11330 | *fabHA* | -1,725 | 1,163 | 0,114 | 77 | 1,183 |
| BKK24640 | *yqxM* | -1,725 | 1,131 | 0,100 | 200 | 1,151 |
| BKK36830 | *atpA* | -1,727 | 1,095 | 0,114 | 135 | 1,114 |
| BKK32460 | *pucM* | -1,728 | 1,128 | 0,139 | 50 | 1,148 |
| BKK12940 | *dppC* | -1,729 | 1,187 | 0,151 | 94 | 1,208 |
| BKK26860 | *yraO* | -1,73 | 1,156 | 0,093 | 212 | 1,177 |
| BKK38240 | *ywcA* | -1,732 | 1,099 | 0,144 | 41 | 1,119 |
| BKK25620 | *yqeL* | -1,74 | 1,143 | 0,093 | 205 | 1,164 |
| BKK17690 | *yncM* | -1,741 | 1,142 | 0,097 | 467 | 1,162 |
| BKK04740 | *rsbX* | -1,745 | 1,145 | 0,137 | 276 | 1,165 |
| BKK31440 | *patB* | -1,745 | 1,128 | 0,096 | 187 | 1,148 |
| BKK32870 | *yusO* | -1,745 | 1,115 | 0,106 | 105 | 1,134 |
| BKK07420 | *yfmM* | -1,75 | 1,187 | 0,136 | 44 | 1,208 |
| BKK34140 | *ganQ* | -1,75 | 1,137 | 0,087 | 145 | 1,158 |
| BKK05990 | *tatCY* | -1,759 | 1,139 | 0,110 | 489 | 1,160 |
| BKK17880 | *ynzC* | -1,766 | 1,141 | 0,100 | 341 | 1,162 |
| BKK21640 | *yokC* | -1,773 | 1,133 | 0,100 | 472 | 1,153 |
| BKK39710 | *iolF* | -1,773 | 1,146 | 0,099 | 194 | 1,166 |
| BKK23830 | *yqjL* | -1,776 | 1,166 | 0,112 | 135 | 1,187 |
| BKK22300 | *yppC* | -1,778 | 1,081 | 0,120 | 153 | 1,100 |
| BKK33500 | *copA* | -1,778 | 1,118 | 0,101 | 103 | 1,138 |
| BKK04270 | *ydaJ* | -1,78 | 1,180 | 0,070 | 54 | 1,201 |
| BKK19999 | *yojW* | -1,781 | 1,109 | 0,134 | 310 | 1,129 |
| BKK24080 | *bcd* | -1,787 | 1,166 | 0,112 | 291 | 1,187 |
| BKK11970 | *yjcS* | -1,79 | 1,142 | 0,111 | 337 | 1,163 |
| BKK30595 | *ytlA* | -1,791 | 1,200 | 0,059 | 243 | 1,222 |
| BKK21110 | *yonF* | -1,793 | 1,134 | 0,116 | 243 | 1,155 |
| BKK32480 | *pucD* | -1,793 | 1,145 | 0,108 | 84 | 1,166 |
| BKK13990 | *kinA* | -1,798 | 1,163 | 0,113 | 98 | 1,184 |
| BKK05780 | *ydhJ* | -1,799 | 1,139 | 0,099 | 444 | 1,160 |
| BKK32360 | *yunC* | -1,799 | 1,127 | 0,115 | 101 | 1,148 |
| BKK03981 | *mtlA* | -1,801 | 1,144 | 0,089 | 125 | 1,165 |
| BKK10940 | *yitC* | -1,801 | 1,162 | 0,112 | 128 | 1,183 |
| BKK15720 | *defA* | -1,801 | 1,114 | 0,115 | 85 | 1,134 |
| BKK26740 | *cypA* | -1,802 | 1,155 | 0,101 | 376 | 1,177 |
| BKK35090 | *yvmA* | -1,802 | 1,129 | 0,097 | 453 | 1,149 |
| BKK08100 | *acoR* | -1,805 | 1,138 | 0,098 | 462 | 1,159 |
| BKK12390 | *uxaA* | -1,805 | 1,147 | 0,105 | 422 | 1,168 |
| BKK39390 | *hutM* | -1,809 | 1,144 | 0,107 | 106 | 1,165 |
| BKK22930 | *sleB* | -1,81 | 1,080 | 0,128 | 100 | 1,100 |
| BKK07430 | *yfmL* | -1,811 | 1,139 | 0,109 | 434 | 1,160 |
| BKK22810 | *yphF* | -1,812 | 1,185 | 0,126 | 147 | 1,206 |
| BKK07530 | *yfmB* | -1,815 | 1,138 | 0,106 | 496 | 1,159 |
| BKK36660 | *ureA* | -1,815 | 1,094 | 0,121 | 218 | 1,114 |
| BKK33850 | *yvbG* | -1,819 | 0,957 | 0,071 | 65 | 0,975 |
| BKK14110 | *ykuK* | -1,821 | 1,141 | 0,111 | 232 | 1,163 |
| BKK34160 | *cycB* | -1,821 | 1,129 | 0,114 | 428 | 1,149 |
| BKK34820 | *bmrA* | -1,823 | 1,128 | 0,094 | 590 | 1,149 |
| BKK25600 | *comER* | -1,825 | 1,142 | 0,094 | 229 | 1,164 |
| BKK01060 | *ybxB* | -1,83 | 1,142 | 0,097 | 388 | 1,164 |
| BKK06079 | *ydzW* | -1,832 | 1,136 | 0,101 | 98 | 1,158 |
| BKK15460 | *rluD* | -1,835 | 1,093 | 0,110 | 129 | 1,113 |
| BKK31850 | *yueC* | -1,837 | 1,127 | 0,115 | 237 | 1,148 |

| BKK name[1] | gene | screening delta[2] (%) | erage width (µ | +/- | nb | ADP |
|---|---|---|---|---|---|---|
| BKK38180 | *ywzA* | -1,84 | 1,136 | 0,100 | 88 | 1,157 |
| BKK03140 | *tmrB* | -1,844 | 1,109 | 0,093 | 113 | 1,130 |
| BKK10080 | *yhfA* | -1,844 | 1,186 | 0,151 | 72 | 1,208 |
| BKK24580 | *yqhH* | -1,844 | 1,136 | 0,093 | 114 | 1,158 |
| BKK11820 | *yjcD* | -1,85 | 1,141 | 0,099 | 347 | 1,163 |
| BKK19350 | *yocS* | -1,853 | 1,184 | 0,144 | 83 | 1,206 |
| BKK40170 | *yydG* | -1,854 | 1,146 | 0,126 | 284 | 1,168 |
| BKK39750 | *iolB* | -1,855 | 1,143 | 0,113 | 74 | 1,165 |
| BKK36350 | *ywpD* | -1,858 | 1,140 | 0,101 | 406 | 1,161 |
| BKK39180 | *yxiH* | -1,859 | 1,143 | 0,095 | 72 | 1,165 |
| BKK39970 | *yxaH* | -1,859 | 1,128 | 0,097 | 551 | 1,149 |
| BKK11790 | *yjcA* | -1,861 | 1,167 | 0,132 | 759 | 1,189 |
| BKK01935 | *skfC* | -1,866 | 1,092 | 0,095 | 102 | 1,113 |
| BKK16650 | *rbfA* | -1,867 | 1,142 | 0,135 | 97 | 1,164 |
| BKK21329 | *youB* | -1,867 | 1,132 | 0,099 | 450 | 1,153 |
| BKK34460 | *levB* | -1,867 | 1,114 | 0,116 | 246 | 1,135 |
| BKK37710 | *bacD* | -1,868 | 1,128 | 0,095 | 609 | 1,149 |
| BKK03890 | *gabR* | -1,873 | 1,143 | 0,129 | 557 | 1,165 |
| BKK06570 | *yerB* | -1,876 | 1,179 | 0,088 | 145 | 1,201 |
| BKK04910 | *yddB* | -1,881 | 1,126 | 0,096 | 417 | 1,147 |
| BKK32000 | *dhbA* | -1,881 | 1,118 | 0,089 | 39 | 1,139 |
| BKK39080 | *licT* | -1,883 | 1,143 | 0,098 | 69 | 1,165 |
| BKK26559 | *yrzN* | -1,884 | 1,155 | 0,108 | 337 | 1,177 |
| BKK20420 | *yorD* | -1,887 | 1,140 | 0,105 | 457 | 1,162 |
| BKK38140 | *qoxD* | -1,887 | 1,135 | 0,089 | 126 | 1,157 |
| BKK01240 | *rpmC* | -1,888 | 1,126 | 0,106 | 154 | 1,147 |
| BKK07070 | *yesY* | -1,888 | 1,157 | 0,135 | 181 | 1,179 |
| BKK32370 | *yunD* | -1,889 | 1,126 | 0,118 | 44 | 1,148 |
| BKK37230 | *ywjA* | -1,892 | 1,126 | 0,106 | 474 | 1,147 |
| BKK20780 | *yopS* | -1,893 | 1,140 | 0,105 | 463 | 1,162 |
| BKK18740 | *yozG* | -1,894 | 1,149 | 0,125 | 113 | 1,172 |
| BKK09259 | *yhzG* | -1,897 | 1,141 | 0,101 | 340 | 1,163 |
| BKK17860 | *yneA* | -1,9 | 1,141 | 0,121 | 79 | 1,164 |
| BKK25960 | *yqcB* | -1,901 | 1,141 | 0,112 | 132 | 1,164 |
| BKK22330 | *ypoC* | -1,905 | 1,129 | 0,101 | 491 | 1,151 |
| BKK37700 | *bacE* | -1,91 | 1,127 | 0,113 | 257 | 1,149 |
| BKK02970 | *yceK* | -1,913 | 1,108 | 0,089 | 99 | 1,130 |
| BKK01750 | *ybbP* | -1,915 | 1,137 | 0,087 | 209 | 1,160 |
| BKK25130 | *nfo* | -1,917 | 1,141 | 0,097 | 250 | 1,164 |
| BKK29450 | *argG* | -1,919 | 1,129 | 0,102 | 330 | 1,151 |
| BKK08960 | *yhbF* | -1,923 | 1,154 | 0,096 | 209 | 1,177 |
| BKK26570 | *yrkB* | -1,923 | 1,131 | 0,102 | 355 | 1,153 |
| BKK16250 | *fliJ* | -1,927 | 1,146 | 0,102 | 344 | 1,168 |
| BKK21590 | *yokH* | -1,929 | 1,133 | 0,109 | 426 | 1,155 |
| BKK26075 | *yqbK* | -1,933 | 1,129 | 0,105 | 474 | 1,151 |
| BKK06420 | *purE* | -1,938 | 1,156 | 0,124 | 258 | 1,179 |
| BKK40310 | *phrG* | -1,938 | 1,129 | 0,097 | 459 | 1,151 |
| BKK13790 | *ykvQ* | -1,939 | 1,144 | 0,107 | 90 | 1,166 |
| BKK33100 | *liaF* | -1,941 | 1,112 | 0,095 | 174 | 1,134 |
| BKK27380 | *yrzB* | -1,946 | 1,129 | 0,102 | 372 | 1,151 |
| BKK02150 | *ybeF* | -1,948 | 1,108 | 0,113 | 74 | 1,130 |
| BKK13240 | *thiU* | -1,95 | 1,144 | 0,117 | 52 | 1,166 |
| BKK17540 | *ynaF* | -1,95 | 1,139 | 0,107 | 360 | 1,162 |
| BKK02070 | *csgA* | -1,952 | 1,108 | 0,093 | 59 | 1,130 |
| BKK28410 | *gerE* | -1,952 | 1,137 | 0,093 | 132 | 1,160 |
| BKK02060 | *ybxG* | -1,953 | 1,108 | 0,078 | 45 | 1,130 |
| BKK30240 | *bioW* | -1,956 | 1,133 | 0,098 | 264 | 1,156 |
| BKK14580 | *pdhA* | -1,958 | 1,156 | 0,161 | 118 | 1,179 |
| BKK35540 | *tuaH* | -1,961 | 1,127 | 0,100 | 691 | 1,149 |
| BKK35630 | *lytB* | -1,961 | 1,127 | 0,100 | 691 | 1,149 |
| BKK19330 | *sodF* | -1,968 | 1,149 | 0,112 | 116 | 1,172 |
| BKK37140 | *ywjG* | -1,968 | 1,097 | 0,112 | 690 | 1,119 |
| BKK31300 | *yugS* | -1,97 | 1,138 | 0,102 | 296 | 1,161 |
| BKK38499 | *ywzH* | -1,972 | 1,097 | 0,122 | 85 | 1,119 |
| BKK24280 | *ispA* | -1,974 | 1,164 | 0,099 | 281 | 1,187 |
| BKK25040 | *yqgB* | -1,974 | 1,141 | 0,114 | 149 | 1,164 |
| BKK36530 | *bcrC* | -1,975 | 1,092 | 0,073 | 51 | 1,114 |
| BKK03730 | *yclH* | -1,976 | 1,178 | 0,116 | 979 | 1,201 |
| BKK27760 | *csbX* | -1,977 | 1,137 | 0,113 | 408 | 1,160 |
| BKK15330 | *sigG* | -1,978 | 1,117 | 0,114 | 66 | 1,139 |
| BKK00350 | *yazA* | -1,982 | 1,156 | 0,155 | 152 | 1,179 |
| BKK14420 | *ykoA* | -1,984 | 1,156 | 0,117 | 450 | 1,179 |
| BKK24710 | *comGC* | -1,985 | 1,140 | 0,091 | 464 | 1,164 |
| BKK32940 | *yusV* | -1,985 | 1,198 | 0,098 | 38 | 1,222 |
| BKK35240 | *ctpB* | -1,988 | 1,113 | 0,131 | 420 | 1,135 |
| BKK13130 | *proA* | -1,989 | 1,143 | 0,113 | 173 | 1,166 |
| BKK16790 | *tepA* | -1,99 | 1,140 | 0,117 | 96 | 1,164 |
| BKK27590 | *dtd* | -1,991 | 1,137 | 0,095 | 316 | 1,160 |
| BKK08570 | *yfhK* | -1,996 | 1,136 | 0,118 | 187 | 1,159 |

| BKK name[1] | gene | screening delta[2] (%) | erage width (µ | +/- | nb | ADP |
|---|---|---|---|---|---|---|
| BKK32020 | *yuiH* | -2,006 | 1,125 | 0,107 | 138 | 1,148 |
| BKK35100 | *yvlD* | -2,007 | 1,112 | 0,106 | 409 | 1,135 |
| BKK37500 | *speE* | -2,01 | 1,096 | 0,135 | 44 | 1,119 |
| BKK12740 | *xkdU* | -2,012 | 1,143 | 0,120 | 135 | 1,166 |
| BKK33140 | *yvqJ* | -2,017 | 1,197 | 0,083 | 140 | 1,222 |
| BKK27040 | *levG* | -2,019 | 1,153 | 0,110 | 255 | 1,177 |
| BKK06036 | *ydzT* | -2,026 | 1,136 | 0,101 | 346 | 1,160 |
| BKK17210 | *pksN* | -2,03 | 1,140 | 0,106 | 81 | 1,164 |
| BKK20600 | *yoqK* | -2,031 | 1,131 | 0,114 | 250 | 1,155 |
| BKK05550 | *cotP* | -2,033 | 1,124 | 0,104 | 346 | 1,147 |
| BKK20230 | *yorW* | -2,033 | 1,144 | 0,114 | 250 | 1,168 |
| BKK02640 | *tatCD* | -2,034 | 1,177 | 0,071 | 79 | 1,201 |
| BKK32140 | *paiB* | -2,035 | 1,125 | 0,098 | 42 | 1,148 |
| BKK20120 | *yosH* | -2,036 | 1,106 | 0,120 | 100 | 1,129 |
| BKK09190 | *yhcR* | -2,037 | 1,153 | 0,104 | 120 | 1,177 |
| BKK12570 | *xtmA* | -2,037 | 1,165 | 0,129 | 436 | 1,189 |
| BKK39900 | *aslA* | -2,041 | 1,142 | 0,108 | 112 | 1,166 |
| BKK24900 | *rpmGA* | -2,043 | 1,140 | 0,095 | 211 | 1,164 |
| BKK34550 | *pgcM* | -2,046 | 1,112 | 0,109 | 267 | 1,135 |
| BKK00560 | *spoVT* | -2,048 | 0,955 | 0,075 | 44 | 0,975 |
| BKK21000 | *yonT* | -2,049 | 1,130 | 0,105 | 585 | 1,153 |
| BKK31319 | *yuzI* | -2,049 | 1,132 | 0,113 | 69 | 1,156 |
| BKK24490 | *yqhQ* | -2,05 | 1,163 | 0,094 | 377 | 1,187 |
| BKK28780 | *araD* | -2,051 | 1,090 | 0,092 | 351 | 1,113 |
| BKK25920 | *yqxG* | -2,054 | 1,140 | 0,130 | 58 | 1,164 |
| BKK18590 | *yoaG* | -2,055 | 1,115 | 0,094 | 104 | 1,138 |
| BKK05840 | *gmuD* | -2,056 | 1,143 | 0,077 | 244 | 1,167 |
| BKK09110 | *yhcJ* | -2,056 | 1,183 | 0,142 | 691 | 1,208 |
| BKK27660 | *comN* | -2,058 | 1,136 | 0,107 | 232 | 1,160 |
| BKK20710 | *yopZ* | -2,06 | 1,131 | 0,104 | 381 | 1,155 |
| BKK21010 | *yonS* | -2,06 | 1,182 | 0,106 | 94 | 1,206 |
| BKK33250 | *yvrL* | -2,06 | 1,137 | 0,104 | 253 | 1,161 |
| BKK02730 | *natK* | -2,062 | 1,177 | 0,059 | 52 | 1,201 |
| BKK30360 | *yttA* | -2,065 | 1,132 | 0,101 | 148 | 1,156 |
| BKK05820 | *gmuA* | -2,068 | 1,124 | 0,095 | 395 | 1,147 |
| BKK26530 | *yrkF* | -2,07 | 1,130 | 0,109 | 398 | 1,153 |
| BKK35239 | *swrAA* | -2,071 | 1,112 | 0,123 | 410 | 1,135 |
| BKK37740 | *bacA* | -2,072 | 1,096 | 0,098 | 90 | 1,119 |
| BKK15630 | *sirC* | -2,074 | 1,124 | 0,112 | 433 | 1,148 |
| BKK13560 | *mtnK* | -2,076 | 1,138 | 0,107 | 430 | 1,163 |
| BKK21130 | *yonD* | -2,079 | 1,131 | 0,097 | 406 | 1,155 |
| BKK30870 | *ytcB* | -2,08 | 1,197 | 0,063 | 445 | 1,222 |
| BKK12240 | *yjkA* | -2,087 | 1,183 | 0,158 | 459 | 1,208 |
| BKK05800 | *pbuE* | -2,088 | 1,176 | 0,051 | 59 | 1,201 |
| BKK06000 | *ydiK* | -2,091 | 1,143 | 0,098 | 204 | 1,167 |
| BKK11890 | *yjcK* | -2,094 | 1,164 | 0,121 | 356 | 1,189 |
| BKK06190 | *ydjG* | -2,096 | 1,154 | 0,103 | 168 | 1,179 |
| BKK05760 | *ydhH* | -2,098 | 1,143 | 0,131 | 139 | 1,167 |
| BKK24700 | *comGD* | -2,099 | 1,127 | 0,104 | 387 | 1,151 |
| BKK33970 | *araR* | -2,101 | 1,133 | 0,091 | 124 | 1,158 |
| BKK12840 | *pit* | -2,103 | 1,183 | 0,132 | 80 | 1,208 |
| BKK34740 | *crh* | -2,105 | 1,111 | 0,094 | 379 | 1,135 |
| BKK26690 | *brnQ* | -2,114 | 1,196 | 0,086 | 222 | 1,222 |
| BKK02320 | *ybfP* | -2,117 | 1,106 | 0,103 | 125 | 1,130 |
| BKK37490 | *speB* | -2,117 | 1,095 | 0,099 | 57 | 1,119 |
| BKK07520 | *yfmC* | -2,119 | 1,183 | 0,125 | 181 | 1,208 |
| BKK12480 | *yjqB* | -2,126 | 1,164 | 0,147 | 729 | 1,189 |
| BKK36090 | *ywrE* | -2,126 | 1,091 | 0,098 | 458 | 1,114 |
| BKK07460 | *yfmI* | -2,131 | 1,135 | 0,110 | 546 | 1,160 |
| BKK25670 | *yqeH* | -2,131 | 1,127 | 0,133 | 130 | 1,151 |
| BKK37800 | *yweA* | -2,137 | 1,125 | 0,105 | 439 | 1,149 |
| BKK08240 | *catE* | -2,141 | 1,134 | 0,109 | 292 | 1,159 |
| BKK04600 | *ydbT* | -2,146 | 1,123 | 0,098 | 421 | 1,147 |
| BKK30220 | *bioF* | -2,15 | 1,131 | 0,103 | 114 | 1,156 |
| BKK31780 | *yueH* | -2,152 | 1,123 | 0,107 | 205 | 1,148 |
| BKK06790 | *yeeD* | -2,157 | 1,135 | 0,096 | 487 | 1,160 |
| BKK21430 | *bhlB* | -2,158 | 1,130 | 0,103 | 474 | 1,155 |
| BKK18450 | *gltA* | -2,159 | 1,113 | 0,087 | 57 | 1,138 |
| BKK10920 | *yitA* | -2,163 | 1,158 | 0,114 | 121 | 1,183 |
| BKK38420 | *sacY* | -2,163 | 1,095 | 0,113 | 183 | 1,119 |
| BKK01610 | *feuC* | -2,166 | 1,151 | 0,095 | 232 | 1,177 |
| BKK38390 | *ywbA* | -2,167 | 1,132 | 0,124 | 61 | 1,157 |
| BKK16010 | *ylqD* | -2,168 | 1,132 | 0,107 | 126 | 1,157 |
| BKK24030 | *bkdB* | -2,168 | 1,161 | 0,108 | 173 | 1,187 |
| BKK22130 | *kduI* | -2,171 | 1,076 | 0,137 | 144 | 1,100 |
| BKK29520 | *yteJ* | -2,178 | 1,089 | 0,105 | 72 | 1,113 |
| BKK09670 | *dat* | -2,179 | 1,182 | 0,127 | 45 | 1,208 |
| BKK32060 | *yuiD* | -2,18 | 1,115 | 0,109 | 82 | 1,139 |
| BKK25850 | *yqzI* | -2,181 | 1,138 | 0,112 | 104 | 1,164 |

| BKK name[1] | gene | screening delta[2] (%) | erage width (µ | +/- | nb | ADP |
|---|---|---|---|---|---|---|
| BKK06350 | *yebA* | -2,185 | 1,153 | 0,110 | 460 | 1,179 |
| BKK13190 | *ispA* | -2,185 | 1,141 | 0,085 | 56 | 1,166 |
| BKK03550 | *ycxC* | -2,186 | 1,105 | 0,097 | 113 | 1,130 |
| BKK36070 | *cotG* | -2,187 | 1,131 | 0,117 | 91 | 1,156 |
| BKK17250 | *ymaE* | -2,189 | 1,136 | 0,107 | 575 | 1,162 |
| BKK26749 | *yrdD* | -2,189 | 1,151 | 0,097 | 166 | 1,177 |
| BKK32210 | *yuzD* | -2,189 | 1,123 | 0,093 | 38 | 1,148 |
| BKK05240 | *ydeL* | -2,19 | 1,114 | 0,105 | 58 | 1,139 |
| BKK01400 | *rpmJ* | -2,192 | 1,134 | 0,129 | 188 | 1,160 |
| BKK11480 | *yjbB* | -2,195 | 1,182 | 0,120 | 87 | 1,208 |
| BKK28580 | *mutSB* | -2,198 | 1,134 | 0,107 | 46 | 1,160 |
| BKK31840 | *yueD* | -2,199 | 1,123 | 0,093 | 145 | 1,148 |
| BKK15030 | *ylbJ* | -2,2 | 1,153 | 0,138 | 261 | 1,179 |
| BKK26220 | *yqaQ* | -2,2 | 1,151 | 0,094 | 408 | 1,177 |
| BKK40090 | *ahpC* | -2,201 | 1,142 | 0,109 | 340 | 1,168 |
| BKK26000 | *yqbR* | -2,206 | 1,126 | 0,092 | 459 | 1,151 |
| BKK37270 | *narH* | -2,206 | 1,094 | 0,108 | 90 | 1,119 |
| BKK25630 | *yqeK* | -2,213 | 1,138 | 0,099 | 142 | 1,164 |
| BKK34830 | *yvzA* | -2,215 | 1,110 | 0,127 | 164 | 1,135 |
| BKK11360 | *appD* | -2,22 | 1,182 | 0,143 | 116 | 1,208 |
| BKK25540 | *gpr* | -2,22 | 1,138 | 0,093 | 189 | 1,164 |
| BKK04470 | *dctP* | -2,221 | 1,132 | 0,095 | 228 | 1,158 |
| BKK05329 | *ydzO* | -2,222 | 1,141 | 0,107 | 146 | 1,167 |
| BKK33770 | *spbC* | -2,223 | 1,135 | 0,097 | 572 | 1,161 |
| BKK13150 | *ohrR* | -2,225 | 1,140 | 0,125 | 195 | 1,166 |
| BKK01760 | *ybbR* | -2,226 | 1,134 | 0,104 | 558 | 1,160 |
| BKK11220 | *argD* | -2,235 | 1,142 | 0,109 | 390 | 1,168 |
| BKK19150 | *yocB* | -2,239 | 1,136 | 0,103 | 468 | 1,162 |
| BKK34510 | *yvdQ* | -2,239 | 1,135 | 0,101 | 418 | 1,161 |
| BKK40200 | *yydD* | -2,239 | 1,125 | 0,102 | 270 | 1,151 |
| BKK30440 | *ytrC* | -2,249 | 1,195 | 0,088 | 239 | 1,222 |
| BKK18530 | *yoaA* | -2,25 | 1,132 | 0,108 | 113 | 1,158 |
| BKK18780 | *yoaW* | -2,251 | 1,145 | 0,095 | 267 | 1,172 |
| BKK00680 | *hprT* | -2,252 | 1,122 | 0,103 | 265 | 1,147 |
| BKK11810 | *spoVIF* | -2,255 | 1,162 | 0,121 | 413 | 1,189 |
| BKK37770 | *rocB* | -2,255 | 1,094 | 0,123 | 46 | 1,119 |
| BKK02890 | *yceC* | -2,256 | 1,131 | 0,095 | 132 | 1,158 |
| BKK29140 | *citZ* | -2,261 | 1,088 | 0,138 | 132 | 1,113 |
| BKK39560 | *yxeG* | -2,262 | 1,138 | 0,092 | 115 | 1,165 |
| BKK34170 | *ganR* | -2,265 | 1,109 | 0,121 | 92 | 1,135 |
| BKK09870 | *khtS* | -2,266 | 1,181 | 0,180 | 175 | 1,208 |
| BKK25750 | *nucB* | -2,267 | 0,953 | 0,064 | 47 | 0,975 |
| BKK26619 | *yrzO* | -2,271 | 1,150 | 0,109 | 114 | 1,177 |
| BKK05510 | *ydfQ* | -2,272 | 1,141 | 0,106 | 647 | 1,167 |
| BKK07640 | *yflL* | -2,272 | 1,132 | 0,106 | 495 | 1,159 |
| BKK13500 | *ktrD* | -2,274 | 1,140 | 0,101 | 289 | 1,166 |
| BKK15050 | *ylbL* | -2,275 | 1,114 | 0,107 | 81 | 1,139 |
| BKK00550 | *mfd* | -2,278 | 1,137 | 0,113 | 86 | 1,164 |
| BKK20810 | *yopP* | -2,28 | 1,125 | 0,108 | 268 | 1,151 |
| BKK27050 | *levF* | -2,282 | 1,150 | 0,094 | 349 | 1,177 |
| BKK07970 | *yfjT* | -2,289 | 1,132 | 0,114 | 252 | 1,159 |
| BKK02720 | *yccF* | -2,29 | 1,174 | 0,082 | 45 | 1,201 |
| BKK36580 | *ywnF* | -2,291 | 1,089 | 0,141 | 214 | 1,114 |
| BKK13980 | *pbpH* | -2,294 | 1,136 | 0,107 | 403 | 1,163 |
| BKK38810 | *msmX* | -2,294 | 1,181 | 0,162 | 257 | 1,208 |
| BKK30930 | *ytaB* | -2,296 | 1,135 | 0,100 | 429 | 1,161 |
| BKK13930 | *splB* | -2,299 | 1,140 | 0,131 | 47 | 1,166 |
| BKK33410 | *yvgO* | -2,3 | 1,108 | 0,079 | 85 | 1,134 |
| BKK21650 | *yokB* | -2,301 | 1,127 | 0,106 | 388 | 1,153 |
| BKK26558 | *yrzM* | -2,302 | 1,127 | 0,104 | 248 | 1,153 |
| BKK39250 | *yxiE* | -2,304 | 1,138 | 0,117 | 85 | 1,165 |
| BKK10470 | *yhjD* | -2,305 | 1,136 | 0,106 | 422 | 1,163 |
| BKK28930 | *lytS* | -2,308 | 1,125 | 0,115 | 236 | 1,151 |
| BKK10090 | *yhgB* | -2,313 | 1,150 | 0,096 | 126 | 1,177 |
| BKK00040 | *recF* | -2,314 | 1,140 | 0,135 | 688 | 1,167 |
| BKK04580 | *cshA* | -2,315 | 1,133 | 0,121 | 225 | 1,160 |
| BKK17960 | *yneK* | -2,316 | 1,135 | 0,112 | 478 | 1,162 |
| BKK33350 | *yvsG* | -2,316 | 1,108 | 0,100 | 127 | 1,134 |
| BKK01820 | *adaB* | -2,321 | 1,133 | 0,101 | 151 | 1,160 |
| BKK26230 | *yqaP* | -2,327 | 1,149 | 0,086 | 152 | 1,177 |
| BKK39000 | *yxjC* | -2,327 | 1,138 | 0,102 | 117 | 1,165 |
| BKK26130 | *yqbF* | -2,33 | 0,952 | 0,105 | 50 | 0,975 |
| BKK10860 | *yisT* | -2,332 | 1,156 | 0,106 | 103 | 1,183 |
| BKK20770 | *yopT* | -2,332 | 1,135 | 0,103 | 225 | 1,162 |
| BKK17150 | *pksG* | -2,333 | 1,136 | 0,105 | 116 | 1,164 |
| BKK38570 | *licA* | -2,335 | 1,138 | 0,092 | 79 | 1,165 |
| BKK22860 | *yphA* | -2,337 | 1,178 | 0,119 | 103 | 1,206 |
| BKK28920 | *lytT* | -2,346 | 1,193 | 0,093 | 69 | 1,222 |
| BKK39870 | *yxbD* | -2,348 | 1,137 | 0,097 | 99 | 1,165 |

### Sup. Table 4: Cell width of mutants of the BKK collection (continued)

| BKK name [1] | gene | screening delta$^2$ (%) | erage width (µ | +/- | nb | ADP |
|---|---|---|---|---|---|---|
| BKK33420 | *nhaK* | -2,351 | 1,134 | 0,101 | 490 | 1,161 |
| BKK14710 | *ylaA* | -2,355 | 1,151 | 0,113 | 150 | 1,179 |
| BKK19990 | *yosV* | -2,356 | 1,141 | 0,108 | 317 | 1,168 |
| BKK32470 | *pucE* | -2,357 | 1,113 | 0,096 | 129 | 1,139 |
| BKK01470 | *ybaF* | -2,358 | 1,136 | 0,092 | 172 | 1,164 |
| BKK05560 | *ydgA* | -2,36 | 1,130 | 0,103 | 167 | 1,158 |
| BKK12110 | *yjfA* | -2,36 | 1,161 | 0,135 | 210 | 1,189 |
| BKK40180 | *yydF* | -2,363 | 1,140 | 0,113 | 245 | 1,168 |
| BKK10280 | *yhfM* | -2,366 | 1,155 | 0,091 | 112 | 1,183 |
| BKK39820 | *htpG* | -2,366 | 1,137 | 0,086 | 112 | 1,165 |
| BKK01350 | *rplO* | -2,372 | 1,136 | 0,099 | 135 | 1,164 |
| BKK38860 | *galE* | -2,375 | 1,137 | 0,119 | 81 | 1,165 |
| BKK29110 | *phoP* | -2,376 | 1,087 | 0,104 | 34 | 1,113 |
| BKK29239 | *ytrH* | -2,378 | 1,087 | 0,080 | 70 | 1,113 |
| BKK16450 | *cheC* | -2,379 | 1,134 | 0,102 | 380 | 1,161 |
| BKK15540 | *pyrD* | -2,384 | 1,128 | 0,122 | 73 | 1,156 |
| BKK19259 | *yoyB* | -2,384 | 1,144 | 0,108 | 70 | 1,172 |
| BKK20220 | *yorX* | -2,386 | 1,112 | 0,112 | 59 | 1,139 |
| BKK14390 | *fruK* | -2,387 | 1,151 | 0,141 | 131 | 1,179 |
| BKK20380 | *yorH* | -2,39 | 1,102 | 0,124 | 48 | 1,129 |
| BKK25980 | *yqbT* | -2,392 | 0,952 | 0,072 | 120 | 0,975 |
| BKK39580 | *yxeE* | -2,394 | 1,137 | 0,089 | 128 | 1,165 |
| BKK17050 | *mutL* | -2,398 | 1,136 | 0,104 | 144 | 1,164 |
| BKK16620 | *rplGA* | -2,399 | 1,130 | 0,100 | 81 | 1,158 |
| BKK06049 | *ydzV* | -2,404 | 1,139 | 0,111 | 125 | 1,167 |
| BKK14640 | *yktA* | -2,404 | 1,135 | 0,110 | 295 | 1,163 |
| BKK30270 | *msmE* | -2,414 | 1,111 | 0,096 | 110 | 1,138 |
| BKK37300 | *ywiC* | -2,416 | 1,092 | 0,110 | 94 | 1,119 |
| BKK12920 | *dppA* | -2,417 | 1,138 | 0,135 | 52 | 1,166 |
| BKK27800 | *yrzH* | -2,42 | 1,123 | 0,114 | 398 | 1,151 |
| BKK31322 | *yugO* | -2,421 | 1,128 | 0,104 | 48 | 1,156 |
| BKK27160 | *cypB* | -2,422 | 1,148 | 0,110 | 280 | 1,177 |
| BKK26610 | *yrkA* | -2,425 | 1,148 | 0,121 | 121 | 1,177 |
| BKK24740 | *yqxL* | -2,426 | 1,177 | 0,128 | 71 | 1,206 |
| BKK20960 | *yopA* | -2,427 | 1,127 | 0,105 | 270 | 1,155 |
| BKK30280 | *amyD* | -2,427 | 1,110 | 0,101 | 68 | 1,138 |
| BKK07040 | *yesV* | -2,431 | 1,150 | 0,111 | 337 | 1,179 |
| BKK22800 | *spoIVA* | -2,434 | 1,112 | 0,092 | 55 | 1,139 |
| BKK25510 | *lepA* | -2,435 | 1,135 | 0,103 | 312 | 1,164 |
| BKK26750 | *yrdD* | -2,437 | 1,148 | 0,105 | 326 | 1,177 |
| BKK35110 | *yvlC* | -2,438 | 1,107 | 0,103 | 428 | 1,135 |
| BKK10590 | *yhjP* | -2,439 | 1,134 | 0,100 | 274 | 1,163 |
| BKK27870 | *nadB* | -2,441 | 1,131 | 0,099 | 246 | 1,160 |
| BKK08040 | *yfjM* | -2,442 | 1,131 | 0,097 | 529 | 1,159 |
| BKK19480 | *yojE* | -2,442 | 1,102 | 0,101 | 828 | 1,129 |
| BKK12310 | *exuM* | -2,445 | 1,179 | 0,120 | 75 | 1,208 |
| BKK18880 | *yobE* | -2,448 | 1,133 | 0,117 | 361 | 1,162 |
| BKK27240 | *yrhC* | -2,448 | 1,148 | 0,109 | 166 | 1,177 |
| BKK03450 | *hxlB* | -2,454 | 1,102 | 0,094 | 30 | 1,130 |
| BKK17900 | *yneE* | -2,456 | 1,135 | 0,085 | 105 | 1,164 |
| BKK12800 | *xhlB* | -2,463 | 1,138 | 0,129 | 184 | 1,166 |
| BKK10840 | *degA* | -2,473 | 1,154 | 0,100 | 142 | 1,183 |
| BKK32650 | *yurS* | -2,474 | 1,137 | 0,105 | 138 | 1,166 |
| BKK34030 | *yvbY* | -2,474 | 1,106 | 0,091 | 88 | 1,134 |
| BKK13660 | *kinD* | -2,476 | 1,177 | 0,160 | 152 | 1,206 |
| BKK30210 | *bioD* | -2,479 | 1,127 | 0,118 | 81 | 1,156 |
| BKK11928 | *yjzF* | -2,488 | 1,139 | 0,105 | 531 | 1,168 |
| BKK17360 | *ymzA* | -2,491 | 1,135 | 0,107 | 132 | 1,164 |
| BKK38740 | *cydC* | -2,491 | 1,172 | 0,129 | 459 | 1,201 |
| BKK14490 | *kinC* | -2,494 | 1,176 | 0,151 | 48 | 1,206 |
| BKK39300 | *yxiD* | -2,497 | 1,137 | 0,094 | 81 | 1,166 |
| BKK12350 | *uxuB* | -2,498 | 1,159 | 0,141 | 177 | 1,189 |
| BKK16350 | *fliP* | -2,498 | 1,133 | 0,108 | 581 | 1,162 |
| BKK15610 | *sumT* | -2,501 | 1,119 | 0,100 | 165 | 1,148 |
| BKK30880 | *ytcC* | -2,505 | 1,132 | 0,107 | 301 | 1,161 |
| BKK40050 | *gntR* | -2,516 | 1,119 | 0,119 | 149 | 1,147 |
| BKK12360 | *exuT* | -2,521 | 1,178 | 0,149 | 104 | 1,208 |
| BKK18940 | *yozK* | -2,523 | 1,142 | 0,105 | 362 | 1,172 |
| BKK19460 | *yojG* | -2,523 | 1,101 | 0,120 | 194 | 1,129 |
| BKK03850 | *ycnC* | -2,524 | 1,136 | 0,098 | 318 | 1,165 |
| BKK40750 | *yyaQ* | -2,524 | 1,119 | 0,106 | 545 | 1,147 |
| BKK26820 | *yrpD* | -2,527 | 1,147 | 0,111 | 280 | 1,177 |
| BKK39730 | *iolD* | -2,527 | 1,135 | 0,108 | 107 | 1,165 |
| BKK20150 | *yosE* | -2,528 | 1,101 | 0,157 | 129 | 1,129 |
| BKK32250 | *thrC* | -2,529 | 1,119 | 0,096 | 75 | 1,148 |
| BKK28520 | *etfA* | -2,532 | 1,128 | 0,107 | 72 | 1,158 |
| BKK21340 | *yomJ* | -2,533 | 1,126 | 0,096 | 315 | 1,155 |
| BKK37020 | *ywkD* | -2,533 | 1,086 | 0,081 | 34 | 1,114 |
| BKK22900 | *ypfB* | -2,534 | 1,072 | 0,109 | 247 | 1,100 |

| BKK name [1] | gene | screening delta$^2$ (%) | erage width (µ | +/- | nb | ADP |
|---|---|---|---|---|---|---|
| BKK27009 | *yrzP* | -2,535 | 1,124 | 0,106 | 366 | 1,153 |
| BKK10670 | *gerPF* | -2,539 | 1,153 | 0,097 | 128 | 1,183 |
| BKK07790 | *yfkQ* | -2,54 | 1,129 | 0,114 | 366 | 1,159 |
| BKK19560 | *yodD* | -2,541 | 1,101 | 0,129 | 112 | 1,129 |
| BKK24260 | *yqxC* | -2,542 | 1,128 | 0,090 | 113 | 1,158 |
| BKK24470 | *yqhS* | -2,542 | 1,128 | 0,090 | 113 | 1,158 |
| BKK05409 | *ydzQ* | -2,543 | 1,118 | 0,098 | 434 | 1,147 |
| BKK26940 | *yraH* | -2,544 | 1,147 | 0,098 | 433 | 1,177 |
| BKK30739 | *ytzL* | -2,548 | 1,127 | 0,091 | 60 | 1,156 |
| BKK10000 | *yhaH* | -2,551 | 1,133 | 0,101 | 400 | 1,163 |
| BKK05310 | *ydeR* | -2,553 | 1,130 | 0,114 | 429 | 1,160 |
| BKK18898 | *yozW* | -2,555 | 1,109 | 0,093 | 41 | 1,138 |
| BKK21770 | *ilvA* | -2,556 | 0,950 | 0,072 | 150 | 0,975 |
| BKK18550 | *yoaC* | -2,562 | 1,109 | 0,109 | 53 | 1,138 |
| BKK12200 | *yjiA* | -2,563 | 1,159 | 0,083 | 1736 | 1,189 |
| BKK08030 | *dusC* | -2,566 | 1,129 | 0,106 | 451 | 1,159 |
| BKK18560 | *yoaD* | -2,568 | 1,132 | 0,110 | 415 | 1,162 |
| BKK18060 | *yneR* | -2,577 | 1,134 | 0,105 | 133 | 1,164 |
| BKK27860 | *nadC* | -2,578 | 1,121 | 0,115 | 422 | 1,151 |
| BKK26600 | *bltD* | -2,579 | 1,146 | 0,099 | 140 | 1,177 |
| BKK28260 | *leuC* | -2,582 | 1,130 | 0,085 | 298 | 1,160 |
| BKK36310 | *ssbB* | -2,59 | 1,085 | 0,114 | 158 | 1,114 |
| BKK40259 | *yyzG* | -2,594 | 1,120 | 0,120 | 232 | 1,149 |
| BKK16810 | *ymfC* | -2,598 | 1,133 | 0,087 | 86 | 1,164 |
| BKK00580 | *yabN* | -2,603 | 1,133 | 0,093 | 136 | 1,164 |
| BKK18170 | *yngA* | -2,612 | 1,141 | 0,125 | 150 | 1,172 |
| BKK36120 | *ywrB* | -2,614 | 1,136 | 0,096 | 96 | 1,166 |
| BKK28900 | *lrgB* | -2,618 | 1,084 | 0,121 | 239 | 1,113 |
| BKK39050 | *katE* | -2,621 | 1,134 | 0,094 | 87 | 1,165 |
| BKK10770 | *wprA* | -2,626 | 1,152 | 0,118 | 131 | 1,183 |
| BKK29170 | *ytzA* | -2,626 | 1,131 | 0,102 | 350 | 1,161 |
| BKK08200 | *malP* | -2,633 | 1,128 | 0,098 | 405 | 1,159 |
| BKK09910 | *yhaO* | -2,633 | 1,153 | 0,102 | 172 | 1,184 |
| BKK17550 | *ynaG* | -2,633 | 1,133 | 0,151 | 89 | 1,164 |
| BKK20630 | *yoqH* | -2,633 | 1,131 | 0,103 | 434 | 1,162 |
| BKK35900 | *pgsB* | -2,633 | 1,119 | 0,103 | 461 | 1,149 |
| BKK06530 | *purD* | -2,634 | 1,148 | 0,118 | 479 | 1,179 |
| BKK40021 | *yxzK* | -2,636 | 1,119 | 0,089 | 601 | 1,149 |
| BKK21440 | *bdbB* | -2,643 | 1,124 | 0,103 | 548 | 1,155 |
| BKK06290 | *yeaA* | -2,644 | 1,148 | 0,116 | 159 | 1,179 |
| BKK35030 | *yvoA* | -2,648 | 1,135 | 0,113 | 78 | 1,166 |
| BKK01380 | *mapA* | -2,649 | 1,133 | 0,104 | 327 | 1,164 |
| BKK22050 | *bcsA* | -2,651 | 1,071 | 0,103 | 79 | 1,100 |
| BKK25140 | *cshB* | -2,654 | 1,133 | 0,107 | 190 | 1,164 |
| BKK33810 | *opuCC* | -2,656 | 1,119 | 0,101 | 243 | 1,149 |
| BKK21680 | *msrB* | -2,662 | 1,124 | 0,093 | 567 | 1,155 |
| BKK03460 | *hxlA* | -2,666 | 1,100 | 0,096 | 72 | 1,130 |
| BKK06800 | *yezA* | -2,671 | 1,147 | 0,112 | 331 | 1,179 |
| BKK14840 | *ylaN* | -2,671 | 1,148 | 0,089 | 337 | 1,179 |
| BKK10570 | *yhjN* | -2,674 | 1,152 | 0,096 | 86 | 1,183 |
| BKK15130 | *mraZ* | -2,674 | 1,108 | 0,100 | 130 | 1,138 |
| BKK04000 | *ycsA* | -2,676 | 1,134 | 0,109 | 110 | 1,165 |
| BKK26748 | *yrdD* | -2,68 | 1,145 | 0,098 | 379 | 1,177 |
| BKK28210 | *lonB* | -2,683 | 1,129 | 0,093 | 223 | 1,160 |
| BKK22460 | *ypjH* | -2,687 | 1,120 | 0,100 | 398 | 1,151 |
| BKK36890 | *upp* | -2,692 | 1,084 | 0,127 | 283 | 1,114 |
| BKK26360 | *yqaD* | -2,695 | 0,949 | 0,088 | 33 | 0,975 |
| BKK33290 | *fhuC* | -2,696 | 1,135 | 0,092 | 100 | 1,166 |
| BKK13370 | *ykoQ* | -2,697 | 1,135 | 0,109 | 154 | 1,166 |
| BKK02080 | *ybxH* | -2,698 | 1,099 | 0,097 | 41 | 1,130 |
| BKK13960 | *ykwC* | -2,7 | 1,135 | 0,118 | 328 | 1,166 |
| BKK31000 | *yuaI* | -2,7 | 1,125 | 0,095 | 139 | 1,156 |
| BKK11280 | *yjaU* | -2,71 | 1,151 | 0,144 | 93 | 1,183 |
| BKK21950 | *ypzA* | -2,71 | 1,124 | 0,107 | 354 | 1,155 |
| BKK40740 | *yyaR* | -2,711 | 1,152 | 0,101 | 266 | 1,184 |
| BKK36610 | *ywnC* | -2,717 | 1,125 | 0,105 | 107 | 1,157 |
| BKK14980 | *ylbE* | -2,718 | 1,147 | 0,130 | 424 | 1,179 |
| BKK23420 | *spoVAC* | -2,718 | 1,120 | 0,110 | 328 | 1,151 |
| BKK37200 | *ywjD* | -2,721 | 0,948 | 0,072 | 47 | 0,975 |
| BKK24880 | *yqgO* | -2,728 | 1,132 | 0,098 | 228 | 1,164 |
| BKK27970 | *spoIVFB* | -2,733 | 1,128 | 0,102 | 269 | 1,160 |
| BKK03470 | *hxlR* | -2,74 | 1,099 | 0,087 | 84 | 1,130 |
| BKK19639 | *yoyE* | -2,741 | 1,098 | 0,141 | 180 | 1,129 |
| BKK12750 | *xkdV* | -2,743 | 1,134 | 0,127 | 291 | 1,166 |
| BKK30860 | *ytcA* | -2,746 | 1,188 | 0,074 | 183 | 1,222 |
| BKK13570 | *mtnU* | -2,747 | 1,134 | 0,099 | 260 | 1,166 |
| BKK19790 | *cgeB* | -2,751 | 1,098 | 0,091 | 379 | 1,129 |
| BKK30260 | *msmR* | -2,757 | 1,124 | 0,102 | 89 | 1,156 |
| BKK34630 | *mdxR* | -2,757 | 1,129 | 0,091 | 497 | 1,161 |

| BKK name [1] | gene | screening delta$^2$ (%) | erage width (µ | +/- | nb | ADP |
|---|---|---|---|---|---|---|
| BKK17800 | *yndJ* | -2,759 | 1,131 | 0,125 | 81 | 1,164 |
| BKK03569 | *sfp* | -2,765 | 1,099 | 0,093 | 46 | 1,130 |
| BKK15210 | *spoVE* | -2,766 | 1,173 | 0,125 | 72 | 1,206 |
| BKK22280 | *yppD* | -2,774 | 1,070 | 0,114 | 281 | 1,100 |
| BKK29770 | *ytxH* | -2,777 | 1,082 | 0,121 | 83 | 1,113 |
| BKK34770 | *yvcJ* | -2,778 | 1,104 | 0,114 | 215 | 1,135 |
| BKK35730 | *tagE* | -2,778 | 1,134 | 0,104 | 114 | 1,166 |
| BKK15750 | *yloN* | -2,779 | 1,103 | 0,097 | 89 | 1,134 |
| BKK23660 | *yqkB* | -2,783 | 1,154 | 0,103 | 246 | 1,187 |
| BKK25240 | *yqfL* | -2,784 | 1,131 | 0,102 | 140 | 1,164 |
| BKK28380 | *gerM* | -2,788 | 1,127 | 0,117 | 165 | 1,160 |
| BKK00980 | *sigH* | -2,792 | 1,154 | 0,109 | 586 | 1,187 |
| BKK00960 | *rlmB* | -2,8 | 1,154 | 0,093 | 297 | 1,187 |
| BKK24910 | *yqgM* | -2,8 | 1,119 | 0,105 | 522 | 1,151 |
| BKK29830 | *ytpQ* | -2,8 | 1,125 | 0,105 | 157 | 1,158 |
| BKK05109 | *ydzN* | -2,801 | 1,127 | 0,101 | 522 | 1,160 |
| BKK13330 | *ykoL* | -2,802 | 1,108 | 0,114 | 39 | 1,139 |
| BKK23870 | *polYA* | -2,803 | 1,154 | 0,100 | 293 | 1,187 |
| BKK27640 | *yrvC* | -2,807 | 1,127 | 0,093 | 225 | 1,160 |
| BKK00830 | *ctsR* | -2,808 | 1,069 | 0,121 | 202 | 1,100 |
| BKK09890 | *natA* | -2,808 | 1,174 | 0,151 | 90 | 1,208 |
| BKK12020 | *manA* | -2,808 | 1,156 | 0,138 | 122 | 1,189 |
| BKK39370 | *hutI* | -2,808 | 0,948 | 0,072 | 30 | 0,975 |
| BKK36180 | *ywqK* | -2,81 | 1,134 | 0,098 | 114 | 1,166 |
| BKK02600 | *cwlJ* | -2,812 | 1,168 | 0,061 | 37 | 1,201 |
| BKK38370 | *ywbC* | -2,813 | 1,087 | 0,222 | 31 | 1,119 |
| BKK32280 | *yutG* | -2,816 | 1,116 | 0,105 | 62 | 1,148 |
| BKK15640 | *yloA* | -2,818 | 1,116 | 0,165 | 31 | 1,148 |
| BKK27850 | *nadA* | -2,823 | 1,127 | 0,098 | 241 | 1,160 |
| BKK35609 | *tuaA* | -2,823 | 1,117 | 0,109 | 518 | 1,149 |
| BKK14360 | *yknY* | -2,829 | 1,106 | 0,096 | 114 | 1,138 |
| BKK27530 | *yrvN* | -2,83 | 1,127 | 0,093 | 414 | 1,160 |
| BKK32719 | *yuzK* | -2,83 | 1,102 | 0,097 | 169 | 1,134 |
| BKK20490 | *yoqW* | -2,843 | 1,097 | 0,126 | 55 | 1,129 |
| BKK09430 | *citR* | -2,849 | 1,143 | 0,109 | 184 | 1,177 |
| BKK27030 | *sacC* | -2,849 | 1,143 | 0,095 | 115 | 1,177 |
| BKK28180 | *ysxD* | -2,85 | 1,127 | 0,099 | 313 | 1,160 |
| BKK02560 | *ycbM* | -2,851 | 1,098 | 0,099 | 170 | 1,130 |
| BKK07920 | *yfkE* | -2,852 | 1,126 | 0,101 | 420 | 1,159 |
| BKK08630 | *yfhQ* | -2,855 | 1,126 | 0,105 | 319 | 1,159 |
| BKK11290 | *yjaV* | -2,86 | 1,149 | 0,123 | 133 | 1,183 |
| BKK08580 | *yfhL* | -2,862 | 1,126 | 0,095 | 507 | 1,159 |
| BKK40320 | *argI* | -2,862 | 1,118 | 0,105 | 351 | 1,151 |
| BKK20640 | *yoqG* | -2,866 | 1,129 | 0,102 | 411 | 1,162 |
| BKK06510 | *purN* | -2,871 | 1,145 | 0,103 | 417 | 1,179 |
| BKK08090 | *acoL* | -2,873 | 1,129 | 0,106 | 386 | 1,163 |
| BKK25400 | *yqeY* | -2,873 | 1,130 | 0,103 | 178 | 1,164 |
| BKK16845 | *ymfF* | -2,877 | 1,130 | 0,114 | 127 | 1,164 |
| BKK36000 | *alsD* | -2,881 | 1,124 | 0,100 | 167 | 1,157 |
| BKK20340 | *yorL* | -2,89 | 1,097 | 0,144 | 56 | 1,129 |
| BKK31760 | *pncA* | -2,894 | 1,106 | 0,096 | 129 | 1,139 |
| BKK20100 | *yosJ* | -2,898 | 1,097 | 0,133 | 51 | 1,129 |
| BKK00160 | *yaaH* | -2,903 | 1,125 | 0,122 | 113 | 1,159 |
| BKK13590 | *mtnW* | -2,908 | 1,132 | 0,083 | 62 | 1,166 |
| BKK28860 | *rpmI* | -2,908 | 1,118 | 0,090 | 208 | 1,151 |
| BKK36200 | *ywqI* | -2,909 | 1,082 | 0,114 | 70 | 1,114 |
| BKK00660 | *yabT* | -2,912 | 1,096 | 0,161 | 81 | 1,129 |
| BKK37940 | *ywdJ* | -2,916 | 1,086 | 0,115 | 64 | 1,119 |
| BKK40430 | *yycE* | -2,918 | 1,132 | 0,099 | 185 | 1,166 |
| BKK01860 | *ybcF* | -2,921 | 1,126 | 0,100 | 422 | 1,160 |
| BKK07690 | *mapB* | -2,928 | 1,125 | 0,111 | 181 | 1,159 |
| BKK17060 | *ymzD* | -2,929 | 1,129 | 0,099 | 74 | 1,164 |
| BKK21670 | *ypqP* | -2,929 | 1,121 | 0,104 | 256 | 1,155 |
| BKK15170 | *spoVD* | -2,931 | 1,129 | 0,117 | 221 | 1,163 |
| BKK31130 | *yubD* | -2,931 | 1,186 | 0,079 | 95 | 1,222 |
| BKK21100 | *yonG* | -2,932 | 1,117 | 0,114 | 347 | 1,151 |
| BKK27468 | *yrzQ* | -2,932 | 1,126 | 0,093 | 407 | 1,160 |
| BKK21570 | *yokJ* | -2,94 | 1,121 | 0,112 | 417 | 1,155 |
| BKK36020 | *alsR* | -2,95 | 1,081 | 0,116 | 336 | 1,114 |
| BKK26410 | *psiE* | -2,953 | 1,142 | 0,103 | 251 | 1,177 |
| BKK10070 | *yhaA* | -2,954 | 1,142 | 0,100 | 150 | 1,177 |
| BKK30370 | *bceB* | -2,959 | 1,117 | 0,122 | 241 | 1,151 |
| BKK39040 | *yxiS* | -2,961 | 1,130 | 0,109 | 60 | 1,165 |
| BKK21330 | *yomK* | -2,963 | 1,119 | 0,098 | 639 | 1,153 |
| BKK21310 | *yozP* | -2,965 | 1,121 | 0,112 | 299 | 1,155 |
| BKK21910 | *metA* | -2,965 | 1,123 | 0,107 | 137 | 1,158 |
| BKK12610 | *xkdG* | -2,968 | 1,132 | 0,119 | 584 | 1,166 |
| BKK18080 | *yneT* | -2,968 | 1,129 | 0,103 | 100 | 1,164 |
| BKK32469 | *yuzJ* | -2,968 | 1,114 | 0,108 | 58 | 1,148 |

| BKK name [1] | gene | screening delta$^2$ (%) | erage width (µ | +/- | nb | ADP |
|---|---|---|---|---|---|---|
| BKK06083 | *ydzW* | -2,972 | 1,133 | 0,102 | 143 | 1,167 |
| BKK32330 | *lipA* | -2,976 | 1,127 | 0,102 | 309 | 1,161 |
| BKK20270 | *yorS* | -2,98 | 1,149 | 0,106 | 101 | 1,184 |
| BKK37360 | *sboX* | -2,982 | 1,085 | 0,096 | 95 | 1,119 |
| BKK28270 | *leuB* | -2,983 | 1,125 | 0,107 | 318 | 1,160 |
| BKK40770 | *tetB* | -2,985 | 1,149 | 0,108 | 84 | 1,184 |
| BKK08620 | *yfhP* | -2,99 | 1,124 | 0,103 | 526 | 1,159 |
| BKK40220 | *yydB* | -2,99 | 1,131 | 0,098 | 113 | 1,166 |
| BKK31480 | *yuxJ* | -2,992 | 1,114 | 0,129 | 184 | 1,148 |
| BKK31490 | *pbpD* | -2,993 | 1,185 | 0,074 | 103 | 1,222 |
| BKK37530 | *ywhC* | -2,999 | 1,085 | 0,097 | 81 | 1,119 |
| BKK08670 | *ygaB* | -3 | 1,124 | 0,115 | 301 | 1,159 |
| BKK13420 | *ykoW* | -3,003 | 1,172 | 0,147 | 169 | 1,208 |
| BKK34750 | *whiA* | -3,003 | 1,101 | 0,144 | 238 | 1,135 |
| BKK23329 | *ypuC* | -3,016 | 1,116 | 0,097 | 316 | 1,151 |
| BKK33600 | *smpB* | -3,02 | 1,100 | 0,093 | 70 | 1,134 |
| BKK20060 | *nrdEB* | -3,021 | 1,133 | 0,108 | 417 | 1,168 |
| BKK02820 | *rapJ* | -3,026 | 1,096 | 0,099 | 54 | 1,130 |
| BKK01570 | *ybaN* | -3,027 | 1,141 | 0,098 | 283 | 1,177 |
| BKK20950 | *yopB* | -3,03 | 1,120 | 0,111 | 297 | 1,155 |
| BKK15510 | *pyrAA* | -3,032 | 1,121 | 0,094 | 126 | 1,156 |
| BKK29130 | *icd* | -3,037 | 1,079 | 0,108 | 216 | 1,113 |
| BKK03560 | *ycxD* | -3,039 | 1,096 | 0,099 | 61 | 1,130 |
| BKK40920 | *engD* | -3,041 | 1,148 | 0,097 | 110 | 1,184 |
| BKK09680 | *nhaC* | -3,044 | 1,141 | 0,116 | 113 | 1,177 |
| BKK29020 | *gapB* | -3,044 | 1,079 | 0,093 | 142 | 1,113 |
| BKK12880 | *ykcB* | -3,053 | 1,131 | 0,111 | 385 | 1,166 |
| BKK07200 | *yetJ* | -3,058 | 1,143 | 0,100 | 309 | 1,179 |
| BKK06550 | *yecA* | -3,059 | 1,143 | 0,108 | 290 | 1,179 |
| BKK17070 | *ymcC* | -3,059 | 1,128 | 0,120 | 77 | 1,164 |
| BKK14630 | *speA* | -3,064 | 1,143 | 0,111 | 578 | 1,179 |
| BKK11780 | *cotV* | -3,066 | 1,153 | 0,125 | 942 | 1,189 |
| BKK24590 | *yqhG* | -3,067 | 0,945 | 0,091 | 78 | 0,975 |
| BKK19190 | *desK* | -3,071 | 1,169 | 0,108 | 71 | 1,206 |
| BKK23700 | *yqjX* | -3,071 | 1,151 | 0,099 | 168 | 1,187 |
| BKK27890 | *nadR* | -3,074 | 1,124 | 0,100 | 193 | 1,160 |
| BKK25720 | *yqeD* | -3,075 | 1,116 | 0,107 | 389 | 1,151 |
| BKK09690 | *nhaX* | -3,08 | 1,141 | 0,099 | 150 | 1,177 |
| BKK26090 | *yqbJ* | -3,086 | 1,116 | 0,102 | 571 | 1,151 |
| BKK06900 | *cotJB* | -3,088 | 1,142 | 0,116 | 414 | 1,179 |
| BKK28820 | *ysdC* | -3,09 | 1,104 | 0,103 | 125 | 1,139 |
| BKK17400 | *ymaB* | -3,092 | 1,126 | 0,106 | 441 | 1,162 |
| BKK15670 | *ylzA* | -3,11 | 1,099 | 0,073 | 125 | 1,134 |
| BKK27250 | *mccB* | -3,11 | 1,140 | 0,114 | 181 | 1,177 |
| BKK20880 | *yopI* | -3,118 | 0,945 | 0,070 | 69 | 0,975 |
| BKK31690 | *comP* | -3,118 | 1,130 | 0,098 | 91 | 1,166 |
| BKK39790 | *yxcE* | -3,119 | 1,128 | 0,075 | 63 | 1,165 |
| BKK27520 | *cymR* | -3,12 | 1,123 | 0,099 | 388 | 1,160 |
| BKK24370 | *spoIIIAG* | -3,121 | 1,115 | 0,095 | 412 | 1,151 |
| BKK40520 | *yybS* | -3,126 | 1,130 | 0,104 | 157 | 1,166 |
| BKK11690 | *thiG* | -3,13 | 1,131 | 0,106 | 235 | 1,168 |
| BKK09040 | *yhcD* | -3,132 | 1,140 | 0,100 | 215 | 1,177 |
| BKK22110 | *kdgK* | -3,133 | 1,066 | 0,117 | 155 | 1,100 |
| BKK08710 | *gsaB* | -3,15 | 1,122 | 0,108 | 543 | 1,159 |
| BKK35640 | *lytA* | -3,153 | 1,125 | 0,107 | 340 | 1,161 |
| BKK05343 | *ydzS* | -3,154 | 1,104 | 0,100 | 137 | 1,139 |
| BKK36730 | *ywmD* | -3,154 | 1,079 | 0,104 | 233 | 1,114 |
| BKK10540 | *ntdB* | -3,157 | 1,170 | 0,147 | 103 | 1,208 |
| BKK19170 | *yocD* | -3,157 | 1,135 | 0,086 | 143 | 1,172 |
| BKK34360 | *epsB* | -3,157 | 1,113 | 0,096 | 410 | 1,149 |
| BKK01010 | *nusG* | -3,16 | 1,150 | 0,133 | 98 | 1,187 |
| BKK08720 | *ygaF* | -3,163 | 1,122 | 0,108 | 460 | 1,159 |
| BKK24480 | *yqhR* | -3,164 | 1,150 | 0,100 | 225 | 1,187 |
| BKK30850 | *ytdA* | -3,165 | 1,183 | 0,059 | 48 | 1,222 |
| BKK27750 | *bofC* | -3,168 | 1,123 | 0,096 | 254 | 1,160 |
| BKK23500 | *drm* | -3,182 | 1,117 | 0,098 | 469 | 1,153 |
| BKK21140 | *yonC* | -3,183 | 1,118 | 0,103 | 283 | 1,155 |
| BKK40573 | *yyzI* | -3,185 | 1,129 | 0,123 | 86 | 1,166 |
| BKK32310 | *yutD* | -3,19 | 1,124 | 0,097 | 399 | 1,161 |
| BKK25020 | *sodA* | -3,192 | 1,126 | 0,100 | 159 | 1,164 |
| BKK19970 | *yosX* | -3,196 | 1,093 | 0,142 | 297 | 1,129 |
| BKK40910 | *rpsF* | -3,199 | 1,147 | 0,108 | 212 | 1,184 |
| BKK14540 | *ykzG* | -3,2 | 0,944 | 0,093 | 45 | 0,975 |
| BKK22420 | *panC* | -3,206 | 1,121 | 0,112 | 139 | 1,158 |
| BKK25280 | *recO* | -3,206 | 1,126 | 0,110 | 82 | 1,164 |
| BKK06930 | *yesK* | -3,208 | 1,163 | 0,082 | 103 | 1,201 |
| BKK09610 | *yhdV* | -3,211 | 1,139 | 0,098 | 139 | 1,177 |
| BKK12580 | *xtmB* | -3,213 | 1,151 | 0,158 | 645 | 1,189 |
| BKK21350 | *yomI* | -3,215 | 1,118 | 0,106 | 363 | 1,155 |

### Sup. Table 4: Cell width of mutants of the BKK collection (continued)

| BKK name [1] | gene | screening delta$^2$ (%) | erage width (µ | +/- | nb | ADP |
|---|---|---|---|---|---|---|
| BKK09790 | *yheB* | -3,22 | 1,139 | 0,102 | 160 | 1,177 |
| BKK26100 | *yqbI* | -3,22 | 1,139 | 0,092 | 273 | 1,177 |
| BKK31790 | *yueG* | -3,222 | 1,124 | 0,102 | 275 | 1,161 |
| BKK29880 | *malS* | -3,223 | 1,077 | 0,100 | 138 | 1,113 |
| BKK27770 | *yrbE* | -3,227 | 1,122 | 0,092 | 155 | 1,160 |
| BKK14040 | *ykuD* | -3,239 | 1,129 | 0,144 | 63 | 1,166 |
| BKK37660 | *pta* | -3,241 | 1,082 | 0,103 | 43 | 1,119 |
| BKK00300 | *yaaR* | -3,253 | 1,110 | 0,098 | 509 | 1,147 |
| BKK06559 | *yezF* | -3,254 | 1,140 | 0,111 | 323 | 1,179 |
| BKK07010 | *yesS* | -3,259 | 1,140 | 0,121 | 322 | 1,179 |
| BKK14569 | *ykzV* | -3,26 | 1,141 | 0,108 | 517 | 1,179 |
| BKK17340 | *ymaH* | -3,269 | 1,125 | 0,100 | 83 | 1,164 |
| BKK17780 | *yndG* | -3,27 | 1,125 | 0,105 | 111 | 1,164 |
| BKK30610 | *ytlC* | -3,287 | 1,182 | 0,097 | 538 | 1,222 |
| BKK14870 | *ctaA* | -3,294 | 1,140 | 0,100 | 700 | 1,179 |
| BKK05970 | *rex* | -3,303 | 1,119 | 0,082 | 66 | 1,158 |
| BKK10660 | *yisB* | -3,316 | 1,144 | 0,111 | 81 | 1,183 |
| BKK32040 | *yuiF* | -3,321 | 1,110 | 0,106 | 107 | 1,148 |
| BKK11320 | *yjzB* | -3,327 | 1,144 | 0,115 | 105 | 1,183 |
| BKK06650 | *sapB* | -3,332 | 1,140 | 0,101 | 446 | 1,179 |
| BKK26970 | *adhB* | -3,333 | 1,113 | 0,113 | 266 | 1,151 |
| BKK33560 | *yvaD* | -3,335 | 1,127 | 0,112 | 111 | 1,166 |
| BKK32580 | *frlM* | -3,337 | 1,110 | 0,135 | 34 | 1,148 |
| BKK09590 | *yhdT* | -3,344 | 1,138 | 0,080 | 117 | 1,177 |
| BKK05020 | *phrI* | -3,351 | 1,109 | 0,099 | 471 | 1,147 |
| BKK34300 | *epsH* | -3,353 | 1,111 | 0,099 | 194 | 1,149 |
| BKK06870 | *yesE* | -3,354 | 1,139 | 0,095 | 217 | 1,179 |
| BKK21470 | *sunT* | -3,362 | 1,116 | 0,106 | 467 | 1,155 |
| BKK10200 | *yhfE* | -3,37 | 1,119 | 0,098 | 274 | 1,158 |
| BKK20190 | *yosA* | -3,372 | 1,091 | 0,129 | 93 | 1,129 |
| BKK33280 | *yvrP* | -3,372 | 1,127 | 0,100 | 87 | 1,166 |
| BKK13710 | *ykvI* | -3,373 | 1,127 | 0,095 | 412 | 1,166 |
| BKK18979 | *yoyA* | -3,38 | 1,123 | 0,105 | 426 | 1,162 |
| BKK22980 | *ypbG* | -3,381 | 1,114 | 0,101 | 575 | 1,153 |
| BKK39400 | *pdp* | -3,384 | 1,125 | 0,092 | 89 | 1,165 |
| BKK09150 | *yhcN* | -3,386 | 1,137 | 0,103 | 110 | 1,177 |
| BKK10930 | *yitB* | -3,389 | 1,143 | 0,117 | 93 | 1,183 |
| BKK25790 | *arsB* | -3,389 | 0,942 | 0,067 | 55 | 0,975 |
| BKK33330 | *lysP* | -3,395 | 1,096 | 0,103 | 76 | 1,134 |
| BKK17267 | *ymzE* | -3,396 | 1,124 | 0,105 | 209 | 1,164 |
| BKK26500 | *yrkI* | -3,397 | 1,114 | 0,105 | 511 | 1,153 |
| BKK03250 | *ycgR* | -3,398 | 1,092 | 0,106 | 46 | 1,130 |
| BKK21700 | *ypoP* | -3,398 | 1,114 | 0,113 | 383 | 1,153 |
| BKK30900 | *cotS* | -3,401 | 1,117 | 0,109 | 124 | 1,156 |
| BKK28890 | *yscB* | -3,405 | 1,180 | 0,075 | 73 | 1,222 |
| BKK03600 | *tcyB* | -3,418 | 1,160 | 0,074 | 478 | 1,201 |
| BKK11730 | *cotO* | -3,419 | 1,144 | 0,087 | 86 | 1,184 |
| BKK11460 | *oppD* | -3,422 | 1,167 | 0,138 | 139 | 1,208 |
| BKK09450 | *yhdF* | -3,424 | 1,137 | 0,090 | 185 | 1,177 |
| BKK04250 | *lrpC* | -3,436 | 1,125 | 0,106 | 439 | 1,165 |
| BKK11140 | *yitU* | -3,455 | 1,142 | 0,113 | 73 | 1,183 |
| BKK23780 | *yqjQ* | -3,467 | 1,146 | 0,097 | 190 | 1,187 |
| BKK05050 | *lrpA* | -3,473 | 1,127 | 0,144 | 150 | 1,167 |
| BKK36960 | *ywlB* | -3,477 | 1,076 | 0,112 | 120 | 1,114 |
| BKK02590 | *ycbP* | -3,478 | 1,091 | 0,094 | 50 | 1,130 |
| BKK07650 | *yflK* | -3,479 | 0,941 | 0,063 | 173 | 0,975 |
| BKK11960 | *yjcR* | -3,489 | 1,148 | 0,130 | 929 | 1,189 |
| BKK38470 | *ywaD* | -3,499 | 1,080 | 0,109 | 176 | 1,119 |
| BKK32610 | *frlB* | -3,5 | 1,108 | 0,106 | 42 | 1,148 |
| BKK41010 | *trmF* | -3,505 | 1,143 | 0,084 | 169 | 1,184 |
| BKK28350 | *ysnB* | -3,509 | 1,119 | 0,091 | 379 | 1,160 |
| BKK19760 | *cgeD* | -3,515 | 1,090 | 0,119 | 447 | 1,129 |
| BKK25550 | *rpsT* | -3,519 | 1,123 | 0,107 | 179 | 1,164 |
| BKK26030 | *yqbO* | -3,521 | 1,135 | 0,098 | 271 | 1,177 |
| BKK20320 | *yorN* | -3,526 | 1,089 | 0,138 | 92 | 1,129 |
| BKK04670 | *rsbRA* | -3,527 | 1,124 | 0,113 | 335 | 1,165 |
| BKK39800 | *yxcD* | -3,532 | 1,124 | 0,094 | 35 | 1,165 |
| BKK06660 | *opuE* | -3,537 | 1,159 | 0,081 | 125 | 1,201 |
| BKK03110 | *ycgH* | -3,54 | 1,159 | 0,079 | 59 | 1,201 |
| BKK21710 | *dinF* | -3,54 | 1,113 | 0,091 | 278 | 1,153 |
| BKK39430 | *deoR* | -3,547 | 1,123 | 0,097 | 111 | 1,165 |
| BKK17678 | *ynzJ* | -3,551 | 1,122 | 0,114 | 97 | 1,164 |
| BKK35010 | *nagA* | -3,553 | 1,095 | 0,121 | 463 | 1,135 |
| BKK03780 | *phrC* | -3,557 | 1,107 | 0,095 | 545 | 1,147 |
| BKK17970 | *cotM* | -3,572 | 1,163 | 0,126 | 121 | 1,206 |
| BKK26240 | *yqaO* | -3,575 | 1,135 | 0,121 | 168 | 1,177 |
| BKK30400 | *bceR* | -3,578 | 1,115 | 0,092 | 136 | 1,156 |
| BKK00800 | *yazB* | -3,586 | 1,061 | 0,121 | 391 | 1,100 |
| BKK08370 | *yfiR* | -3,589 | 0,940 | 0,072 | 60 | 0,975 |

| BKK name [1] | gene | screening delta$^2$ (%) | erage width (µ | +/- | nb | ADP |
|---|---|---|---|---|---|---|
| BKK36080 | *ywrF* | -3,595 | 1,074 | 0,089 | 41 | 1,114 |
| BKK10410 | *yhzC* | -3,599 | 1,141 | 0,102 | 84 | 1,183 |
| BKK09830 | *yhaX* | -3,6 | 1,135 | 0,110 | 118 | 1,177 |
| BKK07150 | *yetG* | -3,602 | 1,136 | 0,119 | 289 | 1,179 |
| BKK27180 | *yrhH* | -3,605 | 1,112 | 0,102 | 274 | 1,153 |
| BKK00710 | *hslO* | -3,607 | 1,113 | 0,099 | 356 | 1,155 |
| BKK27020 | *yraA* | -3,608 | 1,134 | 0,098 | 379 | 1,177 |
| BKK14480 | *abh* | -3,61 | 1,121 | 0,102 | 535 | 1,163 |
| BKK17260 | *aprX* | -3,611 | 1,122 | 0,098 | 158 | 1,164 |
| BKK07540 | *yfmA* | -3,613 | 1,118 | 0,096 | 627 | 1,160 |
| BKK28840 | *ysdA* | -3,614 | 1,073 | 0,116 | 106 | 1,113 |
| BKK39950 | *yxaJ* | -3,614 | 1,123 | 0,101 | 114 | 1,165 |
| BKK20080 | *yosL* | -3,619 | 0,940 | 0,080 | 38 | 0,975 |
| BKK29249 | *ytzJ* | -3,62 | 1,073 | 0,117 | 108 | 1,113 |
| BKK05280 | *ydeO* | -3,645 | 1,125 | 0,088 | 99 | 1,167 |
| BKK23880 | *yqzJ* | -3,651 | 1,144 | 0,098 | 324 | 1,187 |
| BKK02930 | *yceG* | -3,654 | 1,106 | 0,100 | 616 | 1,147 |
| BKK19749 | *yoyG* | -3,667 | 1,088 | 0,152 | 196 | 1,129 |
| BKK36250 | *ptkA* | -3,668 | 1,107 | 0,105 | 358 | 1,149 |
| BKK19720 | *yodR* | -3,67 | 1,088 | 0,093 | 404 | 1,129 |
| BKK32150 | *paiA* | -3,672 | 1,106 | 0,100 | 272 | 1,148 |
| BKK13850 | *zosA* | -3,676 | 1,162 | 0,141 | 155 | 1,206 |
| BKK37370 | *albA* | -3,678 | 1,078 | 0,095 | 136 | 1,119 |
| BKK33760 | *sdpB* | -3,694 | 1,123 | 0,098 | 97 | 1,166 |
| BKK37930 | *ywdK* | -3,709 | 1,107 | 0,090 | 210 | 1,149 |
| BKK14460 | *ykpC* | -3,721 | 1,135 | 0,136 | 765 | 1,179 |
| BKK23410 | *spoVAD* | -3,724 | 1,143 | 0,096 | 457 | 1,187 |
| BKK34310 | *epsG* | -3,724 | 1,093 | 0,119 | 56 | 1,135 |
| BKK21580 | *yokI* | -3,73 | 1,112 | 0,091 | 566 | 1,155 |
| BKK24320 | *nusB* | -3,732 | 0,939 | 0,070 | 33 | 0,975 |
| BKK37870 | *spsE* | -3,735 | 1,107 | 0,098 | 536 | 1,149 |
| BKK36590 | *clsA* | -3,737 | 1,073 | 0,122 | 124 | 1,114 |
| BKK32270 | *yutH* | -3,738 | 1,105 | 0,106 | 90 | 1,148 |
| BKK23590 | *ansR* | -3,751 | 1,143 | 0,095 | 227 | 1,187 |
| BKK12660 | *xkdM* | -3,759 | 1,144 | 0,132 | 612 | 1,189 |
| BKK38280 | *efeU* | -3,772 | 1,077 | 0,104 | 58 | 1,119 |
| BKK04750 | *ydcF* | -3,778 | 1,121 | 0,136 | 564 | 1,165 |
| BKK09700 | *yheJ* | -3,779 | 1,133 | 0,104 | 105 | 1,177 |
| BKK32880 | *yusP* | -3,779 | 1,176 | 0,100 | 100 | 1,222 |
| BKK09940 | *yhaL* | -3,79 | 1,132 | 0,097 | 159 | 1,177 |
| BKK00860 | *clpC* | -3,799 | 1,058 | 0,117 | 107 | 1,100 |
| BKK18700 | *yoaQ* | -3,803 | 1,124 | 0,106 | 271 | 1,168 |
| BKK26960 | *yraF* | -3,807 | 1,132 | 0,108 | 202 | 1,177 |
| BKK11250 | *argF* | -3,819 | 1,138 | 0,104 | 86 | 1,183 |
| BKK40450 | *yycD* | -3,82 | 1,122 | 0,096 | 163 | 1,166 |
| BKK02810 | *cwlK* | -3,825 | 0,938 | 0,082 | 36 | 0,975 |
| BKK36520 | *glnK* | -3,83 | 1,072 | 0,132 | 124 | 1,114 |
| BKK10650 | *sbcC* | -3,832 | 1,138 | 0,096 | 97 | 1,183 |
| BKK15310 | *spoIIGA* | -3,832 | 1,160 | 0,124 | 93 | 1,206 |
| BKK21480 | *sunA* | -3,836 | 1,111 | 0,108 | 268 | 1,155 |
| BKK18899 | *yozX* | -3,839 | 1,123 | 0,110 | 308 | 1,168 |
| BKK27170 | *bscR* | -3,839 | 1,132 | 0,102 | 83 | 1,177 |
| BKK15550 | *pyrF* | -3,84 | 1,096 | 0,098 | 60 | 1,139 |
| BKK14660 | *ykzI* | -3,854 | 1,134 | 0,129 | 292 | 1,179 |
| BKK00090 | *guaB* | -3,863 | 1,133 | 0,135 | 122 | 1,179 |
| BKK35180 | *csbA* | -3,866 | 1,091 | 0,106 | 540 | 1,135 |
| BKK37540 | *ywhB* | -3,866 | 1,075 | 0,114 | 68 | 1,119 |
| BKK21600 | *yokG* | -3,871 | 1,110 | 0,108 | 369 | 1,155 |
| BKK28830 | *ysdB* | -3,876 | 1,070 | 0,094 | 74 | 1,113 |
| BKK19840 | *yotL* | -3,883 | 1,085 | 0,134 | 281 | 1,129 |
| BKK02030 | *ybdM* | -3,888 | 1,086 | 0,090 | 31 | 1,130 |
| BKK03420 | *nin* | -3,897 | 1,086 | 0,104 | 116 | 1,130 |
| BKK23300 | *ypuD* | -3,899 | 1,141 | 0,100 | 84 | 1,187 |
| BKK15740 | *rsmB* | -3,92 | 1,090 | 0,099 | 114 | 1,134 |
| BKK14250 | *yknT* | -3,924 | 1,133 | 0,114 | 285 | 1,179 |
| BKK11055 | *yitO* | -3,936 | 1,137 | 0,110 | 90 | 1,183 |
| BKK35550 | *tuaG* | -3,947 | 1,104 | 0,093 | 553 | 1,149 |
| BKK13440 | *ykoY* | -3,951 | 1,117 | 0,107 | 228 | 1,163 |
| BKK37630 | *ywfM* | -3,953 | 1,104 | 0,104 | 290 | 1,149 |
| BKK21830 | *ypjQ* | -3,954 | 1,112 | 0,097 | 125 | 1,158 |
| BKK26200 | *yqaS* | -3,957 | 1,130 | 0,107 | 234 | 1,177 |
| BKK13320 | *ykzB* | -3,975 | 1,120 | 0,092 | 92 | 1,166 |
| BKK13789 | *ykzQ* | -3,975 | 1,159 | 0,133 | 121 | 1,206 |
| BKK24990 | *pstS* | -3,978 | 1,158 | 0,156 | 108 | 1,206 |
| BKK24360 | *spoIIIAH* | -3,997 | 1,158 | 0,136 | 37 | 1,206 |
| BKK16340 | *fliZ* | -4,004 | 1,115 | 0,095 | 163 | 1,162 |
| BKK14071 | *fadG* | -4,011 | 1,116 | 0,099 | 391 | 1,163 |
| BKK40760 | *yyaP* | -4,018 | 1,137 | 0,098 | 178 | 1,184 |
| BKK04790 | *ydcK* | -4,026 | 1,118 | 0,146 | 53 | 1,165 |

| BKK name [1] | gene | screening delta$^2$ (%) | erage width (µ | +/- | nb | ADP |
|---|---|---|---|---|---|---|
| BKK03010 | *amhX* | -4,043 | 1,084 | 0,074 | 32 | 1,130 |
| BKK17559 | *ynzI* | -4,045 | 1,115 | 0,093 | 526 | 1,162 |
| BKK35620 | *lytC* | -4,049 | 1,114 | 0,103 | 339 | 1,161 |
| BKK13540 | *ogt* | -4,073 | 1,119 | 0,105 | 204 | 1,166 |
| BKK04760 | *ydcG* | -4,077 | 1,117 | 0,152 | 168 | 1,165 |
| BKK04950 | *yddF* | -4,078 | 1,120 | 0,120 | 192 | 1,167 |
| BKK25170 | *yqfO* | -4,091 | 1,116 | 0,097 | 313 | 1,164 |
| BKK16610 | *ylxR* | -4,097 | 1,120 | 0,116 | 327 | 1,168 |
| BKK34490 | *yvdS* | -4,098 | 1,089 | 0,113 | 297 | 1,135 |
| BKK37120 | *fbaA* | -4,113 | 1,073 | 0,088 | 41 | 1,119 |
| BKK10400 | *yhxC* | -4,116 | 1,135 | 0,095 | 95 | 1,183 |
| BKK26850 | *yrpG* | -4,125 | 1,128 | 0,100 | 173 | 1,177 |
| BKK15360 | *ylmC* | -4,127 | 1,130 | 0,156 | 58 | 1,179 |
| BKK36360 | *mscL* | -4,129 | 1,113 | 0,099 | 373 | 1,161 |
| BKK04430 | *ydbD* | -4,132 | 1,112 | 0,098 | 537 | 1,160 |
| BKK09030 | *yhcC* | -4,143 | 1,128 | 0,101 | 105 | 1,177 |
| BKK20560 | *yoqO* | -4,143 | 1,083 | 0,119 | 63 | 1,129 |
| BKK00360 | *yabC* | -4,151 | 1,130 | 0,121 | 206 | 1,179 |
| BKK27360 | *yrrM* | -4,152 | 1,111 | 0,095 | 162 | 1,160 |
| BKK11310 | *comZ* | -4,157 | 1,134 | 0,126 | 120 | 1,183 |
| BKK20240 | *yorV* | -4,158 | 1,082 | 0,152 | 32 | 1,129 |
| BKK14020 | *ykyB* | -4,169 | 1,118 | 0,115 | 263 | 1,166 |
| BKK36970 | *spoIIR* | -4,174 | 1,101 | 0,097 | 368 | 1,149 |
| BKK28420 | *ysmA* | -4,186 | 0,934 | 0,072 | 45 | 0,975 |
| BKK10220 | *gltT* | -4,205 | 1,127 | 0,108 | 206 | 1,177 |
| BKK19260 | *yocM* | -4,211 | 1,122 | 0,115 | 39 | 1,172 |
| BKK10290 | *yhfN* | -4,215 | 1,133 | 0,099 | 135 | 1,183 |
| BKK24390 | *spoIIIAE* | -4,219 | 1,156 | 0,152 | 36 | 1,206 |
| BKK04330 | *mutT* | -4,22 | 1,116 | 0,137 | 405 | 1,165 |
| BKK14400 | *fruA* | -4,227 | 1,129 | 0,124 | 701 | 1,179 |
| BKK19910 | *yotE* | -4,23 | 1,082 | 0,126 | 111 | 1,129 |
| BKK15010 | *ylbH* | -4,237 | 1,129 | 0,093 | 295 | 1,179 |
| BKK37050 | *maeA* | -4,238 | 1,067 | 0,145 | 60 | 1,114 |
| BKK25410 | *rpsU* | -4,24 | 1,114 | 0,102 | 277 | 1,164 |
| BKK37600 | *ywfO* | -4,243 | 1,071 | 0,116 | 355 | 1,119 |
| BKK34810 | *yvcD* | -4,246 | 1,087 | 0,120 | 38 | 1,135 |
| BKK05830 | *gmuC* | -4,253 | 1,118 | 0,098 | 150 | 1,167 |
| BKK35319 | *yvzG* | -4,258 | 1,117 | 0,104 | 141 | 1,166 |
| BKK16710 | *mlpA* | -4,265 | 1,114 | 0,105 | 137 | 1,164 |
| BKK09300 | *glpD* | -4,267 | 1,127 | 0,096 | 306 | 1,177 |
| BKK25710 | *cwlH* | -4,267 | 1,114 | 0,108 | 127 | 1,164 |
| BKK18210 | *yngE* | -4,273 | 1,122 | 0,083 | 93 | 1,172 |
| BKK01560 | *kbaA* | -4,282 | 1,126 | 0,100 | 278 | 1,177 |
| BKK09290 | *glpK* | -4,282 | 1,127 | 0,095 | 239 | 1,177 |
| BKK20430 | *yorC* | -4,284 | 1,081 | 0,134 | 168 | 1,129 |
| BKK16560 | *rseP* | -4,286 | 1,112 | 0,095 | 344 | 1,161 |
| BKK20550 | *yoqP* | -4,289 | 1,081 | 0,113 | 131 | 1,129 |
| BKK23920 | *yqjD* | -4,299 | 1,136 | 0,094 | 539 | 1,187 |
| BKK18819 | *yozU* | -4,31 | 1,121 | 0,121 | 242 | 1,172 |
| BKK31720 | *degQ* | -4,329 | 1,111 | 0,098 | 262 | 1,161 |
| BKK16320 | *fliY* | -4,343 | 1,117 | 0,102 | 233 | 1,168 |
| BKK05960 | *moaC* | -4,346 | 1,116 | 0,143 | 429 | 1,167 |
| BKK22370 | *aspB* | -4,347 | 1,052 | 0,113 | 246 | 1,100 |
| BKK09410 | *phoA* | -4,354 | 1,126 | 0,096 | 124 | 1,177 |
| BKK38750 | *cydB* | -4,356 | 1,115 | 0,093 | 133 | 1,166 |
| BKK17560 | *ynaI* | -4,358 | 1,113 | 0,098 | 106 | 1,164 |
| BKK20660 | *yoqE* | -4,36 | 1,111 | 0,108 | 325 | 1,162 |
| BKK34960 | *yvoF* | -4,365 | 1,086 | 0,132 | 229 | 1,135 |
| BKK05099 | *ydzM* | -4,366 | 1,116 | 0,108 | 265 | 1,167 |
| BKK38700 | *sigY* | -4,369 | 1,114 | 0,126 | 108 | 1,165 |
| BKK12430 | *rapA* | -4,37 | 1,137 | 0,122 | 337 | 1,189 |
| BKK37880 | *spsD* | -4,37 | 1,070 | 0,088 | 48 | 1,119 |
| BKK16400 | *flhF* | -4,375 | 1,117 | 0,097 | 396 | 1,168 |
| BKK24130 | *prpD* | -4,381 | 1,135 | 0,104 | 362 | 1,187 |
| BKK23690 | *yqjY* | -4,394 | 1,135 | 0,091 | 466 | 1,187 |
| BKK14050 | *ykuE* | -4,402 | 1,115 | 0,087 | 38 | 1,166 |
| BKK11920 | *yjcN* | -4,404 | 1,137 | 0,112 | 461 | 1,189 |
| BKK11990 | *yjdB* | -4,405 | 0,932 | 0,069 | 59 | 0,975 |
| BKK22240 | *ypqA* | -4,438 | 1,100 | 0,108 | 384 | 1,151 |
| BKK14800 | *ylaJ* | -4,439 | 1,153 | 0,128 | 77 | 1,206 |
| BKK30290 | *amyC* | -4,441 | 1,168 | 0,100 | 39 | 1,222 |
| BKK29030 | *ytcD* | -4,444 | 1,089 | 0,087 | 120 | 1,139 |
| BKK32430 | *pucJ* | -4,444 | 1,097 | 0,125 | 117 | 1,148 |
| BKK04300 | *ydaM* | -4,447 | 1,113 | 0,116 | 664 | 1,165 |
| BKK18370 | *yoeA* | -4,451 | 1,153 | 0,128 | 99 | 1,206 |
| BKK22470 | *ypjG* | -4,455 | 1,051 | 0,124 | 189 | 1,100 |
| BKK36710 | *fdhD* | -4,455 | 1,065 | 0,100 | 67 | 1,114 |
| BKK36260 | *ywqC* | -4,467 | 1,098 | 0,099 | 537 | 1,149 |
| BKK05010 | *rapI* | -4,478 | 1,096 | 0,098 | 520 | 1,147 |

| BKK name [1] | gene | screening delta$^2$ (%) | erage width (µ | +/- | nb | ADP |
|---|---|---|---|---|---|---|
| BKK25310 | *dgkA* | -4,491 | 1,111 | 0,099 | 473 | 1,164 |
| BKK11430 | *oppA* | -4,499 | 1,131 | 0,091 | 184 | 1,184 |
| BKK36030 | *ywrK* | -4,499 | 1,109 | 0,094 | 476 | 1,161 |
| BKK11030 | *yitL* | -4,506 | 1,130 | 0,105 | 101 | 1,183 |
| BKK01910 | *skfA* | -4,507 | 1,063 | 0,112 | 125 | 1,113 |
| BKK21500 | *uvrX* | -4,507 | 1,099 | 0,087 | 250 | 1,151 |
| BKK36190 | *ywqJ* | -4,507 | 1,064 | 0,100 | 195 | 1,114 |
| BKK34540 | *clpP* | -4,517 | 0,931 | 0,054 | 36 | 0,975 |
| BKK26470 | *yrkL* | -4,523 | 1,123 | 0,100 | 311 | 1,177 |
| BKK38210 | *ywcD* | -4,527 | 1,113 | 0,104 | 138 | 1,166 |
| BKK36470 | *pucI* | -4,534 | 1,064 | 0,108 | 62 | 1,114 |
| BKK33790 | *sdpR* | -4,55 | 1,113 | 0,091 | 226 | 1,166 |
| BKK05860 | *gmuE* | -4,552 | 1,114 | 0,088 | 305 | 1,167 |
| BKK10850 | *yisS* | -4,558 | 1,129 | 0,109 | 73 | 1,183 |
| BKK25970 | *yqcA* | -4,558 | 0,931 | 0,068 | 64 | 0,975 |
| BKK34700 | *yvcR* | -4,564 | 1,097 | 0,098 | 406 | 1,149 |
| BKK09270 | *glpP* | -4,568 | 1,123 | 0,107 | 161 | 1,177 |
| BKK01030 | *rplA* | -4,57 | 1,110 | 0,095 | 223 | 1,164 |
| BKK23450 | *sigF* | -4,571 | 1,133 | 0,090 | 301 | 1,187 |
| BKK27630 | *yrvD* | -4,583 | 1,106 | 0,096 | 377 | 1,160 |
| BKK06950 | *yesM* | -4,585 | 1,146 | 0,096 | 151 | 1,201 |
| BKK29470 | *ackA* | -4,595 | 1,062 | 0,133 | 35 | 1,113 |
| BKK02100 | *cypC* | -4,597 | 1,078 | 0,117 | 37 | 1,130 |
| BKK03370 | *yckA* | -4,601 | 1,078 | 0,094 | 106 | 1,130 |
| BKK35380 | *fliW* | -4,612 | 1,083 | 0,113 | 531 | 1,135 |
| BKK23770 | *dsdA* | -4,62 | 1,132 | 0,095 | 216 | 1,187 |
| BKK19310 | *dhaS* | -4,642 | 1,117 | 0,092 | 136 | 1,172 |
| BKK40670 | *yybE* | -4,648 | 1,129 | 0,094 | 248 | 1,184 |
| BKK16150 | *clpQ* | -4,649 | 1,106 | 0,101 | 470 | 1,160 |
| BKK05220 | *ydeJ* | -4,679 | 1,113 | 0,145 | 91 | 1,167 |
| BKK38190 | *galT* | -4,685 | 1,066 | 0,107 | 79 | 1,119 |
| BKK36330 | *ywpF* | -4,692 | 1,107 | 0,101 | 437 | 1,161 |
| BKK19130 | *yocA* | -4,707 | 1,116 | 0,105 | 99 | 1,172 |
| BKK23480 | *dacF* | -4,71 | 1,150 | 0,165 | 58 | 1,206 |
| BKK29120 | *mdh* | -4,727 | 1,060 | 0,088 | 105 | 1,113 |
| BKK12410 | *yjoA* | -4,729 | 1,133 | 0,143 | 638 | 1,189 |
| BKK32260 | *hom* | -4,73 | 1,086 | 0,079 | 69 | 1,139 |
| BKK11150 | *yitV* | -4,739 | 1,127 | 0,105 | 113 | 1,183 |
| BKK34670 | *yvdA* | -4,739 | 1,081 | 0,111 | 207 | 1,135 |
| BKK04530 | *ydbN* | -4,74 | 1,110 | 0,132 | 519 | 1,165 |
| BKK34640 | *yvdD* | -4,745 | 1,081 | 0,141 | 94 | 1,135 |
| BKK24120 | *prpB* | -4,757 | 1,131 | 0,098 | 252 | 1,187 |
| BKK34890 | *hisH* | -4,766 | 1,081 | 0,123 | 256 | 1,135 |
| BKK00650 | *yabS* | -4,774 | 1,075 | 0,096 | 189 | 1,129 |
| BKK09180 | *yhcQ* | -4,784 | 1,121 | 0,096 | 245 | 1,177 |
| BKK41000 | *rsmG* | -4,785 | 1,128 | 0,101 | 132 | 1,184 |
| BKK37820 | *spsK* | -4,787 | 1,094 | 0,095 | 637 | 1,149 |
| BKK37950 | *ywdI* | -4,791 | 1,065 | 0,104 | 135 | 1,119 |
| BKK12690 | *xkdP* | -4,799 | 1,112 | 0,109 | 284 | 1,168 |
| BKK14620 | *slp* | -4,824 | 1,148 | 0,099 | 30 | 1,206 |
| BKK23490 | *pupG* | -4,83 | 1,130 | 0,095 | 606 | 1,187 |
| BKK00200 | *yaaK* | -4,842 | 1,120 | 0,098 | 112 | 1,177 |
| BKK37890 | *spsC* | -4,844 | 1,094 | 0,109 | 420 | 1,149 |
| BKK07240 | *yetN* | -4,848 | 1,122 | 0,106 | 358 | 1,179 |
| BKK12749 | *xkzA* | -4,85 | 1,110 | 0,111 | 304 | 1,166 |
| BKK10050 | *ecsB* | -4,852 | 1,120 | 0,097 | 133 | 1,177 |
| BKK35610 | *tuaA* | -4,853 | 1,094 | 0,109 | 551 | 1,149 |
| BKK08810 | *senS* | -4,855 | 1,103 | 0,093 | 320 | 1,159 |
| BKK09920 | *yhaN* | -4,864 | 1,120 | 0,104 | 87 | 1,177 |
| BKK13910 | *ptsI* | -4,878 | 1,109 | 0,141 | 37 | 1,166 |
| BKK37100 | *murAB* | -4,883 | 1,093 | 0,107 | 138 | 1,149 |
| BKK10820 | *yisQ* | -4,889 | 1,082 | 0,099 | 56 | 1,138 |
| BKK28000 | *minC* | -4,891 | 1,103 | 0,092 | 148 | 1,160 |
| BKK32850 | *fadM* | -4,908 | 1,079 | 0,092 | 71 | 1,134 |
| BKK03990 | *mtlD* | -4,91 | 1,108 | 0,108 | 181 | 1,165 |
| BKK36668 | *ywzE* | -4,915 | 1,060 | 0,103 | 164 | 1,114 |
| BKK12160 | *yjgC* | -4,947 | 1,130 | 0,138 | 416 | 1,189 |
| BKK33540 | *yvaB* | -4,95 | 0,927 | 0,077 | 67 | 0,975 |
| BKK20140 | *yosF* | -4,958 | 1,073 | 0,118 | 34 | 1,129 |
| BKK19660 | *yozD* | -4,959 | 1,073 | 0,138 | 455 | 1,129 |
| BKK32730 | *metN* | -4,964 | 1,161 | 0,114 | 157 | 1,222 |
| BKK11839 | *yjzE* | -4,985 | 1,130 | 0,103 | 859 | 1,189 |
| BKK04240 | *ydzA* | -4,989 | 1,107 | 0,105 | 284 | 1,165 |
| BKK09800 | *yheA* | -4,993 | 1,118 | 0,103 | 137 | 1,177 |
| BKK18910 | *rapK* | -5,013 | 1,113 | 0,103 | 123 | 1,172 |
| BKK19210 | *yocH* | -5,019 | 1,146 | 0,139 | 59 | 1,206 |
| BKK22200 | *cotD* | -5,019 | 1,045 | 0,142 | 141 | 1,100 |
| BKK12860 | *ykbA* | -5,021 | 1,108 | 0,126 | 173 | 1,166 |
| BKK33020 | *cssS* | -5,025 | 1,108 | 0,098 | 126 | 1,166 |

### Sup. Table 4: Cell width of mutants of the BKK collection (continued)

| BKK name [1] | gene | screening delta[2] (%) | erage width (µ | +/- | nb | ADP |
|---|---|---|---|---|---|---|
| BKK35490 | *degU* | -5,028 | 1,103 | 0,094 | 353 | 1,161 |
| BKK14220 | *ykuU* | -5,047 | 1,120 | 0,137 | 196 | 1,179 |
| BKK19770 | *cgeC* | -5,049 | 1,072 | 0,116 | 409 | 1,129 |
| BKK00640 | *spoIIE* | -5,063 | 1,072 | 0,165 | 177 | 1,129 |
| BKK32230 | *yuxL* | -5,063 | 1,082 | 0,082 | 98 | 1,139 |
| BKK05210 | *ydeI* | -5,069 | 1,108 | 0,096 | 109 | 1,167 |
| BKK28370 | *rph* | -5,069 | 1,099 | 0,101 | 155 | 1,158 |
| BKK39720 | *iolE* | -5,075 | 1,106 | 0,118 | 77 | 1,165 |
| BKK21850 | *ypiP* | -5,086 | 1,096 | 0,100 | 509 | 1,155 |
| BKK13270 | *ykoI* | -5,104 | 1,107 | 0,112 | 287 | 1,166 |
| BKK04080 | *kipI* | -5,122 | 1,089 | 0,094 | 445 | 1,147 |
| BKK07110 | *lplB* | -5,123 | 1,118 | 0,096 | 317 | 1,179 |
| BKK04360 | *mntH* | -5,141 | 1,100 | 0,112 | 447 | 1,160 |
| BKK29090 | *polA* | -5,16 | 1,056 | 0,129 | 60 | 1,113 |
| BKK37250 | *narI* | -5,167 | 1,061 | 0,096 | 52 | 1,119 |
| BKK34265 | *epsK* | -5,179 | 1,076 | 0,097 | 424 | 1,135 |
| BKK23760 | *coaA* | -5,185 | 1,126 | 0,092 | 230 | 1,187 |
| BKK05900 | *thiL* | -5,19 | 1,107 | 0,121 | 370 | 1,167 |
| BKK05980 | *tatAY* | -5,201 | 1,088 | 0,096 | 277 | 1,147 |
| BKK36210 | *ywqH* | -5,201 | 1,056 | 0,105 | 299 | 1,114 |
| BKK36290 | *ywpJ* | -5,219 | 1,056 | 0,121 | 169 | 1,114 |
| BKK38290 | *thiE* | -5,222 | 1,060 | 0,102 | 69 | 1,119 |
| BKK13350 | *ykoN* | -5,223 | 1,105 | 0,103 | 305 | 1,166 |
| BKK06100 | *ydiS* | -5,237 | 1,106 | 0,112 | 395 | 1,167 |
| BKK00740 | *pabB* | -5,238 | 1,043 | 0,145 | 55 | 1,100 |
| BKK14320 | *yknU* | -5,243 | 1,143 | 0,143 | 136 | 1,206 |
| BKK28130 | *hemB* | -5,281 | 1,098 | 0,126 | 56 | 1,160 |
| BKK37080 | *rho* | -5,283 | 1,060 | 0,114 | 150 | 1,119 |
| BKK24040 | *bkdAB* | -5,286 | 1,124 | 0,096 | 57 | 1,187 |
| BKK16970 | *ymdB* | -5,293 | 1,102 | 0,099 | 95 | 1,164 |
| BKK36780 | *ywzB* | -5,303 | 1,055 | 0,123 | 105 | 1,114 |
| BKK38690 | *yxlC* | -5,32 | 1,138 | 0,098 | 184 | 1,201 |
| BKK32130 | *guaC* | -5,331 | 1,087 | 0,117 | 176 | 1,148 |
| BKK18600 | *yozQ* | -5,334 | 1,077 | 0,106 | 38 | 1,138 |
| BKK31945 | *yukJ* | -5,337 | 1,079 | 0,103 | 69 | 1,139 |
| BKK37470 | *phrF* | -5,346 | 1,088 | 0,098 | 284 | 1,149 |
| BKK28730 | *araQ* | -5,352 | 1,078 | 0,106 | 33 | 1,139 |
| BKK00850 | *mcsB* | -5,355 | 1,041 | 0,117 | 384 | 1,100 |
| BKK03080 | *ycgE* | -5,36 | 1,069 | 0,115 | 43 | 1,130 |
| BKK13890 | *ptsG* | -5,372 | 1,125 | 0,113 | 244 | 1,189 |
| BKK36500 | *ywoB* | -5,372 | 1,054 | 0,112 | 54 | 1,114 |
| BKK22690 | *aroH* | -5,383 | 1,041 | 0,110 | 175 | 1,100 |
| BKK36750 | *spoIID* | -5,389 | 1,054 | 0,112 | 82 | 1,114 |
| BKK12120 | *yjfB* | -5,402 | 1,125 | 0,150 | 170 | 1,189 |
| BKK35850 | *ywtE* | -5,418 | 1,054 | 0,109 | 232 | 1,114 |
| BKK32750 | *metQ* | -5,42 | 1,156 | 0,076 | 124 | 1,222 |
| BKK34500 | *yvdR* | -5,445 | 1,103 | 0,089 | 116 | 1,166 |
| BKK31770 | *yueI* | -5,453 | 1,085 | 0,111 | 190 | 1,148 |
| BKK24810 | *yqgV* | -5,456 | 1,120 | 0,085 | 254 | 1,184 |
| BKK38460 | *tyrZ* | -5,457 | 1,058 | 0,120 | 209 | 1,119 |
| BKK13360 | *ykoP* | -5,467 | 1,103 | 0,103 | 133 | 1,166 |
| BKK06090 | *ydiR* | -5,478 | 1,103 | 0,116 | 63 | 1,167 |
| BKK16900 | *ymfK* | -5,503 | 1,104 | 0,095 | 484 | 1,168 |
| BKK31120 | *lytG* | -5,51 | 1,155 | 0,114 | 35 | 1,222 |
| BKK24770 | *mgsR* | -5,511 | 1,099 | 0,120 | 148 | 1,164 |
| BKK14940 | *ylbA* | -5,527 | 1,114 | 0,085 | 267 | 1,179 |
| BKK20370 | *yorI* | -5,528 | 1,067 | 0,130 | 64 | 1,129 |
| BKK11520 | *mecA* | -5,533 | 1,118 | 0,103 | 81 | 1,183 |
| BKK19950 | *sspC* | -5,539 | 1,067 | 0,150 | 182 | 1,129 |
| BKK03760 | *yclK* | -5,543 | 1,135 | 0,108 | 308 | 1,201 |
| BKK21730 | *ypmS* | -5,56 | 1,091 | 0,100 | 321 | 1,155 |
| BKK18790 | *yoaZ* | -5,564 | 1,106 | 0,126 | 34 | 1,172 |
| BKK23040 | *fer* | -5,575 | 1,039 | 0,127 | 159 | 1,100 |
| BKK12540 | *xkdD* | -5,596 | 1,122 | 0,146 | 699 | 1,189 |
| BKK36480 | *ywoD* | -5,598 | 1,052 | 0,104 | 113 | 1,114 |
| BKK35000 | *hprK* | -5,65 | 1,071 | 0,123 | 123 | 1,135 |
| BKK35870 | *pgsE* | -5,661 | 1,051 | 0,132 | 83 | 1,114 |
| BKK40510 | *yybT* | -5,677 | 1,100 | 0,103 | 128 | 1,166 |
| BKK11040 | *yitM* | -5,698 | 1,116 | 0,103 | 82 | 1,183 |
| BKK19579 | *yoyD* | -5,717 | 1,065 | 0,104 | 462 | 1,129 |
| BKK13080 | *ykkB* | -5,729 | 1,100 | 0,122 | 341 | 1,166 |
| BKK19730 | *yodS* | -5,735 | 1,065 | 0,085 | 361 | 1,129 |
| BKK13740 | *queE* | -5,743 | 1,099 | 0,117 | 417 | 1,166 |
| BKK09930 | *yhaM* | -5,753 | 1,109 | 0,088 | 144 | 1,177 |
| BKK05930 | *rimI* | -5,775 | 1,100 | 0,114 | 469 | 1,167 |
| BKK15340 | *ylmA* | -5,795 | 1,137 | 0,138 | 72 | 1,206 |
| BKK00150 | *dgk* | -5,807 | 1,110 | 0,110 | 188 | 1,179 |
| BKK18650 | *pelB* | -5,827 | 1,103 | 0,123 | 37 | 1,172 |
| BKK37860 | *spsF* | -5,842 | 1,053 | 0,133 | 51 | 1,119 |
| BKK37570 | *mmr* | -5,904 | 1,082 | 0,096 | 313 | 1,149 |
| BKK12370 | *exuR* | -5,917 | 1,119 | 0,124 | 591 | 1,189 |
| BKK36770 | *ywmB* | -5,935 | 1,048 | 0,115 | 118 | 1,114 |
| BKK36680 | *ywmF* | -6,051 | 1,047 | 0,104 | 38 | 1,114 |
| BKK09050 | *yhcE* | -6,061 | 1,106 | 0,098 | 147 | 1,177 |
| BKK15420 | *divIVA* | -6,061 | 0,916 | 0,082 | 59 | 0,975 |
| BKK14610 | *pdhD* | -6,065 | 1,108 | 0,135 | 50 | 1,179 |
| BKK37690 | *bacF* | -6,105 | 1,050 | 0,091 | 51 | 1,119 |
| BKK15580 | *cysP* | -6,125 | 1,078 | 0,113 | 180 | 1,148 |
| BKK21840 | *ypjP* | -6,132 | 1,084 | 0,092 | 301 | 1,155 |
| BKK22230 | *ypqE* | -6,15 | 1,033 | 0,094 | 68 | 1,100 |
| BKK28440 | *sdhA* | -6,151 | 0,915 | 0,055 | 47 | 0,975 |
| BKK35830 | *ywtG* | -6,225 | 1,078 | 0,098 | 291 | 1,149 |
| BKK40620 | *yybJ* | -6,234 | 1,094 | 0,142 | 88 | 1,167 |
| BKK28770 | *araL* | -6,238 | 1,044 | 0,104 | 85 | 1,113 |
| BKK22320 | *ponA* | -6,259 | 1,081 | 0,128 | 394 | 1,153 |
| BKK14629 | *ykzW* | -6,274 | 1,105 | 0,123 | 561 | 1,179 |
| BKK18709 | *yozT* | -6,295 | 1,095 | 0,097 | 377 | 1,168 |
| BKK20510 | *yoqU* | -6,361 | 1,057 | 0,110 | 64 | 1,129 |
| BKK18909 | *yozZ* | -6,405 | 1,097 | 0,128 | 169 | 1,172 |
| BKK19820 | *yotN* | -6,465 | 1,056 | 0,100 | 822 | 1,129 |
| BKK14950 | *ylbB* | -6,466 | 1,103 | 0,092 | 122 | 1,179 |
| BKK34980 | *yvoD* | -6,499 | 1,061 | 0,109 | 241 | 1,135 |
| BKK00470 | *purR* | -6,679 | 1,086 | 0,103 | 97 | 1,164 |
| BKK34900 | *hisB* | -6,697 | 1,059 | 0,132 | 224 | 1,135 |
| BKK12330 | *yjmD* | -6,79 | 1,108 | 0,132 | 661 | 1,189 |
| BKK18480 | *proH* | -6,817 | 1,092 | 0,113 | 162 | 1,172 |
| BKK38780 | *yxkI* | -6,825 | 1,085 | 0,104 | 133 | 1,165 |
| BKK19280 | *yocN* | -6,839 | 1,091 | 0,131 | 344 | 1,172 |
| BKK36300 | *glcR* | -6,844 | 1,038 | 0,122 | 159 | 1,114 |
| BKK29550 | *ytcJ* | -6,86 | 1,037 | 0,127 | 55 | 1,113 |
| BKK22510 | *ypjC* | -6,867 | 1,025 | 0,117 | 53 | 1,100 |
| BKK36900 | *glyA* | -6,868 | 0,908 | 0,093 | 50 | 0,975 |
| BKK20520 | *yoqT* | -6,872 | 1,052 | 0,130 | 198 | 1,129 |
| BKK22520 | *ypjB* | -6,873 | 1,025 | 0,127 | 101 | 1,100 |
| BKK13900 | *ptsH* | -6,914 | 1,107 | 0,116 | 41 | 1,189 |
| BKK04710 | *rsbV* | -6,939 | 1,084 | 0,114 | 273 | 1,165 |
| BKK23130 | *resC* | -6,943 | 1,024 | 0,116 | 207 | 1,100 |
| BKK16070 | *ylqG* | -6,983 | 1,023 | 0,148 | 134 | 1,100 |
| BKK13210 | *thiX* | -7,014 | 1,085 | 0,142 | 40 | 1,166 |
| BKK22820 | *yphE* | -7,032 | 1,023 | 0,103 | 46 | 1,100 |
| BKK04510 | *ydbL* | -7,047 | 1,083 | 0,126 | 663 | 1,165 |
| BKK20290 | *yorQ* | -7,086 | 1,049 | 0,132 | 74 | 1,129 |
| BKK35370 | *csrA* | -7,091 | 1,055 | 0,099 | 132 | 1,135 |
| BKK06110 | *ydjA* | -7,15 | 1,084 | 0,102 | 369 | 1,167 |
| BKK33360 | *yvgJ* | -7,2 | 1,134 | 0,104 | 215 | 1,222 |
| BKK29640 | *ytsP* | -7,208 | 1,033 | 0,094 | 123 | 1,113 |
| BKK28760 | *araM* | -7,221 | 1,033 | 0,100 | 80 | 1,113 |
| BKK36460 | *ywoF* | -7,232 | 1,034 | 0,090 | 30 | 1,114 |
| BKK20270 | *yorS* | -7,257 | 1,047 | 0,136 | 40 | 1,129 |
| BKK28120 | *hemL* | -7,261 | 1,075 | 0,093 | 185 | 1,160 |
| BKK22390 | *ypmA* | -7,278 | 1,020 | 0,133 | 164 | 1,100 |
| BKK37440 | *ywhL* | -7,291 | 1,037 | 0,104 | 101 | 1,119 |
| BKK01920 | *skfB* | -7,317 | 1,032 | 0,098 | 110 | 1,113 |
| BKK29350 | *tcyM* | -7,331 | 1,055 | 0,089 | 48 | 1,138 |
| BKK10980 | *yitG* | -7,353 | 1,119 | 0,116 | 43 | 1,208 |
| BKK05680 | *ydgK* | -7,423 | 1,112 | 0,082 | 220 | 1,201 |
| BKK20260 | *yorT* | -7,443 | 1,045 | 0,130 | 38 | 1,129 |
| BKK29440 | *argH* | -7,459 | 1,030 | 0,111 | 30 | 1,113 |
| BKK36430 | *usd* | -7,477 | 1,031 | 0,092 | 84 | 1,114 |
| BKK11770 | *cotW* | -7,497 | 1,100 | 0,099 | 279 | 1,189 |
| BKK37510 | *pbpG* | -7,549 | 1,063 | 0,096 | 228 | 1,149 |
| BKK18730 | *yoaS* | -7,576 | 1,083 | 0,120 | 71 | 1,172 |
| BKK14230 | *ykuV* | -7,642 | 1,089 | 0,123 | 156 | 1,179 |
| BKK00870 | *radA* | -7,706 | 1,015 | 0,122 | 45 | 1,100 |
| BKK14920 | *ctaF* | -7,782 | 1,087 | 0,085 | 361 | 1,179 |
| BKK37640 | *ywfL* | -7,8 | 0,899 | 0,073 | 40 | 0,975 |
| BKK40490 | *yycA* | -7,876 | 1,073 | 0,118 | 255 | 1,165 |
| BKK15600 | *cysC* | -7,935 | 1,057 | 0,118 | 187 | 1,148 |
| BKK32740 | *metP* | -7,952 | 1,074 | 0,096 | 103 | 1,166 |
| BKK35270 | *cccB* | -8,013 | 1,044 | 0,129 | 273 | 1,135 |
| BKK19290 | *yozO* | -8,249 | 1,075 | 0,139 | 549 | 1,172 |
| BKK18760 | *yoaU* | -8,275 | 1,075 | 0,115 | 81 | 1,172 |
| BKK11720 | *fabI* | -8,301 | 1,090 | 0,127 | 90 | 1,189 |
| BKK26710 | *azlC* | -8,397 | 1,054 | 0,122 | 257 | 1,151 |
| BKK05850 | *gmuR* | -8,408 | 1,069 | 0,090 | 154 | 1,167 |
| BKK22480 | *mgsA* | -8,445 | 1,007 | 0,132 | 95 | 1,100 |
| BKK36690 | *rapB* | -8,446 | 1,020 | 0,110 | 88 | 1,114 |
| BKK10140 | *hemY* | -8,489 | 1,077 | 0,097 | 162 | 1,177 |
| BKK38430 | *gspA* | -8,529 | 1,023 | 0,108 | 128 | 1,119 |
| BKK36910 | *ywlG* | -8,643 | 1,018 | 0,105 | 199 | 1,114 |
| BKK00630 | *yabR* | -8,798 | 1,056 | 0,101 | 68 | 1,158 |
| BKK28170 | *hemA* | -8,839 | 1,057 | 0,124 | 323 | 1,160 |
| **BKK29180** | *pyk* | -8,872 | 1,062 | 0,119 | 123 | 1,156 |
| **BKK15790** | *rpe* | -8,905 | 1,034 | 0,115 | 35 | 1,135 |
| **BKK22210** | *yprB* | -8,945 | 1,002 | 0,134 | 110 | 1,100 |
| **BKK13900** | *ptsH* | -8,969 | 1,062 | 0,126 | 58 | 1,166 |
| **BKK19320** | *sqhC* | -9,157 | 1,064 | 0,134 | 55 | 1,172 |
| **BKK34620** | *mdxD* | -9,39 | 1,028 | 0,120 | 61 | 1,135 |
| **BKK03480** | *srfAA* | -9,413 | 1,024 | 0,103 | 38 | 1,130 |
| **BKK20300** | *yorP* | -9,42 | 1,023 | 0,145 | 52 | 1,129 |
| **BKK01889** | *ybzH* | -9,47 | 0,883 | 0,054 | 42 | 0,975 |
| **BKK22340** | *nth* | -9,547 | 0,995 | 0,104 | 85 | 1,100 |
| **BKK19020** | *yobN* | -9,585 | 1,059 | 0,107 | 36 | 1,172 |
| **BKK38660** | *yxlF* | -9,702 | 1,028 | 0,068 | 45 | 1,138 |
| **BKK06360** | *guaA* | -9,94 | 0,878 | 0,055 | 50 | 0,975 |
| **BKK22220** | *yprA* | -10,181 | 0,988 | 0,116 | 38 | 1,100 |
| **BKK22410** | *panD* | -10,738 | 0,982 | 0,183 | 31 | 1,100 |
| **BKK19180** | *des* | -10,806 | 1,045 | 0,088 | 87 | 1,172 |
| **BKK04310** | *ydaN* | -10,896 | 1,038 | 0,123 | 218 | 1,165 |
| **BKK08680** | *ygaC* | -10,965 | 1,032 | 0,117 | 185 | 1,159 |
| **BKK22360** | *asnS* | -12,67 | 0,961 | 0,148 | 37 | 1,100 |
| **BKK40390** | *walH* | -13,873 | 0,973 | 0,069 | 43 | 1,130 |
| BKK00110 | *pdxS* | n/a | n/a | n/a | n/a | n/a |
| BKK00690 | *ftsH* | n/a | n/a | n/a | n/a | n/a |
| BKK01670 | *ybbE* | n/a | n/a | n/a | n/a | n/a |
| BKK27260 | *mccA* | n/a | n/a | n/a | n/a | n/a |
| BKK31290 | *yugT* | n/a | n/a | n/a | n/a | n/a |
| BKK33920 | *tpiA* | n/a | n/a | n/a | n/a | n/a |
| BKK33930 | *pgk* | n/a | n/a | n/a | n/a | n/a |
| BKK38150 | *qoxC* | n/a | n/a | n/a | n/a | n/a |
| BKK41030 | *jag* | n/a | n/a | n/a | n/a | n/a |

1: underscored and bold names indicate the top 10 % and 1% delta, respectively

2: n/a : 9 mutants have no values because they did not grow after several attempts

### Sup. Table 5: Width of the 0.5% largest and thinnest selected strains

| *gene* | | screening step | | | | | checking step | | | | post-backcross step | | | | | backcross strains without $Mg^{2+}$ [5] | | | | | |
|---|---|---|---|---|---|---|---|---|---|---|---|---|---|---|---|---|---|---|---|---|---|
| name | reference | width (µm) | +/- | delta[1] (%) | nb | ADP (µm) | width (µm) | +/- | delta[2] (%) | nb | width (µm) | +/- | delta[3] (%) | nb | *P* -value[4] | width (µm) | +/- | delta (%) | nb | P value[4] | |
| *cwlO* | BKK34800 | 1,42 | 0,14 | 23,36 | 301 | 1,15 | 1,25 | 0,13 | 28,30 | 265 | 1,09 | 0,09 | 13,07 | 438 | *** | 1,10 | 0,08 | 7,68 | 411 | **** | comfirmed positively affected |
| *rodZ* | BKK16910 | 1,29 | 0,11 | 11,40 | 287 | 1,16 | 1,12 | 0,10 | 14,77 | 199 | 1,08 | 0,08 | 12,55 | 331 | **** | 1,07 | 0,09 | 5,52 | 370 | ** | |
| *rpe* | BKK15790 | 1,03 | 0,12 | -8,91 | 35 | 1,14 | 0,86 | 0,07 | -11,39 | 136 | 1,07 | 0,18 | 11,02 | 207 | **** | 1,09 | 0,11 | 6,96 | 357 | **** | |
| *ftsX* | BKK35250 | 1,33 | 0,18 | 15,59 | 286 | 1,15 | 1,18 | 0,12 | 20,97 | 167 | 1,06 | 0,08 | 9,95 | 342 | **** | 1,10 | 0,08 | 7,66 | 341 | **** | |
| *yaaA* | BKK00030 | 1,28 | 0,12 | 9,94 | 189 | 1,17 | 1,02 | 0,07 | 4,87 | 188 | 1,06 | 0,09 | 9,72 | 349 | *** | 1,13 | 0,09 | 11,38 | 294 | **** | |
| *ftsE* | BKK35260 | 1,38 | 0,15 | 19,51 | 109 | 1,16 | 1,21 | 0,11 | 24,20 | 90 | 1,05 | 0,08 | 9,43 | 431 | **** | 1,07 | 0,08 | 5,03 | 335 | ** | |
| *dacA* | BKK00100 | 1,34 | 0,15 | 15,94 | 107 | 1,16 | 1,03 | 0,10 | 5,94 | 260 | 1,05 | 0,08 | 9,33 | 409 | **** | 1,08 | 0,09 | 6,06 | 370 | **** | |
| *yuaC* | BKK31070 | 1,29 | 0,13 | 11,33 | 238 | 1,16 | 0,99 | 0,08 | 2,04 | 150 | 1,02 | 0,08 | 5,48 | 405 | | 1,01 | 0,07 | -0,88 | 281 | | |
| *natA* | BKK02750 | 1,31 | 0,12 | 9,10 | 127 | 1,20 | 0,96 | 0,06 | -0,86 | 129 | 1,00 | 0,07 | 4,03 | 331 | | 1,03 | 0,09 | 1,63 | 90 | | |
| *ymfD* | BKK16825 | 1,31 | 0,17 | 12,67 | 82 | 1,16 | 0,93 | 0,07 | -4,42 | 53 | 1,00 | 0,07 | 3,92 | 383 | | 1,04 | 0,07 | 1,79 | 102 | | |
| *comFC* | BKK35450 | 1,24 | 0,10 | 11,47 | 90 | 1,11 | 1,03 | 0,06 | 6,07 | 238 | 1,00 | 0,07 | 3,64 | 351 | | 1,06 | 0,07 | 3,60 | 97 | | |
| *yvyF* | BKK35440 | 1,22 | 0,11 | 9,58 | 161 | 1,11 | 0,99 | 0,07 | 2,07 | 131 | 1,00 | 0,07 | 3,30 | 367 | | 1,02 | 0,08 | -0,10 | 324 | | |
| *greA* | BKK27320 | 1,10 | 0,10 | 12,65 | 40 | 0,98 | n/a | n/a | n/a | n/a | 0,99 | 0,06 | 3,20 | 154 | | 0,99 | 0,06 | -2,32 | 154 | | |
| *mdxD* | BKK34620 | 1,03 | 0,12 | -9,39 | 61 | 1,14 | 0,95 | 0,07 | -2,70 | 174 | 0,99 | 0,07 | 3,08 | 405 | | 0,99 | 0,07 | -2,56 | 295 | | |
| *yprB* | BKK22210 | 1,00 | 0,13 | -8,95 | 110 | 1,10 | 0,97 | 0,09 | -0,22 | 127 | 0,99 | 0,08 | 3,06 | 180 | | 0,99 | 0,07 | -3,06 | 246 | | |
| *srfAA* | BKK03480 | 1,02 | 0,10 | -9,41 | 38 | 1,13 | 0,92 | 0,08 | -4,89 | 123 | 0,99 | 0,07 | 3,06 | 441 | | 1,00 | 0,08 | -1,46 | 287 | | |
| *yobN* | BKK19020 | 1,06 | 0,11 | -9,59 | 36 | 1,17 | 0,98 | 0,09 | 1,31 | 124 | 0,99 | 0,08 | 3,03 | 324 | | 0,97 | 0,07 | -4,43 | 305 | | |
| *yxlF* | BKK38660 | 1,03 | 0,07 | -9,70 | 45 | 1,14 | 0,97 | 0,06 | -0,12 | 127 | 0,99 | 0,07 | 2,83 | 394 | | 1,00 | 0,08 | -1,91 | 324 | | |
| *ypzH* | BKK22849 | 1,28 | 0,13 | 15,99 | 401 | 1,10 | 1,02 | 0,08 | 4,54 | 78 | 0,99 | 0,08 | 2,73 | 356 | | 0,99 | 0,07 | -2,57 | 116 | | |
| *ykhA* | BKK13030 | 1,27 | 0,18 | 9,17 | 109 | 1,17 | 0,99 | 0,06 | 1,31 | 155 | 0,99 | 0,09 | 2,71 | 355 | | 1,04 | 0,10 | 1,93 | 138 | | |
| *yoqC* [6] | BKK20680 | 1,32 | 0,18 | 13,23 | 411 | 1,17 | 0,94 | 0,07 | -3,46 | 101 | n/a | n/a | n/a | n/a | | n/a | n/a | n/a | n/a | | |
| *yorP* [6] | BKK20300 | 1,02 | 0,15 | -9,42 | 52 | 1,13 | 1,01 | 0,07 | 3,90 | 189 | n/a | n/a | n/a | n/a | | n/a | n/a | n/a | n/a | | |
| *des* | BKK19180 | 1,04 | 0,09 | -10,81 | 87 | 1,17 | 0,98 | 0,07 | 0,72 | 165 | 0,99 | 0,09 | 2,47 | 424 | | 1,02 | 0,07 | 0,29 | 204 | | |
| *yprA* | BKK22220 | 0,99 | 0,12 | -10,18 | 38 | 1,10 | 0,99 | 0,08 | 2,14 | 107 | 0,99 | 0,07 | 2,46 | 276 | | 0,99 | 0,07 | -3,10 | 183 | | |
| *xkdX* | BKK12770 | 1,29 | 0,12 | 10,49 | 135 | 1,17 | 0,96 | 0,07 | -0,86 | 224 | 0,98 | 0,07 | 2,04 | 283 | | 1,02 | 0,08 | -0,30 | 155 | | |
| *yqiW* | BKK23990 | 1,26 | 0,14 | 9,50 | 142 | 1,15 | 0,97 | 0,07 | -0,31 | 237 | 0,98 | 0,07 | 1,61 | 324 | | 0,97 | 0,08 | -4,31 | 131 | | |
| *nth* | BKK22340 | 1,00 | 0,10 | -9,55 | 85 | 1,10 | 1,00 | 0,08 | 2,44 | 160 | 0,98 | 0,07 | 1,33 | 267 | | 0,99 | 0,08 | -2,52 | 277 | | |
| *ygaC* | BKK08680 | 1,03 | 0,12 | -10,97 | 185 | 1,16 | 0,94 | 0,07 | -3,16 | 121 | 0,98 | 0,07 | 1,24 | 366 | | 1,01 | 0,08 | -0,55 | 236 | | |
| *xkdW* | BKK12760 | 1,42 | 0,12 | 21,84 | 150 | 1,17 | 0,99 | 0,08 | 2,14 | 79 | 0,97 | 0,07 | 1,14 | 316 | | 0,98 | 0,06 | -4,25 | 64 | | |
| *sqhC* | BKK19320 | 1,06 | 0,13 | -9,16 | 55 | 1,17 | 0,97 | 0,07 | -0,06 | 163 | 0,97 | 0,08 | 1,07 | 322 | | 0,98 | 0,07 | -3,88 | 270 | | |
| *ydaN* | BKK04310 | 1,04 | 0,12 | -10,90 | 218 | 1,16 | 0,97 | 0,07 | -0,04 | 119 | 0,97 | 0,06 | 0,74 | 376 | | 1,01 | 0,08 | -0,87 | 173 | | |
| *kbl* | BKK17000 | 1,28 | 0,16 | 9,83 | 152 | 1,16 | 0,96 | 0,07 | -1,70 | 161 | 0,97 | 0,07 | 0,62 | 333 | | 0,99 | 0,06 | -2,71 | 75 | | |
| *asnS* | BKK22360 | 0,96 | 0,15 | -12,67 | 37 | 1,10 | 0,97 | 0,07 | -0,24 | 103 | 0,97 | 0,07 | 0,17 | 322 | | 0,98 | 0,08 | -3,34 | 100 | | |
| *minJ* | BKK35220 | 1,24 | 0,13 | 9,38 | 104 | 1,14 | 0,98 | 0,08 | 0,49 | 211 | 0,95 | 0,09 | -1,02 | 677 | | 0,97 | 0,11 | -4,60 | 225 | | |
| *walH* | BKK40390 | 0,97 | 0,07 | -13,87 | 43 | 1,13 | 0,89 | 0,06 | -8,04 | 126 | 0,92 | 0,07 | -4,54 | 366 | | 0,92 | 0,07 | -9,55 | 361 | | |
| *pyk* | BKK29180 | 1,06 | 0,12 | -8,87 | 123 | 1,16 | 0,82 | 0,04 | -15,67 | 105 | 0,88 | 0,06 | -8,32 | 314 | *** | 0,94 | 0,08 | -7,47 | 289 | **** | comfirmed negatively |
| *ybzH* | BKK01889 | 0,88 | 0,05 | -9,47 | 42 | 0,98 | n/a | n/a | n/a | n/a | 0,87 | 0,06 | -9,30 | 369 | **** | 0,86 | 0,06 | -15,19 | 115 | **** | |
| *panD* | BKK22410 | 0,98 | 0,18 | -10,74 | 31 | 1,10 | 0,82 | 0,05 | -15,65 | 135 | 0,85 | 0,06 | -11,39 | 301 | **** | 0,87 | 0,06 | -15,04 | 256 | **** | |
| *guaA* | BKK06360 | 0,88 | 0,05 | -9,94 | 50 | 0,98 | n/a | n/a | n/a | n/a | 0,84 | 0,05 | -12,73 | 351 | **** | 0,81 | 0,05 | -20,58 | 182 | **** | |
| *ptsH* | BKK13900 | 1,06 | 0,13 | -8,97 | 58 | 1,17 | 0,83 | 0,06 | -14,44 | 163 | 0,84 | 0,07 | -13,30 | 392 | **** | 0,91 | 0,06 | -10,27 | 390 | **** | |

1: δ relative to ADP
2: δ relative to wild type width
3: δ relative to wild type strain, average of 3 independant replicates
4: *P* -value (summary) of nested t-tests, comparing the widths of mutants with that of the wild type cells (**** = P<0.0001; *** = 0.0001<P<0.001; ** = 0.001<P<0.01; * = 0.01<P<0.05; ns = P>0.05)
5: width values for confirmed affected mutants are the average of 3 independent replicates
6: PCR checking revealed that the BKK strains are wt for the tested loci. Deletions were therefore not backcrossed into the 168 strain

Sup. Table 6. *B. subtilis* strains used in this study

| Strain (*B. subtilis* ) | Relevant genotype | Source or reference[1] |
|---|---|---|
| 168 | (wt) | Laboratory stock |
| PY79 | (wt) | Laboratory stock |
| RCL413 | Ω*neo3427* - Δ*mreB* | Billaudeau, 2019 |
| PS2062 | Δ*ponA* ::*spc* | Popham, 1995 |
| CcBs351 | Δ*rodZ* ::*cat* | Gibson assembly → 168 |
| CcBs628 | Δ*rodZ* ::*cat* | CcBs351→ PY79 |
| BKK00030 | Δ*yaaA* ::*km* | Koo, 2017 |
| BKK00100 | Δ*dacA* ::*km* | Koo, 2017 |
| BKK01889 | Δ*ybzH* ::*km* | Koo, 2017 |
| BKK02750 | Δ*natA* ::*km* | Koo, 2017 |
| BKK03480 | Δ*srfAA* ::*km* | Koo, 2017 |
| BKK04310 | Δ*ydaN* ::*km* | Koo, 2017 |
| BKK06360 | Δ*guaA* ::*km* | Koo, 2017 |
| BKK08680 | Δ*ygaC* ::*km* | Koo, 2017 |
| BKK12760 | Δ*xkdW* ::*km* | Koo, 2017 |
| BKK12770 | Δ*xkdX* ::*km* | Koo, 2017 |
| BKK13030 | Δ*ykhA* ::*km* | Koo, 2017 |
| BKK13900 | Δ*ptsH* ::*km* | Koo, 2017 |
| BKK15790 | Δ*rpe* ::*km* | Koo, 2017 |
| BKK16825 | Δ*ymfD* ::*km* | Koo, 2017 |
| BKK16910 | Δ*rodZ* ::*km* | Koo, 2017 |
| BKK17000 | Δ*kbl* ::*km* | Koo, 2017 |
| BKK19020 | Δ*yobN* ::*km* | Koo, 2017 |
| BKK19180 | Δ*des* ::*km* | Koo, 2017 |
| BKK19320 | Δ*sqhC* ::*km* | Koo, 2017 |
| BKK20300 | Δ*yorP* ::*km* | Koo, 2017 |
| BKK20680 | Δ*yoqC* ::*km* | Koo, 2017 |
| BKK22210 | Δ*yprB* ::*km* | Koo, 2017 |
| BKK22220 | Δ*yprA* ::*km* | Koo, 2017 |
| BKK22340 | Δ*nth* ::*km* | Koo, 2017 |
| BKK22360 | Δ*asnS* ::*km* | Koo, 2017 |
| BKK22410 | Δ*panD* ::*km* | Koo, 2017 |
| BKK22849 | Δ*ypzH* ::*km* | Koo, 2017 |
| BKK23990 | Δ*yqiw* ::*km* | Koo, 2017 |
| BKK27320 | Δ*greA* ::*km* | Koo, 2017 |
| BKK29180 | Δ*pyk* ::*km* | Koo, 2017 |
| BKK31070 | Δ*yuaC* ::*km* | Koo, 2017 |
| BKK34620 | Δ*mdxD* ::*km* | Koo, 2017 |
| BKK34800 | Δ*cwlo* ::*km* | Koo, 2017 |
| BKK35220 | Δ*minJ* ::*km* | Koo, 2017 |
| BKK35250 | Δ*ftsX* ::*km* | Koo, 2017 |
| BKK35260 | Δ*ftsE* ::*km* | Koo, 2017 |
| BKK35440 | Δ*yvyF* ::*km* | Koo, 2017 |
| BKK35450 | Δ*comFC* ::*km* | Koo, 2017 |
| BKK38660 | Δ*yxlF* ::*km* | Koo, 2017 |
| BKK40390 | Δ*walH* ::*km* | Koo, 2017 |
| RCL0820 | Δ*cwlo* ::*km* | BKK34800 DNA → 168 |
| RCL0821 | Δ*xkdW* ::*km* | BKK12760 DNA → 168 |
| RCL0822 | Δ*ftsE* ::*km* | BKK35260 DNA → 168 |
| RCL0823 | Δ*ypzH* ::*km* | BKK22849 DNA → 168 |
| RCL0824 | Δ*dacA* ::*km* | BKK00100 DNA → 168 |
| RCL0825 | Δ*ftsX* ::*km* | BKK35250 DNA → 168 |
| RCL0826 | Δ*ymfD* ::*km* | BKK16825 DNA → 168 |
| RCL0827 | Δ*comFC* ::*km* | BKK35450 DNA → 168 |
| RCL0828 | Δ*rodZ* ::*km* | BKK16910 DNA → 168 |
| RCL0829 | Δ*yuaC* ::*km* | BKK31070 DNA → 168 |
| RCL0830 | Δ*xkdX* ::*km* | BKK12770 DNA → 168 |
| RCL0831 | Δ*yaaA* ::*km* | BKK00030 DNA → 168 |
| RCL0832 | Δ*kbl* ::*km* | BKK17000 DNA → 168 |
| RCL0833 | Δ*yvyF* ::*km* | BKK35440 DNA → 168 |

| | | |
|---|---|---|
| RCL0834 | Δ*minJ* ::*km* | BKK35220 DNA → 168 |
| RCL0835 | Δ*yqiw* ::*km* | BKK23990 DNA → 168 |
| RCL0837 | Δ*ykhA* ::*km* | BKK13030 DNA → 168 |
| RCL0838 | Δ*natA* ::*km* | BKK02750 DNA → 168 |
| RCL0839 | Δ*ywqA* ::*km* | BKK36280 DNA → 168 |
| RCL0840 | Δ*walH* ::*km* | BKK40390 DNA → 168 |
| RCL0841 | Δ*asnS* ::*km* | BKK22360 DNA → 168 |
| RCL0842 | Δ*ygaC* ::*km* | BKK08680 DNA → 168 |
| RCL0843 | Δ*ydaN* ::*km* | BKK04310 DNA → 168 |
| RCL0844 | Δ*des* ::*km* | BKK19180 DNA → 168 |
| RCL0845 | Δ*panD* ::*km* | BKK22410 DNA → 168 |
| RCL0846 | Δ*yprA* ::*km* | BKK22220 DNA → 168 |
| RCL0848 | Δ*yobN* ::*km* | BKK19020 DNA → 168 |
| RCL0849 | Δ*nth* ::*km* | BKK22340 DNA → 168 |
| RCL0850 | Δ*srfAA* ::*km* | BKK03480 DNA → 168 |
| RCL0851 | Δ*mdxD* ::*km* | BKK34620 DNA → 168 |
| RCL0852 | Δ*sqhC* ::*km* | BKK19320 DNA → 168 |
| RCL0853 | Δ*pyk* ::*km* | BKK29180 DNA → 168 |
| RCL0854 | Δ*ptsH* ::*km* | BKK13900 DNA → 168 |
| RCL0855 | Δ*yprB* ::*km* | BKK22210 DNA → 168 |
| RCL0856 | Δ*rpe* ::*km* | BKK15790 DNA → 168 |
| RCL0858 | Δ*greA* ::*km* | BKK27320 DNA → 168 |
| RCL0859 | Δ*ybzH* ::*km* | BKK01889 DNA → 168 |
| RCL0860 | Δ*guaA* ::*km* | BKK06360 DNA → 168 |

1: Arrows indicate construction by transformation with chromosomal DNA
